# Proposal of two new genera and seventy-seven new species of ascomycetous yeasts isolated from China

**DOI:** 10.1101/2025.06.10.658801

**Authors:** H.-H. Zhu, A.-H. Li, M.-M. Liu, Y.-L. Jiang, X.-M. Zhao, S. Pan, J.-H. Liu, X.-H. Zhang, J.-C. Wang, Z.-Y. Tian, G.-S. Wang, Y.-T. Guo, D. Phurbu, F.-Y. Bai, Q.-M. Wang

## Abstract

Ascomycetous yeasts belong to *Saccharomycotina* and *Taphrinomycotina,* with about 1,200 described species distributed worldwide. More than 300 ascomycetous yeast species have been reported from China, which indicates that China possesses a hotspot with high yeast diversity. However, our knowledge of yeast diversity and distribution remains limited, and this area needs to be explored in depth. Here we describe 77 new ascomycetous yeast species with 195 isolates obtained over the past two decades during the continuous yeast diversity surveys in China, based on the phylogenetic analyses of the sequences of D1/D2 domains of the large subunit (LSU) and the internal transcribed spacer region including the 5.8S rDNA (ITS) of the nuclear ribosomal DNA (rDNA). The average nucleotide identity (ANI) analysis was used to delimitate closely related species with low sequence heterogeneity in the ITS and D1/D2 regions. Seventy-five of these new species belong to 35 genera in the *Saccharaomycotina*, while the other two are distributed in *Taphrina* of the *Taphrinomycotina*. In addition, we propose two new genera, namely *Gaboromyces gen. nov.* and *Yurkovozyma gen. nov*.

## INTRODUCTION

Yeasts are defined as a kind of fungi whose asexual reproduction is predominantly in a manner of budding or fission, and most of them form sexual states without a fruiting body (Kurtzman *et al*. 2011a). The members of yeasts belong to *Agaricomycotina*, *Pucciniomycotina* and *Ustilaginomycotina* (*Basidiomycota*, basidiomycetous yeasts) and *Saccharomycotina* and *Taphrinomycotina* (*Ascomycota*, ascomycetous yeasts). A few ascomycetous yeasts are classified into *Neolectomycetes*, *Pneumocystidomycetes*, *Schizosaccharomycetes* and *Taphrinomycetes* (*Taphrinomycotina*), and most of them belong to *Saccharomycetales* (*Saccharomycetes*, *Saccharomycotina*) (Kurtzman 2011). Recently, Groenewald *et al*. (2023) reclassified this order and divided it into seven classes and 12 orders based on a genome-scale analysis.

More than 2,000 yeast species, including about 1,200 ascomycetous yeasts, have been described, and more than half of them have been reported over the past 20 years (Boekhout *et al*. 2022). Among those yeasts, about 353 species were described as new species isolated from China (Li *et al*. 2020, 2022, Boekhout *et al*. 2022, Jiang *et al*. 2024). Only one-quarter of these 353 novelties (about 88 species) belong to ascomycetous yeasts; the others are basidiomycetous yeasts, of which 200 were reported mainly in three papers (Li *et al*. 2020, 2022, Jiang *et al*. 2024). The imbalance between the numbers of newly described ascomycetous and basidiomycetous yeasts from China may be caused by the bias of sampling, isolating, and describing methods. For example, the published yeast species by our team were mostly isolated from the leaves (i.e., Li *et al*. 2020, 2022, Jiang *et al*. 2024) using the ballistoconidia-fall method (Nakase & Takashima 1993), which is a biased method for basidiomycetous yeasts isolation, especially for ballistoconidia yeasts. In order to present a clear outline of the yeast diversity in China, we describe in this study 77 new ascomycetous yeast species that had been isolated from more than 9 various substrates over the past decades based on the nucleotide sequence analyses of the internal transcribed spacer including the 5.8S rDNA (ITS) and the D1/D2 domains of the large subunit (LSU) of rDNA, and the genomic metric analysis, i.e., the average nucleotide identity (ANI).

## MATERIALS AND METHODS

### Strains and phenotypic characterization

The strains studied are listed in Table 1. Strains were isolated from a variety of substrates (including bark, rotting wood, soil, flowers, fruits, and insects) collected from China (Tibet, Yunnan, Beijing, Hunan, Hainan and other provinces).

**Table 1.**
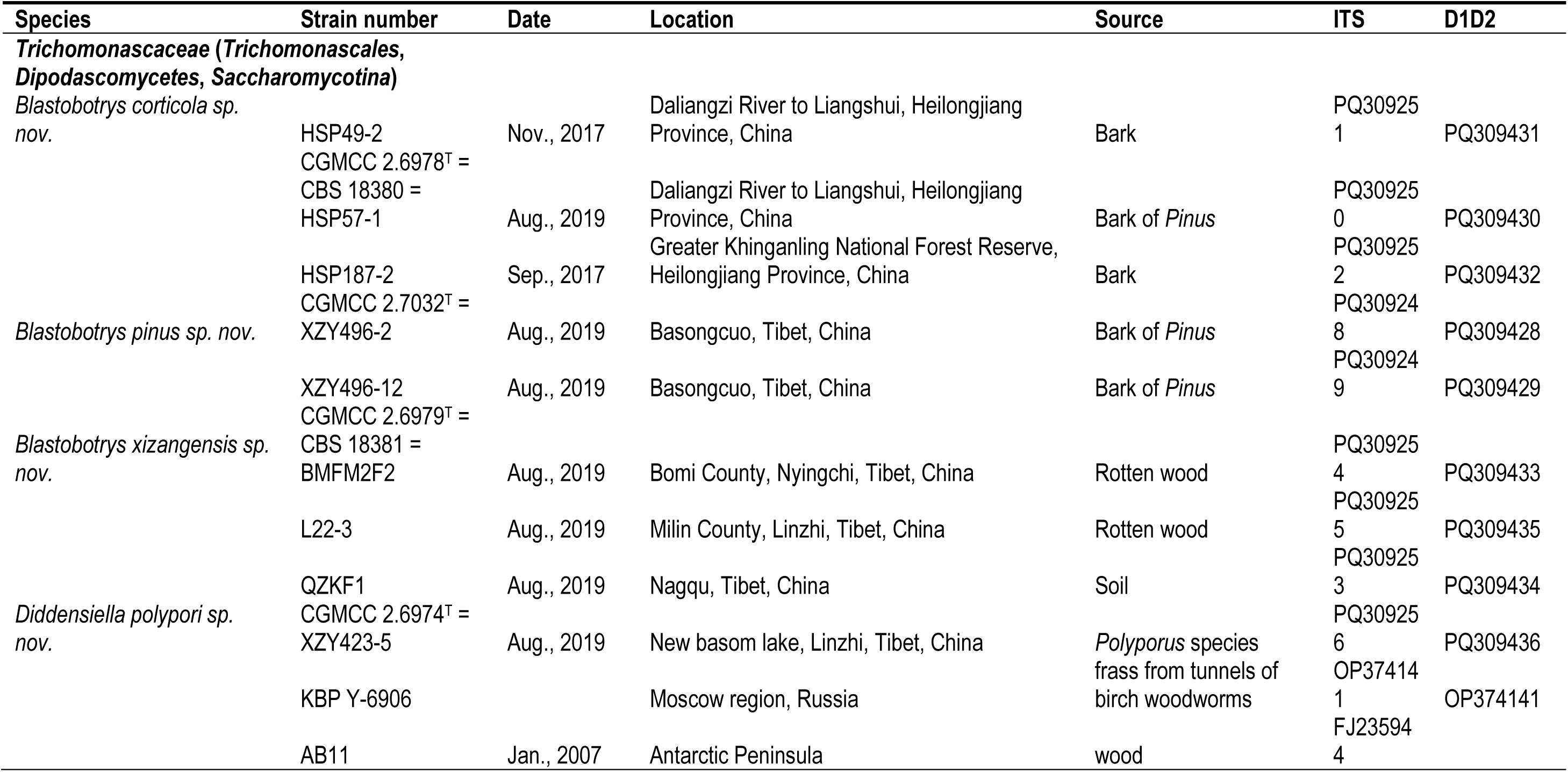
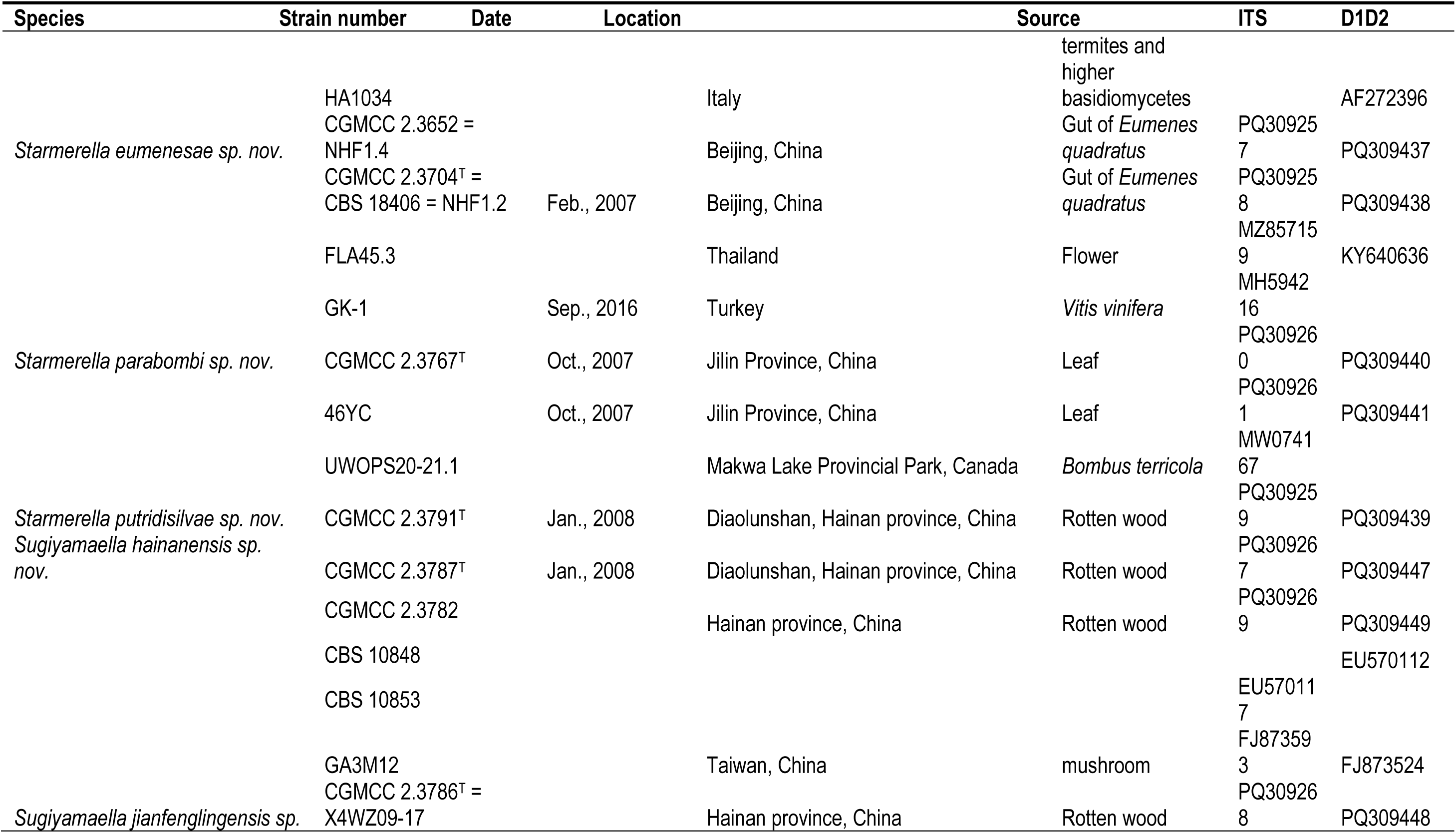
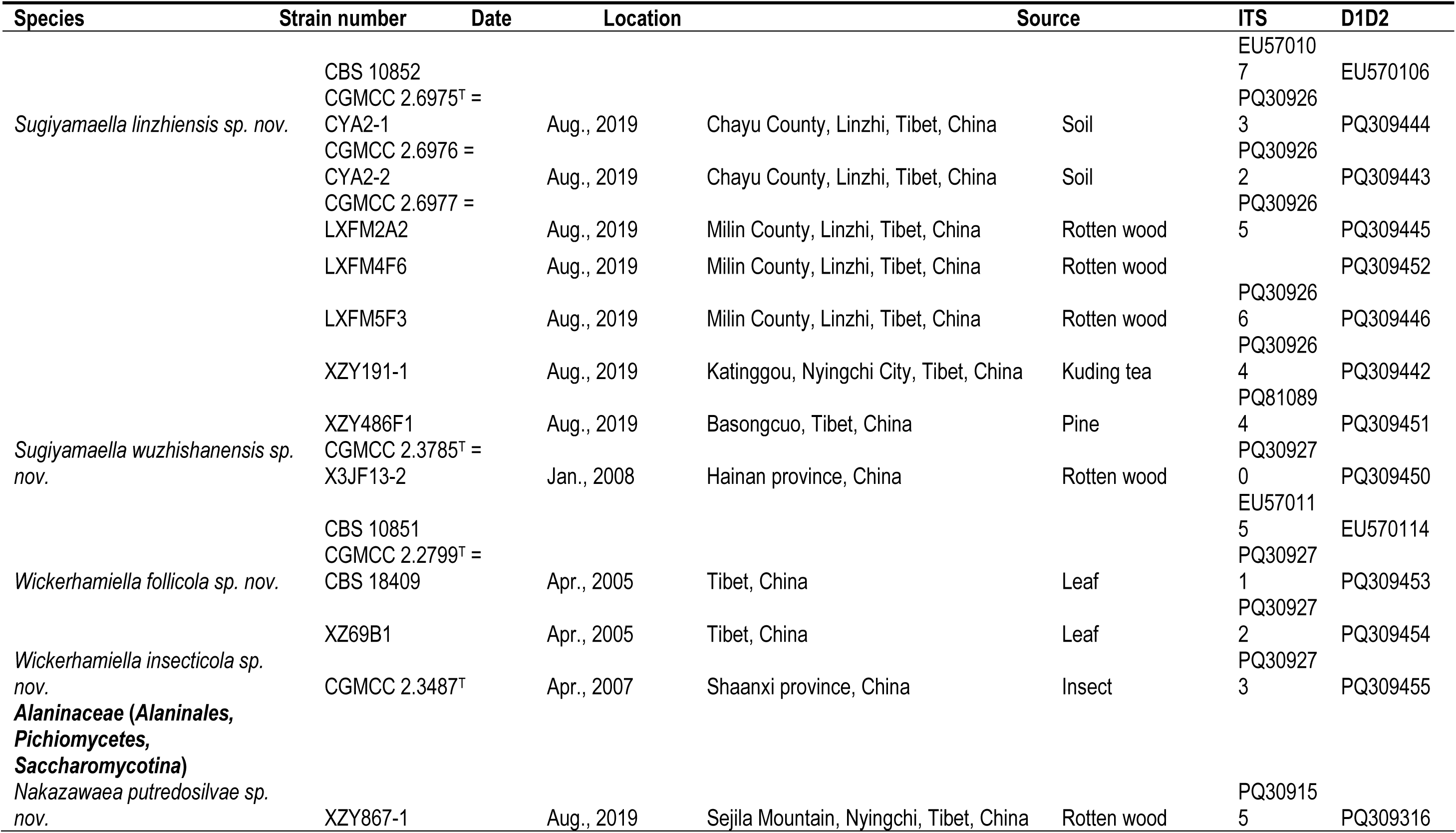
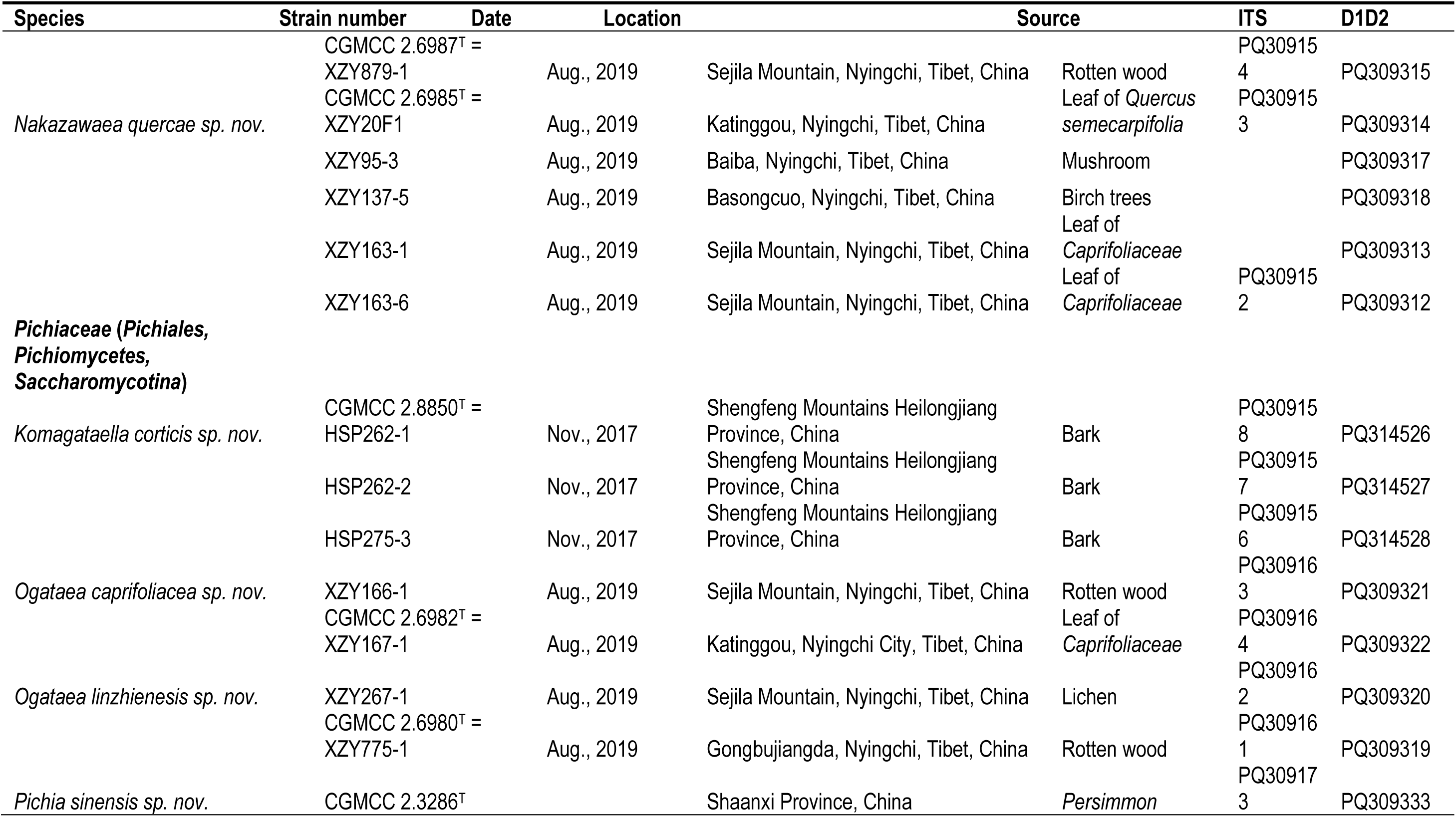
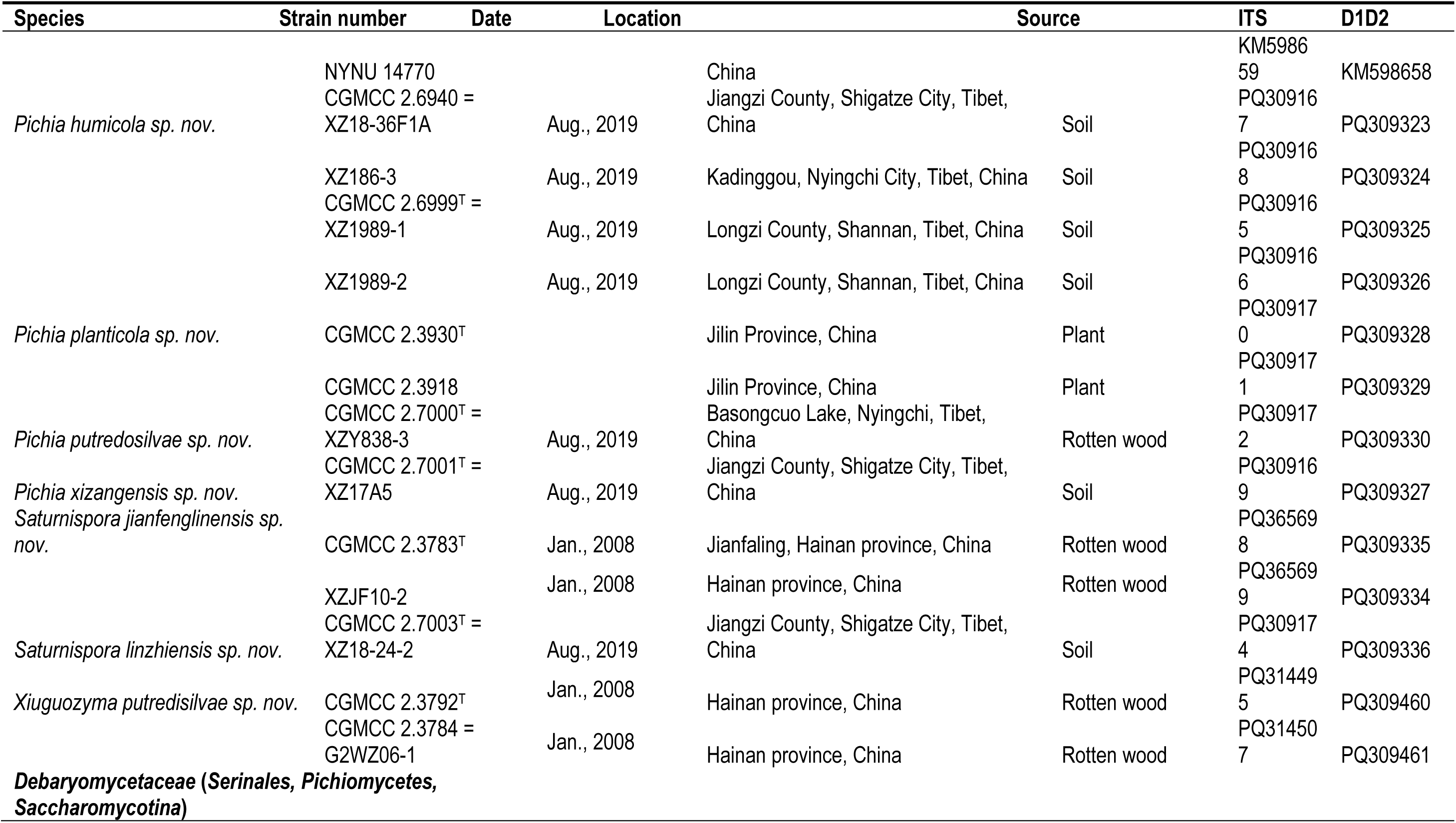
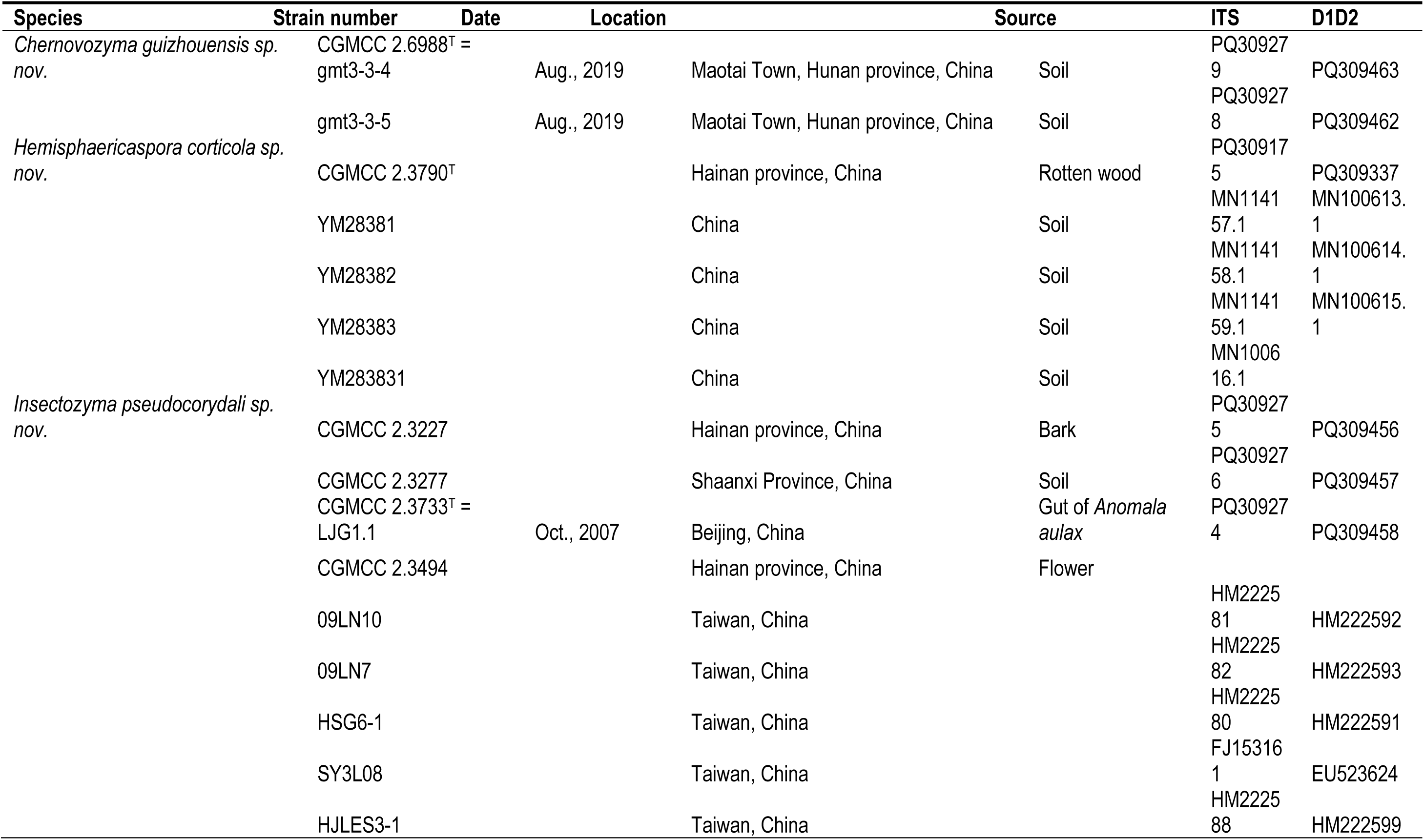
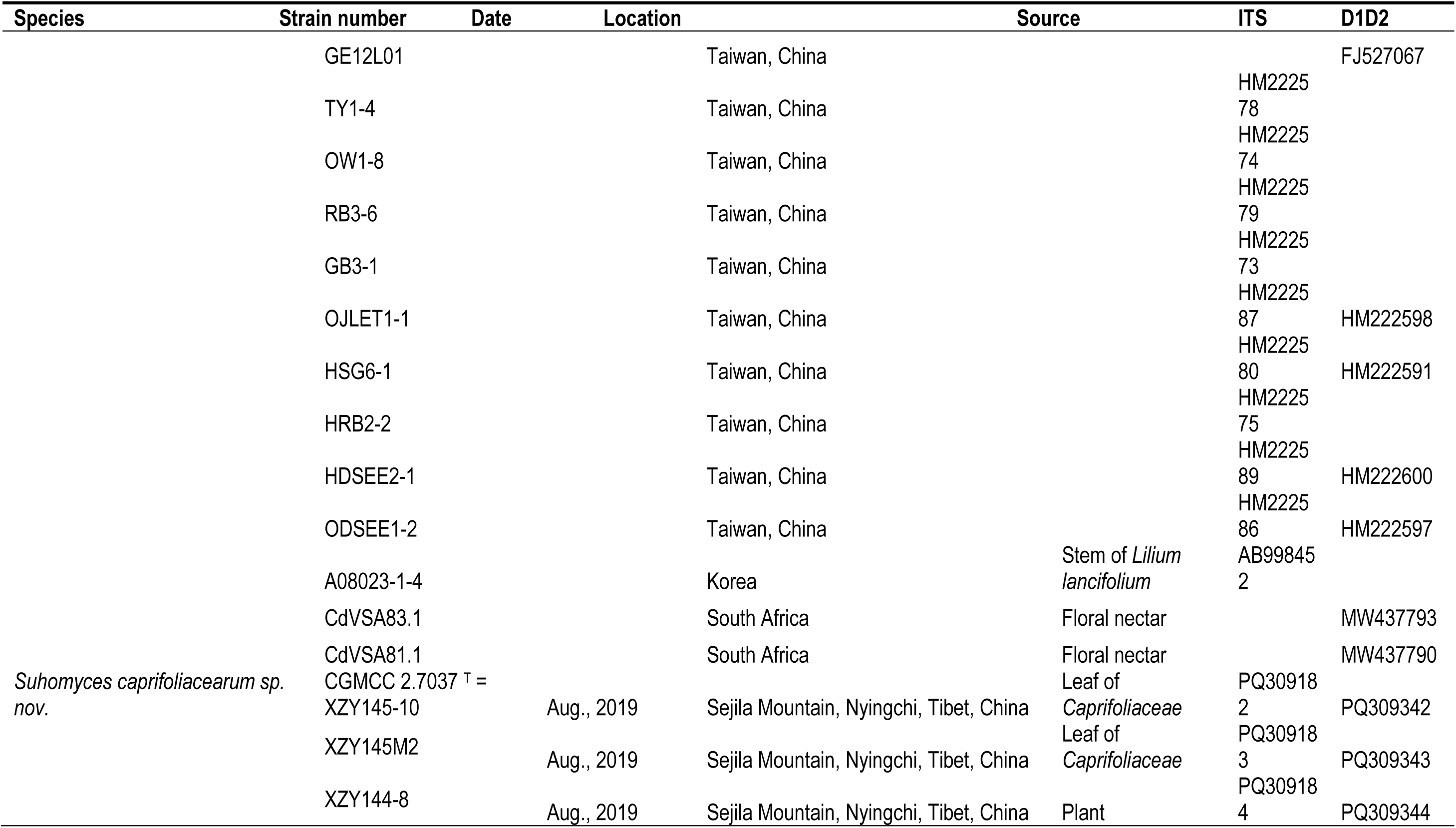
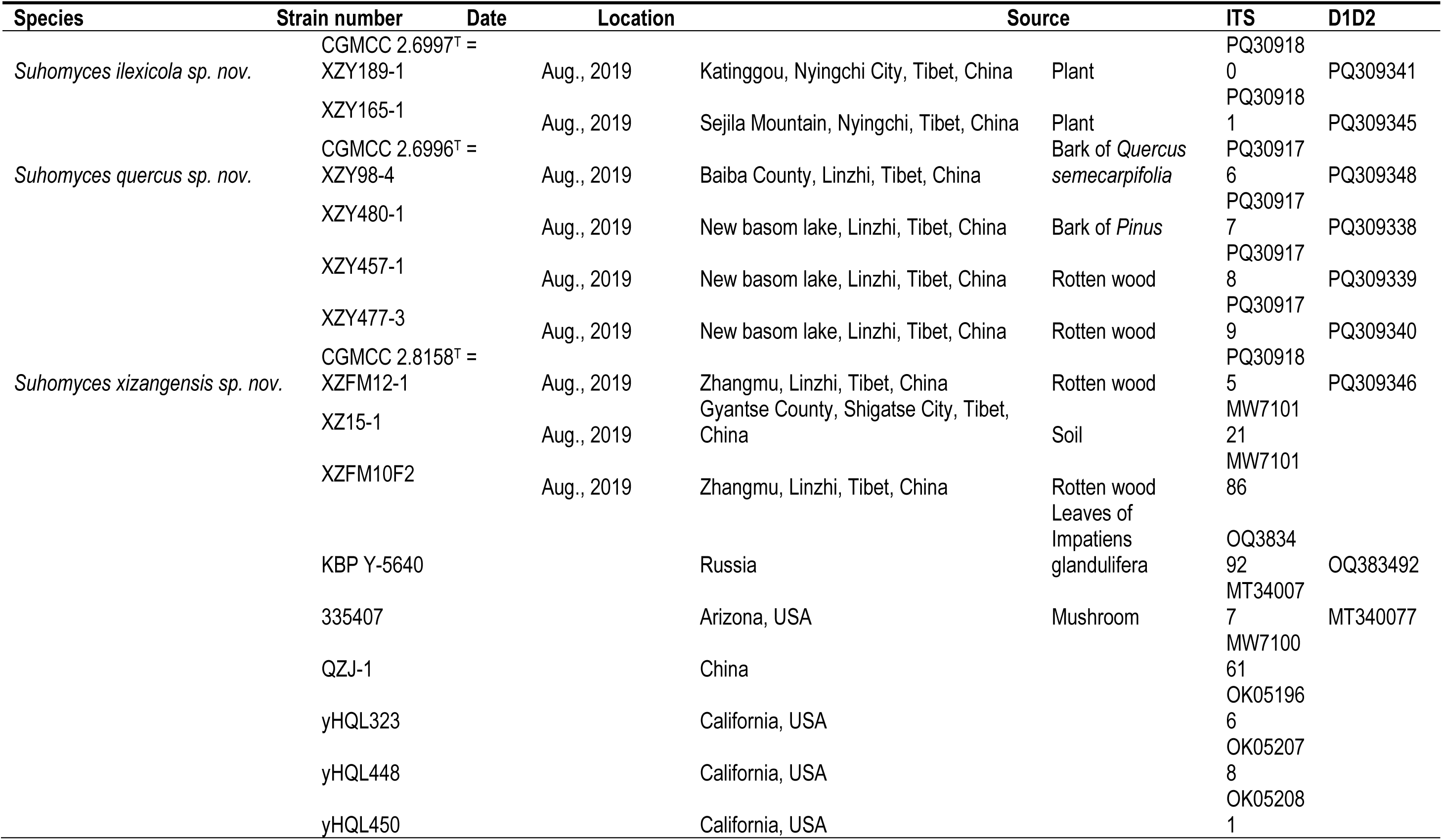
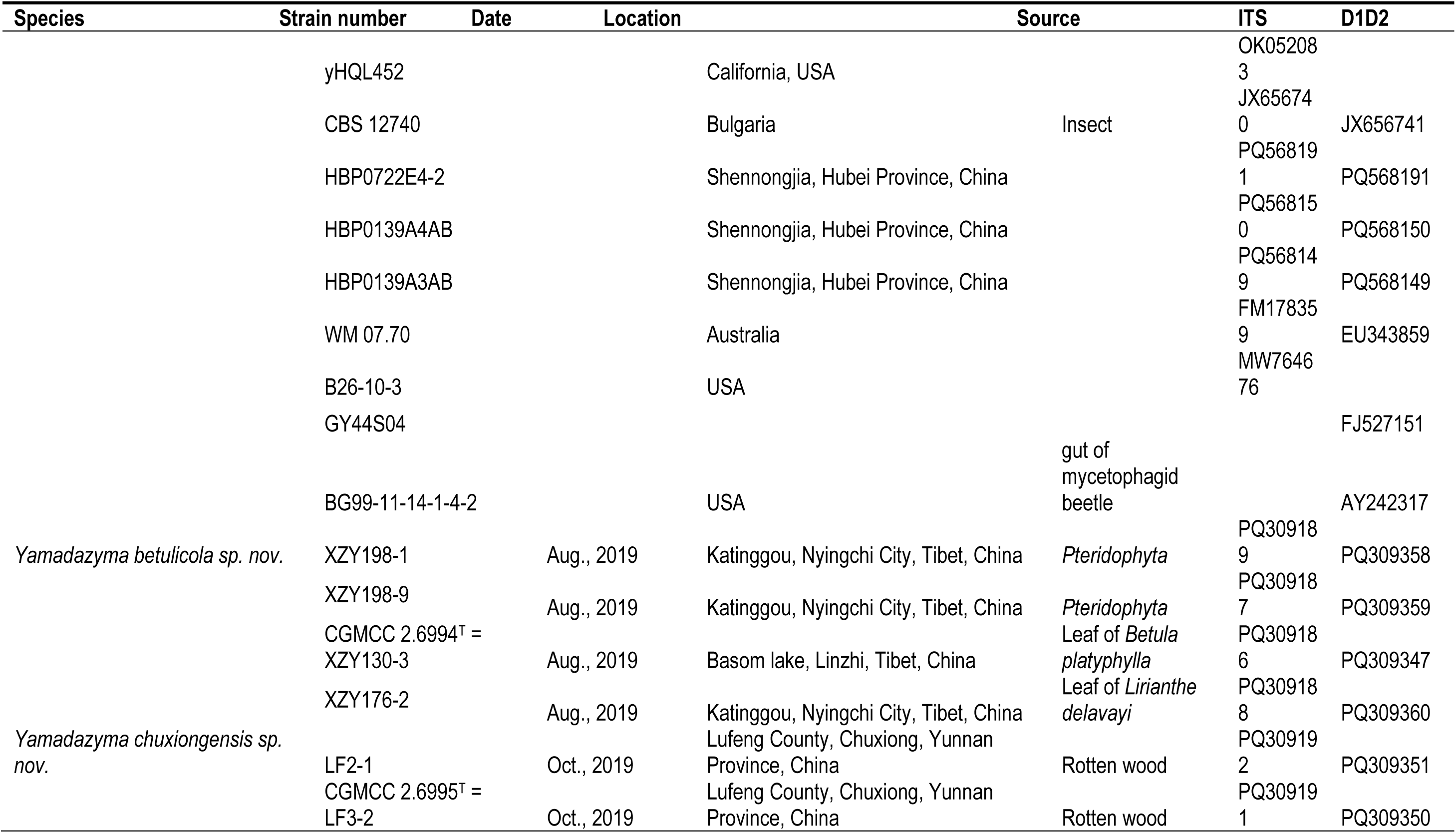
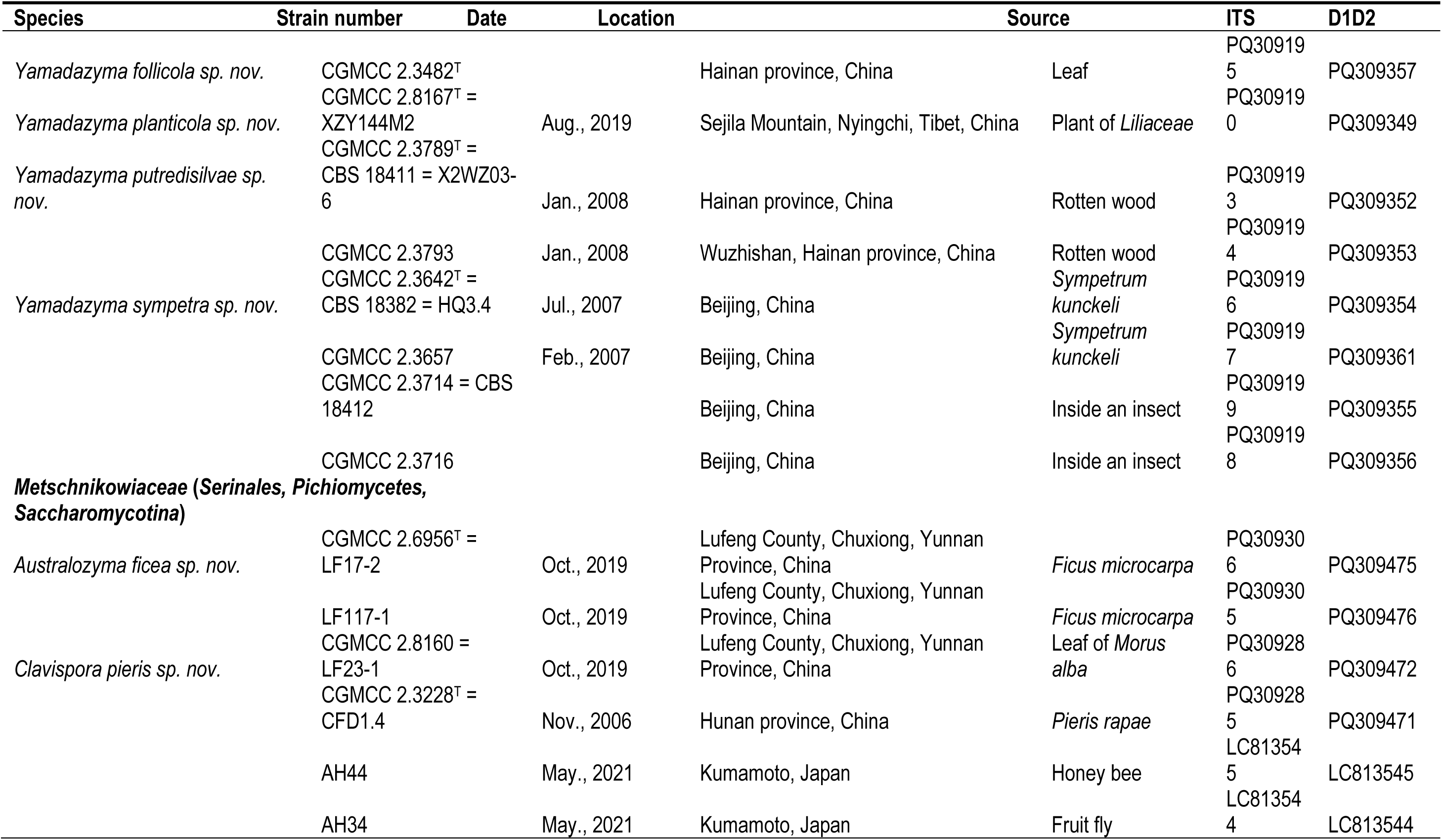
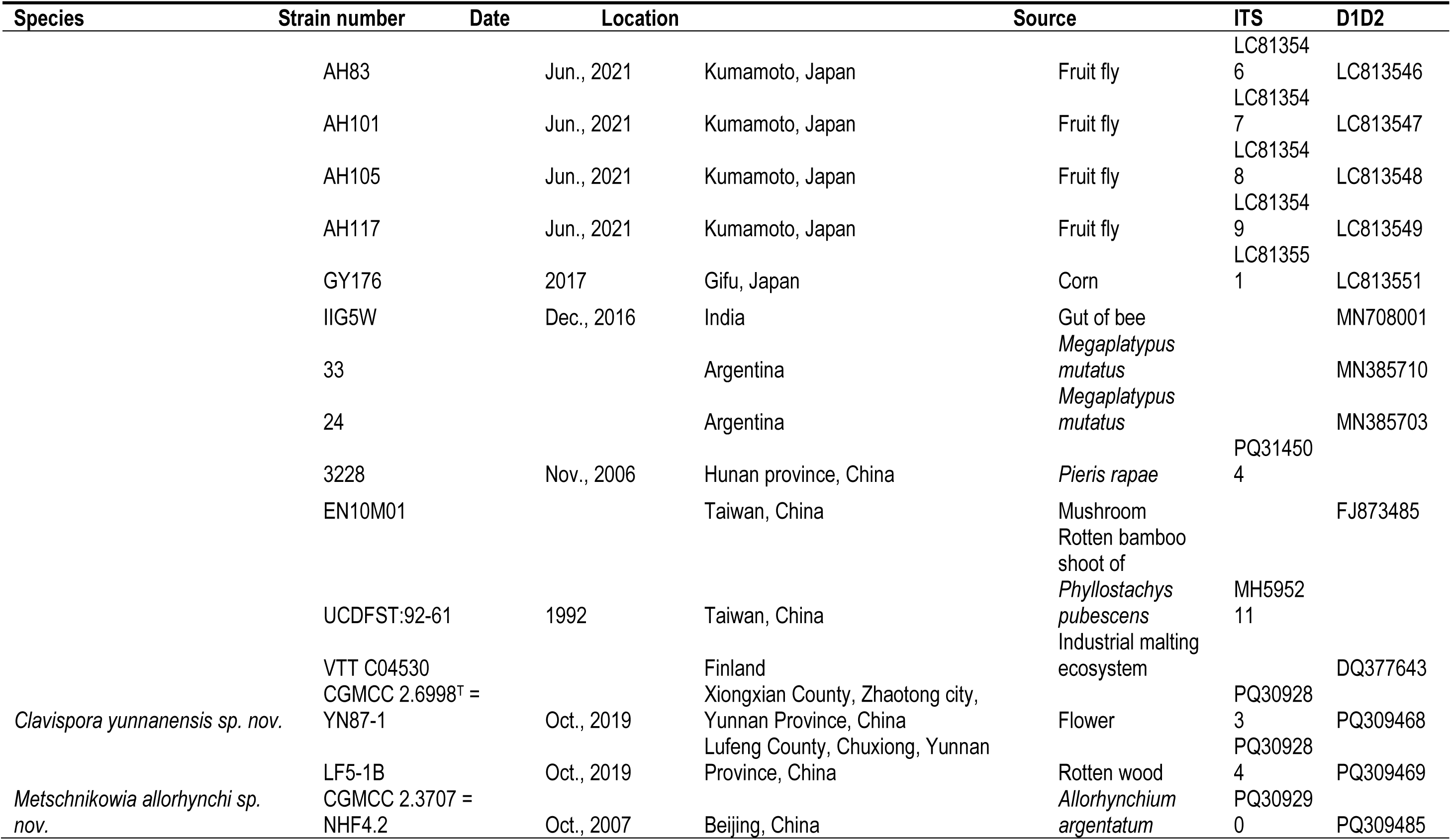
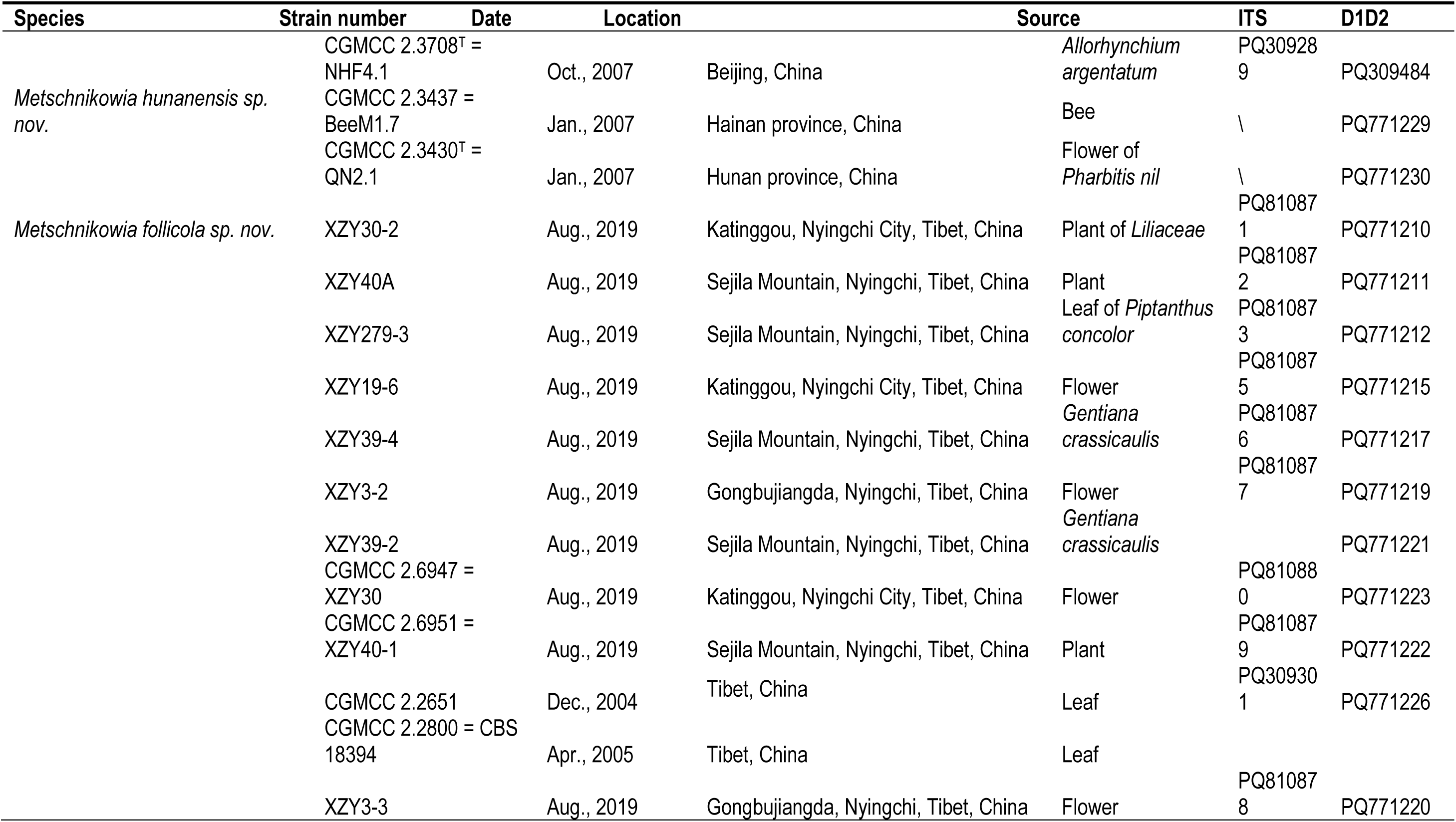
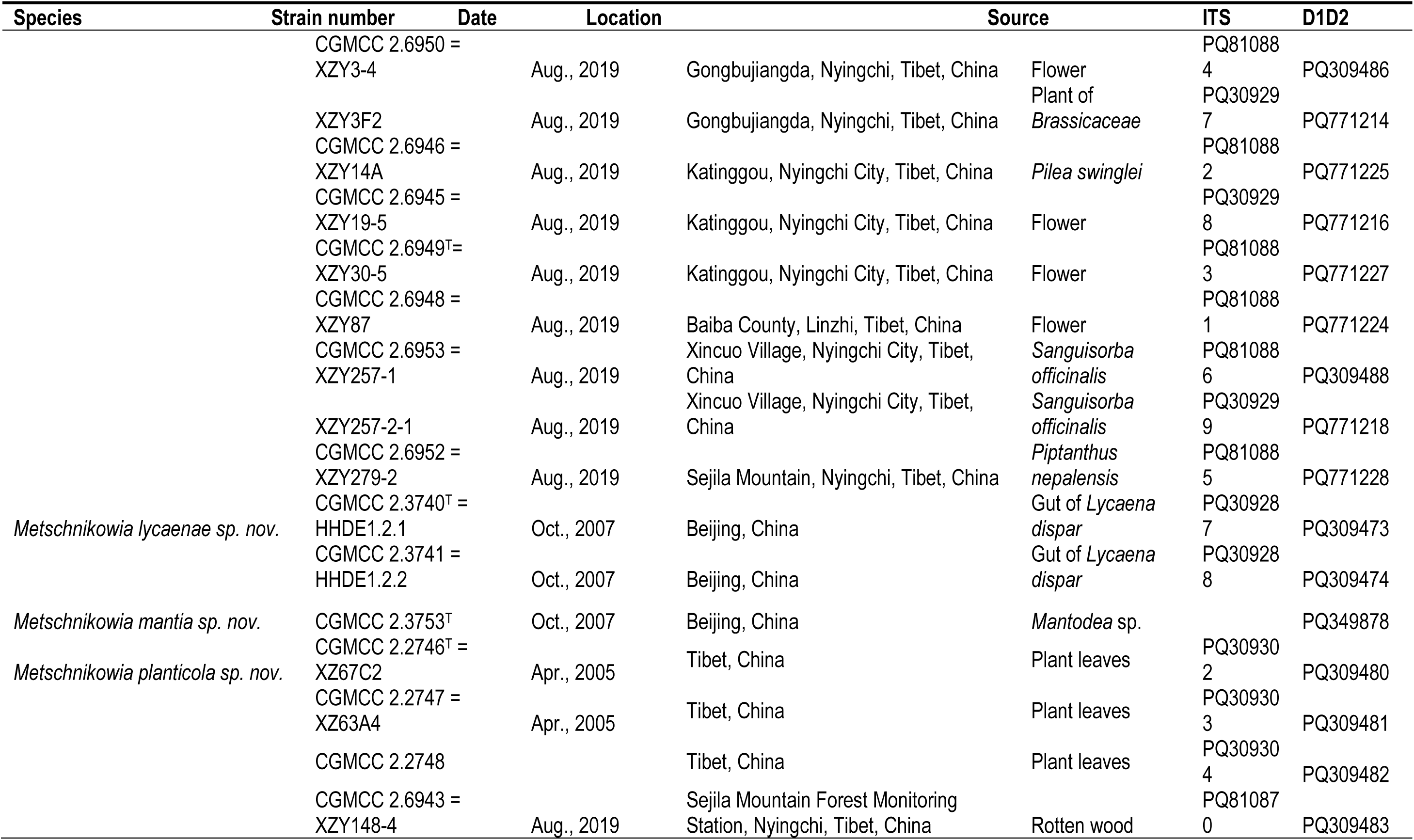
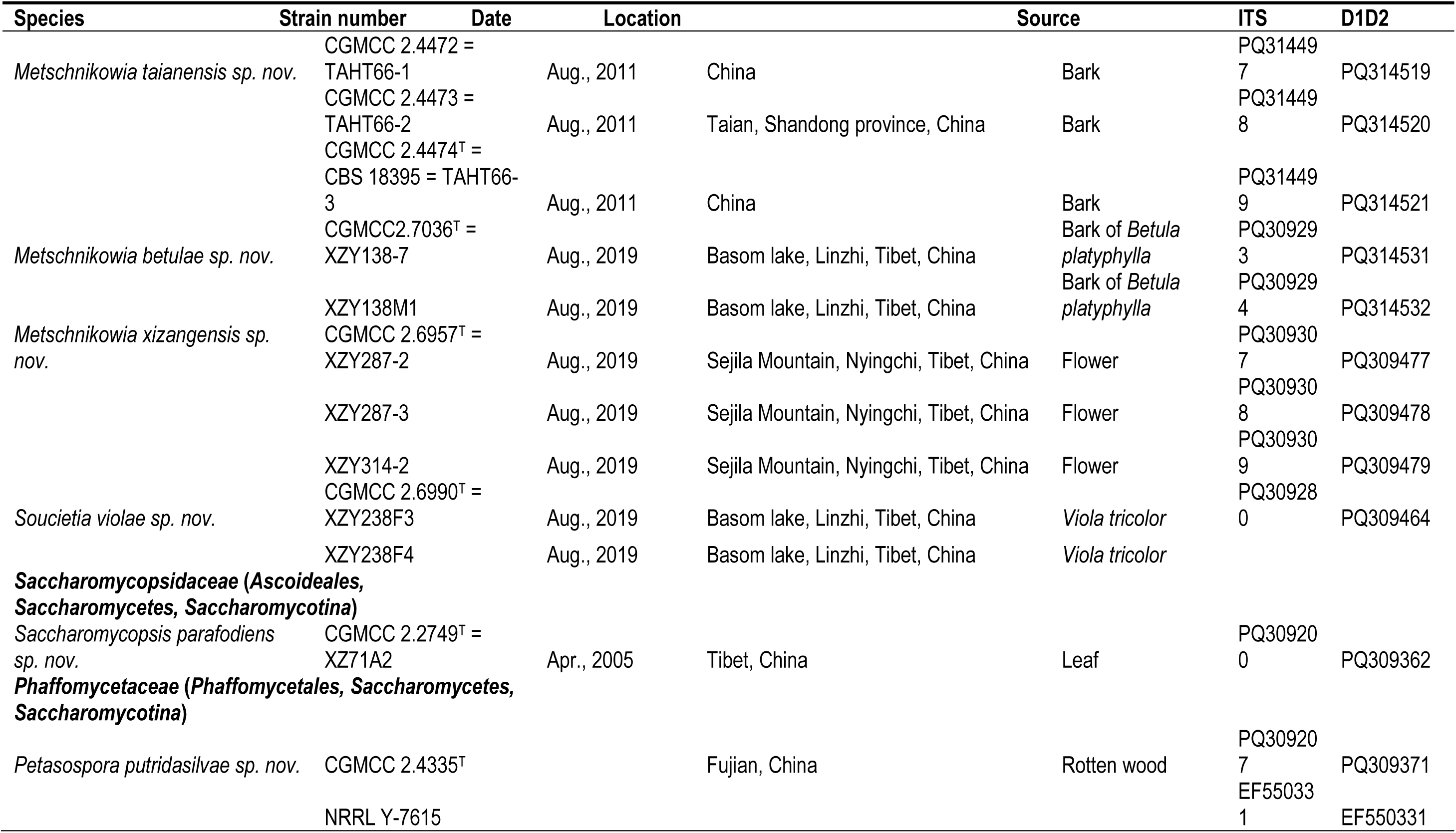
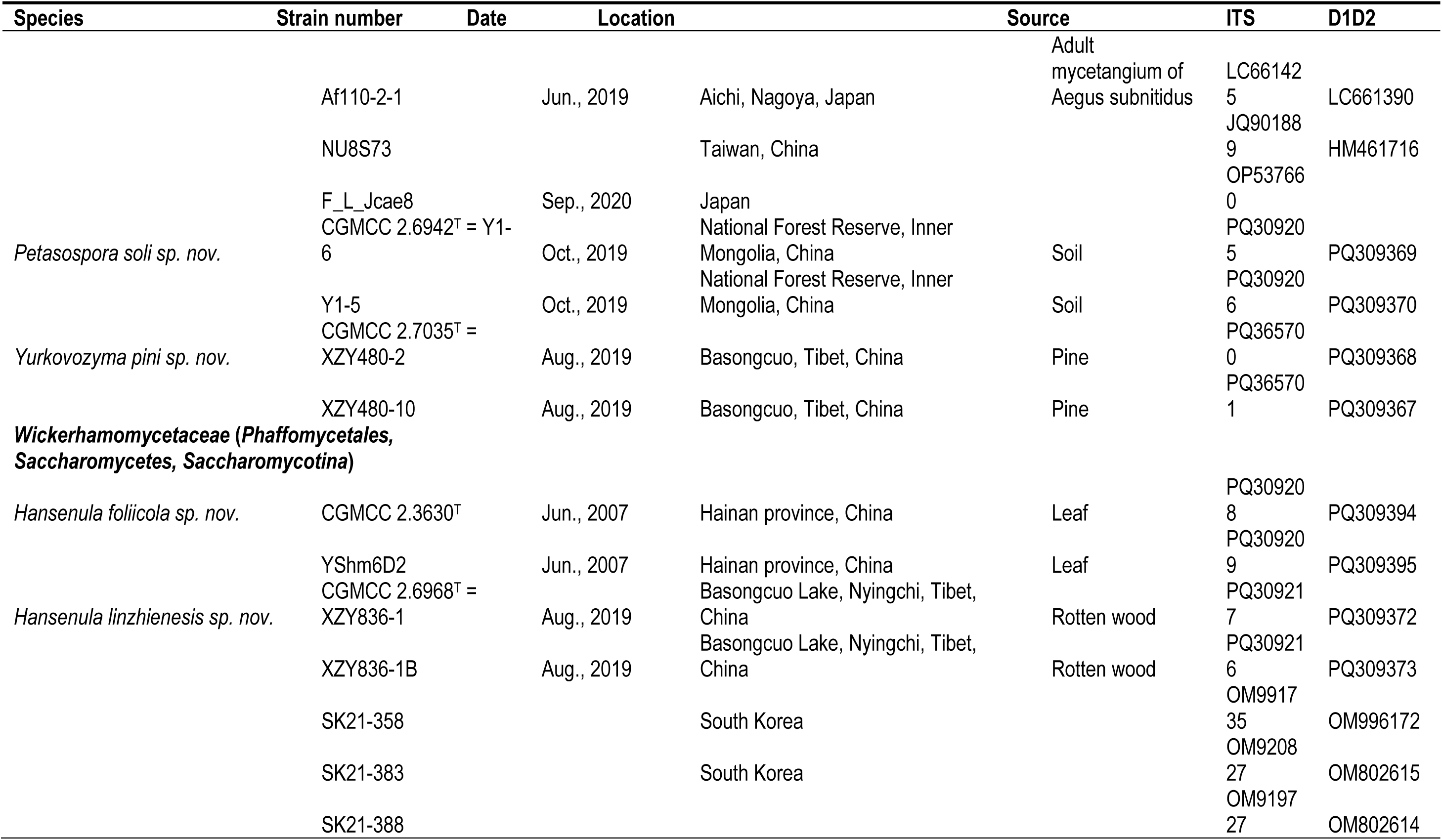
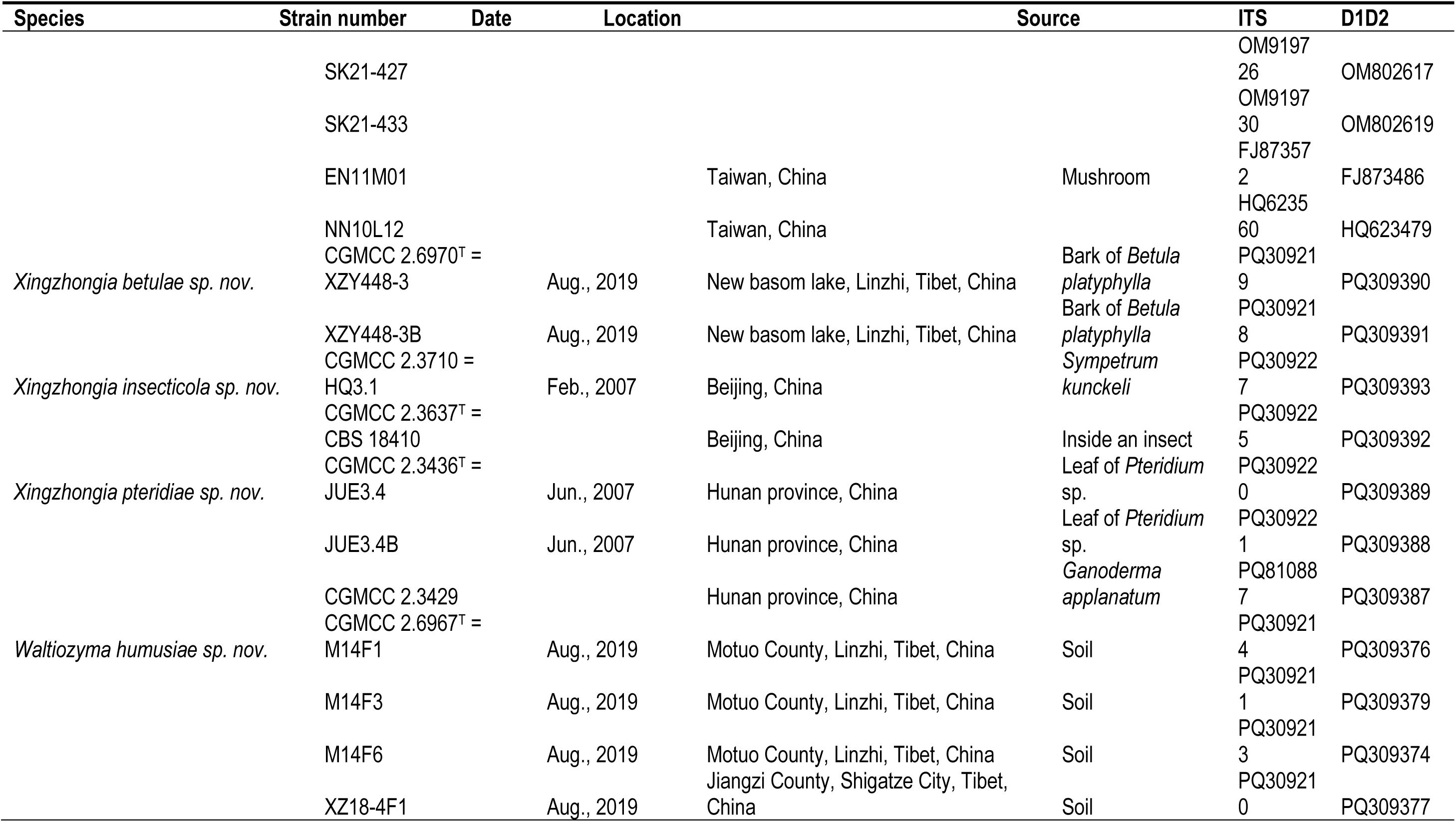
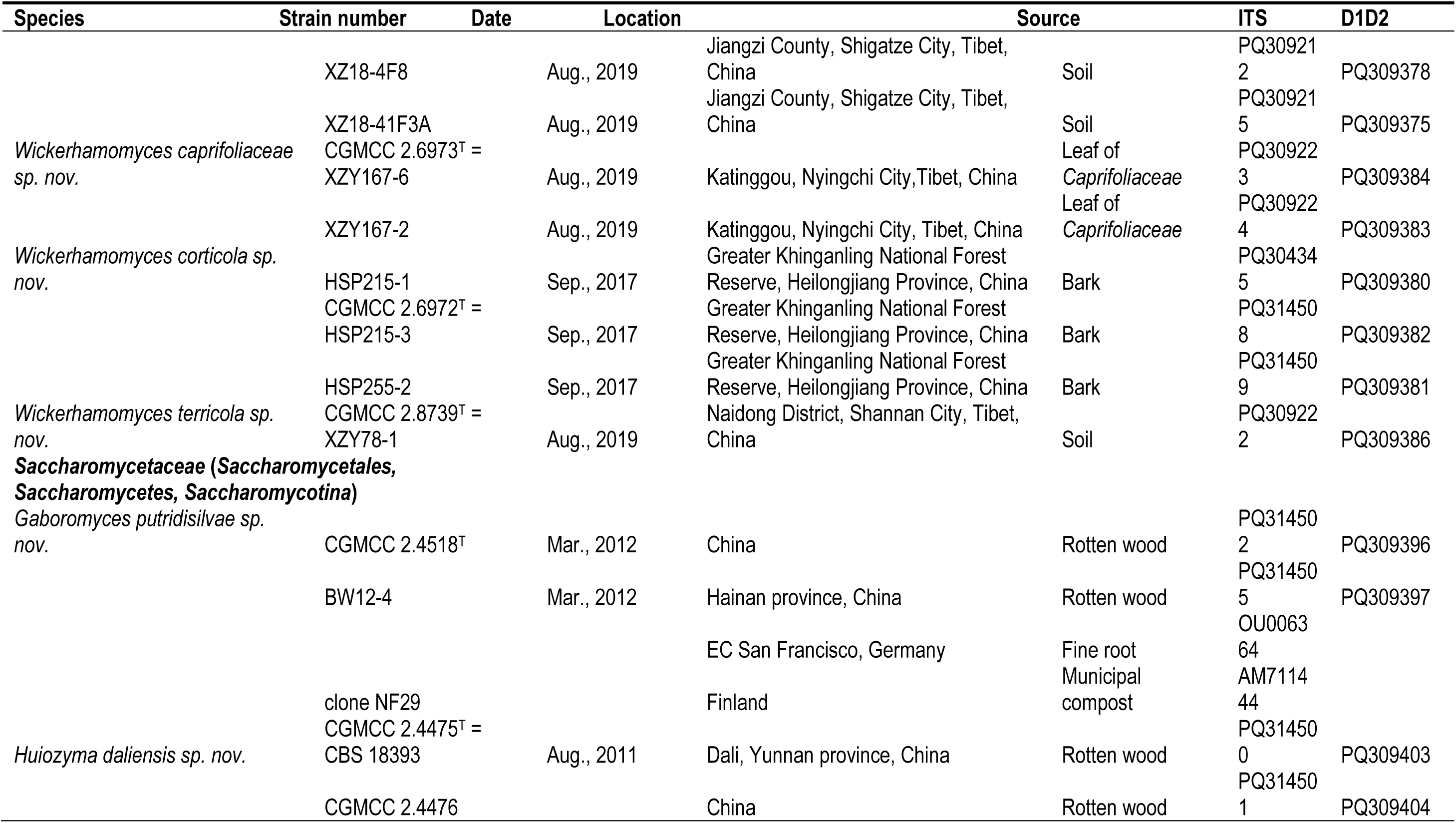
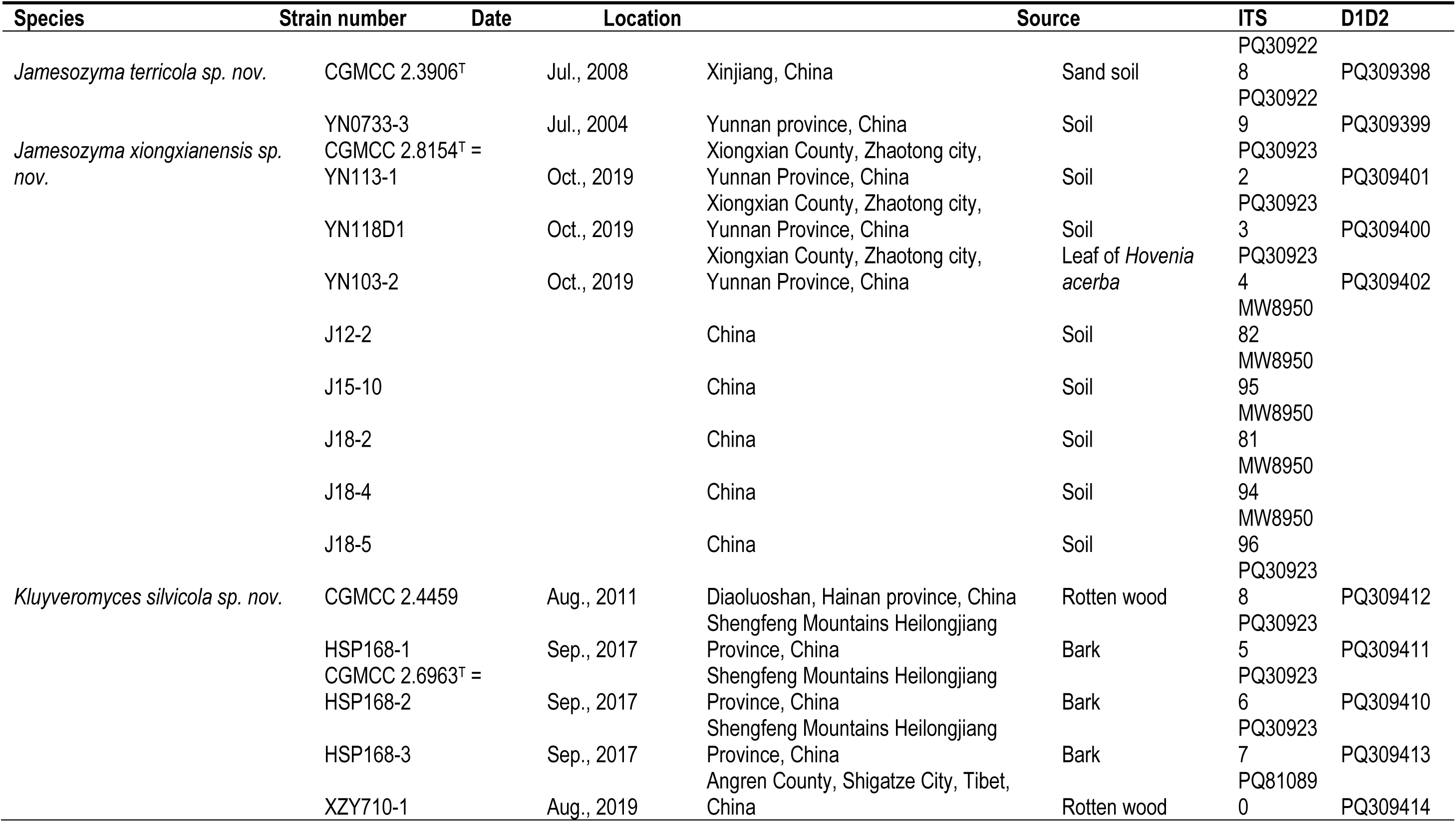
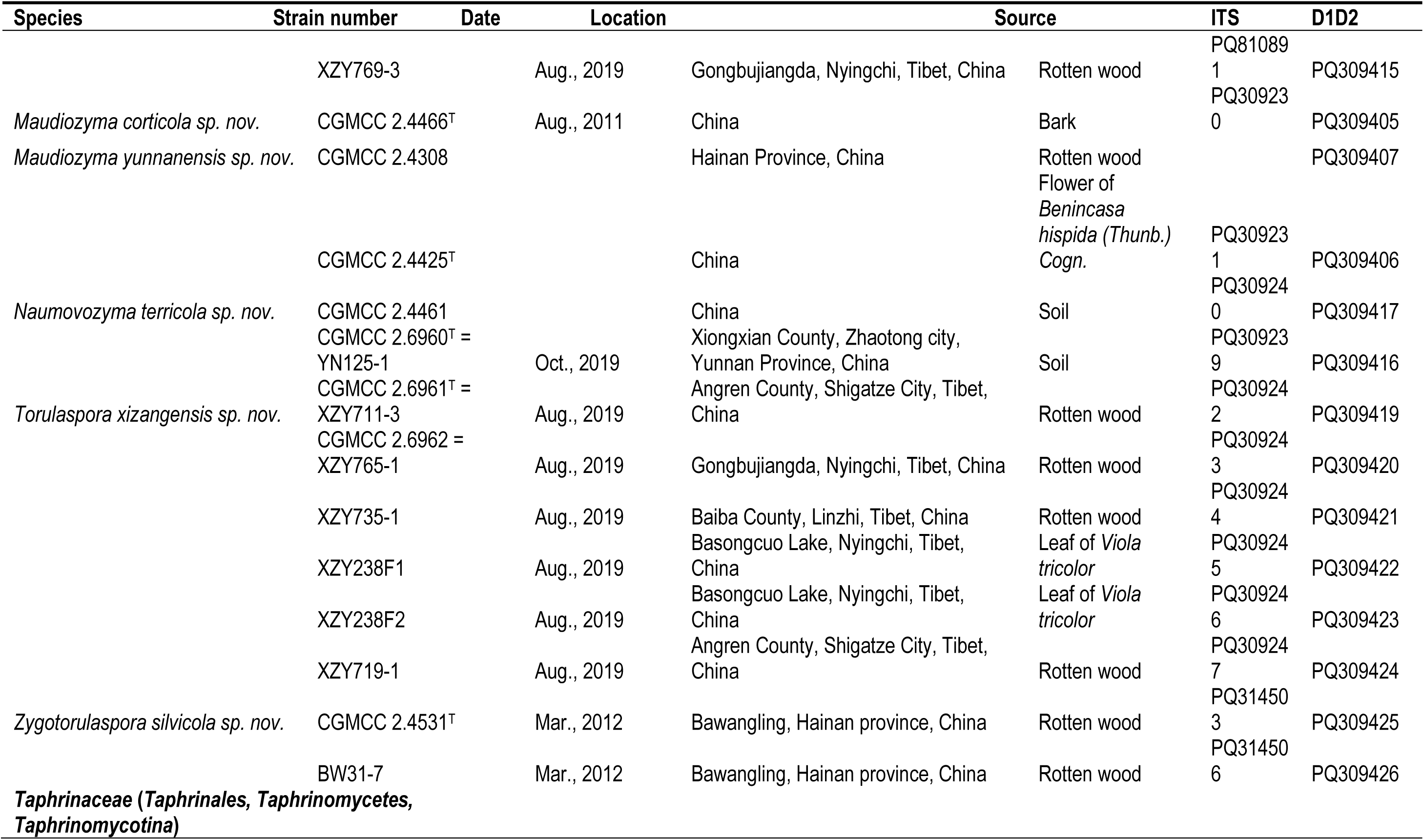
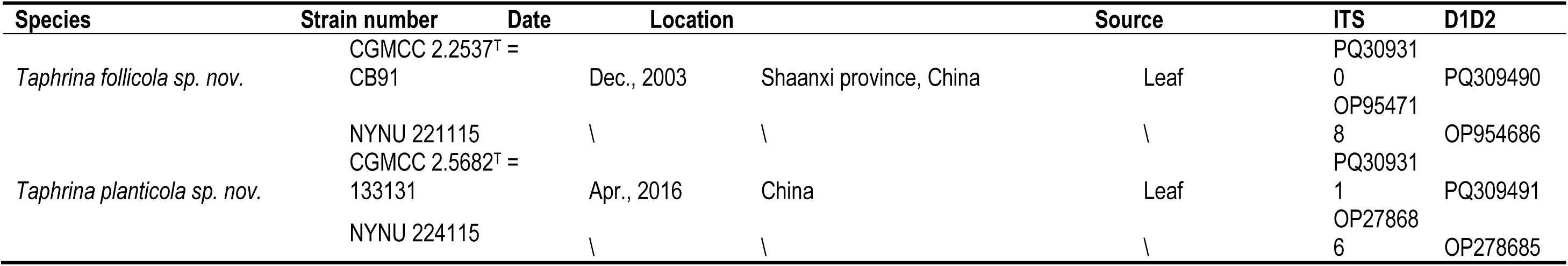
List of yeast and sequence numbers used for the new species described in this study (provided as a separate Excel file).

Samples were collected in 15 mL sterile centrifuge tubes. Cells were enriched by YPD (1% yeast extract, 2% peptone, 2% glucose, 7% ethanol, and 200 mg/L chloramphenicol) for one month at 25 °C. 0.1 ml of enrichment medium was diluted tenfold gradually to a suitable concentration and evenly spread onto the YPD medium at 25 °C for 4 days. Yeast and yeast-like colonies on the plates were isolated and purified 2 times, identified, and stored in 25 % glycerol at −80 °C.

The morphological characteristics and physiological and biochemical tests were examined using standard methods (Kurtzman *et al*. 2011b). Carbon source and nitrogen compound assimilation experiments were examined in a liquid medium. Potential sexual cycles of strains possibly representing new species were investigated using McClary medium 1 (MC1, 0.5 % sodium acetate trihydrate and 2 % agar, pH 6.5–7.0), McClary medium 2 (MC2, 0.82% sodium acetate trihydrate, 0.18 % potassium chloride, 0.1 % glucose, 0.25 % yeast extract, and 2 % agar, pH 6.5–7.0) and V8 agar (10 % V8 juice and 2 % agar). The inoculated agar plates were incubated at 25 °C for up to one month.

### DNA extraction and ribosomal DNA sequencing

DNA extraction from yeast cells was performed using the previously described method (Wang & Bai 2008). The nuclear ribosomal internal transcribed spacers (ITS1, ITS2 and 5.8S) region and the 26 S rDNA D1/D2 domains were amplified and sequenced using the previously described method (Wang *et al*. 2014). Pairwise comparisons of sequences were performed by BLAST search (Altschul *et al*. 1997). The newly produced sequences and those of related species retrieved from the GenBank database were aligned using the multiple alignment program MEGA7 (Kumar *et al*. 2016). The best nucleotide substitution models for each dataset were determined with the program MEGA7. Phylogenetic trees were reconstructed based on the sequences in the D1/D2 domains of LSU, the internal transcribed spacers (ITS1 and ITS2) and the 5.8S ribosomal DNA gene. Evolutionary distance data were calculated in the maximum likelihood (ML) analysis. Confidence limits were estimated from the bootstrap analysis (1000 replicates) (Felsenstein *et al*. 1985), and only values >50 % were recorded on the resulting trees. The ITS and D1/D2 sequences determined in this study were deposited in GenBank (Table 1). GenBank accession numbers for other known species are listed in Table S1.

### Genome sequencing, assembly, gene prediction

Genomic libraries (150 bp paired-end) were constructed following the manufacturer’s protocols of TruSeq DNA Nano library prep kit (Illumina) and sequenced on an Illumina HiSeq 2000 platform using TruSeq SBS Kit (Illumina). The adapter sequence and low-quality reads were removed with default parameters using Fastp v. 0.20.1 (Chen et al. 2018). SPAdes v. 3.15.0 (Bankevich et al. 2012) was used to assemble the genomes of the above yeast strains with the following parameters: “--memory 800 -k 21,33,55,77,99 --care ful --cov-cutoff auto”. Gene prediction was done using GeneMark-ES (Ter-Hovhannisyan et al. 2008). The information about the assembly and annotation genomes was listed in Table 2.

**Table 2.** List of genomes used in this study (provided as a separate Excel file).

| Source | Species | Strain | Assembly | Nums of<br>contig | Largest<br>contig | Total length<br>(Mb) | GC (%) | N50 | Nums of<br>gene |
| --- | --- | --- | --- | --- | --- | --- | --- | --- | --- |
| NCBI | *Blastobotrys buckinghamii* | CBS 13900 | GCA_030579975.1 | 949 | 317086 | 25820393 | 31.59 | 85955 | 7466 |
| this study | *Blastobotrys corticola* | HSP49-2 | NMDC60203897 | 368 | 1015553 | 25712679 | 29.35 | 219566 | 7387 |
| this study | *Blastobotrys corticola* | HSP57-1 | NMDC60203898 | 471 | 1001401 | 37487288 | 31.67 | 239153 | 13556 |
| this study | *Blastobotrys corticola* | HSP187-2 | NMDC60203899 | 540 | 808955 | 35863958 | 32.75 | 169811 | 12438 |
| this study | *Blastobotrys xizangensis* | BMFM2F2 | NMDC60192684 | 9 | 2061164 | 3898518 | 46.43 | 2061164 | 1680 |
| this study | *Chernovozyma guizhouensis* | gmt3-3-4 | NMDC60203901 | 77 | 1060152 | 14161975 | 45.61 | 410398 | 5591 |
| this study | *Chernovozyma guizhouensis* | gmt3-3-5 | NMDC60203902 | 62 | 1555233 | 14161366 | 45.61 | 595897 | 5593 |
| this study | *Chernovozyma guizhouensis* | yHMH407 | NMDC60203900 | 586 | 510040 | 14119388 | 45.15 | 63756 | 5583 |
| this study | *Clavispora pieris* | LF23-1 | NMDC60203903 | 117 | 1353122 | 18126445 | 53.46 | 531460 | 6287 |
| this study | *Clavispora pieris* | CGMCC 2.3228 | NMDC60203904 | 65 | 1669373 | 11297394 | 49.11 | 898037 | 5155 |
| this study | *Gaboromyces putridisilvae* | CGMCC 2.4518 | NMDC60192670 | 73 | 751305 | 10643309 | 34.1 | 344664 | 5126 |
| this study | *Starmerella parabombi* | CGMCC 2.3767 | NMDC60203905 | 86 | 1055779 | 10490506 | 41.61 | 475113 | 4977 |
| NCBI | *Jamesozyma jinghongensis* | CBS 15232 | GCA_030572895.1 | 562 | 264844 | 11642923 | 35.51 | 55136 | 5361 |
| this study | *Jamesozyma terricola* | CGMCC 2.3906 | NMDC60192668 | 239 | 621306 | 11963063 | 36.56 | 123606 | 5411 |
| this study | *Jamesozyma xiongxiensis* | YN113-1 | NMDC60203865 | 398 | 352385 | 12180590 | 36.46 | 74584 | 5473 |
| this study | *Jamesozyma xiongxiensis* | YN118D1 | NMDC60209938 | 532 | 402218 | 19049559 | 45.12 | 75880 | 6888 |
| NCBI | *Kazachstania psychrophila* | CBS 12689 | GCA_030579255.1 | 99 | 844606 | 10504644 | 33.29 | 324962 | 5127 |
| NCBI | *Kluyveromyces dobzhanskii* | CBS 2104 | GCA 000820885.1 | 86 | 1379240 | 10741898 | 41.25 | 493016 | 4962 |
| this study | *Kluyveromyces silvicola* | CGMCC 2.4459 | NMDC60203906 | 33 | 2113277 | 10821049 | 40.21 | 952130 | 4848 |
| this study | *Kluyveromyces silvicola* | HSP168-1 | NMDC60203907 | 44 | 1441059 | 10828139 | 40.22 | 962594 | 4840 |
| this study | *Kluyveromyces silvicola* | HSP168-2 | NMDC60192672 | 37 | 1400756 | 10829617 | 40.21 | 1140079 | 4843 |
| NCBI | *Komagataella pastoris* | CBS 704 | GCA 001708105.1 | 11 | 3325186 | 9597535 | 41.47 | 2687137 | 5305 |
| this study | *Komagataella corticis* | HSP262-1 | NMDC60203908 | 35 | 1867074 | 9377848 | 41 | 1389144 | 4989 |
| this study | *Komagataella corticis* | HSP262-2 | NMDC60203909 | 105 | 1867055 | 21149723 | 38.64 | 652490 | 11121 |
| this study | *Komagataella corticis* | HSP275-3 | NMDC60203910 | 38 | 1867372 | 9373887 | 41 | 1411070 | 4989 |
| this study | *Maudiozyma corticola* | CGMCC 2.4466 | NMDC60203911 | 390 | 320282 | 13337270 | 31.44 | 73520 | 5394 |
| NCBI | *Maudiozyma exigua* | CBS 379 | GCA 030580595.1 | 1185 | 272987 | 25075894 | 32.33 | 53120 | 10433 |
| this study | *Maudiozyma yunnanensis* | CGMCC 2.4425 | NMDC60203912 | 966 | 268187 | 25479012 | 31.61 | 56970 | 10412 |
| this study | *Pichia planticola* | CGMCC 2.3930 | NMDC60203913 | 697 | 260889 | 23545030 | 41.43 | 70548 | 7385 |
| this study | *Pichia xizangensis* | XZ17A5 | NMDC60203914 | 82 | 1279905 | 11764612 | 45.72 | 568322 | 5359 |
| NCBI | *Saccharomycopsis fodiens* | CBS 8332 | GCA 002564235.1 | 22 | 2742214 | 14948151 | 51.3 | 2605998 | 6113 |
| this study | *Saccharomycopsis*<br>*parafodiens* | CGMCC 2.2749 | NMDC60203915 | 475 | 441996 | 17848487 | 46.98 | 79669 | 6072 |
| NCBI | *Starmerella bombi* | NRRL Y-17081 | GCA 030579915.1 | 382 | 1483250 | 11158732 | 47.92 | 407262 | 4470 |
| this study | *Sugiyamaella hainanensis* | CGMCC 2.3787 | NMDC60203916 | 651 | 393861 | 19900182 | 34.74 | 119315 | 6761 |
| this study | *Sugiyamaella*<br>*jianfenglingensis* | CGMCC 2.3786 | NMDC60203917 | 609 | 260889 | 23471924 | 41.45 | 78766 | 7329 |
| this study | *Sugiyamaella*<br>*wuzhishanensis* | CGMCC 2.3785 | NMDC60203918 | 461 | 577079 | 17996893 | 37.47 | 92077 | 6610 |
| this study | *Suhyomyces caprifoliacearum* | XZY144-8 | NMDC60203919 | 166 | 738262 | 14086505 | 41.3 | 200467 | 5638 |
| this study | *Suhyomyces caprifoliacearum* | XZY145-10 | NMDC60203868 | 1120 | 545079 | 21967813 | 37.94 | 147837 | 9617 |
| this study | *Suhyomyces ilexicola* | XZY189-1 | NMDC60192679 | 63 | 1381949 | 13513791 | 38.38 | 553980 | 5555 |
| this study | *Suhyomyces quercus* | XZY457-1 | NMDC60203920 | 285 | 1389170 | 14743506 | 42.18 | 423038 | 5934 |
| this study | *Suhyomyces quercus* | XZY477-3 | NMDC60203921 | 82 | 1389315 | 14396411 | 41.69 | 482657 | 5714 |

| Source | Species | Strain | Assembly | Nums of contig | Largest contig | Total length (Mb) | GC (%) | N50 | Nums of gene |
| --- | --- | --- | --- | --- | --- | --- | --- | --- | --- |
| this study | *Suhyomyces quercus* | XZY98-4 | NMDC60192678 | 80 | 2358830 | 14391396 | 41.7 | 406889 | 5713 |
| this study | *Suhyomyces quercus* | XZY480-1 | NMDC60203922 | 88 | 1389693 | 14413301 | 41.71 | 482555 | 5725 |
| NCBI | *Trichomonascus apis* | CBS 10922 | GCA 030557665.1 | 355 | 677297 | 14285932 | 43.13 | 155056 | 7154 |
| this study | *Yamadazyma betulicola* | XZY130-3 | NMDC60192654 | 48 | 1731235 | 10760569 | 33 | 677229 | 5615 |
| this study | *Yamadazyma planticola* | XZY144M2 | NMDC60203923 | 35 | 2193411 | 11201566 | 36.92 | 1196883 | 5705 |

### ANI analysis

Pyani v.0.2.x (https://widdowquinn.github.io/pyani/) was used for the average nucleotide identity (ANI) to evaluate genomic similarity between microbial strains (Table 3). High-precision forward and backward alignment of the whole genome sequence was performed based on the ANIb algorithm to generate raw ANI alignment data. Then, the average ANI between each pair was calculated as the final ANI of these two strains to construct a multidimensional similarity matrix with species-level resolution (Goris et al. 2007).

**Table 3.**
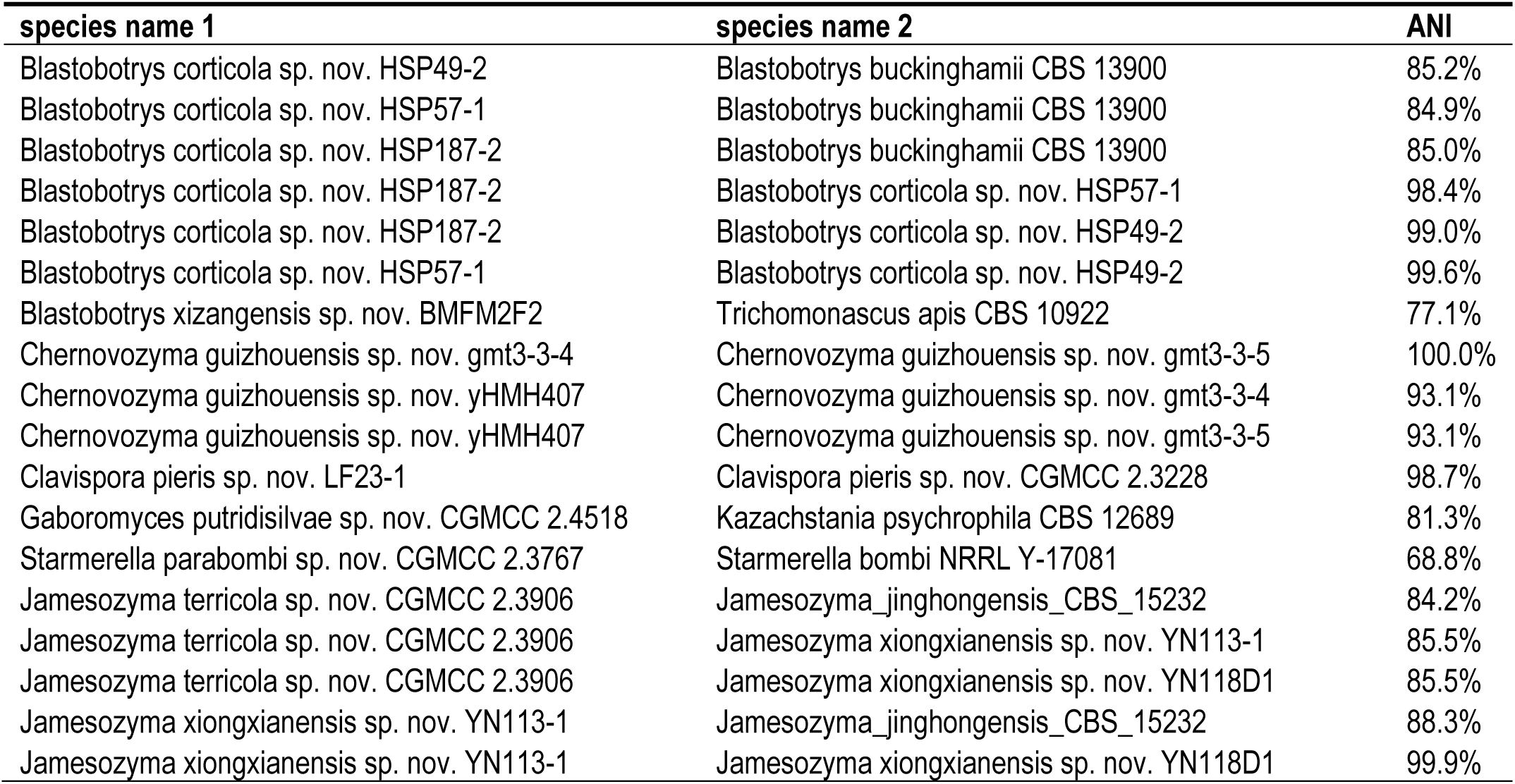
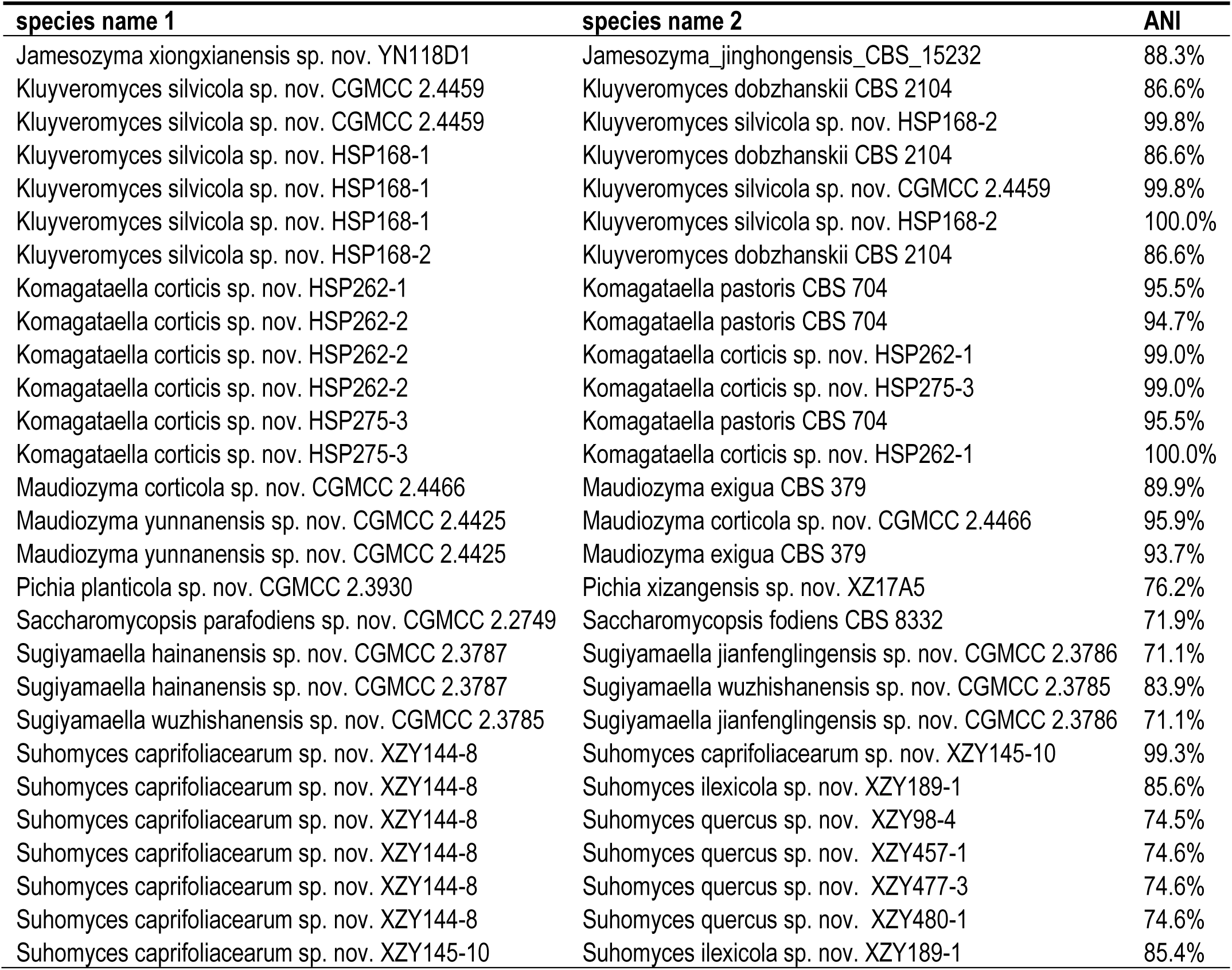
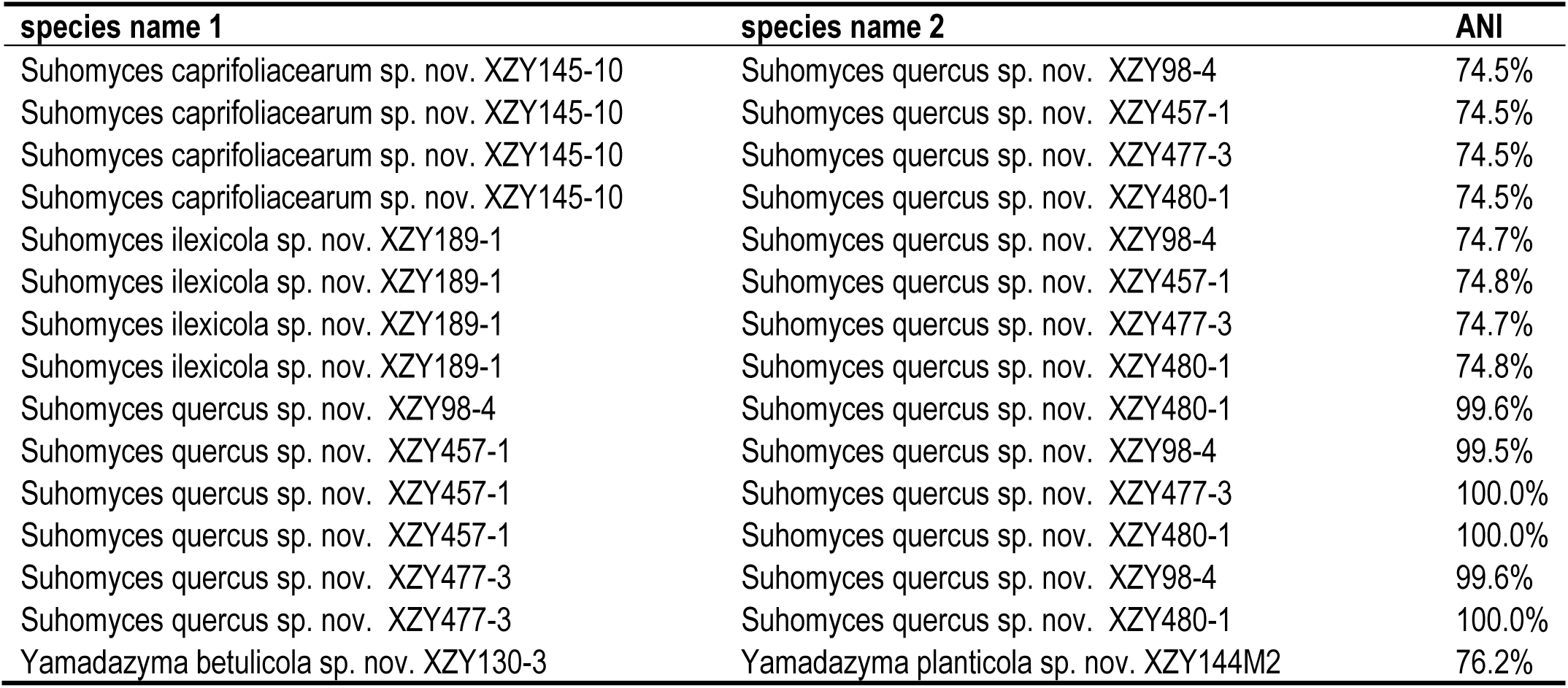
List of the AAI values in this study (provided as a separate Excel file).

## RESULTS AND DISCUSSION

### Samples collection, strain isolation and species delimitation

Over the past 30 years, a series of yeast diversity surveys in China have been done, from which 195 unidentified ascomycetous yeast strains were isolated from various substrates, including bark, rotting wood, soil, flowers, leaves and insects (Table 1). Among these unidentified yeasts, the largest proportion of isolates came from plant material, i.e., rot wood at 24.1 %, followed by leaf (15.9%), soil (14.4%), and bark (12.8%) (Fig. S1). The samples were mainly collected from 12 provinces in China (Fig. 1), with the isolates obtained from Tibet accounting for the largest proportion. These unidentified yeasts offer valuable resources for further exploring yeast species diversity and ecological distribution patterns in China.

**Fig. 1.**
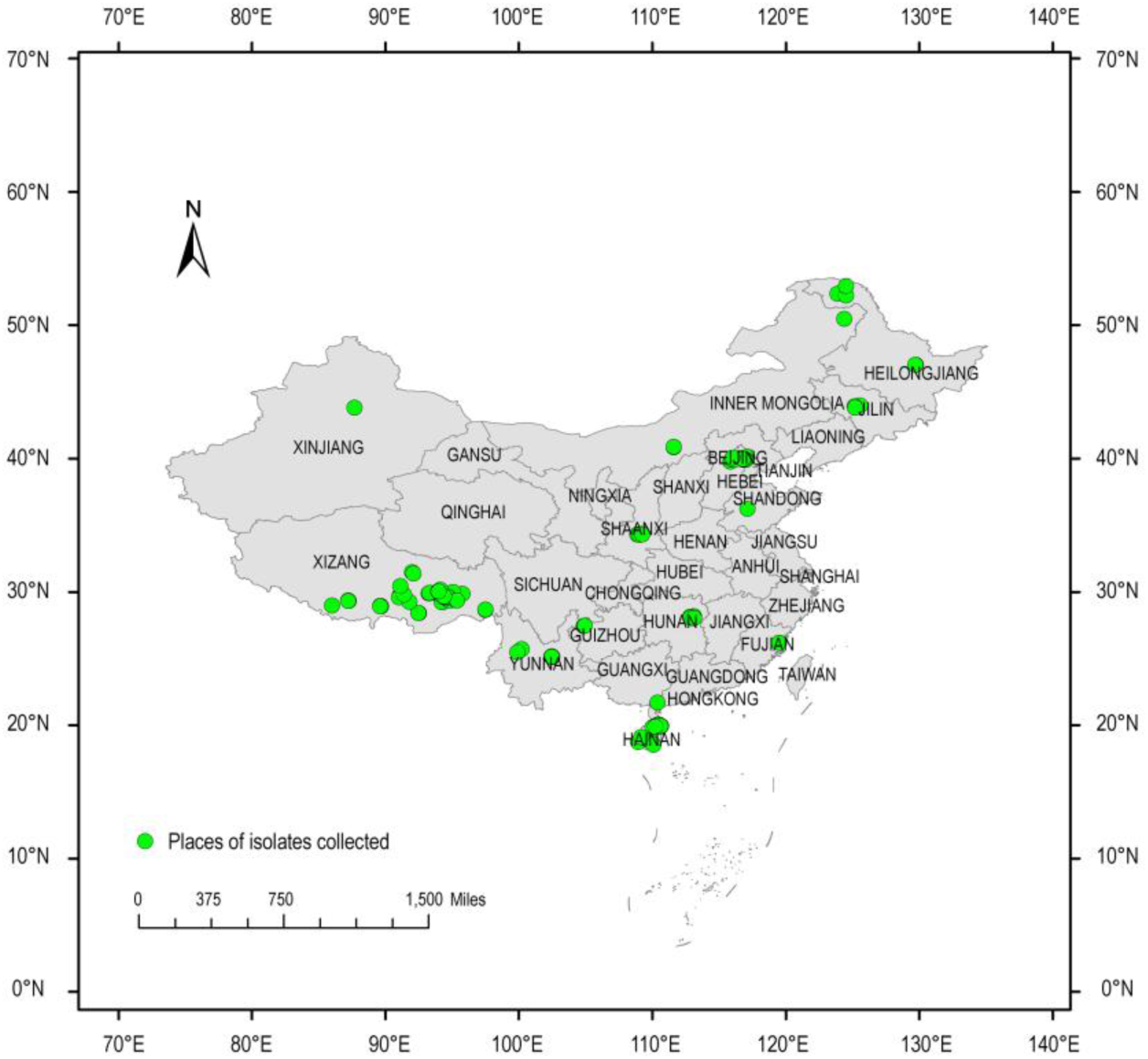
Localisation of sampling sites in China.

Sequence analyses of the D1/D2 and ITS regions have been widely used for yeast species delineation (Kurtzman & Robnett 1998, Fell *et al*. 2000, Scorzetti *et al*. 2002, Kurtzman 2011, Vu *et al*. 2016, Li *et al*. 2020, Boekhout *et al*. 2021, Jiang *et al*. 2024). Kurtzman & Robnett (1998) demonstrated that nucleotide sequences in the LSU rDNA D1/D2 domains from various species tend to show more than 1% sequence divergence. However, strains with three or fewer nucleotide differences are likely to belong to the same or closely related sister species. For most ascomycetous yeasts, there is more sequence diversity in the ITS region than in the D1/D2 domains (Li *et al*. 2020), which means that ascomycetous yeasts with three to six nucleotide differences usually have more than 1 % sequence divergence in the ITS region. Therefore, both ITS and D1/D2 domains serve essential roles in the identification of yeast species (Kurtzman 2011, Boekhout *et al*. 2021, Vu *et al*. 2016, Li *et al*. 2020, Jiang *et al*. 2024). Recently, genome-based metrics, i.e., ANI, have been used to delimitate yeast species, especially for ascomycetous yeasts (Lachance *et al*. 2020, Libkind *et al*. 2020, Troiano *et al*. 2023). Cortimiglia *et al*. (2024) recommended a cutoff of 94–96 % ANI value for yeast species boundaries based on the analysis of 644 assemblies from 12 ascomycetous yeast genera. To confirm the strain’s taxonomic position with low sequence heterogeneity in the ITS and D1/D2 regions, the ANI analysis was performed in this study (Table 3).

### New taxa delineation and phylogenetic placement

#### New species identification in the Trichomonascaceae (Trichomonascales, Dipodascomycetes, Saccharomycotina)

##### Blastobotrys

Eight strains representing three groups are located in the *Blastobotrys/Trichomonascus* lineage (Fig. 2). Strains BMFM2F2, L22-3 and QZKF1 with the same ITS and D1/D2 sequences are closely related to *Trichomonascus apis* (Fig. 2), and differ from it by 1 nucleotide (nt) in the D1/D2 domains and 13 nt (3 %) in the ITS region. The ANI value of those three new isolates and *T. apis* is 77.1 %, which indicates that MFM2F2, L22-3 and QZKF1 represent a new species.

**Fig. 2.**
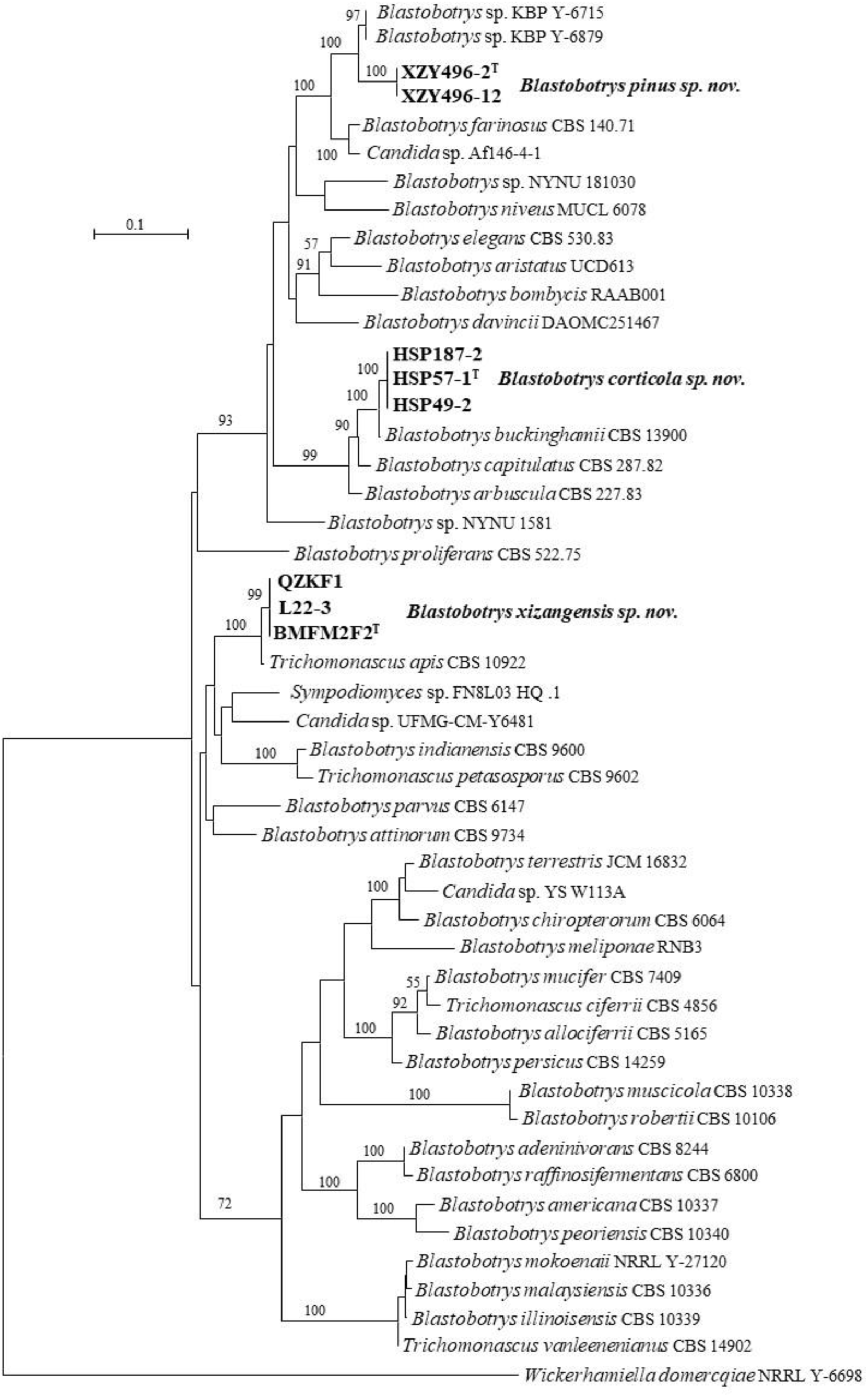
Phylogeny of new taxa (in bold) in the *Blastobotrys* inferred from the combined sequences of the LSU rDNA D1/D2 domains and ITS region (including 5.8S rDNA) by maximum likelihood analysis and over 50 % from 1 000 bootstrap replicates is shown. Bar = 0.1 substitutions per nucleotide position.

Strains HSP49-2, HSP57-1 and HSP187-2 have identical sequences in the ITS region and D1/D2 domains. They have affinity with *Blastobotrys buckinghamii* (Fig. 2), and differed from *B. buckinghamii* by 3 nt in the D1/D2 domains and by 19 nt (3.6 %) ITS region, respectively. The ANI analysis showed that HSP49-2, HSP57-1 and HSP187-2 should represent a distinct species from *B. buckinghamii* due to their ANI value of 84.9 %.

Strains XZY496-2 and XZY496-12 have 1 nt with *Blastobotrys* sp. KBP Y-6715 and KBP Y-6879 in the D1/D2 domains, however, they have 29 nt (6 %) differences in the ITS region, which indicates that they are not conspecific. Those two strains differ from the known species by more than 4.8 % in the D1/D2 domains and by more than 6 % in the ITS region.

Based on the above analysis *Blastobotrys corticola sp. nov.*, *Blastobotrys pinus sp. nov.* and *Blastobotrys xizangensis sp. nov.* were proposed to accommodate groups HSP49-2, XZY496-2 and BMFM2F2, respectively.

##### Diddensiella

Strain XZY423-5 has identical or similar ITS and D1/D2 sequences with *Diddensiella* sp. KBP Y-6906 (OP374141) and Fungal sp. AB11 (FJ235944), *Candida* sp. HA1034 (AF272396), which indicates that they are the same species. They are closely related to *Diddensiella caesifluorescens* and *Diddensiella cacticola* (Fig. 3), and differ from them by more than 17 nt (4 %) in the ITS region and by more than 10 nt (2 %) in the D1/D2 domains. Therefore, we proposed *Diddensiella polypori sp. nov.* to accommodate strains XZY423-5, KBP Y-6906, AB11 and HA1034.

**Fig. 3.**
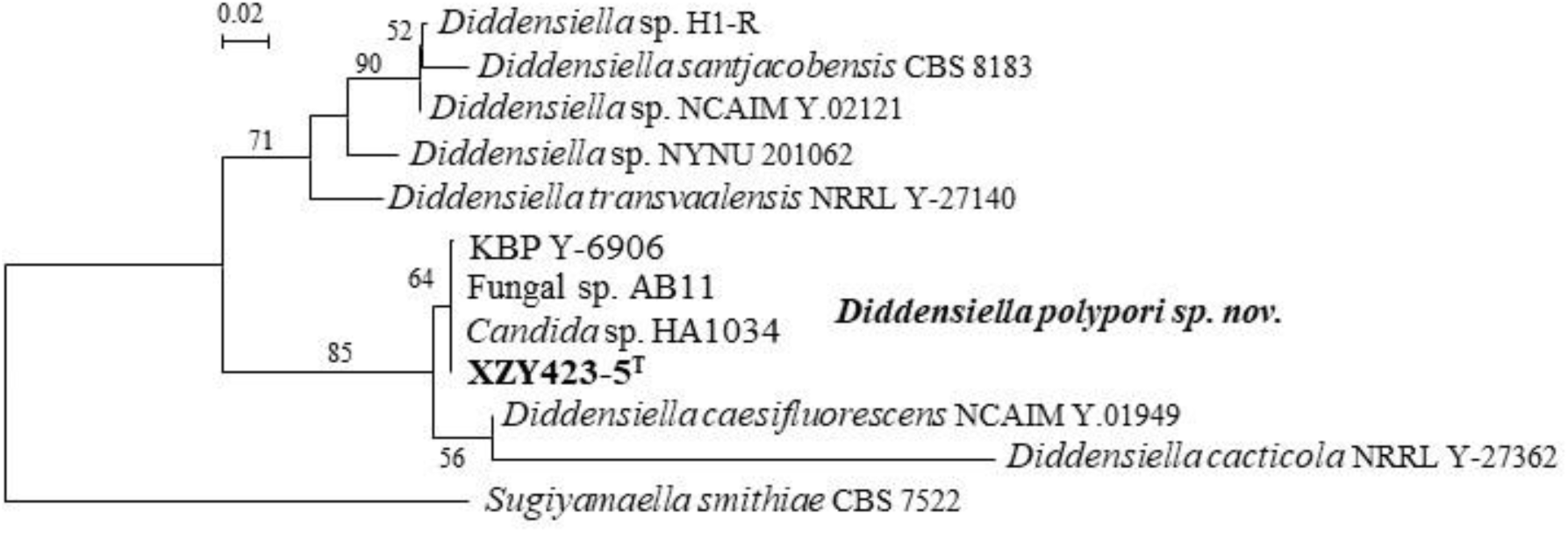
Phylogeny of new taxa (in bold) in the *Diddensiella* inferred from the combined sequences of the LSU rDNA D1/D2 domains and ITS region (including 5.8S rDNA) by maximum likelihood analysis and over 50 % from 1 000 bootstrap replicates is shown. Bar = 0.02 substitutions per nucleotide position.

##### Starmerella

Liu *et al*. (2025) showed that the genus *Starmerella* is heterogeneous and assigned two lineages, namely the *Starmerella* clade 1 and the *Starmerella* clade 2. The *Starmerella* clade 1 has been placed in the reinstated genus *Entelexis*, whereas the *Starmerella* clade 2 is still heterogeneous and was split into four clades and one single-species lineage, namely the *Starmerella sensu stricto* clade, the *Starmerella apicola* clade, the *Starmerella cellae* clade, the *Starmerella stellata* clade and single-species lineage *Starmerella sirachaensis*, based on genome-scale analyses, i.e., the average amino acid identity (AAI) values, the percentage of conserved proteins (POCP) and the presence-absence patterns of orthologs (PAPO). Although the *Starmerella* clade 2 is heterogeneous, Liu *et al*. (2025) suggested that its reclassification would be addressed with more consensus in the yeast community. Here, we followed the suggestion by Liu *et al*. (2025) and assigned three new species in the *Starmerella* clade 2 below.

Five strains, 46YC, CGMCC 2.3767, CGMCC 2.3652, CGMCC 2.3704 and CGMCC 2.3791, are located in the *Starmerella* clade 2. Strain CGMCC 2.3791 is situated in the *Starmerella sensu stricto* clade (Fig. 4) and differs from the known species of *Starmerella* by 4–6.3 % in the D1/D2 domains and by more than 7.4 % in the ITS region. Strains 46YC, CGMCC 2.3767, CGMCC 2.3652 and CGMCC 2.3704, representing two groups, are placed in the *Starmerella apicola* clade (Fig. 4). CGMCC 2.3767 and 46YC have identical sequences with ‘*Starmerella bombi’* UWOPS20-21.1, and they differ from the type strain of *Starmerella bombi* (CBS 5836 =NCYC 2431 =NRRL Y-17081) by 70 nt (15 %) in the ITS region and 4 nt (0.8 %) in the D1/D2 domains. The ANI value of CGMCC 2.3767 and NRRL Y-17081 is 68.8 %, which indicates that CGMCC 2.3767, 46YC and ‘*Starmerella bombi’* UWOPS20-21.1 represent a distinct species from *G. bombi*. Strains CGMCC 2.3652, CGMCC 2.3704 and *Starmerella* sp. FLA45.3 has 0–3 nt difference in the D1/D2 and ITS regions, which indicates that they are conspecific. They differ from the known *Starmerella* species by more than 24 nt (4.8 %) in the D1/D2 domains and by more than 5 % in the ITS region. ‘*Starmerella apicola*’ GK-1 (MH594216) isolated from *Vitis vinifera* in Turkey differs from strains CGMCC 2.3652 and CGMCC 2.3704 by 3 nt in the ITS region, which indicates that they are conspecific.

**Fig. 4.**
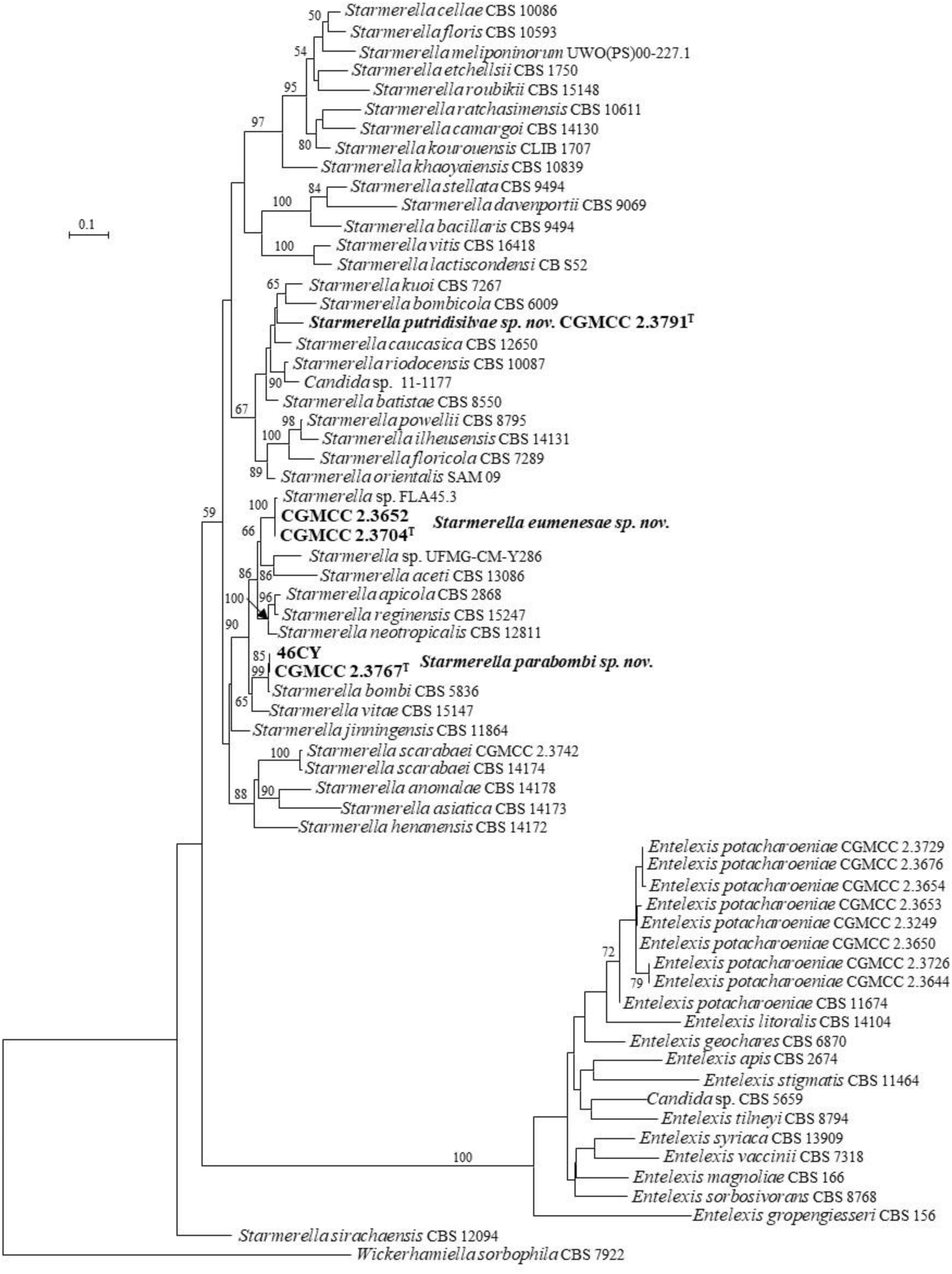
Phylogeny of new taxa (in bold) in the *Starmerella* inferred from the combined sequences of the LSU rDNA D1/D2 domains and ITS region (including 5.8S rDNA) by maximum likelihood analysis and over 50 % from 1 000 bootstrap replicates is shown. Bar = 0.1 substitutions per nucleotide position.

Based on the above analysis, *Starmerella eumenesae sp. nov.*, *Starmerella parabombi sp. nov.* and *Starmerella putridisilvae sp. nov.* are proposed for group CGMCC 2.3704, group CGMCC 2.3767, and CGMCC 2.3791, respectively.

##### Sugiyamaella

Eleven Chinese strains are placed in the *Sugiyamaella* (Fig. 5). CYA2-1, CYA2-2, LXFM2A2, LXFM4F6, LXFM5F3, XZY191-1 and XZY486F1 with identical D1/D2 and ITS sequences are closely related to *Sugiyamaella paludigena* and *Sugiyamaella castrensis* (Fig. 5), and differ from them by 0–2 nt in the D1/D2 domains, but by 12–34 nt (2.6–7.3 %) in the ITS region, which indicates that those seven strains represent a new species.

**Fig. 5.**
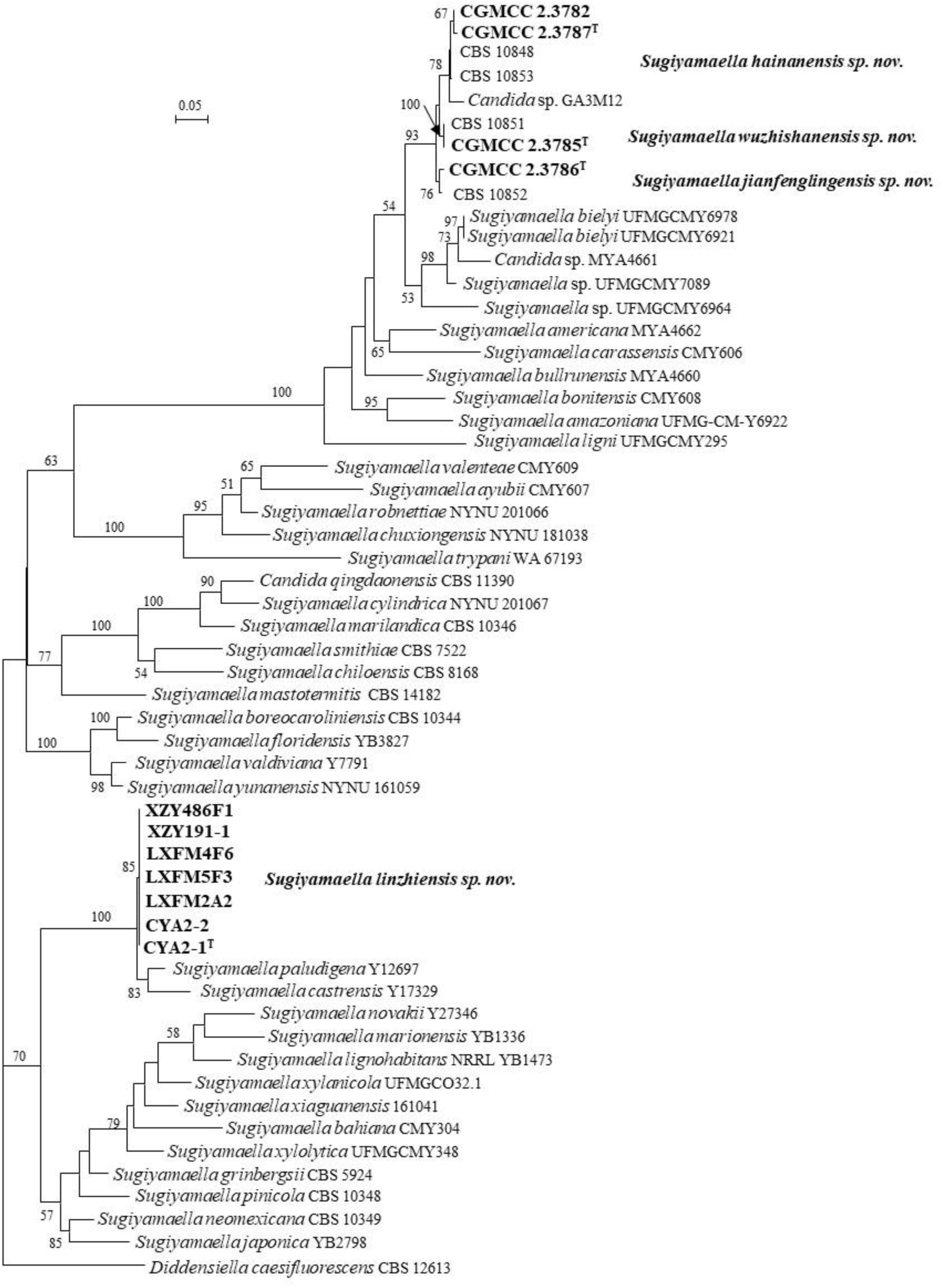
Phylogeny of new taxa (in bold) in the *Sugiyamaella* inferred from the combined sequences of the LSU rDNA D1/D2 domains and ITS region (including 5.8S rDNA) by maximum likelihood analysis and over 50 % from 1 000 bootstrap replicates is shown. Bar = 0.05 substitutions per nucleotide position.

Strains CGMCC 2.3782, CGMCC 2.3785, CGMCC 2.3786, CGMCC 2.3787, CBS 10848 CBS 10851, CBS 10852, and CBS 10853 form three groups represented by CGMCC 2.3782, CGMCC 2.3785 and CGMCC 2.3786, respectively, and cluster with *Sugiyamaella amazoniana*, *Sugiyamaella americana*, *Sugiyamaella bielyi*, *Sugiyamaella bullrunensis*, *Sugiyamaella bonitensis*, *Sugiyamaella carassensis* and *Sugiyamaella ligni* (Fig. 5). Group CGMCC 2.3782 with four strains have same D1/D2 sequences and 1 nt difference in the ITS region. Group CGMCC 2.3786 with two strains has the same D1/D2 sequences and 1 nt difference in the ITS region. Group CGMCC 2.3785 includes CGMCC 2.3785 and CBS 10851 with identical ITS and D1/D2 sequences and differs from group CGMCC 2.3786 by 6 nt in the D1/D2 domains, and 3 nt in the ITS region. Group CGMCC 2.3785 differs from group CGMCC 2.3782 by 9 nt in the D1/D2 domains, and 5 nt in the ITS region. Group CGMCC 2.3786 differed from group CGMCC 2.3782 by 7 nt in the D1/D2 domains, and 7 nt in the ITS region. They differ from the known *Sugiyamaella* species by 28 nt (4.2 %) in the D1/D2 domains, and by more than 11.8 % in the ITS region. The ANI values between groups CGMCC 2.3782, CGMCC 2.3785 and CGMCC 2.3786 are 71.1–83.9 %, which indicates that they represent three distinct species, although they have low ITS and D1/D2 sequence diversity, especially lower ITS sequence differences. *Candida* sp. GA3M12 isolated from the mushroom in Taiwan, China clusters with group CGMCC 2.3782, but differs from them by 3 nt and 11–12 nt (∼2.5 %) in the D1/D2 domains and the ITS region, which indicates that they may be different species.

Based on the above analysis, *Sugiyamaella hainanensis sp. nov.*, *Sugiyamaella jianfenglingensis sp. nov.*, *Sugiyamaella linzhiensis sp. nov.* and *Sugiyamaella wuzhishanensis sp. nov.* are proposed to accommodate groups CGMCC 2.3782, CGMCC 2.3786, CYA2-1 and CGMCC 2.3785, respectively.

##### Wickerhamiella

Three strains are located in the genus *Wickerhamiella* (Fig. 6). CGMCC 2.2799 and XZ69B1 are closely related to *Wickerhamiella cachassae*, *Wickerhamiella dulcicola*, *Wickerhamiella natalensis* and some unpublished strains (Fig. 6). Those two strains differed from them by more than 11 nt (2 %) in the D1/D2 domains and more than 11.8 % in the ITS region. Strain CGMCC 2.3487 is placed in a separate branch in our ITS+D1/D2 tree (Fig. 6). This strain differs from its closely related *Wickerhamiella* species by more than 5.1 % and 9.2 % in the D1/D2 domains and ITS region, respectively. The above rDNA analysis indicates that those three strains represent two new species in *Wickerhamiella*, therefore, *Wickerhamiella follicola sp. nov.* and *Wickerhamiella insecticola sp. nov.* are proposed for them.

**Fig. 6.**
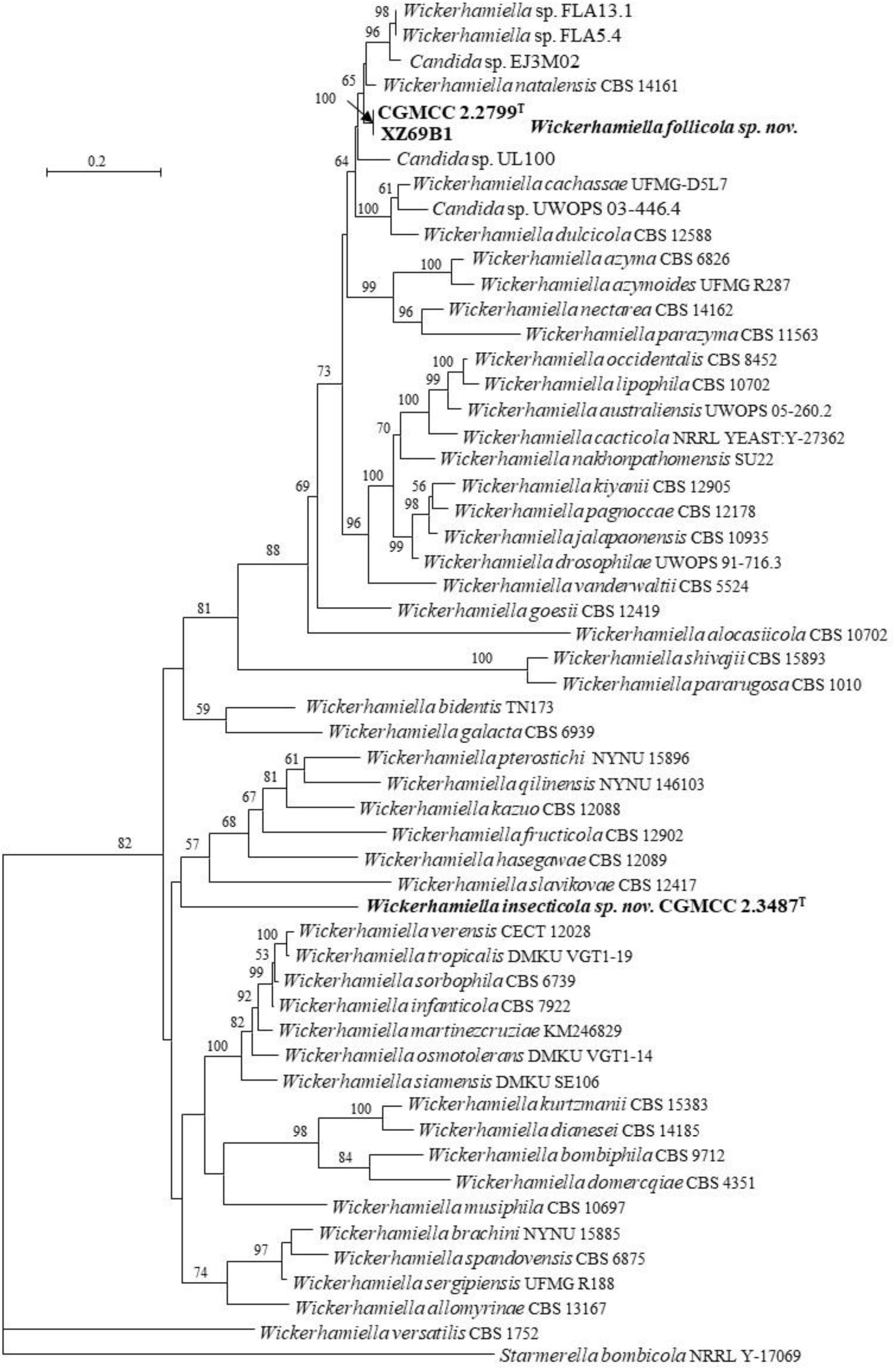
Phylogeny of new taxa (in bold) in the *Wickerhamiella* inferred from the combined sequences of the LSU rDNA D1/D2 domains and ITS region (including 5.8S rDNA) by maximum likelihood analysis and over 50 % from 1 000 bootstrap replicates is shown. Bar = 0.2 substitutions per nucleotide position.

#### New species identification in the Alaninaceae (Alaninales, Pichiomycetes, Saccharomycotina)

##### Nakazawaea

Strains XZY20F1, XZY95-3, XZY137-5, XZY163-1 and XZY163-6 with 0–8 nt ITS sequence difference and identical D1/D2 sequence are closely related to *Nakazawaea ernobii*, *Nakazawaea holstii* and *Nakazawaea tricholomae* (Fig. 7), and differ from them by 0–7 nt in the D1/D2 domains and by 12–25 nt (2–3.9 %) in the ITS region. Strains XZY867-1 and XZY879-1 have 2 nt ITS and 1 nt D1/D2 sequence difference, which are closely related to *Nakazawaea anatomiae*, *Nakazawaea populi* and *Nakazawaea wickerhamii* (Fig. 7), and differ from them by 8–9 nt (1.4–1.7 %) in the D1/D2 domains and by more than 33 nt (4 %) in the ITS region. Based on the above analysis, two new *Nakazawaea* species are proposed to accommodate those groups, namely *Nakazawaea quercae sp. nov.* and *Nakazawaea putredosilvae sp. nov*.

**Fig. 7.**
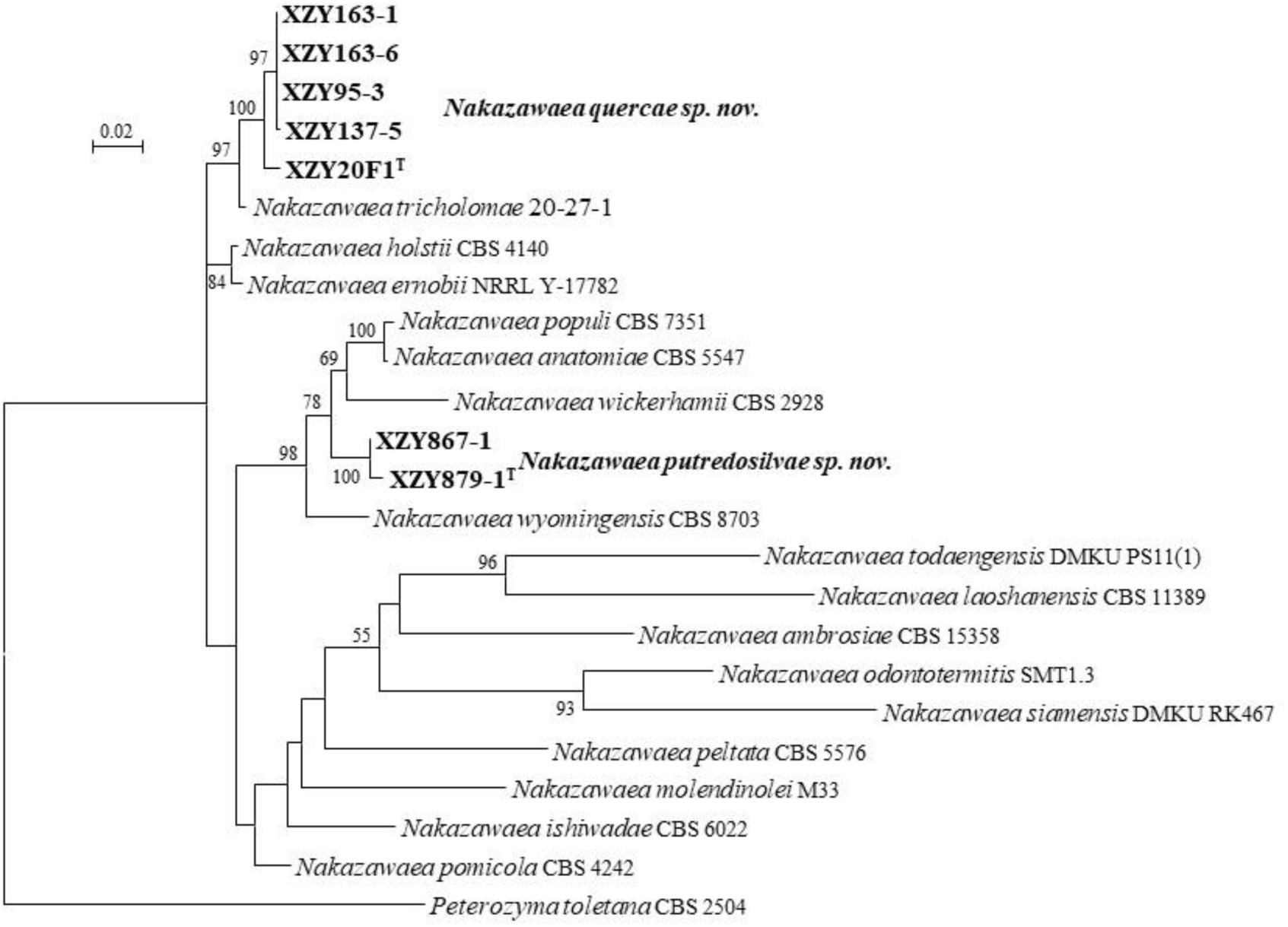
Phylogeny of new taxa (in bold) in the *Nakazawaea* inferred from the combined sequences of the LSU rDNA D1/D2 domains and ITS region (including 5.8S rDNA) by maximum likelihood analysis and over 50 % from 1 000 bootstrap replicates is shown. Bar = 0.02 substitutions per nucleotide position.

#### New species identification in the Pichiaceae (Pichiales, Pichiomycetes, Saccharomycotina)

##### Komagataella

Strains HSP262-1, HSP262-2 and HSP275-3 form a clade that is located in the *Komagataella,* which is more closely related to *Komagataella pastoris* (Fig. 8), and differs from it by four nt in the D1/D2 domains, but by more than 6.8 % in the ITS region. Those three strains have a 94.7 % ANI value with *K. pastoris*. The above rDNA sequence analysis and ANI comparison showed that those three strains represent one distinct species of *Komagataella*. Therefore, *Komagataella corticis sp. nov.* was proposed to accommodate those three strains.

**Fig. 8.**
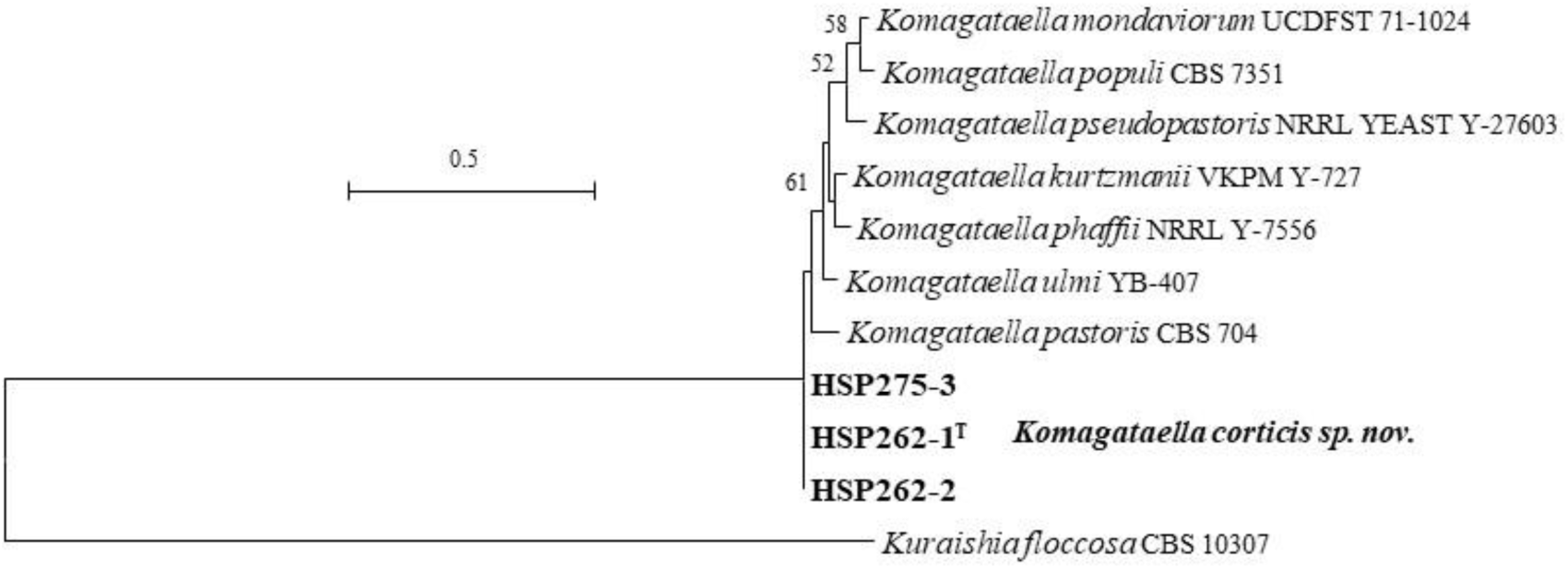
Phylogeny of new taxa (in bold) in the *Komagataella* inferred from the combined sequences of the LSU rDNA D1/D2 domains and ITS region (including 5.8S rDNA) by maximum likelihood analysis and over 50 % from 1 000 bootstrap replicates is shown. Bar = 0.5 substitutions per nucleotide position.

##### Ogataea

Liu *et al*. (2025) demonstrated that the genus *Ogataea* is polyphyletic based on the phylogenomic analysis, and genome-based metrics, i.e., AAI, POCP and PAPO. Two large clades, namely the *Ogataea* clade 1 and the *Ogataea* clade 2, were assigned to the genus *Ogataea*. The *Ogataea* clade 2 was phylogenetically separated from the *Ogataea* clade 1 contains the type species of *Ogataea*, and was placed in a newly described genus *Wenyingozyma* (Liu *et al*. 2025). Although the *Ogataea* clade 1 is still heterogeneous based on the genome-based metric analysis, Liu *et al*. (2025) argued that the reclassification of the *Ogataea* clade 1 needs more robust analyses to address their taxonomic conclusions in the future. Therefore, we also follow the above treatment and assigned two new species in the *Ogataea sensu stricto* subclade and *Ogataea pilisensis* subclade.

Strains XZY267-1 and XZY775-1 are closely related to *Ogataea paradorogensis*, *Ogataea xyloterini* and an unpublished strain *Cyberlindnera* sp. GG3L01 in the *Ogataea sensu stricto* subclade (Fig. 9), and differ from them by 8–10 nt (1.3–1.4 %) in the D1/D2 domains and 17–23 nt (2–2.2 %) in the ITS region. XZY166-1 and XZY167-1 have an affinity to *Ogataea nitratoaversa*, *Ogataea pilisensis*, and *Candida piceae* in the *Ogataea pilisensis* subclade, and differ from them by 6–8 nt (1.2–1.6 %) in the D1/D2 domains and 54–79 nt (7.4–10.7 %) in the ITS region. Therefore, *Ogataea linzhienesis sp. nov.* is proposed to accommodate XZY267-1 and XZY775-1 and *Ogataea caprifoliacea sp. nov.* is created to include XZY166-1 and XZY167-1.

**Fig. 9.**
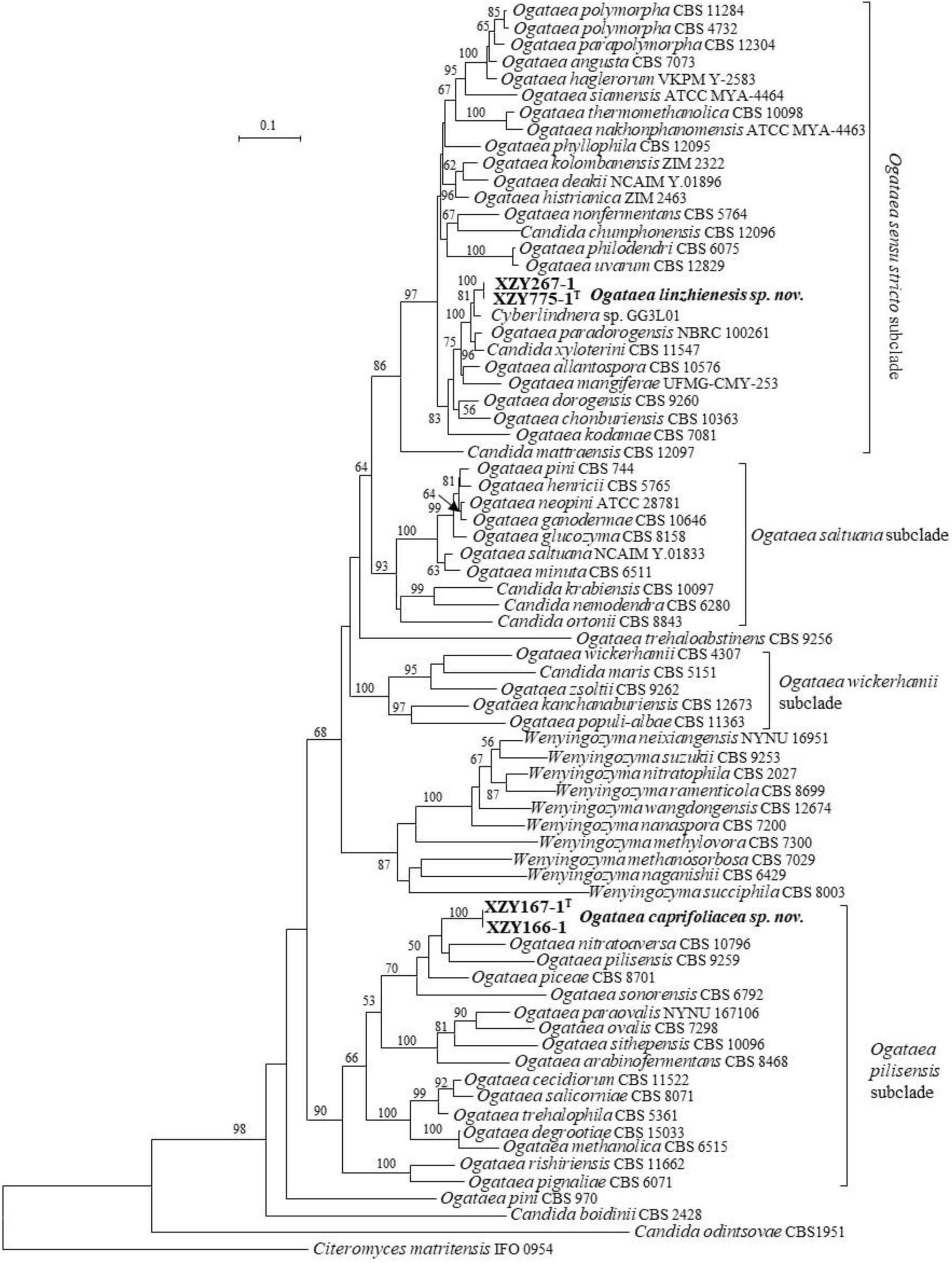
Phylogeny of new taxa (in bold) in the *Ogataea* inferred from the combined sequences of the LSU rDNA D1/D2 domains and ITS region (including 5.8S rDNA) by maximum likelihood analysis and over 50 % from 1 000 bootstrap replicates is shown. Bar = 0.1 substitutions per nucleotide position.

##### Pichia

Nine strains are located in the genus *Pichia*, and assigned to five groups based on the analysis of the ITS and D1/D2 sequences (Fig. 10). Group XZ1989-1 includes four strains isolated from Tibet, namely XZ18-36F1A, XZ186-3, XZ1989-1 and XZ1989-2, have 0 to 2 nt different ITS and D1/D2 sequences. They are closely related to *Pichia pseudolambica* (Fig. 10), and differ from it by 3 nt in the D1/D2 domains, but by 19–20 nt (4.3 %) in the ITS region.

**Fig. 10.**
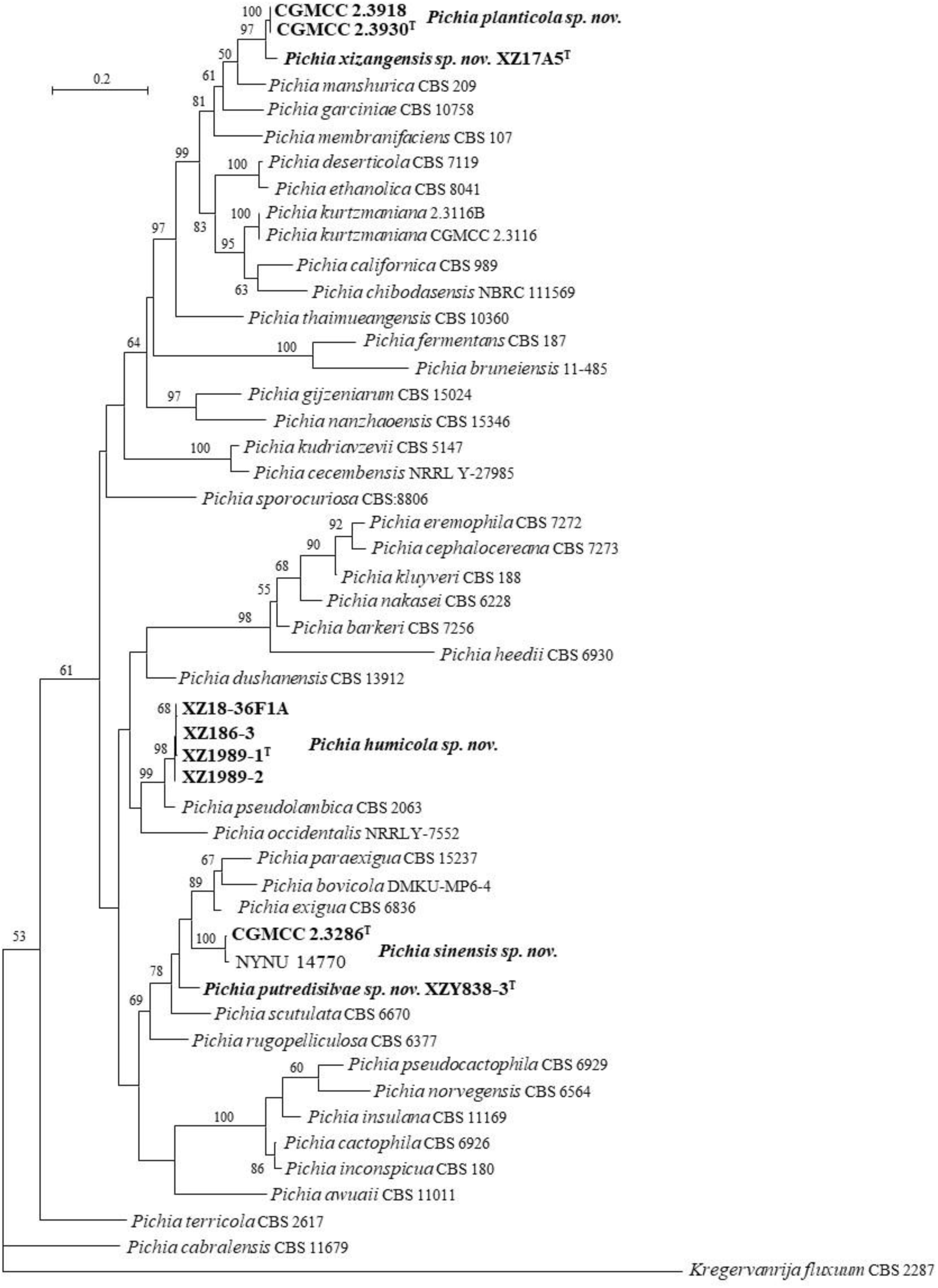
Phylogeny of new taxa (in bold) in the *Pichia* inferred from the combined sequences of the LSU rDNA D1/D2 domains and ITS region (including 5.8S rDNA) by maximum likelihood analysis and over 50 % from 1 000 bootstrap replicates is shown. Bar = 0.2 substitutions per nucleotide position.

Group CGMCC 2.3930 and group XZ17A5 have identical D1D2 sequences, but the latter differs from the former by 14 nt (3.1 %) in the ITS region, which indicates that they may be different species. Those three strains differ from the known species *Pichia garciniae*, *Pichia manshurica* and *Pichia membranifaciens* by 6–25 nt (1–4.1 %) in the D1/D2 domains, and by 47–66 nt (10–14.2 %) in the ITS region. The ANI value between CGMCC 2.3930 and XZ17A5 is 76.2 %, which supports the separation of those strains as two species.

Groups CGMCC 2.3286 and XZY838-3 differ from each other by 27 nt (4.4 %) in the D1/D2 domains and by 44 nt (9.4 %) in the ITS region. They are closely related to *Pichia bovicola*, *Pichia exigua*, *Pichia paraexigua* and *Pichia scutulata*, and differ from them by 22–35 nt (4–6.1 %) in the D1/D2 domains, by more than 8 % in the ITS region. CGMCC 2.3286 and *Pichia* sp. NYNU 14770 (KM598659) have identical D1/D2 sequences and 1 nt difference in the ITS region, which indicates that they are conspecific. CGMCC 2.3286 has an identical D1/D2 sequence with *Candida* sp. JW3-2 (KU316732), *Candida* sp. HU19CS (LC602868), *Candida* sp. HU20CS (LC602869) and *Pichia* sp. NRRL Y-12827 (EF550245), which indicates that they may be the same species. XZY838-3 and *Pichia* sp. UASWS2914 (ON428565) isolated from the bark of *Vitis vinifera* in Switzerland have 99.3% similarity in the D1/D2 domains, which indicates that they may be conspecific.

Based on the above analysis, five new *Pichia* species, namely *Pichia humicola sp. nov.* for group XZ1989-1, *Pichia planticola sp. nov.* for group CGMCC 2.3930, *Pichia putredosilvae sp. nov.* for group XZY838-3, *Pichia sinensis sp. nov.* for group CGMCC 2.3286 and *Pichia xizangensis sp. nov.* for group XZ17A5, are proposed to accommodate those five groups.

##### Saturnispora

Three strains are placed in the genus *Saturnispora* based on the ITS and D1/D2 sequence analysis. CGMCC 2.3783 and XZJF10-2 have identical D1/D2 sequences and 1 nt different ITS sequences. They are closely related to *Saturnispora zaruensis* and two unpublished strains *Saturnispora* sp. GY16S01 and *Saturnispora* sp. ES3M03 (Fig. 11), and differ from them by 3 nt in the D1/D2 domains, but by 11–14 nt (2.4–6 %) in the ITS region, which indicates that CGMCC 2.3783 and XZJF10-2 represent a new species of *Saturnispora*. Strain XZ18-24-2 and two unpublished strains *Saturnispora* sp. NYNU 14639 and *Candida* sp. ES13S05 have 0–1 nt difference in the D1/D2 domains but differs from each other by 19–20 nt (4.8–5 %) in the ITS region, which indicates that they are not conspecific. Strain XZ18-24-2 differs from the known *Saturnispora* species by more than 12 nt (2.5 %) in the D1/D2 domains and more than 3.8 % in the ITS region. Based on the above analysis, *Saturnispora jianfenglingensis sp. nov.* and *Saturnispora linzhiensis sp. nov.* are proposed to accommodate CGMCC 2.3783 and XZJF10-2, and XZ18-24-2, respectively.

**Fig. 11.**
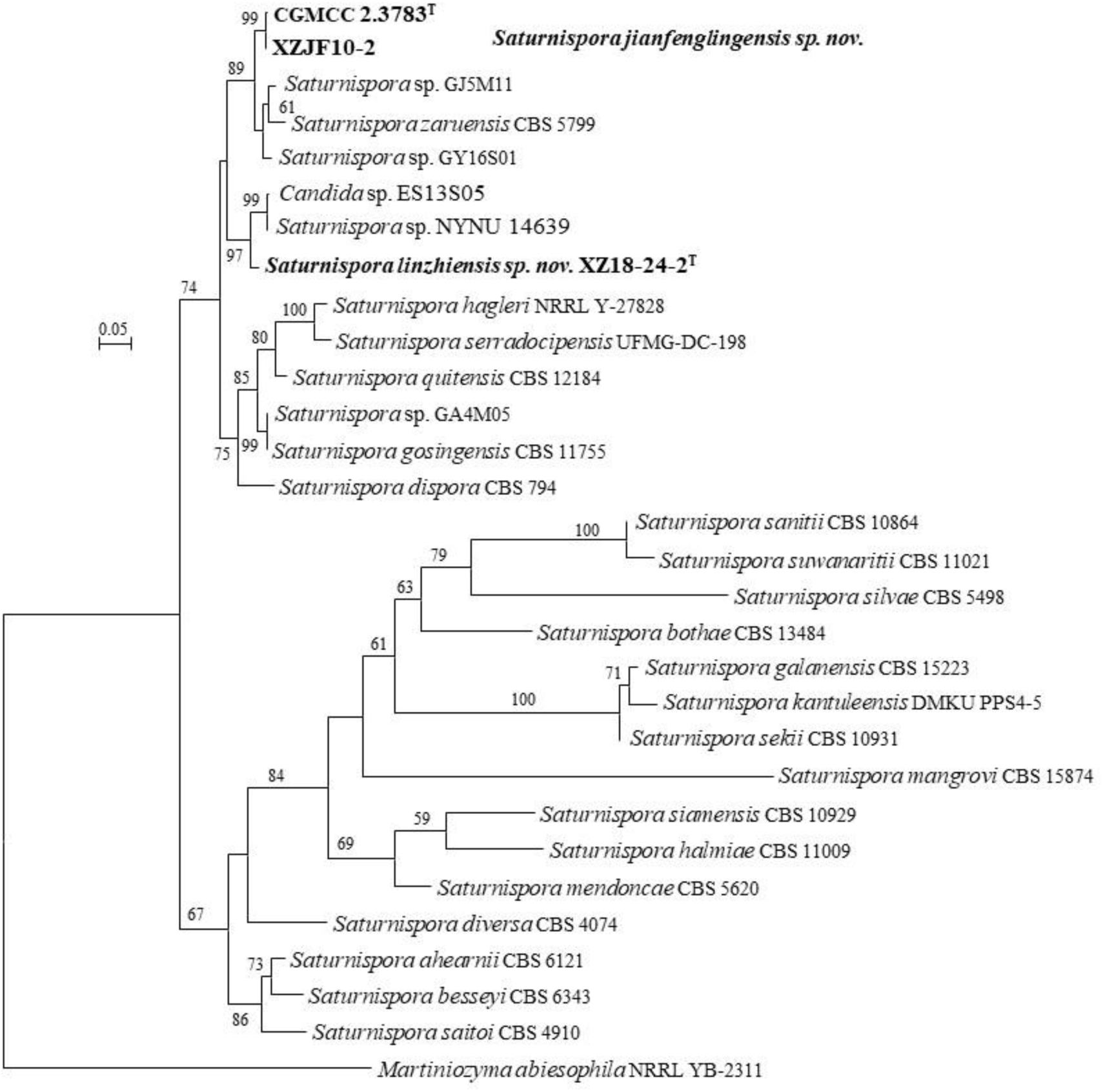
Phylogeny of new taxa (in bold) in the *Saturnispora* inferred from the combined sequences of the LSU rDNA D1/D2 domains and ITS region (including 5.8S rDNA) by maximum likelihood analysis and over 50 % from 1 000 bootstrap replicates is shown. Bar = 0.05 substitutions per nucleotide position.

##### Xiuguozyma

The species *Candida silvatica* has an unstable phylogenetic position based on different gene analyses (Kurtzman & Robnett 1998, Sugita & Nakase 1999, Lachance *et al*. 2011), but possible affinities with the genus *Dekkera* (*Brettanomyces*) (Kurtzman & Robnett 1998, Sugita & Nakase 1999). The ITS+D1/D2 phylogenetic analysis showed *C. sylvatica* located in a long separate branch closely related to *Brettanomyces* (Liu *et al*. 2025). The phylogenomic analyses from two studies showed that *C. silvatica* clustered with *Candida insectalens* and formed a well-supported *Candida insectalens* clade, which is closely related to the genus *Brettanomyces* (Opulente *et al*. 2024, Liu *et al*. 2025). Based on the phylogenomic analysis and the AAI, POCP and PAPO genome-based metric analyses, the *Candida insectalens* clade was assigned to the newly proposed genus *Xiuguozyma* (Liu *et al*. 2025).

Strains CGMCC 2.3792 and G2WZ06-1 have the same ITS and one nt different D1/D2 sequences. They are closely related to *Xiuguozyma sylvatica* (formerly *C. silvatica*) and an unpublished strain *Candida* sp. JCM 16747 (AB552927) isolated from the beetle gallery in Japan in the ITS+D1/D2 rDNA analysis with 97% bootstrap support (Fig. 12), and differ from them by 34–38 nt (6.2–6.7 %) in the D1/D2 domains and 75 nt (15 %) in the ITS region. *Xiuguozyma insectalens* (formerly *C. insectalens*) formed a long branch without bootstrap support with *X. silvatica* and CGMCC 2.3792, G2WZ06-1 and JCM 16747. Considering the above analyses, we propose a new species *Xiuguozyma putredisilvae sp. nov.* to accommodate two Chinese strains. JCM 16747 seems to represent the fourth member of *Xiuguozyma* because it differs from *X. silvatica* by 24 nt (4 %) in the D1/D2 domains.

**Fig. 12.**
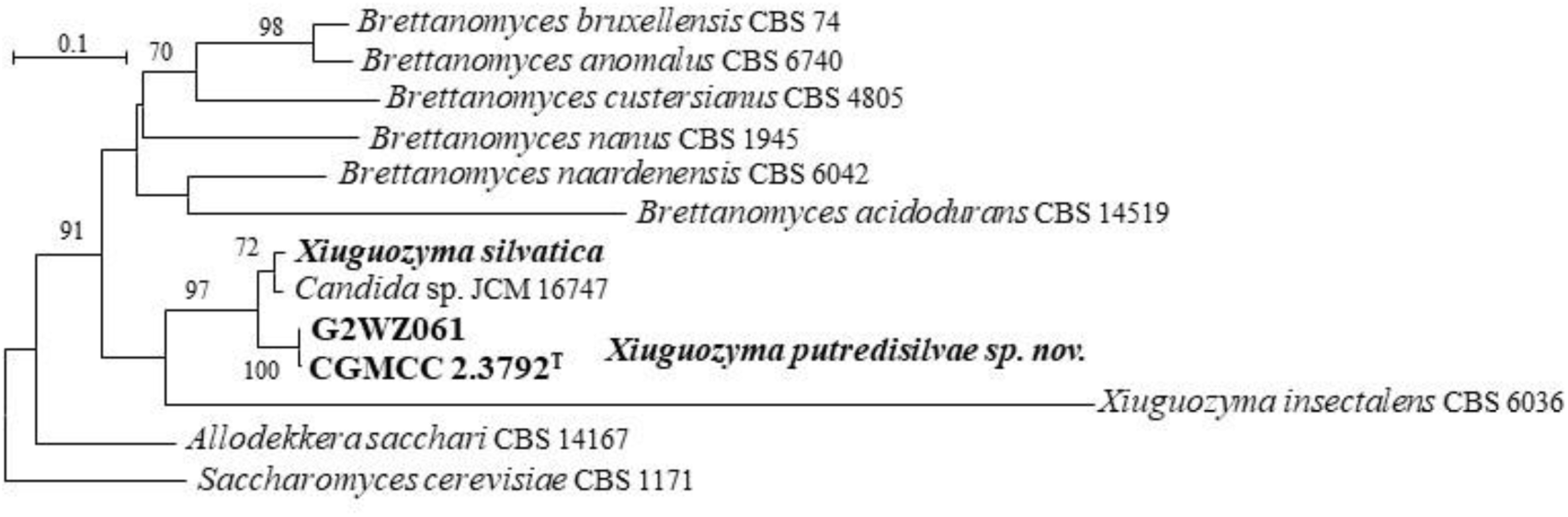
Phylogeny of new taxa (in bold) in the *Xiuguozyma* inferred from the combined sequences of the LSU rDNA D1/D2 domains and ITS region (including 5.8S rDNA) by maximum likelihood analysis and over 50 % from 1 000 bootstrap replicates is shown. Bar = 0.1 substitutions per nucleotide position.

#### New species identification in the Debaryomycetaceae (Serinales, Pichiomycetes, Saccharomycotina)

##### Chernovozyma

The Blast of the ITS and D1/D2 sequences against the NCBI nucleotide database showed that gmt3-3-4 and gmt3-3-5 have affinity with *Candida sophiae-reginae*. They differ from each other by 16 nt (2.7 %) in the D1/D2 domains and 55 nt (9 %) in the ITS region. gmt3-3-4 and gmt3-3-5 have 4 nt ITS and 6 nt D1/D2 sequence difference with *Candida sp.* EJ7S08 (HQ623544/EF653945), which indicates that they may be conspecific. The phylogenomic analysis in Liu *et al*. (2025) and the ITS+D1/D2 phylogenetic analysis in this study (Fig. 13) showed that gmt3-3-4 is located in the *Candida aurita* clade. Liu *et al*. (2025) assigned this clade to a new genus *Chernovozyma*. Therefore, we proposed *Chernovozyma guizhouensis sp. nov.* to accommodate gmt3-3-4 and gmt3-3-5. The strain *Schwanniomyces* sp. yHMH407 isolated from the USA has the same ITS and D1/D2 sequence as strain gmt3-3-4, which indicates that *Chernovozyma guizhouensis* occurs outside of China.

**Fig. 13.**
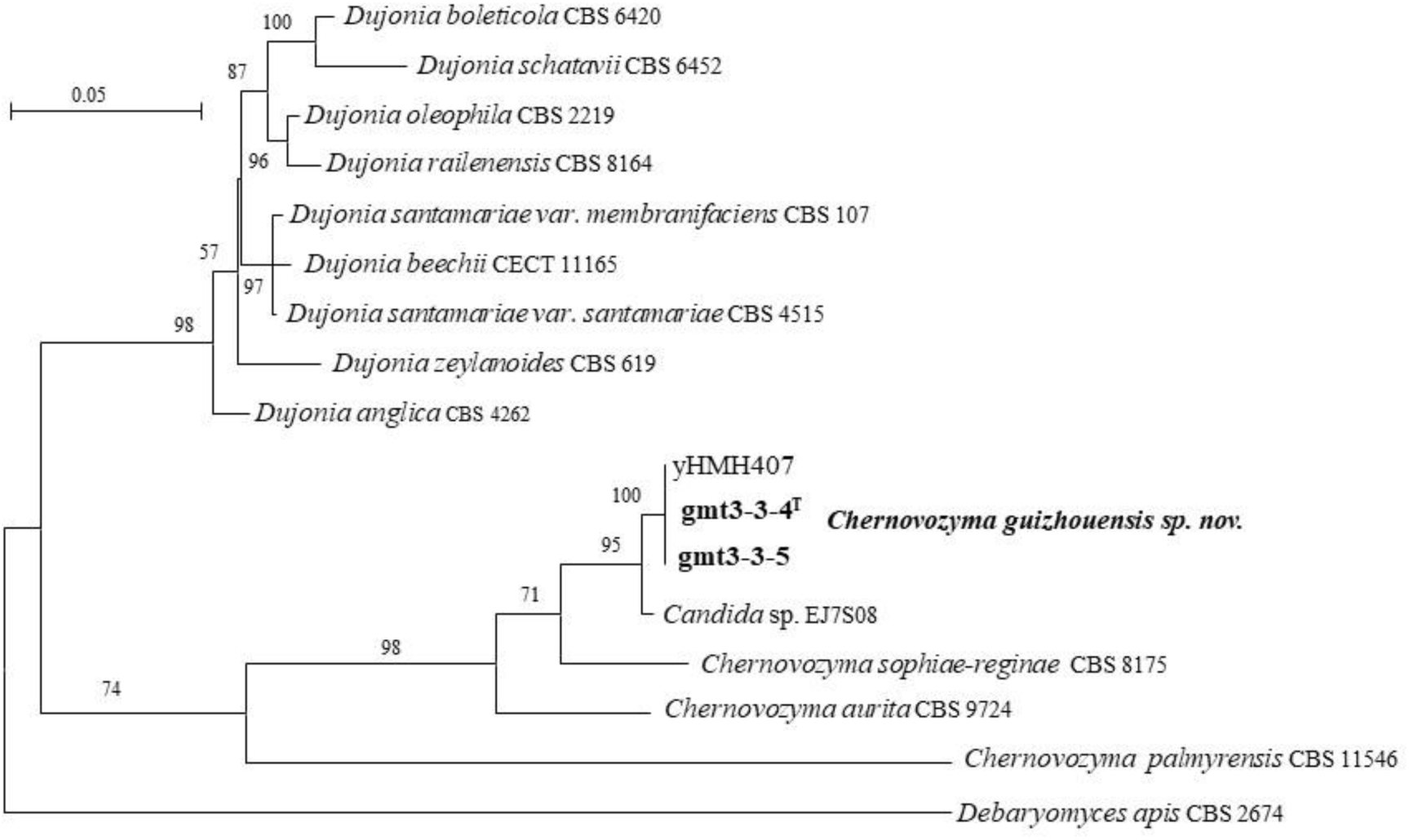
Phylogeny of new taxa (in bold) in the *Chernovozyma* inferred from the combined sequences of the LSU rDNA D1/D2 domains and ITS region (including 5.8S rDNA) by maximum likelihood analysis and over 50 % from 1 000 bootstrap replicates is shown. Bar = 0.05 substitutions per nucleotide position.

##### Hemisphaericaspora

Strain CGMCC 2.3790 and four strains YM28381, YM28382, YM28383 and YM283831 isolated from the rhizosphere and bulk soil of *Magnolia sinica* in China have the same ITS and D1/D2 sequence. They are closely related to *Spathaspora elongata* and *Spathaspora mengyangensis* (Fig. 14), and differ from them by 14–17 nt (2.7–3.3 %) and 30 nt (6.3 %) in the D1/D2 domains and the ITS region, respectively. Liu *et al*. (2025) showed that *S. elongata* and *S. mengyangensis* are located in the *Hemisphaericaspora* clade and separated from the other *Spathaspora* species, therefore, Liu *et al*. (2025) transferred those *Spathaspora* and *Candida* species in the *Hemisphaericaspora* clade to *Hemisphaericaspora.* The above analysis indicates that strains CGMCC 2.3790, YM28381, YM28382, YM28383 and YM283831 represent a new species of *Hemisphaericaspora*, for which the name *Hemisphaericaspora corticola sp. nov.* is proposed.

**Fig. 14.**
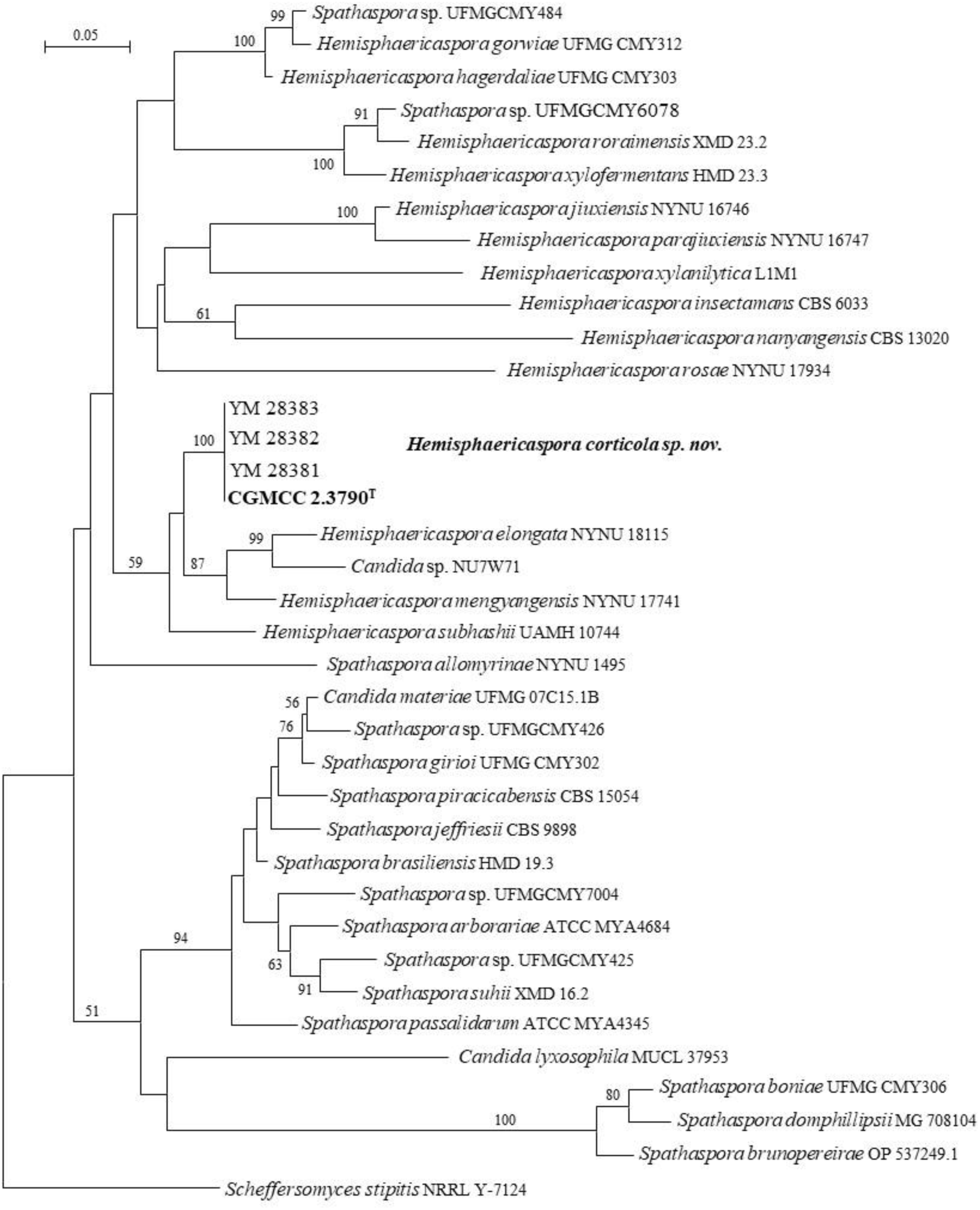
Phylogeny of new taxa (in bold) in the *Hemisphaericaspora* inferred from the combined sequences of the LSU rDNA D1/D2 domains and ITS region (including 5.8S rDNA) by maximum likelihood analysis and over 50 % from 1 000 bootstrap replicates is shown. Bar = 0.05 substitutions per nucleotide position.

##### Insectozyma

Liu *et al*. (2025) recently reclassified *Candida* species in the *Saccharomycotina* and placed the *Candida corydali* clade into a new genus *Insectozyma*. Strains CGMCC 2.3227, CGMCC 2.3277, CGMCC 2.3494 and CGMCC 2.3733 are closely related to *Insectozyma corydali* (Fig. 15), and differ from it by 10 nt (1.7 %) and 8 nt (1.5 %) in the D1/D2 domains and ITS region, respectively. They have 0–2 nt with 15 strains isolated from Taiwan, China. i.e., 09LN10 (HM222592/HM222581), 09LN7 (HM222593/HM222582), HSG6-1 (HM222591/HM222580), SY3L08 (EU523624/FJ153161), HJLES3-1 (HM222599/HM222588), GE12L01 (FJ527067), TY1-4 (HM222578), OW1-8 (HM222574), RB3-6 (HM222579), GB3-1 (HM222573), OJLET1-1 (HM222598/HM222587), HSG6-1 (HM222591/HM222580), HRB2-2 (HM222575), HDSEE2-1 (HM222600/HM222589), ODSEE1-2 (HM222597/HM222586), and CdVSA83.1 (MW437793) and CdVSA81.1 (MW437790) isolated from the floral nectar in South Africa and *Metschnikowia* sp. A08023-1-4 (AB998452) isolated from the stem of *Lilium lancifolium* in Koreain the D1/D2 domains or ITS region (Table 1). The above data indicated that this new species had a worldwide distribution. Therefore, the name *Insectozyma pseudocorydali sp. nov.* is proposed for them.

**Fig. 15.**
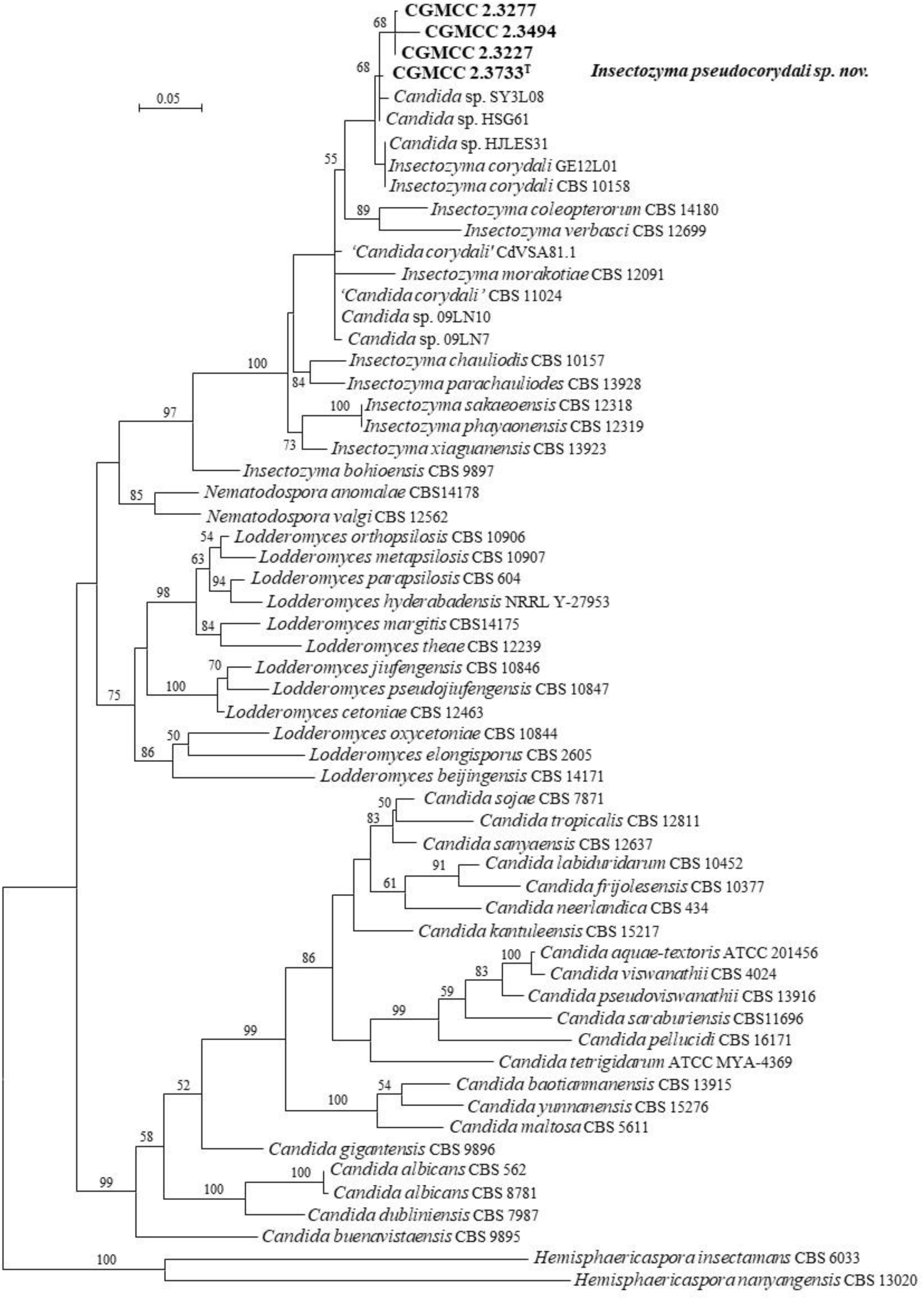
Phylogeny of new taxa (in bold) in the *Insectozyma* inferred from the combined sequences of the LSU rDNA D1/D2 domains and ITS region (including 5.8S rDNA) by maximum likelihood analysis and over 50 % from 1 000 bootstrap replicates is shown. Bar = 0.05 substitutions per nucleotide position.

##### Suhomyces

Twelve strains isolated from Tibet form four groups representing XZY98-4, XZY189-1, XZY145-10 and XZFM12-1, respectively, in the genus *Suhomyces* and separate from the known *Suhomyces* species (Fig. 16). Group XZY98-4 includes four strains, which have identical ITS and D1/D2 sequences. They have 3 nt with *Candida* sp. EN13M07 in the D1/D2 domains, but 37 nt (10 %) in the ITS region. They differ from the other known species by more than 6 % in the D1/D2 domains and more than 14% in the ITS region. Group XZY189-1 contains two strains with one nt difference in D1/D2 sequences and identical ITS sequences. Group XZY145-10 is composed of three strains having identical D1/D2 sequences and 2 nt different ITS sequences. Those two groups are closely related and differ from each other by 4 nt in the D1/D2 domains and 11 nt (2.5 %) in the ITS region. The ANI value between groups XZY189-1 and XZY145-10 is 74.8 %, which supports the separation of those two groups. Group XZY189-1 has 2 nt in the D1/D2 domains with *Candida* sp. CBS 7170, which indicates that they may be conspecific. Groups XZY189-1 and XZY145-10 have more than 7 % and 10 % sequence diversity from the other known species in the D1/D2 domains and the ITS region, respectively.

**Fig. 16.**
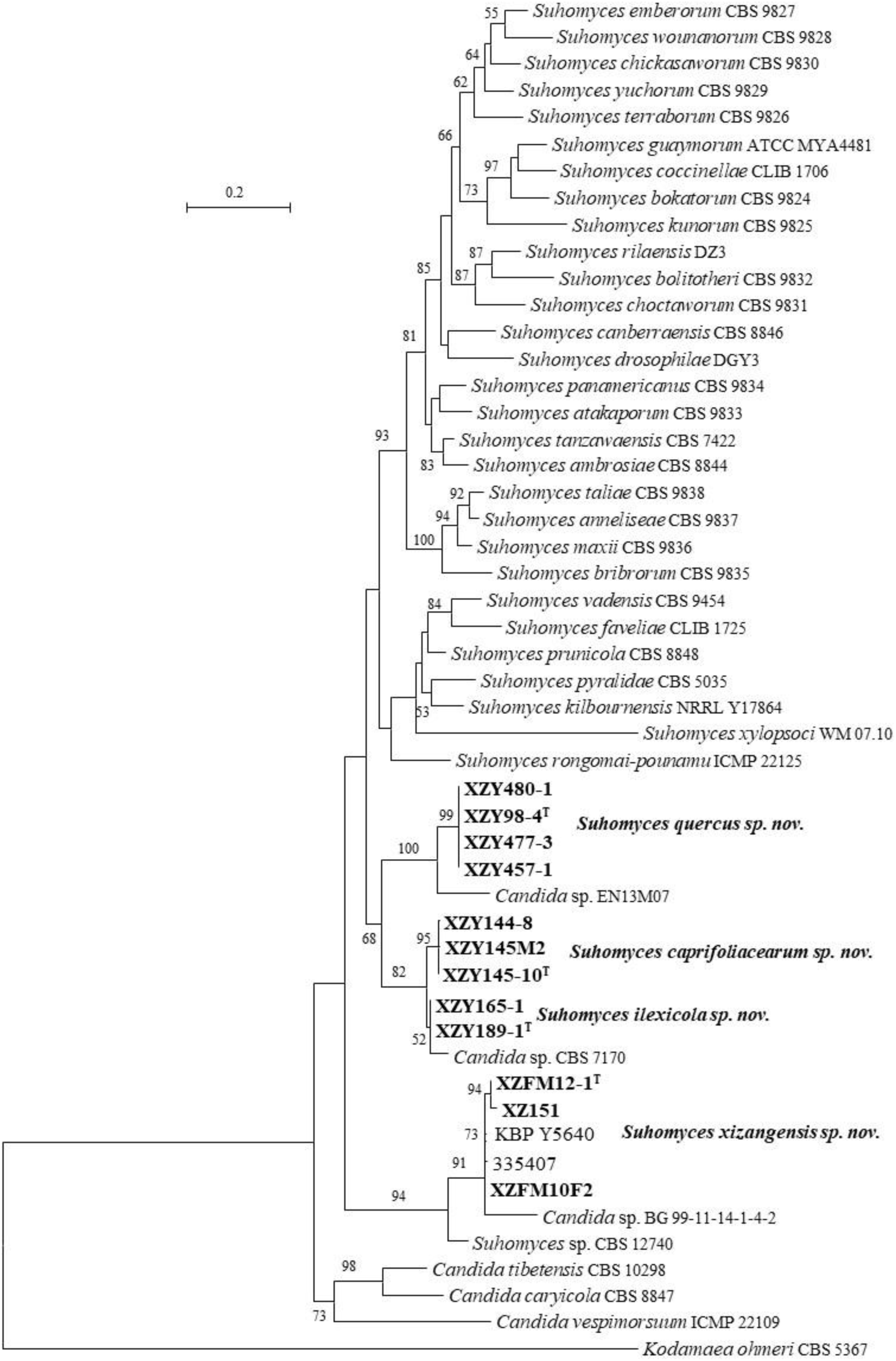
Phylogeny of new taxa (in bold) in the *Suhomyces* inferred from the combined sequences of the LSU rDNA D1/D2 domains and ITS region (including 5.8S rDNA) by maximum likelihood analysis and over 50 % from 1 000 bootstrap replicates is shown. Bar = 0.2 substitutions per nucleotide position.

Group XZFM12-1 contains three strains with 2–6 nt ITS sequence differences. They have 0–1 nt in the D1/D2 domains or 2–7 nt in the ITS region with fifteen strains, namely KBP Y-5640 (OQ383492), 335407 (MT340077), QZJ-1 (MW710061), BG99-11-14-1-4-2 (MH745214), GY44S04 (FJ527151), B26-10-3 (MW764676), WM 07.70 (FM178359/EU343859), yHQL452 (OK052083), yHQL448 (OK052078), yHQL450 (OK052081), yHQL323 (OK051966), HBP0722E4-2 (PQ568191), HBP0139A4AB (PQ568150), HBP0139A3AB (PQ568149) and CBS 12740 (JX656740/JX656741). Those strains form a separate branch from the other known *Suhomyces* species and may be conspecific (Fig. 16 and Fig. S2).

Based on the above analysis, four *Suhomyces* new species, namely *Suhomyces ilexicola sp. nov.* for group XZY189-1, *Suhomyces caprifoliacearum sp. nov.* for group XZY145-10, *Suhomyces quercus sp. nov.* for group XZY98-4 and *Suhomyces xizangensis sp. nov.* for group XZFM12-1, are proposed.

##### Yamadazyma

Liu *et al*. (2025) demonstrated that the genus *Yamadazyma* was heterogeneous and split it into four clades, namely *Yamadazyma sensu stricto* clade*, Yamadazyma epiphylla* clade, *Yamadazyma olivae* clade, and *Yamadazyma triangularis* clade, based on the genome-scale analyses. Liu *et al*. (2025) transferred *Candida* species in the *Yamadazyma sensu stricto* clade to *Yamadazyma*, whereas they placed *Candida* species in the *Yamadazyma olivae* clade as *pro tempore* in *Candida* and suggested reclassifying the *Yamadazyma epiphylla* clade, *Yamadazyma olivae* clade, and *Yamadazyma triangularis* clade after discussing in the yeast taxonomist’s community. Therefore, we currently follow this suggestion below when we propose the new species in the genus *Yamadazyma*.

Nine strains form four groups in the *Yamadazyma sensu stricto* clade (Fig. 17). Strains CGMCC 2.3789 and CGMCC 2.3793 are closely related to *Yamadazyma ubonensis*, *Yamadazyma phyllophila* and *Yamadazyma paraphyllophila*. They differ from the three known species by 5–10 nt (1–1.7 %) in the D1/D2 domains, and 9–22 nt (1.5–3.5 %) in the ITS region. Strain CGMCC 2.3482 forms a clade with *Yamadazyma laniorum* and *Yamadazyma tenuis* (Fig. 17). It differs from the two known species by more than 11 nt (2 %) in the D1/D2 domains and by more than 27 nt (4.5 %) in the ITS region. Strains CGMCC 2.3642, CGMCC 2.3657, CGMCC 2.3714 and CGMCC 2.3716 form a clade separated from the known species of *Yamadazyma* (Fig. 17), and differ from them by more than 10–11 nt (1.7–2 %) in the D1/D2 domains and more than 34 nt (5.3 %) in the ITS region. Strains LF2-1 and LF3-2 with 1 nt different D1/D2 sequence and identical ITS sequence are closely related to *Yamadazyma insectorum*, *Yamadazyma riverae* and *Candida* sp. UFMG-CM-Y6947 (Fig. 17). Those two Chinese strains differed from them by 8– 13 nt (1.4–2.3 %) in the D1/D2 domains, and more than 7 % in the ITS region. The above analyses indicate that those four groups represent four new *Yamadazyma* species, for which the names *Yamadazyma chuxiongensis sp. nov.*, *Yamadazyma follicola sp. nov.*, *Yamadazyma putredisilvae sp. nov.* and *Yamadazyma sympetra sp. nov.* are proposed to accommodate the groups LF3-2, CGMCC 2.3482, CGMCC 2.3789 and CGMCC 2.3642, respectively.

**Fig. 17.**
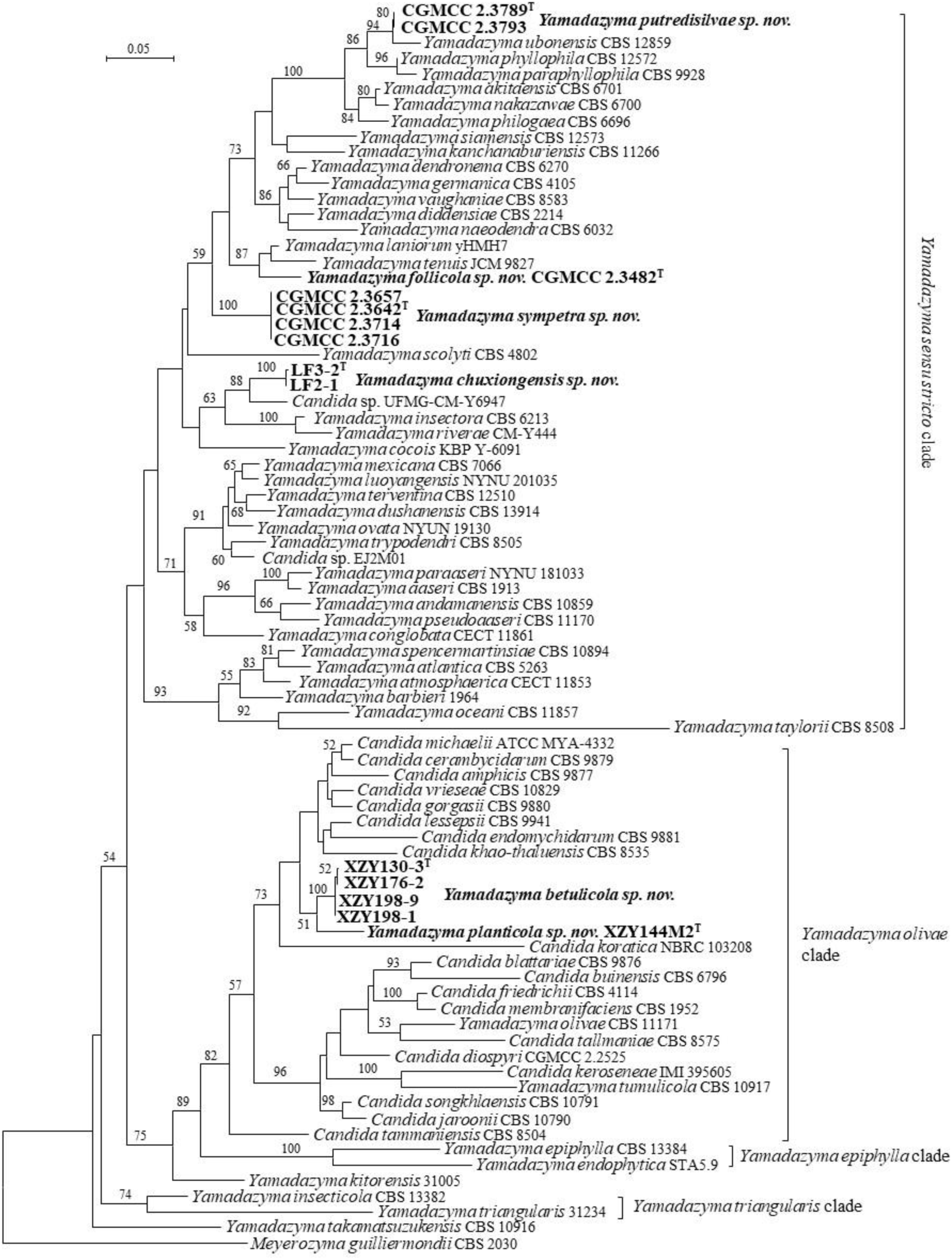
Phylogeny of new taxa (in bold) in the*Yamadazyma* inferred from the combined sequences of the LSU rDNA D1/D2 domains and ITS region (including 5.8S rDNA) by maximum likelihood analysis and over 50 % from 1 000 bootstrap replicates is shown. Bar = 0.05 substitutions per nucleotide position.

XZY144M2, XZY198-1, XZY198-9, XZY130-3 and XZY176-2 form two groups in the *Yamadazyma olivae* clade (Fig. 17). Group XZY130-3 has four strains with identical D1/D2 sequences and 1 nt different ITS sequence, which indicates that they are conspecific. Group XZY144M2 differs from group XZY198-1 by 2 nt in the D1/D2 domains, but by 20 nt (3.3%) in the ITS region. The ANI analysis showed that groups XZY144M2 and XZY130-3 have 76.2 % ANI value, which demonstrates that those two groups represent two different species. Group XZY144M2 and XZY130-3 are closely related to *Candida amphicis*, *Candida cerambycidarum*, *Candida endomychidarum*, *Candida gorgasii*, *Candida khao-thaluensis*, *Candida koratica*, *Candida lessepsii*, *Candida michaelii* and *Candida vrieseae* (Fig. 17), and differed from them by 3–9 nt (0.5–1.5 %) in the D1/D2 domains, and more than 4 % in the ITS region. Based on the above analysis, we will propose *Yamadazyma betulicola sp. nov.* and *Yamadazyma planticola sp. nov.* to accommodate groups XZY130-3 and XZY144M2, respectively.

#### New species identification in the Metschnikowiaceae (Serinales, Pichiomycetes, Saccharomycotina)

##### Australozyma

Liu *et al*. (2024b) reclassified the *Candida* species and related taxa in the *Metschikowiaceae* and proposed 13 new genera. Seven *Candida* species and one *Metschnikowia* species were transferred into the newly described genus *Australozyma* (Liu *et al*. 2024b). Two strains, LF17-2 and LF117-1, isolated from Yunnan Province, are placed in the *Australozyma* (Fig. 18). They are closely related to *Australozyma picinguabensis*, *Australozyma tocantinsensis* and *Australozyma saccharicola* and differ from those known species by 23–30 nt (11.6–15.1%) in the D1/D2 domains, and more than 60 nt (11 %) in the ITS region. LF17-2 and LF117-1 have identical D1/D2 sequences and 2 nt different ITS sequences, which indicates that they are conspecific. Therefore, *Australozyma ficea sp. nov.* is proposed to accommodate LF17-2 and LF117-1.

**Fig. 18.**
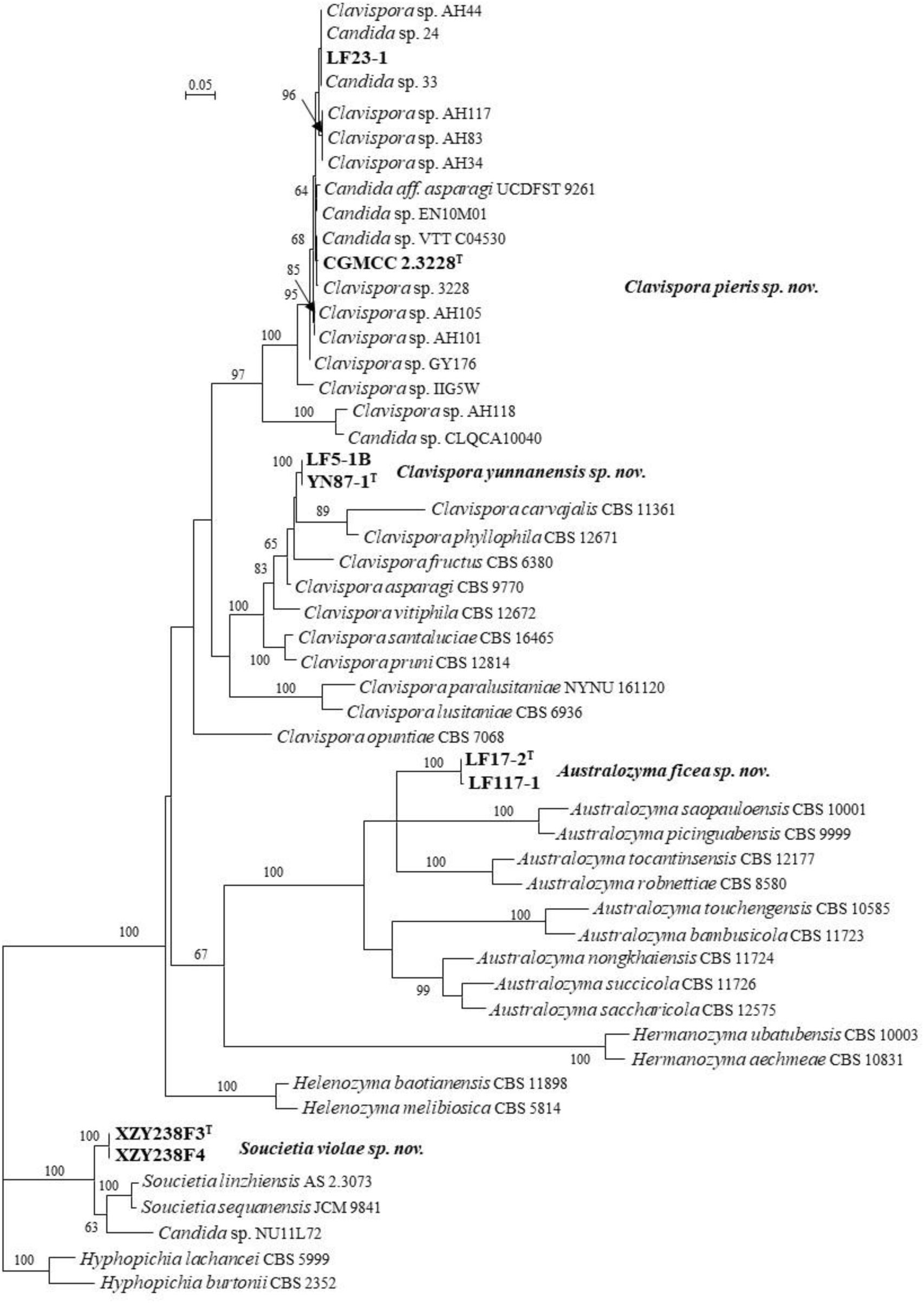
Phylogeny of new taxa (in bold) in the *Australozyma*, *Clavispora* and *Soucietia* inferred from the combined sequences of the LSU rDNA D1/D2 domains and ITS region (including 5.8S rDNA) by maximum likelihood analysis and over 50 % from 1 000 bootstrap replicates is shown. Bar = 0.05 substitutions per nucleotide position.

##### Clavispora

Strains YN87-1 and LF5-1B have an affinity with *Clavispora asparagi*, *Clavispora carvajalis*, *Clavispora fructus* and *Clavispora phyllophila* (Fig. 18), and differ from them by more than 8 nt (1.4 %) in the D1/D2 domains and more than 11 nt (3.3 %) in the ITS region, which indicates that those two strains represent a new species. Therefore, *Clavispora yunnanensis sp. nov.* is proposed to accommodate them. The two strains have 4 nt D1/D2 sequence differences with *Clavispora* sp. UFMG-CM-Y6393 (MH348140), which indicates that the taxonomic position of them needs to be confirmed in the future based on more sequence analysis.

LF23-1 and CGMCC 2.3228 have the same D1/D2 sequence and differ from each other by 11 nt in the ITS region, however, the ANI value between LF23-1 and CGMCC 2.3228 is 98.7 %, which indicates that they are conspecific. LF23-1 and CGMCC 2.3228 differ from seven Japanese strains *Clavispora* sp. AH44 (LC813545) from honey bee, AH34 (LC813544), AH83 (LC813546), AH101 (LC813547), AH105 (LC813548) and AH117 (LC813549) from fruit fly and GY176 (LC813551) from corn, *Clavispora* sp. IIG5W (MN708001) from the insect gut in India, *Candida* sp. 24 (MN385703) and *Candida* sp. 33 (MN385710) from *Megaplatypus mutatus* in Argentina, *Clavispora* sp. 3228 (PQ314504) from *Pieris rapae* in Hunan province, China, *Candida* sp. EN10M01 (FJ873485) from the mushroom and *Candida aff. asparagi* UCDFST:92-61 (MH595211) from a rotten bamboo shoot of *Phyllostachys pubescens* in Taiwan, China, *Candida* sp. VTT C04530 (DQ377643) from the industrial malting ecosystem in Finland, by 1–5 nt in the D1/D2 domains and 5–10 nt in the ITS region. Those 14 strains differ from the known *Clavispora* species by more than 12 % in the D1/D2 domains and more than 15 % in the ITS region. The above analysis indicated that LF23-1 and CGMCC 2.3228 and those unpublished strains represent new *Clavispora* species, for which *Clavispora pieris sp. nov.* was proposed for them.

##### Metschnikowia

Forty strains are located in the genus *Metschnikowia* (Table 1, Fig. 19). Among them, twenty-three strains, namely CGMCC 2.2651, CGMCC 2.2800, CGMCC 2.3430, CGMCC 2.3437, XZY3-3, XZY3-4, XZY3F2, XZY14A, XZY19-5, XZY30-5, XZY87, XZY257-1, XZY257-2-1, XZY279-2, XZY30-2, XZY40A, XZY279-3, XZY19-6, XZY39-4, XZY3-2, XZY39-2, XZY30 and XZY40-1 formed two groups representing by CGMCC 2.3430 and XZY30-5, respectively. Groups CGMCC 2.3430 and XZY30-5 differ from each other by 4–5 nt in the D1/D2 domain. They cluster with *Metschnikowia cibodasensis*, *Metschnikowia reukaufii*, *Metschnikowia vanudenii* and *Candida magnifica*. They differ from those known species by 4–7 nt in the D1/D2 domains and by 9–16 nt in the ITS region.

**Fig. 19.**
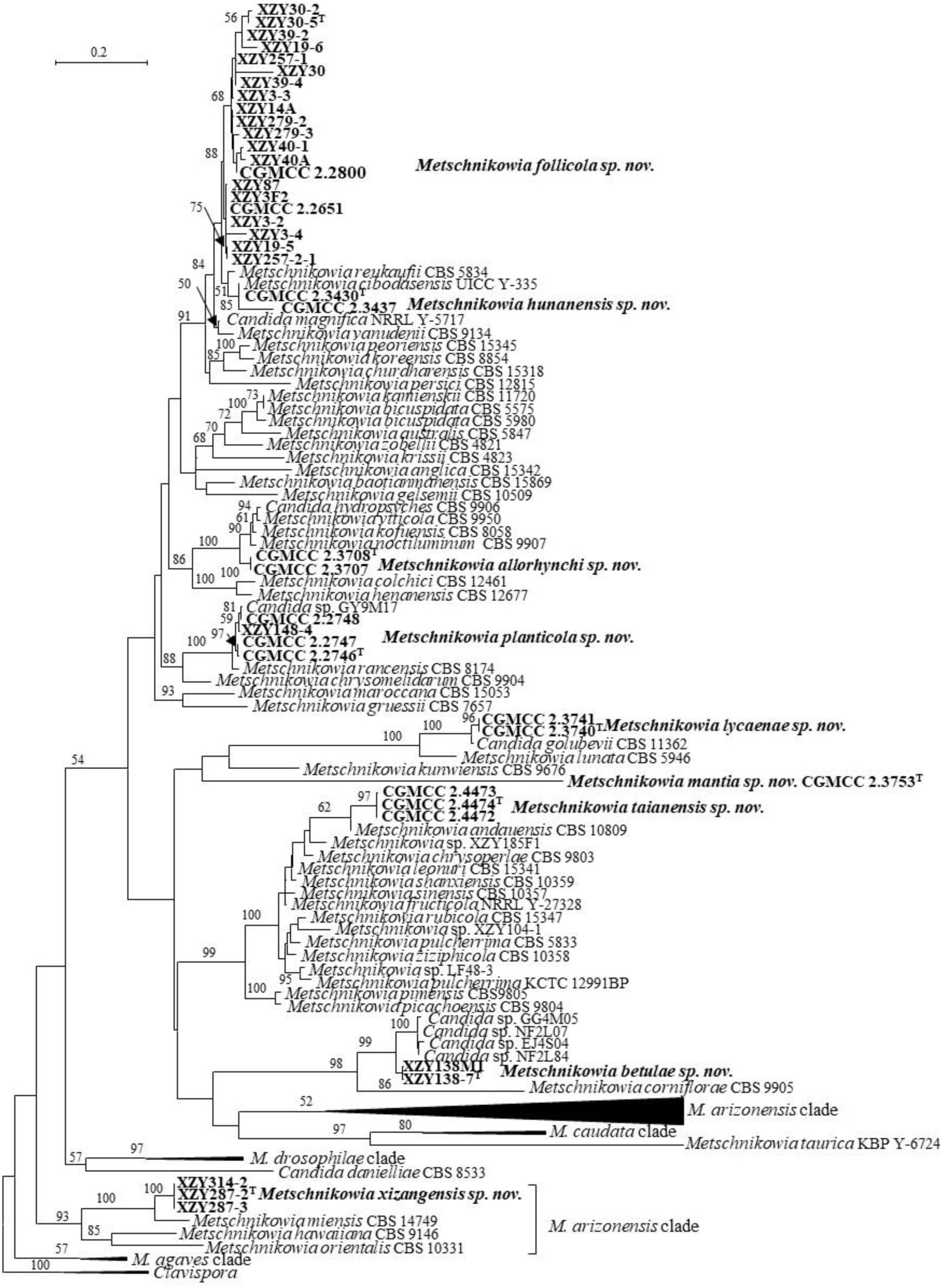
Phylogeny of new taxa (in bold) in the *Metschnikowia* inferred from the combined sequences of the LSU rDNA D1/D2 domains and ITS region (including 5.8S rDNA) by maximum likelihood analysis and over 50 % from 1 000 bootstrap replicates is shown. Bar = 0.2 substitutions per nucleotide position.

CGMCC 2.3707 and CGMCC 2.3708 with identical ITS and D1/D2 sequences are closely related to *Metschnikowia kofuensis*, *Metschnikowia noctiluminum*, *Metschnikowia viticola* and *Candida hydropsyches*, and differ from them by 10–13 nt (2.6–3.3 %) in the ITS region, and by 16 nt (2.8 %) in the D1/D2 domains.

CGMCC 2.2746, CGMCC 2.2747, CGMCC 2.2748 and XZY148-4 are closely related to *Metschnikowia rancensis* and *Metschnikowia chrysomelidarum*, and differ from them by 7–15 nt in the ITS region and by 6–15 nt in the D1/D2 domains. CGMCC 2.2746 and CGMCC 2.2747 have the same D1/D2 and 1nt different ITS sequences, and differ from CGMCC 2.2748 and XZY148-4 by 7-8 in the ITS region, and by 3–5 nt in the D1/D2 domains. CGMCC 2.2746, CGMCC 2.2747, CGMCC 2.2748 and XZY148-4 differed from *Candida* sp. GY9M17 (HM461581/FJ873539) by 4–8 nt in the ITS region, and by 0–5 nt in the D1/D2 domains, which indicates that they may be conspecific.

Strains CGMCC 2.3740 and CGMCC 2.3741 have identical D1/D2 and ITS sequences. They have an affinity to *Candida golubevii* and *Metschnikowia lunata* by more than 57 nt (10.8%) in the D1/D2 domains and by more than 11.5% diversity in the ITS region.

The D1/D2 sequences blast result showed that CGMCC 2.3753 is more closely related to *Metschnikowia shanxiensis* and *Candida gelsemii*, and differs from them by more than 100 nt (8 %) in the D1/D2 domains. But this strain is placed in a separate long branch closely related to *Metschnikowia kunwiensis* without bootstrap support in the *Metschnikowiaceae* (Fig. 19).

CGMCC 2.4472, CGMCC 2.4473 and CGMCC 2.4474 with identical ITS and D1/D2 sequences, and differ from *Metschnikowia andauensis*, *Metschnikowia chrysoperlae, Metschnikowia leonuri*, *Metschnikowia shanxiensis* and *Metschnikowia sinensis* by 6–19 nt (1.3–2.7 %) in the D1/D2 domains, and by 26-28 nt (∼9.3 %) in the ITS region.

XZY138-7 and XZY138M1 have identical D1/D2 sequences and two nt different ITS sequences. They are closely related to four unpublished strains *Candida* sp. NF2L84, NF2L07, EJ4S04, GG4M05 and differ from them by 8 nt in the D1/D2 domains, and by 22-23 nt in the ITS region. They differ from *Metschnikowia corniflorae* by 40 nt (10.3 %) in the ITS region, and by more than 45 nt (11%) in the D1/D2 domains.

XZY287-2, XZY287-3 and XZY314-2 have an affinity with *Metschnikowia miensis*, *Metschnikowia hawaiiana* and *Metschnikowia orientalis*, and differ from them by 30–41 nt (7.6–10.5 %) in the ITS region and by 39–92 nt (7–15.3 %) in the D1/D2 domains.

Based on the above analysis, we will propose 9 *Metschnikowia* new species to accommodate those 40 strains isolated from different regions in China (see details in the Taxonomy section).

##### Soucietia

Strains XZY238F3 and XZY238F4 are clustered with *Soucietia sequanensis*, *Soucietia linzhiensis* and *Candida* sp. NU11L72 (Fig. 18). Those strains differ from the two known species by more than 29 nt (5 %) in the D1/D2 domains, and more than 9 % in the ITS region. Therefore, they represent a new species of *Soucietia*, for which *Soucietia violae sp. nov.* is proposed to accommodate XZY238F3 and XZY238F4.

#### New species identification in the Saccharomycopsidaceae (Ascoideales, Saccharomycetes, Saccharomycotina)

##### Saccharomycopsis

Strain CGMCC 2.2749 has an identical D1/D2 sequence with *Candida* sp. 91 (JN581104), which indicates that they may be the same species. CGMCC 2.2749 is located in the *Saccharomycopsis* (Fig. 20) and differs from *Saccharomycopsis fodiens* by 1 nt in the D1/D2 domains and by 9 nt (1.5 %) in the ITS region. The ANI analysis showed that CGMCC 2.2749 and *S. fodiens* have a 71.9 % ANI value, which indicates that they are different species. Therefore, *Saccharomycopsis parafodiens sp. nov.* is proposed for CGMCC 2.2749.

**Fig. 20.**
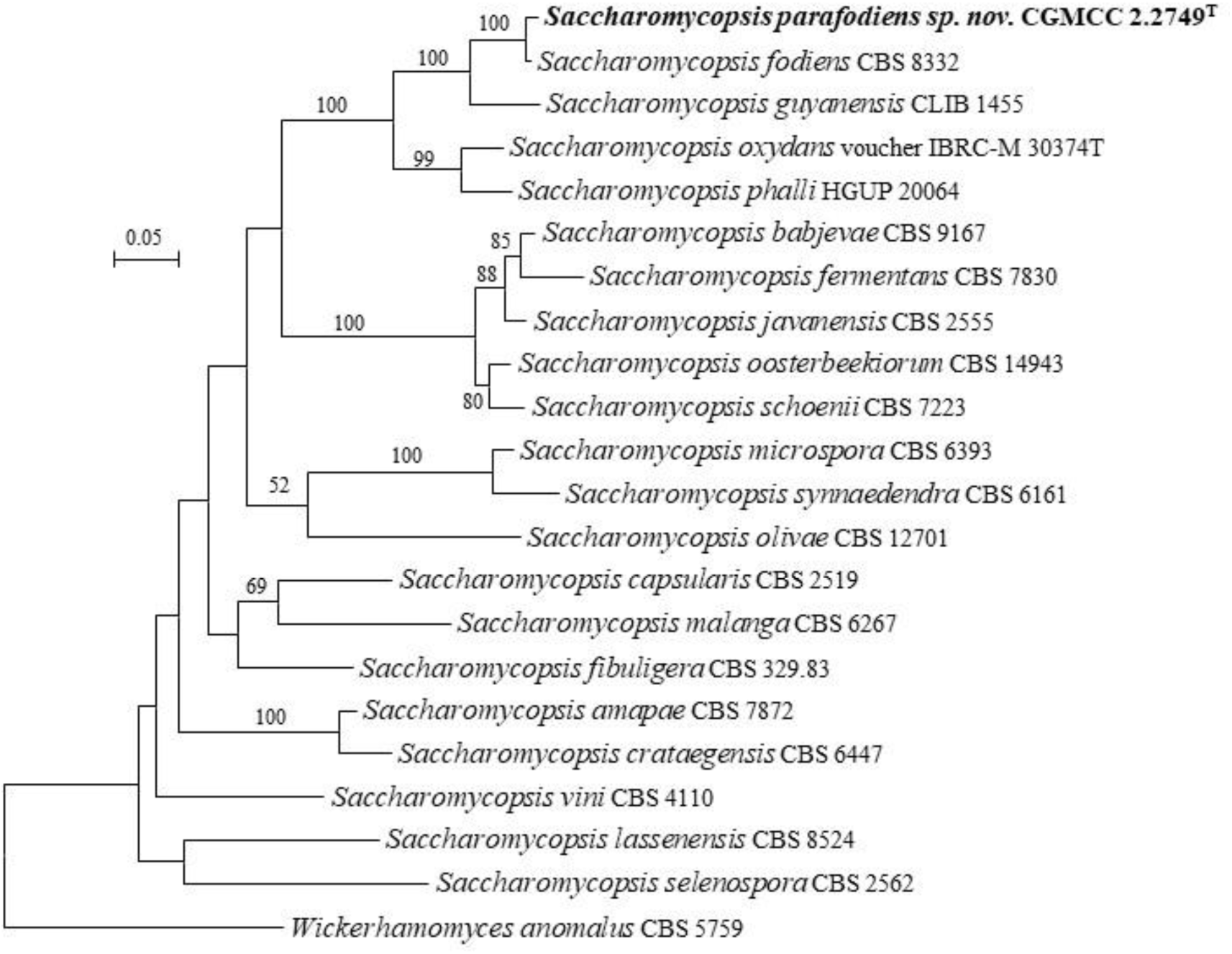
Phylogeny of new taxa (in bold) in the *Saccharomycopsis* inferred from the combined sequences of the LSU rDNA D1/D2 domains and ITS region (including 5.8S rDNA) by maximum likelihood analysis and over 50 % from 1 000 bootstrap replicates is shown. Bar = 0.05 substitutions per nucleotide position.

#### New species identification in the Phaffomycetaceae (Phaffomycetales, Saccharomycetes, Saccharomycotina)

##### Petasospora

Strains Y1-6 and Y1-5 have the same sequences in the D/D2 domain, but there is a 1 nt difference in the ITS regions. They are closely related to *Petasospora japonica*, *Petasospora xylosilytica*, *Petasospora maesa* and *Petasospora easanensis* (Fig. 21), and differ from them by 9–12 nt (2 %) in the D1/D2 domains, and more than 51 nt (8 %) in the ITS region. Strain CGMCC 2.4335 has identical ITS and D1/D2 sequences with *Candida* sp. NRRL Y-7615 (EF550331), *Candida* sp. Af110-2-1 (LC661425/LC661390), *Candida* sp. NU8S73 (JQ901889/HM461716) and Fungal sp. isolate F_L_Jcae8 (OP537660), which indicates that they are conspecific. Those strains are closely related to *Petasospora maritima*, and differ from it by 5 nt and 15 nt (3 %) in the D1D2 domain and ITS region, respectively. Based on the above analysis *Petasospora soli sp. nov.* for Y1-6 and Y1-5 and *Petasospora putridisilvae sp. nov.* for CGMCC 2.4335 are proposed to accommodate those two groups.

**Fig. 21.**
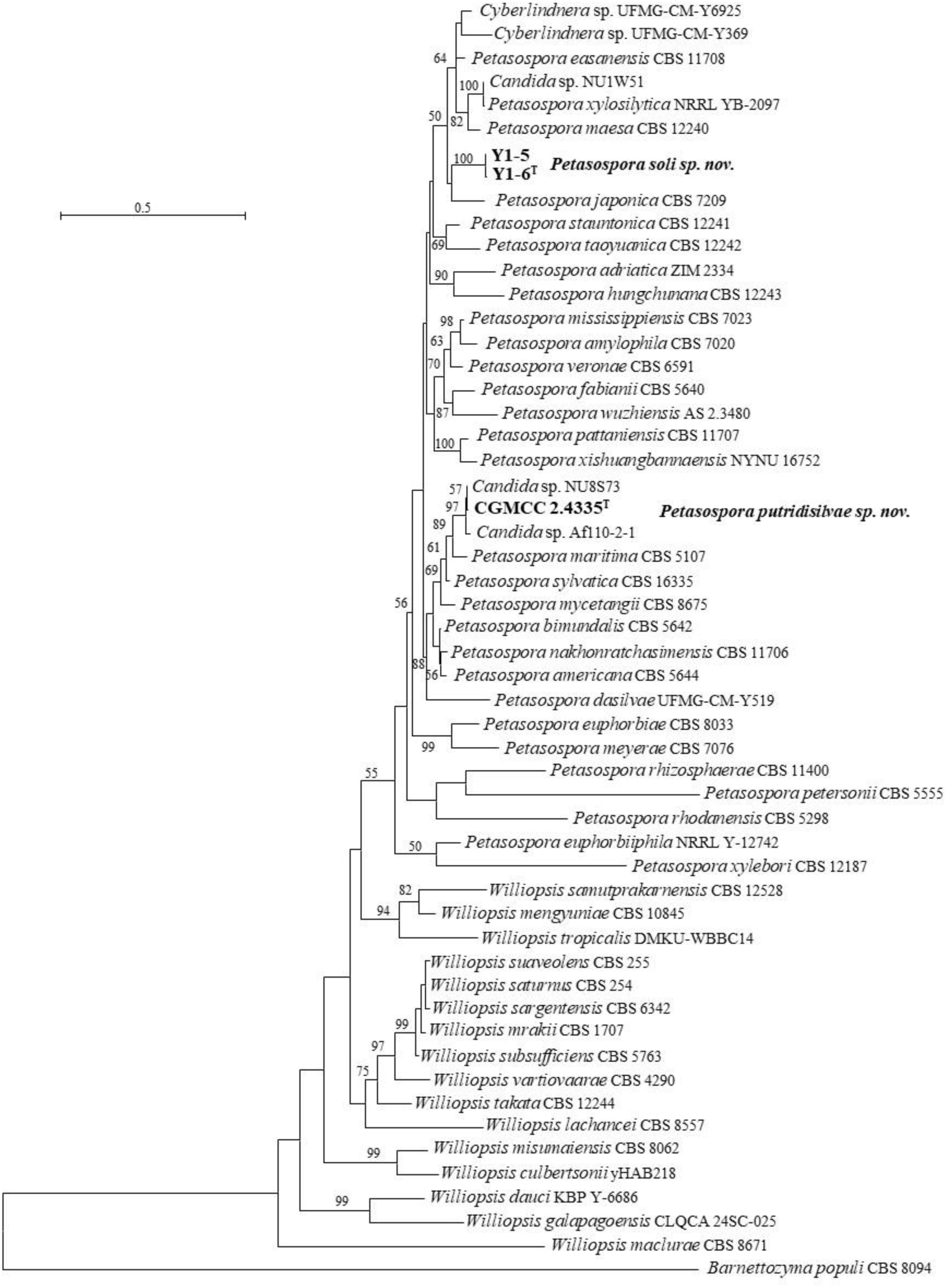
Phylogeny of new taxa (in bold) in the *Petasospora* inferred from the combined sequences of the LSU rDNA D1/D2 domains and ITS region (including 5.8S rDNA) by maximum likelihood analysis and over 50 % from 1 000 bootstrap replicates is shown. Bar = 0.5 substitutions per nucleotide position.

##### Yurkovozyma gen. nov

The phylogenomic analyses showed that the genus *Barnettozyma* is polyphyletic (Shen *et al*. 2018, Opulente *et al*. 2024, Liu *et al*. 2025). Two separate clades of *Barnettozyma*, namely the *Barnettozyma siamensis* clade (including *Barnettozyma salicaria*) and *Barnettozyma wickerhamii* clade, have been split from *Barnettozyma* and placed in the newly proposed genus *Gotozyma* and the reinstated genus *Komagataea* (Liu *et al*. 2025). The phylogenomic analysis and the ITS+LSU rDNA analysis from Liu *et al*. (2025) showed that XZY480-2 was located in a separate branch from *Gotozyma*, *Millerago* and *Komagataea*.

XZY480-2 and XZY480-10 have identical ITS and D1/D2 sequences. The blast analysis based on the ITS and D1/D2 sequences against the NCBI nucleotide database shows that those two strains have more sequence similarity with *Barnettozyma* than other genera in *Phaffomycetaceae*. XZY480-2 and XZY480-10 differed from the known species of *Barnettozyma* by 39 nt (11 %) in the D1/D2 domains and had less than 94.3% similarity with 46–75 % coverage in the ITS region. Those strains form a long branch closely related to *Gotozyma botsteinii*, *Gotozyma siamensis*, *Gotozyma montana* and *Komagataea salicaria* in the combined ITS and D1/D2 tree (Fig. 22). The ITS+LSU rDNA analysis from Liu *et al*. (2025) showed that XZY480-2 is located in a separate branch closely related to *Gotozyma*. However, the phylogenomic analysis showed that XZY480-2 was located in a basal branch related to *Millerago* and *Gotozyma*. The above data indicate that XZY480-2 and XZY480-10 do not belong to any known genera or clades in *Phaffomycetaceae*. Therefore, *Yurkovozyma gen. nov.* and *Yurkovozyma pini sp. nov.* are proposed to accommodate XZY480-2 and XZY480-10.

**Fig. 22.**
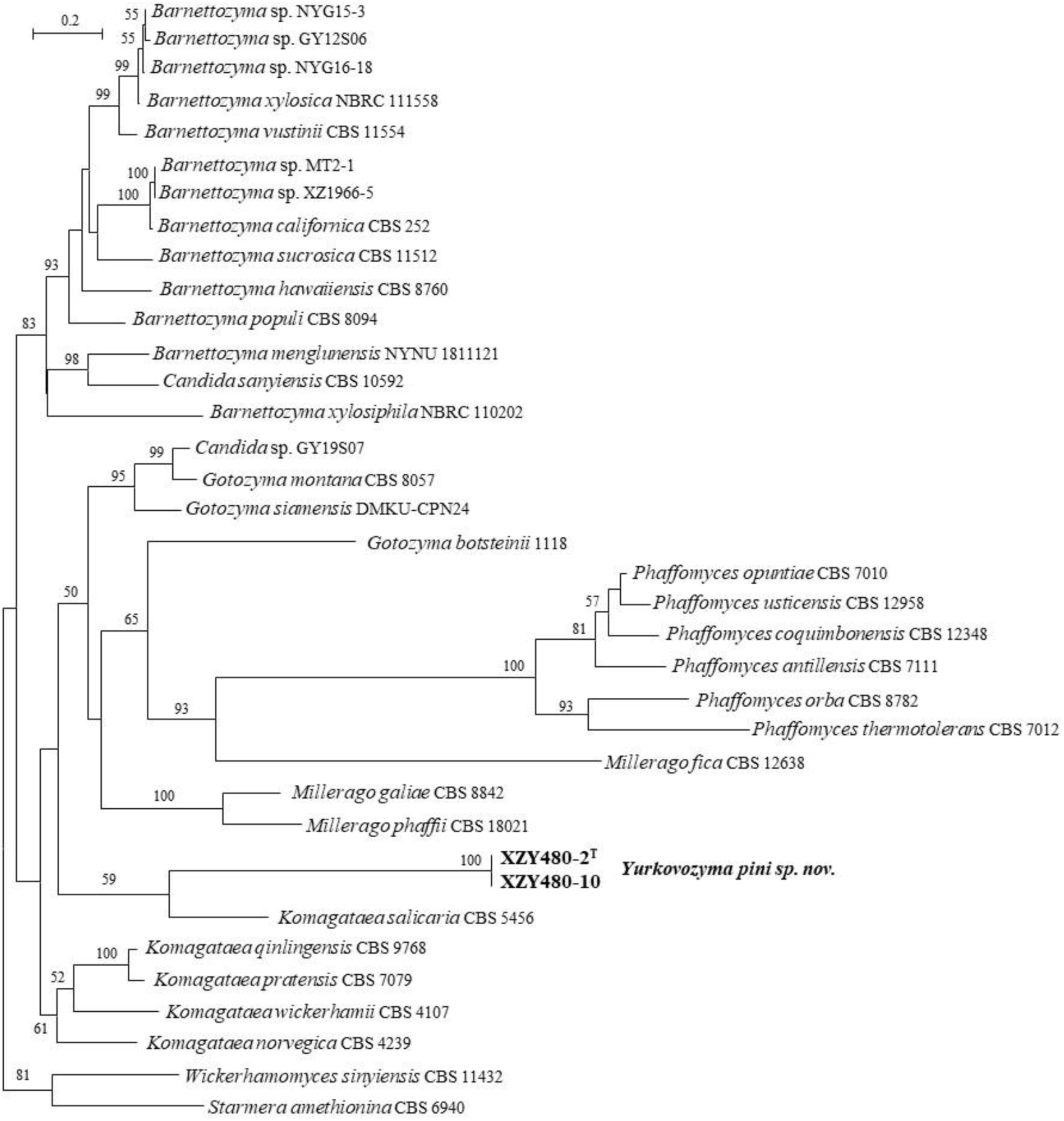
Phylogeny of new taxa (in bold) in the *Yurkovozyma* inferred from the combined sequences of the LSU rDNA D1/D2 domains and ITS region (including 5.8S rDNA) by maximum likelihood analysis and over 50 % from 1 000 bootstrap replicates is shown. Bar = 0.2 substitutions per nucleotide position.

#### New species identification in the Wickerhamomycetaceae (Phaffomycetales, Saccharomycetes, Saccharomycotina)

##### Hansenula

Liu *et al*. (2025) demonstrated that the genus *Wickerhamomyces* was polyphyletic and heterogeneous based on the phylogenomic and genome-based metric analyses and assigned six clades and one single-species lineage, namely, the *Wickerhamomyces sensu stricto* clade, the *Wickerhamomyces bovis* clade, the *Wickerhamomyces hampshirensis* clade, the *Wickerhamomyces chambardii* clade, the *Wickerhamomyces pijperi* clade, the *Hansenula* clade, and three single-species lineages *Wickerhamomyces kurtzmanii*, *Wickerhamomyces mucosus* and *Wickerhamomyces silvicola* to place *Wickerhamomyces* species and related *Candida* species. They reclassified this genus and proposed four new genera, namely *Lacusozyma* for *Wickerhamomyces kurtzmanii*, *Ruyongia* for the *Wickerhamomyces chambardii* clade, *Sucozym* for *Wickerhamomyces silvicola*, *Taiozyma* for the *Wickerhamomyces bovis* clade and *Xingzhongia* for the *Wickerhamomyces hampshirensis* clade, and reinstated the genus *Hansenula* and *Waltiozyma* for the *Hansenula* clade and *Wickerhamomyces mucosus+Wickerhamomyces pijperi* clade, respectively.

Three strains representing two groups are located in *Hansenula* (Fig. 23). Strains XZY836-1 and XZY836-1B have identical ITS sequences, differing from *Wickerhamomyces* sp. SK21-358 and SK21-383 by 3 nt and 5 nt in the D1/D2 domains and the ITS region, respectively, which indicates that they are conspecific. Those two strains differ from *Wickerhamomyces* sp. SK21-388, SK21-427, SK21-433, EN11M01, and NN10L12 by 3 nt and 9 nt in the D1/D2 domains and ITS region, respectively, which indicated that they may be the same species, but need more robust data, i.e., ANI, to confirm. XZY836-1 and XZY836-1B differ from the known *Wickerhamomyces* species by more than 7 nt and 14 nt (2.2 %) in the D1/D2 domains and the ITS region, respectively. Stains CGMCC 2.3630 and YShm6D2 have the same ITS and D1/D2 sequences. They are closely related to *Hansenula myanmarensis*, and differ from it by four nt (0.6 %) and more than 35 nt (5.4 %) in the D1/D2 domains and the ITS region, respectively. Based on the above analysis, we propose *Hansenula foliicola sp. nov.* and *Hansenula linzhienesis sp. nov.* for groups CGMCC 2.3630 and XZY836-1, respectively.

**Fig. 23.**
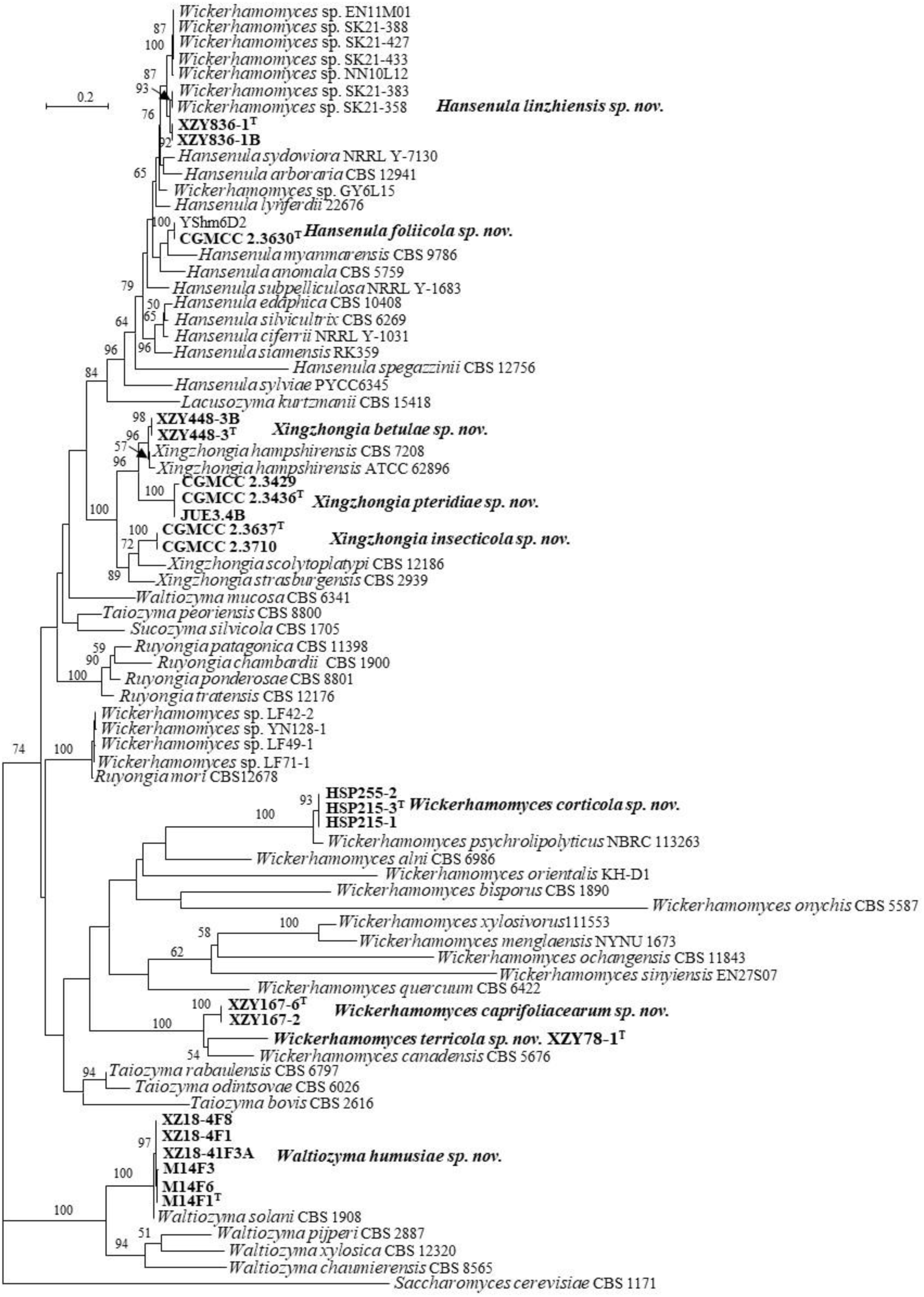
Phylogeny of new taxa (in bold) in the *Hansenula*, *Waltiozyma*, *Wickerhamomyces* and *Xingzhongia* inferred from the combined sequences of the LSU rDNA D1/D2 domains and ITS region (including 5.8S rDNA) by maximum likelihood analysis and over 50 % from 1 000 bootstrap replicates is shown. Bar = 0.2 substitutions per nucleotide position.

##### Waltiozyma

M14F1, M14F3, M14F6, XZ18-4F1, XZ18-4F8 and XZ18-41F3A have 1 nt both in the D1/D2 domains and ITS region. Those six strains are closely related to *Waltiozyma solani* (Fig. 23), and differ from it by 2 nt in the D1/D2 domains, but 18 nt (3 %) in the ITS region. Therefore, *Waltiozyma humusiae sp. nov.* is proposed to accommodate those six strains.

##### Xingzhongia

CGMCC 2.3429, CGMCC 2.3436, CGMCC 2.3637, CGMCC 2.3710, JUE3.4B, XZY448-3 and XZY448-3B form three groups and are located in the genus *Xingzhongia* (Fig. 23). CGMCC 2.3436, JUE3.4B and CGMCC 2.3429 with the same ITS and D1/D2 sequences differ from XZY448-3 and XZY448-3B by 10 nt (1.6 %) and 49 nt (8 %) in the D1/D2 domains and the ITS region, respectively. They are closely related to *Xingzhongia hampshirensis* (Fig. 23). XZY448-3 and XZY448-3B differ from *X. hampshirensis* by 2 nt and 11 nt (2.2 %) in the D1/D2 domains and ITS region, respectively. Strains CGMCC 2.3436, JUE3.4B and CGMCC 2.3429 differ from *X. hampshirensis* by 7 nt (1.2 %) and more than 9.4 % in the D1/D2 domains and the ITS region, respectively. Strains CGMCC 2.3637 and CGMCC 2.3710 have an affinity to *Xingzhongia scolytoplatypi* and *Xingzhongia strasburgensis*. They have 12–13 nt (2–2.1 %) difference in the D1/D2 domains, and more than 7.7 % in the ITS region. The above analyses indicate that those three groups represent three new species of *Xingzhongia*, therefore, *Xingzhongia betulae sp. nov.*, *Xingzhongia insecticola sp. nov.* and *Xingzhongia pteridiae sp. nov.* are proposed to accommodate groups XZY448-3, CGMCC 2.3637 and CGMCC 2.3436, respectively.

##### Wickerhamomyces

Six strains form three groups and are placed in *Wickerhamomyces* (Fig. 23). XZY78-1, XZY167-6 and XZY167-2 are closely related to *Wickerhamomyces canadensis* and differ from it by 10–11 nt in the D1/D2 domains, and more than 55–65 nt (9.6–12%) in the ITS region. Group XZ78-1 differs from group XZY167-6 by 4 nt in the D1/D2 domains, but by 70 nt (13 %) in the ITS region. HSP215-1, HSP215-3 and HSP255-2 with identical ITS and D1/D2 sequences have five nt in the ITS region with *Ascomycota* sp. KS83-4 (AB922629), which indicates that they may be the same species. They differ from the closely related species *Wickerhamomyces psychrolipolyticus* by more than 7 % and in the D1/D2 domains and 20 % ITS region, respectively. Based on the above analysis, we propose *Wickerhamomyces caprifoliaceae sp. nov.* for group XZY167-6, *Wickerhamomyces corticola sp. nov.* for group HSP215-1 and *Wickerhamomyces terricola sp. nov.* for group XZY78-1.

#### New species identification in the Saccharomycetaceae (Saccharomycetales, Saccharomycetes, Saccharomycotina)

##### Gaboromyces gen. nov

Liu *et al*. (2024a) reclassified the genus *Kazachstania* based on the genome-scale analysis and proposed five new genera and reinstated the genus *Arxiozyma* in the *Kazachstania* lineage. However, six single-species lineages of *Kazachstania* lineage, namely *Kazachstania bromeliacearum*, *Kazachstania kunashirensis*, *Kazachstania martiniae*, *Kazachstania molopis*, *Kazachstania psychrophila* and *Kazachstania taianensis*, were still placed in the *Kazachstania*, which were suggested to be separated from *Kazachstania* with the discovery of more related species (Liu *et al*. 2024a). Strains CGMCC 2.4518 and BW12-4 are closely related to *K. psychrophila* (Fig. 24) and differ from it by 6 nt (1 %) in the D1/D2 domains and by 9 nt (1.3 %) in the ITS region, respectively, which indicates that those two strains may represent a new species closely related to *K. kunashirensis*. The ANI value between CGMCC 2.4518 and *K. psychrophila* is 81.3 % (Table 3), which supports that group CGMCC 2.4518 represents a different species from *K. psychrophila*. Taking together the above analysis and the indication of (Liu *et al*. 2024a), a new genus *Gaboromyces gen. nov.* will be proposed to accommodate the new species and *K. psychrophila* in the Taxonomy section. Consequently, *Gaboromyces putridisilvae sp. nov.* is proposed to accommodate CGMCC 2.4518 and BW12-4. Strains CGMCC 2.4518 and BW12-4 have 0–4 nt ITS sequence difference with the uncultured fungus genomic DNA sequence (OU006364) isolated from fine root in EC San Francisco, Germany, and clone NF29 (AM711444) isolated from full-scale municipal compost in Finland, which indicates that the new genus and species occur in Europe.

**Fig. 24.**
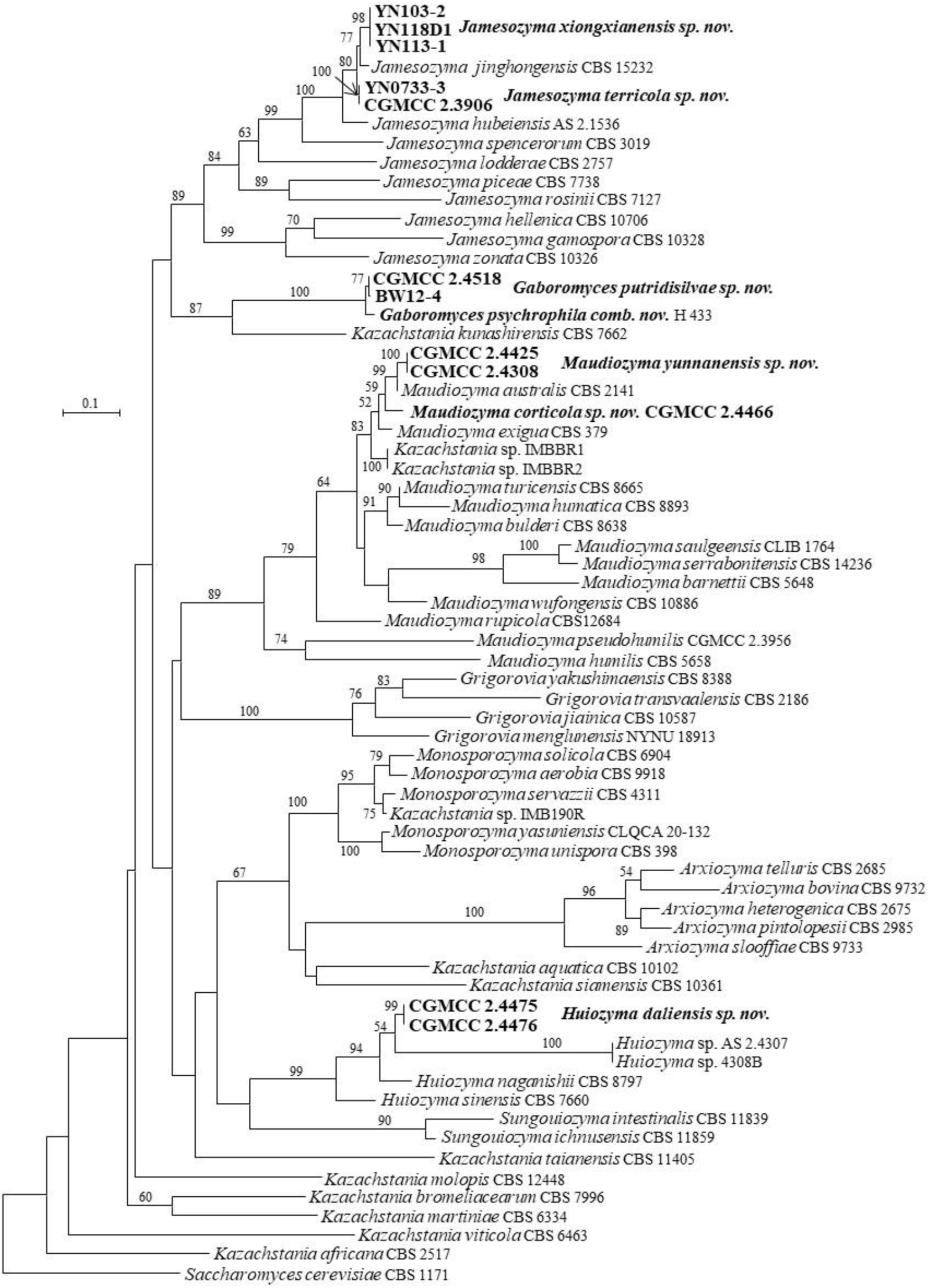
Phylogeny of new taxa (in bold) in the *Gaboromyces*, *Huiozyma*, *Jamesozyma* and *Maudiozyma* inferred from the combined sequences of the LSU rDNA D1/D2 domains and ITS region (including 5.8S rDNA) by maximum likelihood analysis and over 50 % from 1 000 bootstrap replicates is shown. Bar = 0.1 substitutions per nucleotide position.

##### Huiozyma

Strains CGMCC 2.4475 and CGMCC 2.4476 with identical ITS and D1/D2 sequences are closely related to *Huiozyma naganishii* and *Huiozyma* sp. AS 2.4307 and 4308B (Fig. 24). They differ from the two known species by 3–11 nt in the D1/D2 domains and more than 25 nt (3.8 %) in the ITS region, respectively. Thus, we propose *Huiozyma daliensis sp. nov.* to accommodate those two strains.

##### Jamesozyma

Strains CGMCC 2.3906, YN0733-3, YN113-1, YN118D1 and YN103-2 form two groups, and are closely related to *Jamesozyma jinghongensis* (Fig. 24). They differ from *J. jinghongensis* by 10–13 nt (1.9 %) in the ITS region and by 4–5 nt in the D1/D2 domains. YN113-1, YN118D1 and YN103-2 have identical ITS sequences with *Candida* sp. J12-2 (MW895082), J15-10 (MW895095), J18-2 (MW895081), J18-4 (MW895094) and J18-5 (MW895096) isolated from soil in China, which indicates that they are conspecific. They differ from CGMCC 2.3906 and YN0733-3 by 9 nt (1.2 %) in the ITS region and one nt in the D1/D2 domains. The ANI values between CGMCC 2.3906, YN113-1 and *J. jinghongensis* are 84.22–88.30 % (Table 3), which supports that groups CGMCC 2.3906 and YN113-1 are two distinct species from *J. jinghongensis.* Based on the above analysis, *Jamesozyma terricola sp. nov.* and *Jamesozyma xiongxianensis sp. nov.* are proposed to accommodate groups CGMCC 2.3906 and YN113-1, respectively.

##### Kluyveromyces

Strains CGMCC 2.4459, HSP168-1, HSP168-2, HSP168-3, XZY710-1 and XZY769-3 with identical ITS and D1/D2 sequences are closely related to *Kluyveromyces dobzhanskii* (Fig. 25), and differ from it by 9 nt in the ITS region and by 6 nt in the D1/D2 domains. The ANI value of *K. dobzhanskii* and those six isolates is 86.6 % (Table 3). Taken together, those six isolates should represent a new species of *Kluyveromyces*, for which *Kluyveromyces silvicola sp. nov.* is proposed to accommodate those six strains.

**Fig. 25.**
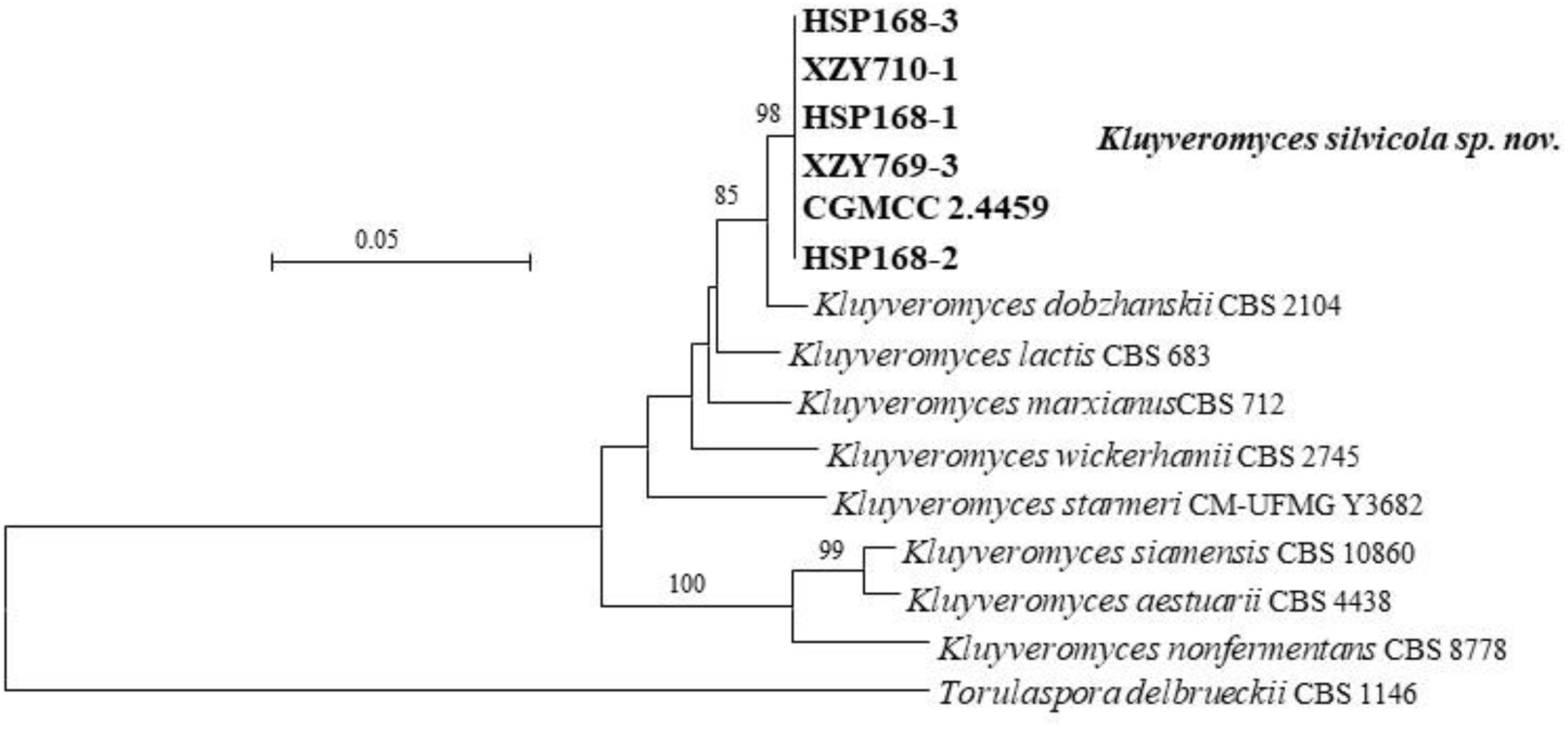
Phylogeny of new taxa (in bold) in the *Kluyveromyces* inferred from the combined sequences of the LSU rDNA D1/D2 domains and ITS region (including 5.8S rDNA) by maximum likelihood analysis and over 50 % from 1 000 bootstrap replicates is shown. Bar = 0.05 substitutions per nucleotide position.

##### Maudiozyma

Strains CGMCC 2.4308, CGMCC 2.4425 and CGMCC 2.4466 form two groups and have an affinity to *Maudiozyma exigua* (Fig. 24). They differ from it by 3–8 nt in the D1/D2 domains and by 18–23 nt (3–3.8 %) in the ITS region. CGMCC 2.4308 and CGMCC 2.4425 differed from CGMCC 2.4466 by 1 nt in the D1/D2 domains and by 18 nt (2.7 %) in the ITS region. Our ANI analysis showed that the ANI values are 95.9 % (pair of CGMCC 2.4425 and CGMCC 2.4466), 93.7 % (pair of CGMCC 2.4425 and *M. exigua*) and 89.9 % (pair of CGMCC 2.4466 and *M. exigua*), respectively, which indicates that CGMCC 2.4308, CGMCC 2.4425, and CGMCC 2.4466 represent two distinct species from *M. exigua*. Therefore, we propose *Maudiozyma corticola sp. nov.* for CGMCC 2.4466 and *Maudiozyma yunnanensis sp. nov.* for CGMCC 2.4308 and CGMCC 2.4425, respectively.

##### Naumovozyma

Strains CGMCC 2.4461 and YN125-1 with identical D1/D2 and four nt different ITS sequences are closely related to *Naumovozyma castellii* (Fig. 26), and differ from it by 1 nt in the D1/D2 domains, but by 20–23 nt (3–3.2 %) in the ITS region. Therefore, *Naumovozyma terricola sp. nov.* is proposed for CGMCC 2.4461 and YN125-1.

**Fig. 26.**
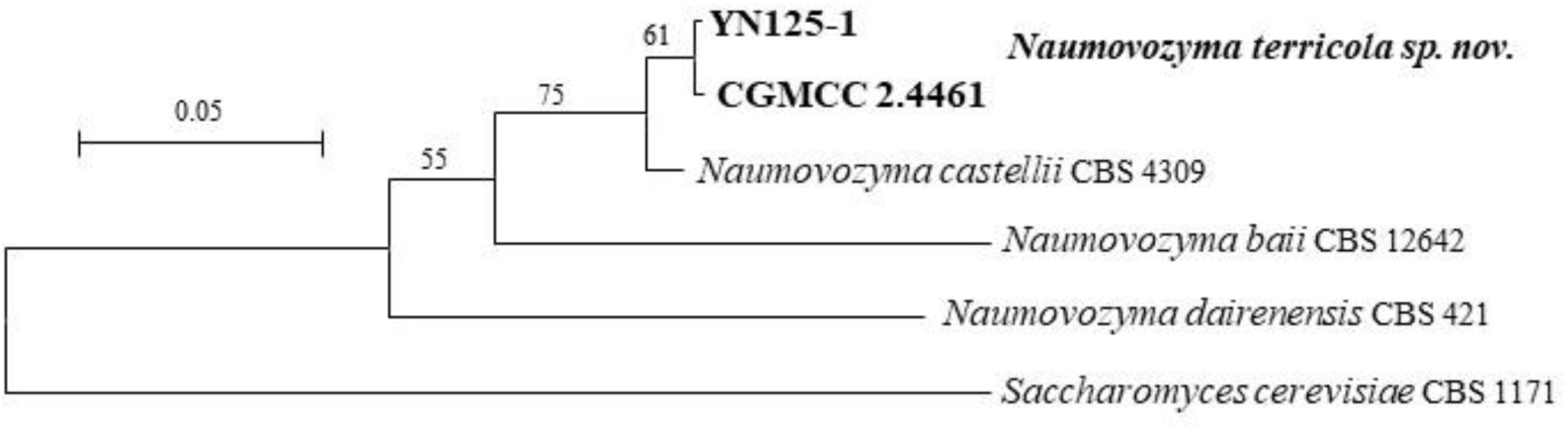
Phylogeny of new taxa (in bold) in the *Blastobotrys* inferred from the combined sequences of the LSU rDNA D1/D2 domains and ITS region (including 5.8S rDNA) by maximum likelihood analysis and over 50 % from 1 000 bootstrap replicates is shown. Bar = 0.05 substitutions per nucleotide position.

##### Torulaspora

XZY711-3, XZY765-1, XZY735-1, XZY238F1, XZY238F2 and XZY719-1 have identical ITS and D1/D2 sequences. They are closely related to *Torulaspora delbrueckii* (Fig. 27) and differ from it by 6 nt (1 %) and 23 nt (2.9 %) in the D1/D2 domains and the ITS region, respectively, which indicates that those six strains represent a new species of *Torulaspora*. Therefore, *Torulaspora xizangensis sp. nov.* is proposed to accommodate them.

**Fig. 27.**
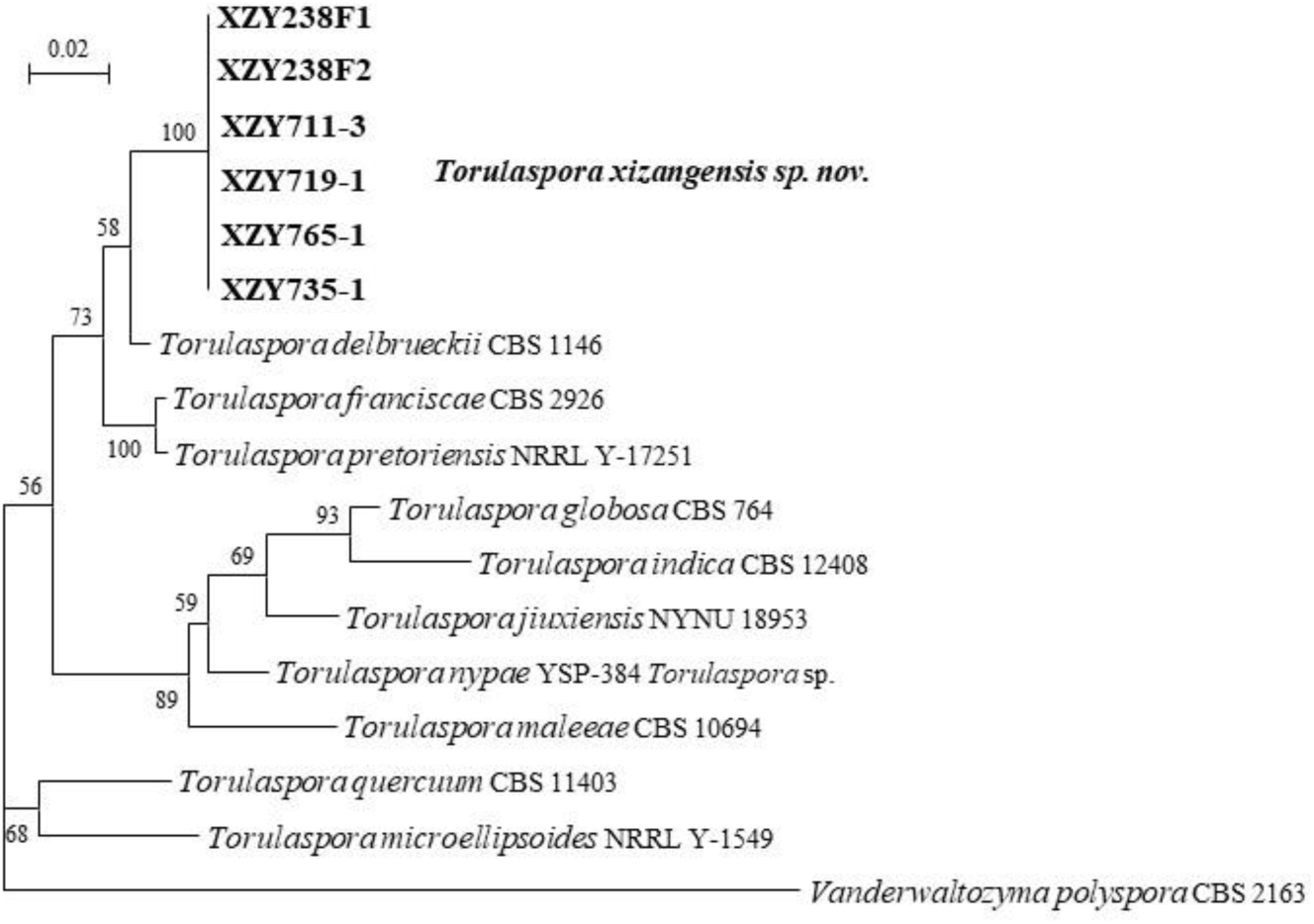
Phylogeny of new taxa (in bold) in the *Torulaspora* inferred from the combined sequences of the LSU rDNA D1/D2 domains and ITS region (including 5.8S rDNA) by maximum likelihood analysis and over 50 % from 1 000 bootstrap replicates is shown. Bar = 0.02 substitutions per nucleotide position.

##### Zygotorulaspora

Strains CGMCC 2.4531 and BW31-7 are closely related to *Zygotorulaspora cariocana* (Fig. 28) and differ from it by 1–6 nt in the D1/D2 domains, but 19–28 nt (4.6–6 %) in the ITS region, which indicates that they belong to different species. Those two strains differ from the other known *Zygotorulaspora* species by 31 nt (5 %) in the D1/D2 domains and by 124 nt (18.9 %) in the ITS region. Thus, we propose *Zygotorulaspora silvicola sp. nov.* to accommodate CGMCC 2.4531 and BW31-7.

**Fig. 28.**
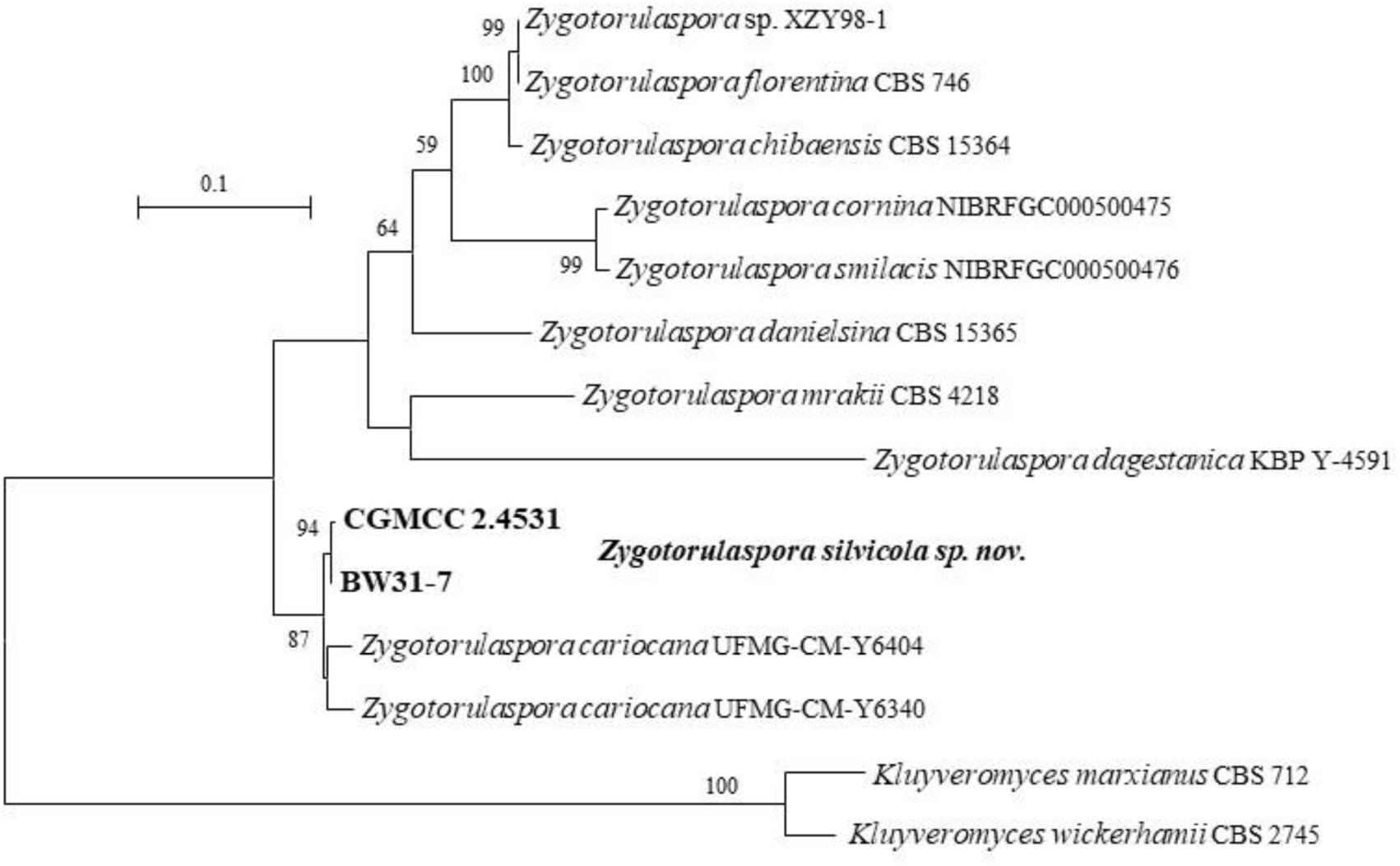
Phylogeny of new taxa (in bold) in the *Zygotorulaspora* inferred from the combined sequences of the LSU rDNA D1/D2 domains and ITS region (including 5.8S rDNA) by maximum likelihood analysis and over 50 % from 1 000 bootstrap replicates is shown. Bar = 0.1 substitutions per nucleotide position.

#### New species identification in the *Taphrina* (*Taphrinaceae*, *Taphrinales*, *Taphrinomycetes*, Taphrinomycotina)

##### Taphrina

Strain CGMCC 2.2537 has an identical D1/D2 sequence and two nt ITS differences with *Taphrina sp.* NYNU 221115. They are closely related to *Taphrina americana*, *Taphrina carpini*, *Taphrina betulina*, and *Taphrina polystichi* (Fig. 29), and differ from them by 3–4 nt in the D1D2 domain, but more than 34 nt (6.3 %) in the ITS region. Strain CGMCC 2.5682 has an identical D1/D2 sequence and two nt ITS differences with *Taphrina sp.* NYNU 224115. They cluster with *Taphrina caerulescens* and *Taphrina kurtzmanii* (Fig. 29), and differ from them by 11 (2 %) in the D1/D2 domains and by more than 81 nt (14%) in the ITS region. The above analysis indicates that CGMCC 2.2537 and CGMCC 2.5682 represent two new *Taphrina* species, therefore, *Taphrina follicola sp. nov.* and *Taphrina planticola sp. nov.* are proposed to accommodate them.

**Fig. 29.**
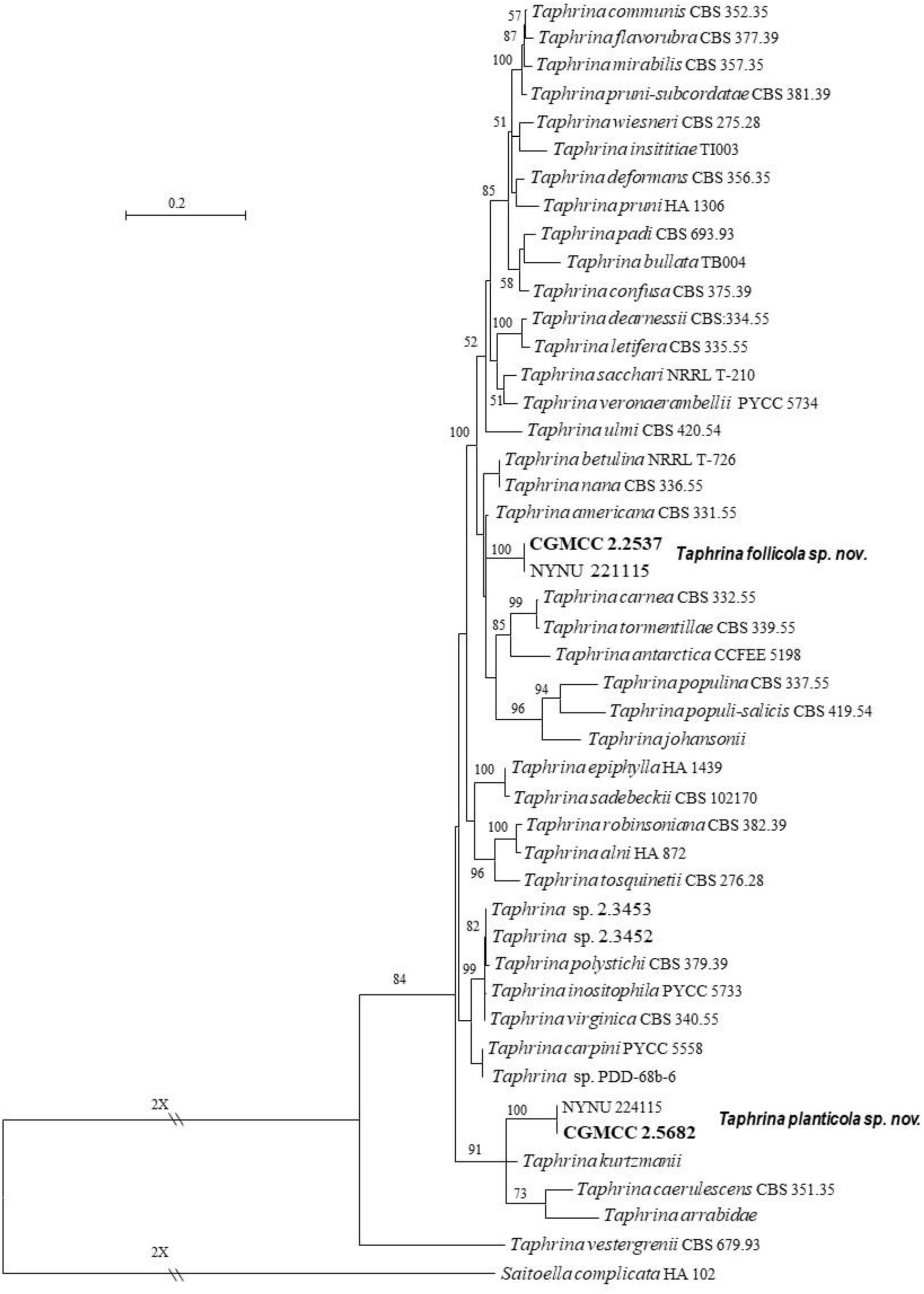
Phylogeny of new taxa (in bold) in the *Taphrina* inferred from the combined sequences of the LSU rDNA D1/D2 domains and ITS region (including 5.8S rDNA) by maximum likelihood analysis and over 50 % from 1 000 bootstrap replicates is shown. Bar = 0.2 substitutions per nucleotide position.

##### Taxonomy

New taxa in the *Trichomonascaceae* (*Trichomonascales*, *Dipodascomycetes*, *Saccharomycotina*) *Blastobotrys corticola* Q.-M. Wang, A.-H. Li, M.-M. Liu, H.-H. Zhu & F.-Y. Bai, *sp. nov.* Fungal Names FN 572745. Fig. 30A, B.

**Fig. 30.**
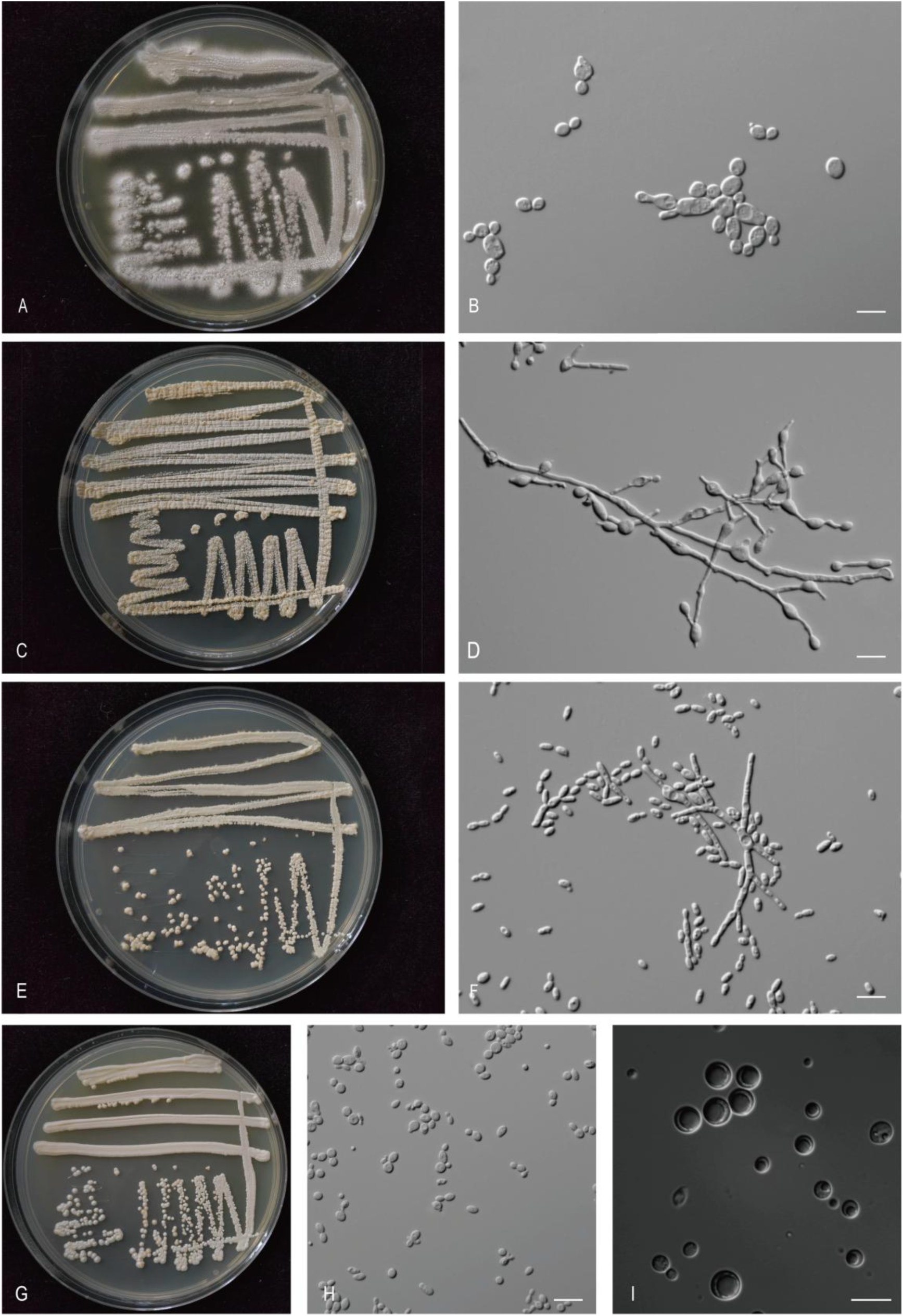
The streak culture grown in YM agar and vegetative cells grown in YM broth for 7 d at 25 °C. A, B. *Bla. corticola* CGMCC 2.6978^T^. C, D. *Bla. pinus* CGMCC 2.7032^T^. E, F. *Bla. xizangensis* 2.6979^T^. G-I. *Did. polypori* CGMCC 2.6974^T^. Scale bars: B, D, F, H, I = 10 μm.

###### Etymology

the specific epithet *corticola* refers to the type strain isolated from bark.

###### Culture characteristics

On YM agar, after 1 month at 25 °C, the streak culture is white, rough and dimmed, with dry powder. The margin is fringed (Fig. 30A). In YM broth, after 3 days at 25 °C, cells are subglobosal and oval, 2.42–6.30 × 3.89–10.47 µm and single, budding is polar (Fig. 30B), after 7 days at 25 °C, sediment and pellicle are formed. After 1 month at 25 °C, sediment and pellicle are present, a ring does not appear. In Dalmau plate culture on corn meal agar, hyphae are formed. Asci and ascospores are not observed on V8 and Acetate agar.

###### Physiological and biochemical characteristics

Glucose, galactose, sucrose (variable), maltose (variable) and raffinose (variable) fermentation are present. Lactose fermentation is absent. Glucose, galactose, sucrose (variable), maltose, cellobiose, trehalose, lactose (variable), melibiose (variable), raffinose (variable), melezitose (variable), inulin (weak), soluble starch (variable), D-xylose (variable), L-rhamnose (variable), N-acetyl-D-glucosamine, ethanol, glycerol (variable), erythritol (variable), ribitol (variable), galactitol (variable), D-mannitol (weak), D-glucitol (variable), methyl-α-D-glucoside (variable), salicin (variable), DL-lactic acid (variable) and succinic acid (variable) are assimilated as sole carbon sources. L-sorbose, L-arabinose, D-arabinose, D-ribose, D-glucosamine, methanol, citric acid, inositol, hexadecane and D-glucuronic acid are not assimilated. Ammonium sulfate, potassium nitrate (variable), sodium nitrite (variable), L-lysine, ethylamine (latent) and cadaverine (weak) are assimilated as sole nitrogen sources. The maximum growth temperature is 42 °C. Growth in the vitamin-free medium is positive. Starch-like substances are not produced. Growth on 50 % (w/w) glucose-yeast extract agar is positive (weak).

Physiologically, *B. corticola* differs from the closely related species *B. buckinghamii*, *B. capitulates* and *B. arbuscula* (Fig. 2) in its inability to assimilate L-sorbose and L-arabinose (Table S2.1).

###### Typus

**China**, Daliangzi River to Liangshui, Heilongjiang Province, obtained from bark of *Pinus*, Aug. 2019, *Q.-M. Wang* (**holotype** CGMCC 2.6978^T^ preserved in a metabolically inactive state, ex-type cultures are preserved in culture collections under the numbers CBS 18380 and HSP57-1).

###### Other culture examined

**China**, Daliangzi River to Liangshui, Heilongjiang Province, obtained from bark, Nov. 2017, isolated by *Q.M. Wang*, culture HSP49-2; **China**, Greater Khinganling National Forest Reserve, Heilongjiang Province, obtained from bark, Sep. 2017, isolated by *Q.M. Wang*, culture HSP187-2.

***Blastobotrys pinus*** Q.-M. Wang, A.-H. Li, M.-M. Liu, H.-H. Zhu & F.-Y. Bai, ***sp. nov.*** Fungal Names FN 572746. Fig. 30C, D.

###### Etymology

the specific epithet *pinus* refers to *Pinus*, the plant genus from which the type strain was isolated.

###### Culture characteristics

On YM agar, after 1 month at 25 °C, the streak culture is white to yellowish-cream, butyrous, and rough. The margin is entire (Fig. 30C). In YM broth, after 3 days at 25 °C, cells are subglobose and oval, 2.42–3.85 × 2.68–4.68 µm and single, budding is polar (Fig. 30D), after 7 days at 25 °C, sediment and pellicle are formed. After 1 month at 25 °C, sediment and complete pellicle are present, and a ring does not appear. In Dalmau plate culture on corn meal agar, hyphae are formed. Asci and ascospores are not observed on V8 and Acetate agar.

###### Physiological and biochemical characteristics

Glucose and galactose fermentation are present. Lactose, sucrose, maltose and raffinose fermentation are absent. Glucose, galactose (latent), sucrose (latent, weak), maltose (latent), cellobiose (latent), trehalose (weak), lactose (latent, weak), raffinose (latent, weak), Inulin (latent, weak), soluble starch (latent), D-xylose (latent, weak), L-rhamnose (latent), ribitol (latent, weak), D-mannitol (latent, weak) and D-glucuronic acid (latent, weak) are assimilated as sole carbon sources. L-sorbose, melibiose, melezitose, L-arabinose, D-arabinose, D-ribose, D-glucosamine, N-acetyl-D-glucosamine, methanol, ethanol, glycerol, erythritol, galactitol, D-glucitol, methyl-α-D-glucoside, salicin, DL-lactic acid, succinic acid, citric acid, inositol and hexadecane are not assimilated. Ammonium sulfate (latent, weak) and cadaverine (latent, weak) are assimilated as sole nitrogen sources. Potassium nitrate, sodium nitrite, L-lysine and ethylamine are not assimilated. The maximum growth temperature is 20 °C. Growth in the vitamin-free medium is positive (latent). Starch-like substances are not produced. Growth on 50 % (w/w) glucose-yeast extract agar is positive (weak).

Physiologically, *B. pinus* differs from the closely related species *B. farinosus* (Fig. 2) in its inability to assimilate D-glucitol (Table S2.1).

###### Typus

**China**, Basongcuo, Tibet, obtained from bark of *Pinus*, Aug. 2019, *Q.-M. Wang* (**holotype** CGMCC 2.7032^T^ preserved in a metabolically inactive state, ex-type cultures are preserved in culture collections under the numbers XZY496-2).

###### Other culture examined

**China**, Basongcuo, Tibet, obtained from bark of *Pinus*, Aug. 2019, isolated by *Q.M. Wang*, *G.S. Wang*, culture XZY496-12.

***Blastobotrys xizangensis*** Q.-M. Wang, A.-H. Li, M.-M. Liu, H.-H. Zhu & F.-Y. Bai, ***sp. nov.*** Fungal Names FN 572747. Fig. 30E, F.

###### Etymology

the specific epithet *xizangensis* refers to the geographic origin of the type strain, Xizang.

###### Culture characteristics

On YM agar, after 1 month at 25 °C, the streak culture is white, friable, rough and dimmed, with dry powder. The margin is a punctate protrusion (Fig. 30E). In YM broth, after 3 days at 25 °C, cells are ellipsoidal and rod, 1.64–3.03 × 3.50–5.31 µm, and single, budding is polar (Fig. 30F), after 7 days at 25 °C, sediment and pellicle are formed. After 1 month at 25 °C, sediment and pellicle are present, but a ring does not appear. In Dalmau plate culture on corn meal agar, pseudohyphae are formed. Asci and ascospores are not observed on V8 and Acetate agar.

###### Physiological and biochemical characteristics

Glucose, galactose (variable) and maltose (variable) fermentation are present. Lactose, sucrose and raffinose fermentation are absent. Glucose, galactose (latent), L-sorbose (latent), sucrose (variable), maltose (latent), cellobiose (latent), trehalose (latent), lactose (variable), inulin (latent), soluble starch, D-xylose, L-arabinose (latent), D-arabinose (variable), D-ribose (latent), L-rhamnose (latent, weak), D-glucosamine, N-acetyl-D-glucosamine, ethanol (variable), glycerol (latent, weak), erythritol (latent, weak), ribitol (latent, weak), galactitol (latent, weak), D-mannitol, D-glucitol, methyl-α-D-glucoside(latent), succinic acid (variable), inositol (latent, weak) and D-glucuronic acid (variable) are assimilated as sole carbon sources. Melibiose, raffinose, melezitose, methanol, salicin, DL-lactic acid, citric acid and hexadecane are not assimilated. Ammonium sulfate, potassium nitrate, sodium nitrite, L-lysine, ethylamine and cadaverine are assimilated as sole nitrogen sources. The maximum growth temperature is 32 °C. Growth in the vitamin-free medium is positive. Starch-like substances are not produced. Growth on 50 % (w/w) glucose-yeast extract agar is positive (variable).

Physiologically, *B. xizangensis* differs from the closely related species *T. apis* (Fig. 2) in its inability to assimilate melibiose, raffinose, salicin, DL-lactic acid and citric acid, the ability to assimilate inulin, potassium nitrate, sodium nitrite and grow in vitamin-free medium (Table S2.1).

###### Typus

**China**, Bomi County, Nyingchi City, Tibet, obtained from rotten wood, Aug. 2019, *Q.-M. Wang* (**holotype** CGMCC 2.6979^T^ preserved in a metabolically inactive state, ex-type cultures are preserved in culture collections under the numbers CBS 18381 and BMFM2F2).

###### Other culture examined

**China**, Nagqu, Tibet, obtained from soil, Aug. 2019, isolated by *Q.M. Wang*, *G.S. Wang*, culture QZKF1. **China**, Milin County, Linzhi, Tibet, obtained from rotten wood, Aug. 2019, isolated by *Q.M. Wang*, *G.S. Wang*, culture L22-3.

***Diddensiella polypori*** Q.-M. Wang, A.-H. Li, M.-M. Liu, H.-H. Zhu & F.-Y. Bai, ***sp. nov.*** Fungal Names FN 572748. Fig. 30G-I.

###### Etymology

the specific epithet *polypori* refers to *Polyporus*, the mushroom genus from which the type strain was isolated.

###### Culture characteristics

On YM agar, after 1 month at 25 °C, the streak culture is off-white, butyrous, smooth and pale glossy. The margin is entire (Fig. 30G). In YM broth, after 3 days at 25 °C, cells are subcircular and oval, 1.74–3.67 × 2.92–5.64 µm and single, budding is polar (Fig. 30H), after 7 days at 25 °C, a sediment is formed. After 1 month at 25 °C, sediment is present, and the ring and pellicle do not appear. In Dalmau plate culture on corn meal agar, pseudohyphae are not formed. On V8 agar, the ascus is oval and contains one ascospore (Fig. 30I). The ascospores are spherical or subspherical, 2.24–4.56 × 2.16–4.45 µm.

###### Physiological and biochemical characteristics

Glucose, lactose, galactose, sucrose, maltose and raffinose fermentation are absent. Glucose, galactose (latent), L-sorbose, sucrose (latent, weak), maltose, cellobiose (latent), trehalose, lactose (latent), melibiose, raffinose, melezitose, inulin, soluble starch (latent), D-xylose, L-arabinose, D-arabinose, D-ribose (latent, weak), L-rhamnose, D-glucosamine, N-acetyl-D-glucosamine, ethanol, glycerol (latent), erythritol (latent), ribitol, galactitol (latent, weak), D-mannitol, D-glucitol (weak),methyl-α-D-glucoside, salicin, DL-lactic acid (latent), succinic acid (latent), citric acid, inositol (latent) and D-glucuronic acid (latent, weak) are assimilated as sole carbon sources. Methanol and hexadecane are not assimilated. Ammonium sulfate, potassium nitrate, sodium nitrite (latent, weak), L-lysine, ethylamine and cadaverine are assimilated as sole nitrogen sources. The maximum growth temperature is 25 °C. Growth in the vitamin-free medium is positive. Starch-like substances are not produced. Growth on 50 % (w/w) glucose-yeast extract agar is negative.

Physiologically, *D. polypori* differs from the closely related species *D. caesifluorescens* and *D. cacticola* (Fig. 3) in its ability to assimilate lactose, soluble starch, galactitol, DL-lactic acid, D-glucuronic acid and grow in vitamin-free medium (Table S2.2).

###### Typus

**China**, New basom lake, Linzhi, Tibet, obtained from *Polyporus* species, Aug. 2019, *Q.-M. Wang* (**holotype** CGMCC 2.6974^T^ preserved in a metabolically inactive state, ex-type cultures are preserved in culture collections under the numbers XZY423-5).

***Starmerella eumenesae*** Q.-M. Wang, A.-H. Li, M.-M. Liu, H.-H. Zhu & F.-Y. Bai, ***sp. nov.*** Fungal Names FN 572749. Fig. 31A, B.

**Fig. 31.**
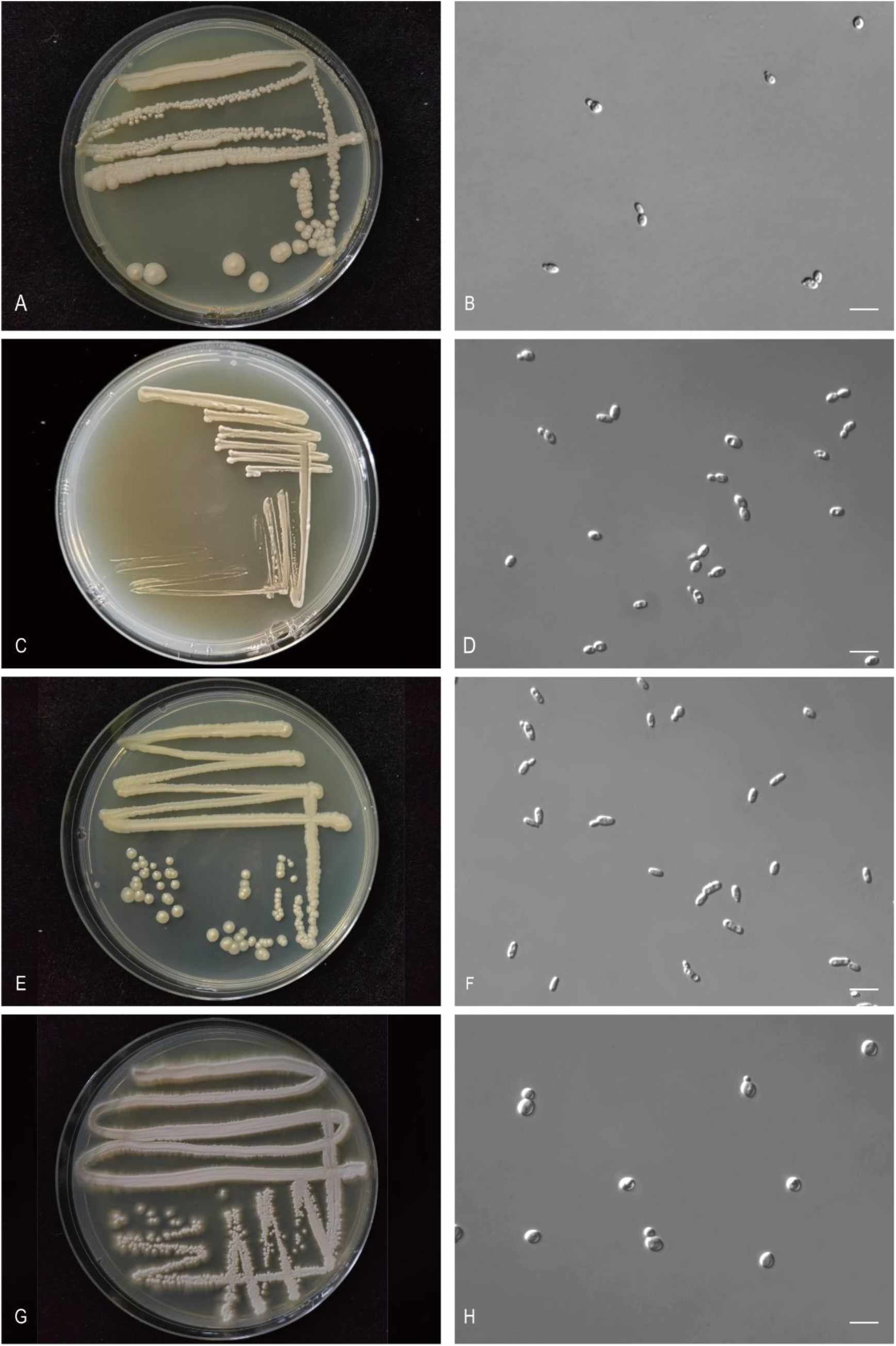
The streak culture grown in YM agar and vegetative cells grown in YM broth for 7 d at 25 °C. A, B. *Sta. eumenesae* CGMCC 3704^T^. C, D. *Sta. parabombi* CGMCC 2.3767^T^. E, F. *Sta. putridisilvae* CGMCC 2.3791^T^. G, H. *Sug. hainanensis* CGMCC 2.3787^T^. Scale bars: B, D, F, H = 10 μm.

###### Etymology

the specific epithet *eumenesae* refers to the type strain isolated from a bee. Epithet *polypori* refers to *Eumenes*, the insect genus from which the type strain was isolated.

###### Culture characteristics

On YM agar, after 1 month at 25 °C, the streak culture is off-white, butyrous, smooth, and pale glossy. The margin is entire (Fig. 31A). In YM broth, after 3 days at 25 °C, cells are fusiform, ellipsoidal and oval, 1.99–3.42 × 3.47–6.59 µm and single, budding is polar (Fig. 31B), after 7 days at 25 °C, a sediment is formed. After 1 month at 25 °C, sediment is present, but a ring and pellicle do not appear. In Dalmau plate culture on corn meal agar, pseudohyphae are not formed. Asci and ascospores are not observed on V8 and Acetate agar.

###### Physiological and biochemical characteristics

Glucose and sucrose (weak) fermentation are present. Lactose, galactose, maltose and raffinose fermentation are absent. Glucose, galactose (variable), L-sorbose, sucrose (variable), maltose (variable), melibiose, raffinose, inulin, D-xylose, L-arabinose, D-arabinose, D-ribose, ribitol (weak), galactitol, D-mannitol, D-glucitol, methyl-α-D-glucoside (variable), salicin, DL-lactic acid (variable), succinic acid and citric acid are assimilated as sole carbon sources. Cellobiose, trehalose, lactose, melezitose, soluble starch, L-rhamnose, D-glucosamine, N-acetyl-D-glucosamine, methanol, ethanol, glycerol, erythritol, inositol, hexadecane and D-glucuronic acid are not assimilated. Ammonium sulfate, potassium nitrate, L-lysine, ethylamine and cadaverine are assimilated as sole nitrogen sources. Sodium nitrite is not assimilated. The maximum growth temperature is 37 °C.

Growth in the vitamin-free medium is positive. Starch-like substances are not produced. Growth on 50 % (w/w) glucose-yeast extract agar is negative.

Physiologically, *S. eumenesae* differs from the closely related species *S. aceti*, *S. apicola*, *S. reginensis* and *S. neotropicalis* (Fig. 4) in its ability to assimilate D-xylose and citric acid (Table S2.3).

###### Typus

**China**, Beijing, obtained from the gut of *Eumenes quadratus*, Feb. 2007, *F.-Y. Bai* (**holotype** CGMCC 2.3704^T^ preserved in a metabolically inactive state, ex-type cultures are preserved in culture collections under the numbers CBS 18406 = NHF1.2).

###### Other culture examined

**China**, Beijing, obtained from the gut of *Eumenes quadratus*, isolated by *F.-Y. Bai*, culture CGMCC 2.3652 (= NHF1.4).

***Starmerella parabombi*** Q.-M. Wang, A.-H. Li, M.-M. Liu, H.-H. Zhu & F.-Y. Bai, ***sp. nov.*** Fungal Names FN 572750. Fig. 31C, D.

###### Etymology

the specific epithet *parabombi* refers to a similar colony morphology to that of *Starmerella bombi*.

###### Culture characteristics

On YM agar, after 1 month at 25 °C, the streak culture is white, butyrous, smooth and glossy. The margin is entire (Fig. 31C). In YM broth, after 3 days at 25 °C, cells are ellipsoidal and fusiform, 2.27–4.03 × 3.90–5.53 µm and single, budding is polar (Fig. 31D), after 7 days at 25 °C, a sediment is formed. After 1 month at 25 °C, sediment is present, but a ring and pellicle do not appear. In Dalmau plate culture on corn meal agar, pseudohyphae are not formed. Asci and ascospores are not observed on V8 and Acetate agar.

###### Physiological and biochemical characteristics

Glucose, galactose, sucrose and maltose fermentation are present. Lactose and raffinose fermentation are absent. Glucose, L-sorbose, sucrose, maltose (weak), cellobiose (weak), trehalose, lactose (weak), melibiose, raffinose, melezitose (weak), inulin (weak), soluble starch (weak), D-xylose, D-ribose, L-rhamnose (weak), ethanol (latent), glycerol, ribitol (latent, weak), D-mannitol, D-glucitol, methyl-α-D-glucoside (weak), DL-lactic acid (weak), succinic acid and citric acid are assimilated as sole carbon sources. Galactose, L-arabinose, D-arabinose, D-glucosamine, N-acetyl-D-glucosamine, methanol, erythritol, galactitol, salicin, inositol, hexadecane and D-glucuronic acid are not assimilated. Ammonium sulfate, potassium nitrate, sodium nitrite, L-lysine, ethylamine and cadaverine are assimilated as sole nitrogen sources. The maximum growth temperature is 42 °C. Growth in the vitamin-free medium is positive. Starch-like substances are not produced. Growth on 50 % (w/w) glucose-yeast extract agar is positive.

Physiologically, *S. parabombi* differs from the closely related species *S. bombi* (Fig. 4) in its inability to assimilate D-glucuronic acid, and the ability to ferment galactose and maltose and assimilate maltose, cellobiose, lactose, melibiose, melezitose, inulin, D-xylose, D-ribose, L-rhamnose, ethanol, ribitol, methyl-α-D-glucoside, DL-lactic acid, succinic acid, citric acid, potassium nitrate, sodium nitrite and grow in vitamin-free medium (Table S2.3).

###### Typus

**China**, Jilin province, obtained from leaf, Oct. 2007, *Q.-M. Wang* (**holotype** CGMCC 2.3767^T^ preserved in a metabolically inactive state).

###### Other culture examined

**China**, Jilin province, obtained from leaf, Oct. 2007, isolated by *Q.M. Wang*, culture 46YC.

***Starmerella putridisilvae*** Q.-M. Wang, A.-H. Li, M.-M. Liu, H.-H. Zhu & F.-Y. Bai, *sp. nov.* Fungal Names FN 572751. Fig. 31E, F.

###### Etymology

the specific epithet *putridisilvae* refers to the type strain isolated from rotten wood.

###### Culture characteristics

On YM agar, after 1 month at 25 °C, the streak culture is yellowish, butyrous, smooth, and pale glistening. The margin is entire (Fig. 31E). In YM broth, after 3 days at 25 °C, cells are ellipsoidal and cylindrical, 1.93–3.15 × 4.74–7.62 µm and single, budding is polar (Fig. 31F), after 7 days at 25 °C, a sediment is formed. After 1 month at 25 °C, sediment is present, but a ring and pellicle do not appear. In Dalmau plate culture on corn meal agar, pseudohyphae are not formed. Asci and ascospores are not observed on V8 and Acetate agar.

###### Physiological and biochemical characteristics

Glucose and sucrose fermentation are present. Lactose, galactose, maltose and raffinose fermentation are absent. Glucose, galactose, L-sorbose, sucrose, melibiose, raffinose, inulin (weak), D-xylose (latent), D-ribose, glycerol, D-mannitol, D-glucitol, succinic acid and citric acid are assimilated as sole carbon sources. Maltose, cellobiose, trehalose, lactose, melezitose, soluble starch, L-arabinose, D-arabinose, L-rhamnose, D-glucosamine, N-acetyl-D-glucosamine, methanol, ethanol, erythritol, ribitol, galactitol, methyl-α-D-glucoside, salicin, DL-lactic acid, inositol, hexadecane and D-glucuronic acid are not assimilated. Ammonium sulfate, potassium nitrate, sodium nitrite, L-lysine, ethylamine and cadaverine are assimilated as sole nitrogen sources. The maximum growth temperature is 35 °C. Growth in the vitamin-free medium is positive. Starch-like substances are not produced. Growth on 50 % (w/w) glucose-yeast extract agar is negative.

Physiologically, *S. putridisilvae* differs from the closely related species *S. caucasica*, *S. bombicola* and *S. kuoi* (Fig. 4) in its inability to assimilate ethanol (Table S2.3).

###### Typus

**China**, Diaolunshan, Hainan province, obtained from rotten wood, Jan. 2008, *Q.-M. Wang* (**holotype** CGMCC 2.3791^T^ preserved in a metabolically inactive state).

***Sugiyamaella hainanensis*** Q.-M. Wang, A.-H. Li, M.-M. Liu, H.-H. Zhu & F.-Y. Bai, ***sp. nov.*** Fungal Names FN 572752. Fig. 31G, H.

###### Etymology

the specific epithet *hainanensis* refers to the geographic origin of the type strain, Hainan province.

###### Culture characteristics

On YM agar, after 1 month at 25 °C, the streak culture is whitish, butyrous, rough, and pale dimmed. The margin is fimbriate (Fig. 31G). In YM broth, after 3 days at 25 °C, cells are ellipsoidal and oval, 3.48–5.26 × 4.50–6.25 µm and single, budding is polar (Fig. 31H), after 7 days at 25 °C, a sediment is formed. After 1 month at 25 °C, sediment is present, pellicle and ring do not appear. In Dalmau plate culture on corn meal agar, hyphae are formed. Asci and ascospores are not observed on V8 and Acetate agar.

###### Physiological and biochemical characteristics

Glucose (weak) fermentation is present. Lactose, galactose, sucrose, maltose and raffinose fermentation are absent. Glucose, galactose, L-sorbose, sucrose, maltose, cellobiose, trehalose, lactose, melibiose, raffinose, melezitose, inulin, soluble starch (weak), D-xylose, L-arabinose, D-arabinose (variable), D-ribose, L-rhamnose, D-glucosamine, N-acetyl-D-glucosamine, ethanol, ribitol, galactitol, D-mannitol, D-glucitol, salicin, DL-lactic acid, succinic acid, citric acid, inositol (variable), hexadecane (variable) and D-glucuronic acid are assimilated as sole carbon sources. Methanol, glycerol, erythritol and methyl-α-D-glucoside are not assimilated. Ammonium sulfate, potassium nitrate (variable), sodium nitrite, L-lysine, ethylamine and cadaverine are assimilated as sole nitrogen sources. The maximum growth temperature is 37 °C. Growth in the vitamin-free medium is positive. Starch-like substances are not produced. Growth on 50 % (w/w) glucose-yeast extract agar is negative.

Physiologically, *S. hainanensis* does not differ from the closely related species *S. jianfenglingensis* and *S. wuzhishanensis* (Fig. 5) in its ability to assimilate carbon sources and nitrogen sources (Table S2.4).

###### Typus

**China**, Diaoluoshan, Hainan province, obtained from rotten wood, Jan. 2008, *Q.-M. Wang* (**holotype** CGMCC 2.3787^T^ preserved in a metabolically inactive state).

###### Other culture examined

**China**, Hainan province, obtained from rotten wood, isolated by *Q.M. Wang*, culture CGMCC 2.3782.

***Sugiyamaella jianfenglingensis*** Q.-M. Wang, A.-H. Li, M.-M. Liu, H.-H. Zhu & F.-Y. Bai, ***sp. nov.*** Fungal Names FN 572753. Fig. 32A, B.

**Fig. 32.**
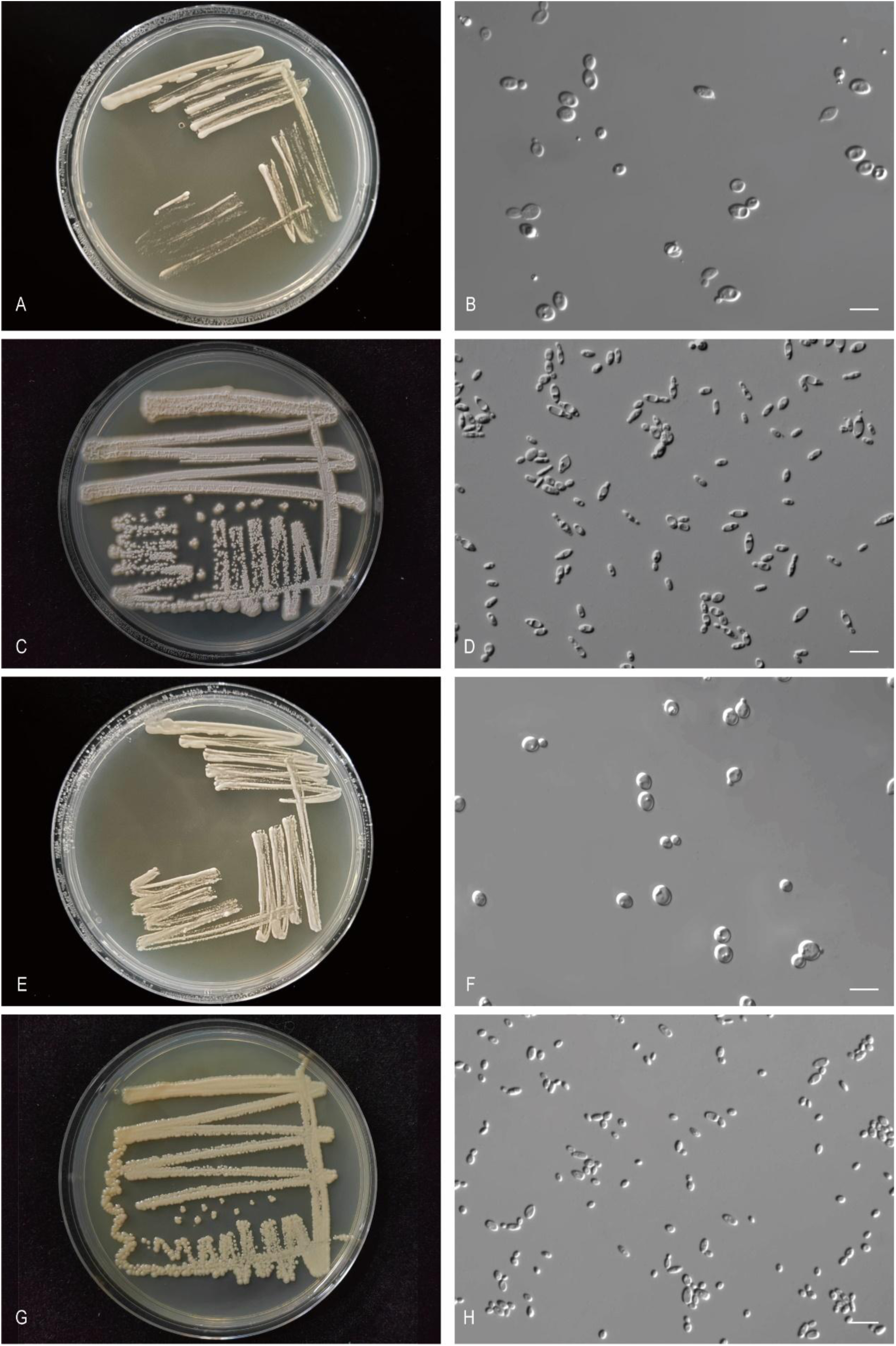
The streak culture grown in YM agar and vegetative cells grown in YM broth for 7 d at 25 °C. A, B. *Sug. jianfenglingensis* CGMCC 2.3786^T^. C, D. *Sug. linzhiensis* CGMCC 2.6975^T^. E, F. *Sug. wuzhishanensis* CGMCC 2.3785^T^. G, H. *Wic. follicola* CGMCC 2.2799^T^. Scale bars: B, D, F, H = 10 μm.

###### Etymology

the specific epithet *jianfenglingensis* refers to the geographic origin of the type strain, Jianfengling mountain, Hainan province.

###### Culture characteristics

On YM agar, after 1 month at 25 °C, the streak culture is whitish-cream, butyrous, rough, smooth and glossy. The margin is entire (Fig. 32A). In YM broth, after 3 days at 25 °C, cells are subglobose and oval, 3.19–5.95 × 2.52–7.02 µm and single, budding is polar (Fig. 32B), after 7 days at 25 °C, a sediment is formed. After 1 month at 25 °C, sediment is present, pellicle and ring do not appear. In Dalmau plate culture on corn meal agar, pseudohyphae are not formed. Asci and ascospores are not observed on V8 and Acetate agar.

###### Physiological and biochemical characteristics

Glucose (weak) fermentation is present. Lactose, galactose, sucrose, maltose and raffinose fermentation are absent. Glucose, galactose, L-sorbose, sucrose, maltose, cellobiose (weak), trehalose, lactose, melibiose, raffinose, melezitose, inulin, soluble starch, D-xylose, L-arabinose, D-arabinose, D-ribose, L-rhamnose, D-glucosamine, N-acetyl-D-glucosamine, ethanol, ribitol, galactitol, D-mannitol, D-glucitol, salicin, DL-lactic acid, succinic acid, citric acid and D-glucuronic acid are assimilated as sole carbon sources. Methanol, glycerol, erythritol, methyl-α-D-glucoside, inositol and hexadecane are not assimilated. Ammonium sulfate (weak), potassium nitrate, sodium nitrite, L-lysine, ethylamine and cadaverine are assimilated as sole nitrogen sources. The maximum growth temperature is 37 °C. Growth in the vitamin-free medium is positive. Starch-like substances are not produced. Growth on 50 % (w/w) glucose-yeast extract agar is negative.

Physiologically, *S. jianfenglingensis* does not differ from the closely related species *S. hainanensis* and *S. wuzhishanensis* (Fig. 5) in its ability to assimilate carbon sources and nitrogen sources (Table S2.4).

###### Typus

**China**, Hainan province, obtained from rotten wood, *Q.-M. Wang* (**holotype** CGMCC 2.3786^T^ preserved in a metabolically inactive state, ex-type cultures are preserved in culture collections under the numbers X4WZ09-17).

***Sugiyamaella linzhiensis*** Q.-M. Wang, A.-H. Li, M.-M. Liu, H.-H. Zhu & F.-Y. Bai, ***sp. nov.*** Fungal Names FN 572754. Fig. 32C, D.

###### Etymology

the specific epithet *linzhiensis* refers to the geographic origin of the type strain, Linzhi county, Tibet.

###### Culture characteristics

On YM agar, after 1 month at 25 °C, the streak culture is white, butyrous, rough, hairlike and pale dimmed with white powder. The margin is fringed (Fig. 32C). In YM broth, after 3 days at 25 °C, cells are ellipsoidal and fusiform, 1.02–3.72 × 4.10–8.76 µm and single, budding is polar (Fig. 32D), after 7 days at 25 °C, a sediment is formed. After 1 month at 25 °C, sediment is present, pellicle and ring does not appear. In Dalmau plate culture on corn meal agar, hyphae are formed. Asci and ascospores are not observed on V8 and Acetate agar.

###### Physiological and biochemical characteristics

Glucose (variable), galactose (variable), sucrose (variable) and raffinose (variable) fermentation are present. Lactose and maltose fermentation are absent. Glucose, galactose, L-sorbose (weak), sucrose, maltose, cellobiose, trehalose (variable), lactose (variable), melibiose (variable), raffinose, melezitose (variable), inulin, soluble starch (weak), D-xylose (latent), L-arabinose (variable), D-arabinose (variable), D-ribose (variable), L-rhamnose (variable), D-glucosamine (latent), N-acetyl-D-glucosamine, ethanol (latent), glycerol (variable), erythritol (variable), ribitol (variable), D-mannitol, D-glucitol (variable), methyl-α-D-glucoside (latent), salicin (latent, weak), DL-lactic acid (variable), succinic acid (variable), citric acid, inositol (variable) and D-glucuronic acid (variable) are assimilated as sole carbon sources. Methanol, galactitol and hexadecane are not assimilated. Ammonium sulfate, potassium nitrate (variable), sodium nitrite (variable), L-lysine (variable), ethylamine and cadaverine are assimilated as sole nitrogen sources. The maximum growth temperature is 37 °C. Growth in the vitamin-free medium is positive (variable). Starch-like substances are not produced. Growth on 50 % (w/w) glucose-yeast extract agar is negative.

Physiologically, *S. linzhiensis* differs from the closely related species *S. paludigena* and *S. castrensis* (Fig. 5) in its ability to assimilate inulin and soluble starch (Table S2.4).

###### Typus

**China**, Chayu County, Linzhi, Tibet, obtained from soil, Aug. 2019, *Q.-M. Wang* (**holotype** CGMCC 2.6975^T^ preserved in a metabolically inactive state, ex-type cultures are preserved in culture collections under the numbers CYA2-1).

###### Other culture examined

**China**, Chayu County, Linzhi, Tibet, obtained from soil, Aug. 2019, isolated by *Q.M. Wang*, culture CGMCC 2.6976 (= CYA2-2); **China**, Milin County, Linzhi, Tibet, obtained from rotten wood, Aug. 2019, isolated by *Q.M. Wang*, culture CGMCC 2.6977 (= LXFM2A2); **China**, Katinggou, Nyingchi City, Tibet, obtained from kuding tea, Aug. 2019, isolated by *Q.M. Wang*, culture XZY191-1; **China**, Milin County, Linzhi, Tibet, obtained from rotten wood, Aug. 2019, isolated by *Q.M. Wang*, culture LXFM5F3. **China**, Milin County, Linzhi, Tibet, obtained from rotten wood, Aug. 2019, isolated by *Q.M. Wang*, culture LXFM4F6. **China**, Basongcuo, Tibet, obtained from pine, Aug. 2019, isolated by *Q.M. Wang*, culture XZY486F1.

***Sugiyamaella wuzhishanensis*** Q.-M. Wang, A.-H. Li, M.-M. Liu, H.-H. Zhu & F.-Y. Bai, ***sp. nov.*** Fungal Names FN 572755. Fig. 32E, F.

###### Etymology

the specific epithet *wuzhishanensis* refers to the geographic origin of the type strain, Wuzhishan mountain, Hainan province.

###### Culture characteristics

On YM agar, after 1 month at 25 °C, the streak culture is white, butyrous, smooth and glossy. The margin is entire (Fig. 32E). In YM broth, after 3 days at 25 °C, cells are subglobose, ellipsoidal and oval, 2.88–6.37 × 4.35–7.42 µm and single, budding is polar (Fig. 32F), after 7 days at 25 °C, a sediment is formed. After 1 month at 25 °C, sediment is present, pellicle and ring do not appear. In Dalmau plate culture on corn meal agar, pseudohyphae are not formed. Asci and ascospores are not observed on V8 and Acetate agar.

###### Physiological and biochemical characteristics

Raffinose (weak) fermentation is present. Glucose, lactose, galactose, sucrose and maltose fermentation are absent. Glucose, galactose (latent, weak), L-sorbose (latent), cellobiose (weak), trehalose, inulin (weak), D-xylose (latent), N-acetyl-D-glucosamine (latent), ethanol (weak), galactitol (weak), D-mannitol, D-glucitol (latent), salicin and succinic acid (latent) are assimilated as sole carbon sources. Sucrose, maltose, lactose, melibiose, raffinose, melezitose, soluble starch, L-arabinose, D-arabinose, D-ribose, L-rhamnose, D-glucosamine, methanol, glycerol, erythritol, ribitol, methyl-α-D-glucoside, DL-lactic acid, citric acid, inositol, hexadecane and D-glucuronic acid are not assimilated. L-lysine, ethylamine (latent) and cadaverine (latent) are assimilated as sole nitrogen sources. Ammonium sulfate, potassium nitrate and sodium nitrite are not assimilated. The maximum growth temperature is 38 °C. Growth in the vitamin-free medium is positive (weak). Starch-like substances are not produced. Growth on 50 % (w/w) glucose-yeast extract agar is positive (weak).

Physiologically, *S. wuzhishanensis* differs from the closely related species *S. hainanensis* and *S. jianfenglingensis* (Fig. 5) in its inability to ferment glucose, the inability to assimilate sucrose, maltose, lactose, melibiose, raffinose, melezitose, soluble starch, L-arabinose, D-arabinose, D-ribose, L-rhamnose, D-glucosamine, ribitol, DL-lactic acid, citric acid, D-glucuronic acid, ammonium sulfate and sodium nitrite, and the ability to ferment raffinose and grow with 50% glucose (Table S2.4).

###### Typus

**China**, Hainan province, obtained from rotten wood, Jan. 2008, *Q.-M. Wang* (**holotype** CGMCC 2.3785^T^ preserved in a metabolically inactive state, ex-type cultures are preserved in culture collections under the numbers X3JF13-2).

***Wickerhamiella follicola*** Q.-M. Wang, A.-H. Li, M.-M. Liu, H.-H. Zhu & F.-Y. Bai, ***sp. nov.*** Fungal Names FN 572756. Fig. 32G, H.

###### Etymology

the specific epithet *follicola* refers to the type strain isolated from the leaf of an unidentified plant.

###### Culture characteristics

On YM agar, after 1 month at 25 °C, the streak culture is yellowish-cream, butyrous, smooth and glossy. The margin is entire (Fig. 32G). In YM broth, after 3 days at 25 °C, cells are oval and fusiform, 1.67–2.58 × 3.16–5.82 µm and single, budding is polar (Fig. 32H), after 7 days at 25 °C, a sediment is formed. After 1 month at 25 °C, sediment and pellicle are present, but a ring does not appear. In Dalmau plate culture on corn meal agar, pseudohyphae are formed. Asci and ascospores are not observed on V8 and Acetate agar.

###### Physiological and biochemical characteristics

Glucose, lactose, galactose, sucrose, maltose and raffinose fermentation are absent. Glucose, galactose, L-sorbose, raffinose, inulin (weak), D-xylose, L-arabinose, ribitol, D-mannitol, D-glucitol, succinic acid and citric acid are assimilated as sole carbon sources.

Sucrose, maltose, cellobiose, trehalose, lactose, melibiose, melezitose, soluble starch, D-arabinose, D-ribose, L-rhamnose, D-glucosamine, N-acetyl-D-glucosamine, methanol, ethanol, glycerol, erythritol, galactitol, methyl-α-D-glucoside, salicin, DL-lactic acid, inositol, hexadecane and D-glucuronic acid are not assimilated. Ammonium sulfate, potassium nitrate, sodium nitrite (latent), L-lysine, ethylamine and cadaverine are assimilated as sole nitrogen sources. The maximum growth temperature is 37 °C. Growth in the vitamin-free medium is positive. Starch-like substances are not produced. Growth on 50 % (w/w) glucose-yeast extract agar is negative.

Physiologically, *W. follicola* differs from the closely related species *W. natalensis* (Fig. 6) in its inability to assimilate ethanol, glycerol, and grow with 50% glucose and the ability to assimilate galactose, raffinose, inulin, D-xylose, L-arabinose, ribitol, and grow in vitamin-free medium (Table S2.5).

###### Typus

**China**, Tibet, obtained from leaf, Apr. 2005, *Q.-M. Wang* (**holotype** CGMCC 2.2799^T^ preserved in a metabolically inactive state, ex-type cultures are preserved in culture collections under the numbers CBS 18409).

###### Other culture examined

**China**, Tibet, obtained from leaf, Apr. 2005, isolated by *Q.M. Wang*, culture XZ69B1.

***Wickerhamiella insecticola*** Q.-M. Wang, A.-H. Li, M.-M. Liu, H.-H. Zhu & F.-Y. Bai, ***sp. nov.*** Fungal Names FN 572757. Fig. 33A, B.

**Fig. 33.**
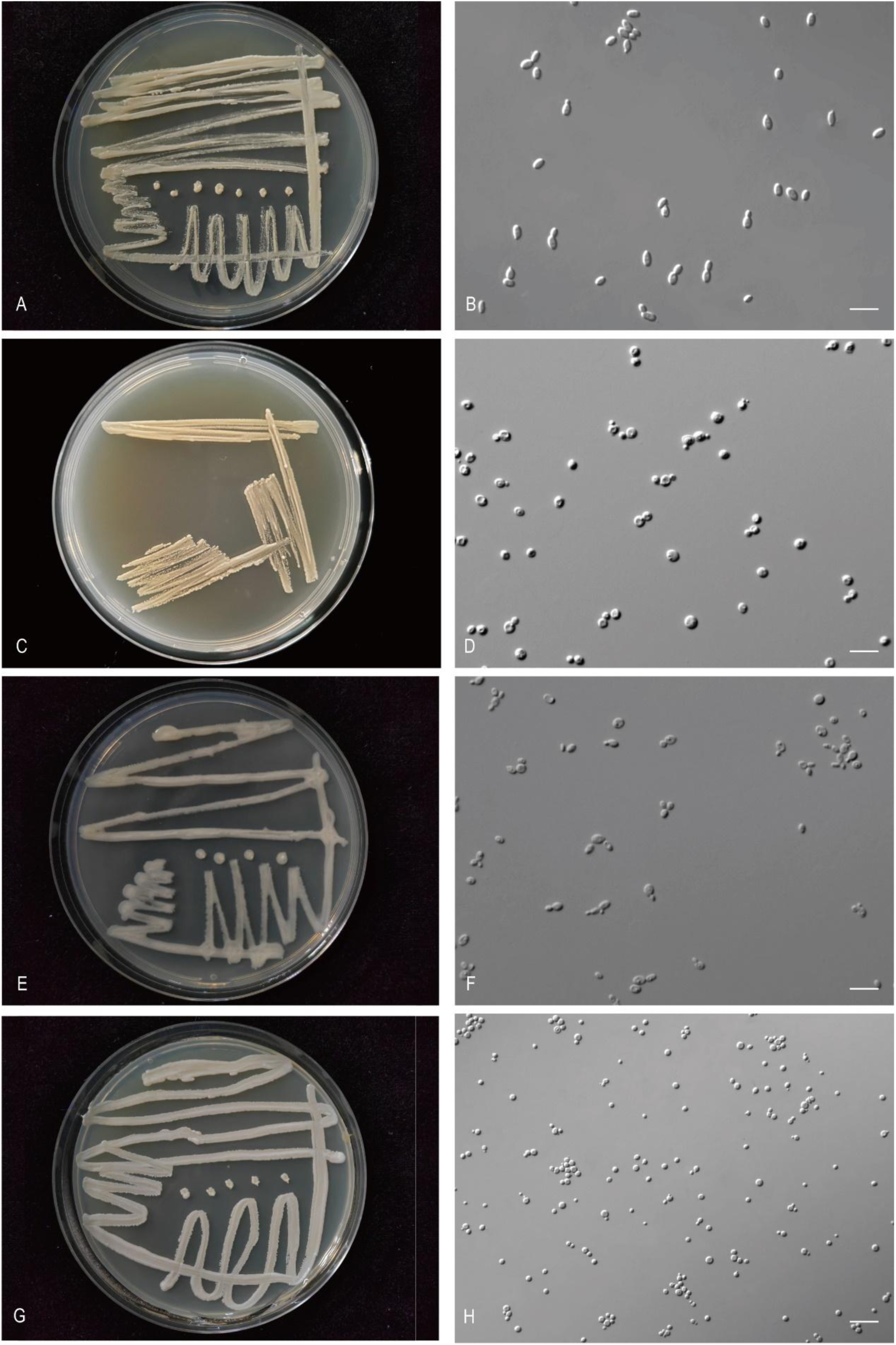
The streak culture grown in YM agar and vegetative cells grown in YM broth for 7 d at 25 °C. A, B. *Wic. insecticola* CGMCC 2.3487^T^. C, D. *Nak. putredosilvae* CGMCC 2.6987^T^. E, F. *Nak. Quercae* CGMCC 2.6985^T^. G, H. *Kom. corticis* CGMCC 2.8850^T^. Scale bars: B, D, F, H = 10 μm.

###### Etymology

the specific epithet *insecticola* refers to the type strain isolated from insect.

###### Culture characteristics

On YM agar, after 1 month at 25 °C, the streak culture is yellowish-cream, butyrous, smooth and glossy. The margin is fringed (Fig. 33A). In YM broth, after 3 days at 25 °C, cells are ellipsoidal and fusiform, 2.06–3.05 × 2.93–5.28 µm and single, budding is polar (Fig. 33B), after 7 days at 25 °C, a sediment is formed. After 1 month at 25 °C, sediment and ring are present, but a pellicle does not appear. In Dalmau plate culture on corn meal agar, pseudohyphae are formed. Asci and ascospores are not observed on V8 and Acetate agar.

###### Physiological and biochemical characteristics

Glucose, lactose, galactose, sucrose, maltose and raffinose fermentation are absent. Glucose, galactose, L-sorbose, melibiose, raffinose, inulin, soluble starch (weak), D-xylose, glycerol, D-mannitol, D-glucitol and succinic acid are assimilated as sole carbon sources.

Sucrose, maltose, cellobiose, trehalose, lactose, melezitose, L-arabinose, D-arabinose, D-ribose, L-rhamnose, D-glucosamine, N-acetyl-D-glucosamine, methanol, ethanol, erythritol, ribitol, galactitol, methyl-α-D-glucoside, salicin, DL-lactic acid, citric acid, inositol, hexadecane and D-glucuronic acid are not assimilated. Ammonium sulfate, potassium nitrate, sodium nitrite, L-lysine, ethylamine and cadaverine are assimilated as sole nitrogen sources. The maximum growth temperature is 37 °C. Growth in the vitamin-free medium is positive. Starch-like substances are not produced. Growth on 50 % (w/w) glucose-yeast extract agar is negative.

Physiologically, *W. insecticola* differs from the closely related species *W. slavikovae*, *W. hasegawae*, *W. fructicola*, *W. kazuoi*, *W. qilinensis* and *W. pterostichi* (Fig. 6) in its ability to assimilate melibiose, raffinose, soluble starch, potassium nitrate, and sodium nitrite (Table S2.5).

###### Typus

**China**, Shaanxi province, obtained from an insect, Apr. 2007, *Q.-M. Wang* (**holotype** CGMCC 2.3487^T^ preserved in a metabolically inactive state).

###### New taxa in the Alaninaceae (Alaninales, Pichiomycetes, Saccharomycotina)

***Nakazawaea putredisilvae*** Q.-M. Wang, A.-H. Li, M.-M. Liu, H.-H. Zhu & F.-Y. Bai, ***sp. nov.*** Fungal Names FN 572758. Fig. 33C, D.

###### Etymology

the specific epithet *putredisilvae* refers to the type strain isolated from rotten wood.

###### Culture characteristics

On YM agar, after 1 month at 25 °C, the streak culture is white-cream, butyrous, smooth and pale glossy. The margin is entire (Fig. 33C). In YM broth, after 3 days at 25 °C, cells are globular and subglobose, 2.25–4.21 × 3.27–5.06 µm and single, budding is polar (Fig. 33D), after 7 days at 25 °C, a sediment is formed. After 1 month at 25 °C, sediment and ring are present, but a pellicle does not appear. In Dalmau plate culture on corn meal agar, pseudohyphae are not formed. Asci and ascospores are not observed on V8 and Acetate agar.

###### Physiological and biochemical characteristics

Glucose fermentation is present. Lactose, galactose, sucrose, maltose and raffinose fermentation are absent. Glucose, L-sorbose (latent), sucrose (variable), cellobiose (latent), trehalose, inulin (latent, weak), D-xylose (latent, weak), L-arabinose (latent, weak), D-arabinose (variable), D-ribose (latent), L-rhamnose (latent, weak), N-acetyl-D-glucosamine, ethanol (latent), glycerol, ribitol (latent), D-mannitol, D-glucitol (latent), salicin and succinic acid (variable) are assimilated as sole carbon sources. Galactose, maltose, lactose, melibiose, raffinose, melezitose, soluble starch, D-glucosamine, methanol, erythritol, galactitol, methyl-α-D-glucoside, DL-lactic acid, citric acid, inositol, hexadecane and D-glucuronic acid are not assimilated. Ammonium sulfate, potassium nitrate, sodium nitrite, L-lysine, ethylamine and cadaverine are assimilated as sole nitrogen sources. The maximum growth temperature is 32 °C. Growth in the vitamin-free medium is positive. Starch-like substances are not produced. Growth on 50 % (w/w) glucose-yeast extract agar is negative.

Physiologically, *N. putredosilvae* differs from the closely related species *N. wickerhamii*, *N. anatomiae* and *N. populi* (Fig. 7) in its inability to assimilate citric acid and the ability to assimilate L-sorbose, inulin, and grow in vitamin-free medium (Table S2.6).

###### Typus

**China**, Sejila Mountain, Nyingchi, Tibet, obtained from rotten wood, Aug. 2019, *Q.-M. Wang* (**holotype** CGMCC 2.6987^T^ preserved in a metabolically inactive state, ex-type culture is preserved in culture collections under the numbers XZY879-1).

###### Other culture examined

**China**, Sejila Mountain, Nyingchi, Tibet, obtained from rotten wood, Aug. 2019, isolated by *Q.-M. Wang*, *G.-S. Wang*, culture XZY867-1.

***Nakazawaea quercae*** Q.-M. Wang, A.-H. Li, M.-M. Liu, H.-H. Zhu & F.-Y. Bai, ***sp. nov.*** Fungal Names FN 572759. Fig. 33E, F.

###### Etymology

the specific epithet *quercae* refers to *Quercus*, the plant genus from which the type strain was isolated.

###### Culture characteristics

On YM agar, after 1 month at 25 °C, the streak culture is white, butyrous, smooth and glossy. The margin is entire (Fig. 33E). In YM broth, after 3 days at 25 °C, cells are fusiform, 1.17–3.27 × 2.28–4.33 µm and single, budding is polar (Fig. 33F), after 7 days at 25 °C, a sediment is formed. After 1 month at 25 °C, sediment and ring are present, but a pellicle does not appear. In Dalmau plate culture on corn meal agar, pseudohyphae are not formed. Asci and ascospores are not observed on V8 and Acetate agar.

###### Physiological and biochemical characteristics

Glucose, sucrose (variable) and maltose (variable) fermentation are present. Lactose, galactose and raffinose fermentation are absent. Glucose, galactose (latent, weak), L-sorbose, sucrose (weak), maltose, cellobiose, trehalose, melibiose (variable), raffinose (variable), melezitose, inulin, soluble starch (latent), D-xylose (latent, weak), L-arabinose (variable), D-arabinose (variable), D-ribose, L-rhamnose (latent, weak), D-glucosamine (variable), N-acetyl-D-glucosamine (latent), ethanol (latent), glycerol, ribitol (latent, weak), D-mannitol, D-glucitol (latent),methyl-α-D-glucoside (variable), salicin, DL-lactic acid (variable), succinic acid (latent, weak), citric acid and inositol (variable) are assimilated as sole carbon sources. Lactose, methanol, erythritol, galactitol, hexadecane and D-glucuronic acid are not assimilated. Ammonium sulfate, potassium nitrate, sodium nitrite, L-lysine, ethylamine and cadaverine are assimilated as sole nitrogen sources. The maximum growth temperature is 35 °C. Growth in the vitamin-free medium is positive. Starch-like substances are not produced. Growth on 50 % (w/w) glucose-yeast extract agar is positive.

Physiologically, *N. quercae* does not differ from the closely related species *N. holstii*, *N. ernobii* and *N. tricholomae* (Fig. 7) in its ability to assimilate carbon sources and nitrogen sources (Table S2.6).

###### Typus

**China**, Katinggou, Nyingchi, Tibet, obtained from the leaf of *Quercus semicarpifolia*, Aug. 2019, *Q.-M. Wang* (**holotype** CGMCC 2.6985^T^ preserved in a metabolically inactive state, ex-type cultures are preserved in culture collections under the numbers XZY20F1).

###### Other culture examined

**China**, Sejila Mountain, Nyingchi, Tibet, obtained from leaf of *Caprifoliaceae*, Aug. 2019, isolated by *Q.M. Wang*, culture XZY163-6. **China**, Sejila Mountain, Nyingchi, Tibet, obtained from leaf of *Caprifoliaceae*, Aug. 2019, isolated by *Q.M. Wang*, culture XZY163-1. **China**, Baiba, Nyingchi, Tibet, obtained from mushroom, Aug. 2019, isolated by *Q.M. Wang*, culture XZY95-3. **China**, Basongcuo, Nyingchi, Tibet, obtained from birch trees, Aug. 2019, isolated by *Q.M. Wang*, culture XZY137-5.

###### New taxa in the Pichiaceae (Pichiales, Pichiomycetes, Saccharomycotina)

***Komagataella corticis*** Q.-M. Wang, A.-H. Li, M.-M. Liu, H.-H. Zhu & F.-Y. Bai, ***sp. nov.*** Fungal Names FN 572760. Fig. 33G, H.

###### Etymology

the specific epithet *corticis* refers to the type strain isolated from bark.

###### Culture characteristics

On YM agar, after 1 month at 25 °C, the streak culture is ivory, butyrous, and smooth. The margin is entire (Fig. 33G). In YM broth, after 3 days at 25 °C, cells are globular and subglobose, 2.53–4.86 × 2.69–6.98 µm and single, budding is polar (Fig. 33H), after 7 days at 25 °C, a sediment is formed. After 1 month at 25 °C, sediment is present, but a ring and pellicle do not appear. In Dalmau plate culture on corn meal agar, pseudohyphae are not formed. Asci and ascospores are not observed on V8 and Acetate agar.

###### Physiological and biochemical characteristics

Glucose and raffinose (variable) fermentation are present. Lactose, galactose, sucrose and maltose fermentation are absent. Glucose, galactose, L-sorbose (latent, weak), sucrose (variable), maltose, cellobiose, trehalose, melezitose (variable), inulin, soluble starch (latent, weak), D-xylose (latent), D-ribose (variable), L-rhamnose (latent), D-glucosamine (variable), N-acetyl-D-glucosamine (latent), methanol (latent), ethanol, glycerol, ribitol (variable), galactitol (variable), D-mannitol, D-glucitol (latent, weak), methyl-α-D-glucoside (variable), salicin (variable), DL-lactic acid (variable), succinic acid (latent, weak), citric acid (variable) and D-glucuronic acid (variable) are assimilated as sole carbon sources. Lactose, melibiose, raffinose, L-arabinose, D-arabinose, erythritol, inositol and hexadecane are not assimilated. Ammonium sulfate, potassium nitrate, sodium nitrite (latent, weak), L-lysine (latent), ethylamine (latent) and cadaverine are assimilated as sole nitrogen sources. The maximum growth temperature is 40 °C. Growth in the vitamin-free medium is positive. Starch-like substances are not produced. Growth on 50 % (w/w) glucose-yeast extract agar is negative.

Physiologically, *K. corticis* differs from the closely related species *K. pastoris*, *K. ulmi*, *K. phaffii*, *K. kurtzmanii*, *K. pseudopastoris*, *K. populi* and *K. mondaviorum* (Fig. 8) in its ability to assimilate N-acetyl-D-glucosamine, potassium nitrate, and grow in vitamin-free medium (Table S2.7).

###### Typus

**China**, Shengfeng Mountains Heilongjiang Province, obtained from bark, Nov. 2017, *Q.-M. Wang* (**holotype** CGMCC 2.8850^T^ preserved in a metabolically inactive state, ex-type cultures are preserved in culture collections under the numbers HSP262-1).

###### Other culture examined

**China**, Shengfeng Mountains Heilongjiang Province, obtained from bark, Nov. 2017, isolated by *Q.M. Wang*, culture HSP275-3. **China**, Shengfeng Mountains Heilongjiang Province, obtained from bark, Nov. 2017, isolated by *Q.M. Wang*, culture HSP262-2.

***Ogataea caprifoliacea*** Q.-M. Wang, A.-H. Li, M.-M. Liu, H.-H. Zhu & F.-Y. Bai, ***sp. nov.*** Fungal Names FN 572761. Fig. 34A, B.

**Fig. 34.**
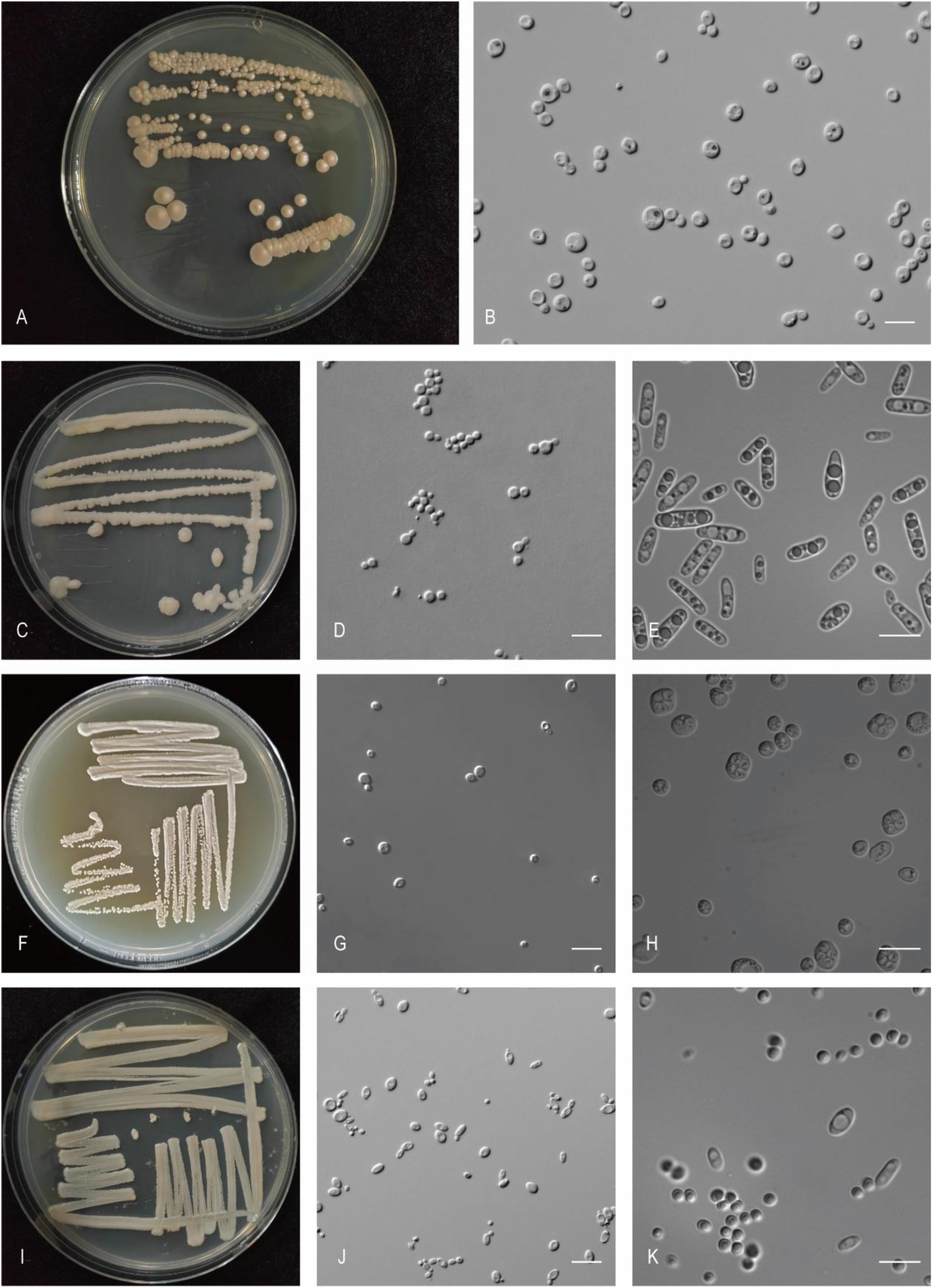
The streak culture grown in YM agar and vegetative cells grown in YM broth for 7 d at 25 °C. A, B. *Oga. caprifoliacea* CGMCC 2.6982^T^. C-E. *Oga. linzhienesis* CGMCC 2.6980^T^. F-H. *Pic. Sinensis* CGMCC 2.3286^T^. I-K. *Pic. humicola* CGMCC 2.6999^T^. Scale bars: B, D, E, G, H, J, K = 10 μm.

###### Etymology

the specific epithet *caprifoliacea* refers to *Caprifoliacea*, the plant family from which the type strain was isolated.

###### Culture characteristics

On YM agar, after 1 month at 25 °C, the streak culture is yellowish, butyrous, smooth and pale glossy. The margin is entire (Fig. 34A). In YM broth, after 3 days at 25 °C, cells are ellipsoidal, globular and subglobose, 3.00–6.81 × 3.66–8.42 µm and single, budding is polar (Fig. 34B), after 7 days at 25 °C, a sediment is formed. After 1 month at 25 °C, sediment is present, but a ring and pellicle do not appear. In Dalmau plate culture on corn meal agar, pseudohyphae are not formed. Asci and ascospores are not observed on V8 and Acetate agar.

###### Physiological and biochemical characteristics

Glucose, sucrose fermentation are present. Lactose, galactose, maltose and raffinose fermentation are absent. Glucose, galactose (variable), L-sorbose (latent, weak), sucrose (latent), maltose (latent), cellobiose (latent), trehalose (latent), lactose (variable), melibiose (variable), raffinose (latent), inulin (variable), soluble starch (variable), D-xylose (variable), L-arabinose (latent), D-ribose (weak), N-acetyl-D-glucosamine (variable), methanol (latent, weak), ethanol (latent), glycerol (variable), erythritol (variable), ribitol (latent, weak), D-mannitol, D-glucitol (variable) and salicin (latent) are assimilated as sole carbon sources. Melezitose, D-arabinose, L-rhamnose, D-glucosamine, galactitol, methyl-α-D-glucoside, DL-lactic acid, succinic acid, citric acid, inositol, hexadecane and D-glucuronic acid are not assimilated. Ammonium sulfate, potassium nitrate (variable), L-lysine (latent, weak), ethylamine and cadaverine (latent) are assimilated as sole nitrogen sources. Sodium nitrite is not assimilated. The maximum growth temperature is 25 °C. Growth in the vitamin-free medium is positive (latent). Starch-like substances are not produced. Growth on 50 % (w/w) glucose-yeast extract agar is negative.

Physiologically, *O. caprifoliacea* differs from the closely related species *O. nitratoaversa*, *O. pilisensis*, *O. piceae* and *O. sonorensis* (Fig. 9) in its ability to ferment sucrose, the ability to assimilate L-sorbose, raffinose, and grow in vitamin-free medium (Table S2.8).

###### Typus

**China**, Katinggou, Nyingchi city, Tibet, obtained from the leaf of *Caprifoliaceae*, Aug. 2019, *Q.-M. Wang* (**holotype** CGMCC 2.6982^T^ preserved in a metabolically inactive state, ex-type cultures are preserved in culture collections under the numbers XZY167-1).

###### Other culture examined

**China**, Sejila Mountain, Nyingchi, Tibet, obtained from rotten wood, Aug. 2019, isolated by *Q.M. Wang*, culture XZY166-1.

***Ogataea linzhienesis*** Q.-M. Wang, A.-H. Li, M.-M. Liu, H.-H. Zhu & F.-Y. Bai, ***sp. nov.*** Fungal Names FN 572762. Fig. 34C-E.

###### Etymology

the specific epithet *linzhienesis* refers to the geographic origin of the type strain, Linzhi, Tibet.

###### Culture characteristics

On YM agar, after 1 month at 25 °C, the streak culture is whitish-cream, butyrous, smooth and glossy. The margin is entire (Fig. 34C). In YM broth, after 3 days at 25 °C, cells are ellipsoidal and oval, 2.56–4.18 × 2.92–4.47 µm and single, budding is polar (Fig. 34D), after 7 days at 25 °C, a sediment is formed. After 1 month at 25 °C, sediment is present, but a ring and pellicle do not appear. In Dalmau plate culture on corn meal agar, pseudohyphae are not formed. On V8 agar, the ascus is fusiform and contains two ascospores (Fig. 34E). The ascospores are spherical or subspherical, 1.55–3.73 × 1.64–4.26 µm.

###### Physiological and biochemical characteristics

Glucose, lactose, galactose, sucrose, maltose and raffinose fermentation are absent. Glucose, L-sorbose (latent), cellobiose (latent), inulin (latent), D-xylose (latent, weak), D-ribose(latent), methanol (latent, weak), ethanol (latent), glycerol (latent), erythritol (latent), ribitol (latent), D-mannitol (latent), D-glucitol (latent) and salicin (latent) are assimilated as sole carbon sources. Galactose, sucrose, maltose, trehalose, lactose, melibiose, raffinose, melezitose, soluble starch, L-arabinose, D-arabinose, L-rhamnose, D-glucosamine, N-acetyl-D-glucosamine, galactitol, methyl-α-D-glucoside, DL-lactic acid, succinic acid, citric acid, inositol, hexadecane and D-glucuronic acid are not assimilated. Ammonium sulfate, potassium nitrate, L-lysine, ethylamine and cadaverine are assimilated as sole nitrogen sources. Sodium nitrite is not assimilated. The maximum growth temperature is 35 °C. Growth in the vitamin-free medium is positive. Starch-like substances are not produced. Growth on 50 % (w/w) glucose-yeast extract agar is negative.

Physiologically, *O. linzhienesis* differs from the closely related species *O. paradorogensis* and *O. xyloterini* (Fig. 9) in its inability to assimilate trehalose and the ability to assimilate inulin, potassium nitrate, and grow in vitamin-free medium (Table S2.8).

###### Typus

**China**, Gongbujiangda, Nyingchi, Tibet, obtained from rotten wood, Aug. 2019, *Q.-M. Wang* (**holotype** CGMCC 2.6980^T^ preserved in a metabolically inactive state, ex-type cultures are preserved in culture collections under the numbers XZY775-1).

###### Other culture examined

**China**, Sejila mountain, Nyingchi, Tibet, obtained from lichen, Aug. 2019, isolated by *Q.M. Wang*, culture XZY267-1.

***Pichia sinensis*** Q.-M. Wang, A.-H. Li, M.-M. Liu, H.-H. Zhu & F.-Y. Bai, ***sp. nov.*** Fungal Names FN 572763. Fig. 34F-H.

###### Etymology

the specific epithet *sinensis* refers to the geographic origin of the type strain, Shaanxi province, China.

###### Culture characteristics

On YM agar, after 1 month at 25 °C, the streak culture is white, butyrous and smooth. The margin is eroded (Fig. 34F). In YM broth, after 3 days at 25 °C, cells are globose and oval, 3.01–5.70 × 3.03–5.39 µm and single, budding is polar (Fig. 34G), after 7 days at 25 °C, a sediment is formed. After 1 month at 25 °C, sediment is present, the ring and a pellicle do not appear. In Dalmau plate culture on corn meal agar, pseudohyphae are not formed. On Acetate agar, the ascus is oval and contains four ascospores (Fig. 34H). The ascospores are spherical or subspherical, 1.96–2.45 × 1.88–2.64 µm.

###### Physiological and biochemical characteristics

Glucose, galactose (latent), sucrose (weak) and maltose fermentation are present. Lactose and raffinose fermentation are absent. Glucose, galactose, sucrose, raffinose (latent), melezitose, inulin (latent, weak), ethanol (weak), glycerol (weak), galactitol (weak), D-mannitol, methyl-α-D-glucoside and succinic acid (latent, weak) are assimilated as sole carbon sources. L-sorbose, maltose, cellobiose, trehalose, lactose, melibiose, soluble starch, D-xylose, L-arabinose, D-arabinose, D-ribose, L-rhamnose, D-glucosamine, N-acetyl-D-glucosamine, methanol, erythritol, ribitol, D-glucitol, salicin, DL-lactic acid, citric acid, inositol, hexadecane and D-glucuronic acid are not assimilated. Ammonium sulfate (latent), potassium nitrate (latent, weak), L-lysine, ethylamine and cadaverine are assimilated as sole nitrogen sources. Sodium nitrite is not assimilated. The maximum growth temperature is 38 °C. Growth in the vitamin-free medium is positive. Starch-like substances are not produced. Growth on 50 % (w/w) glucose-yeast extract agar is positive.

Physiologically, *P. sinensis* differs from the closely related species *P. putredosilvae*, *P. paraexigua*, *P. bovicola*, *P. exigua*, *P. scutulata* and *P. rugopelliculosa* (Fig. 10) in its ability to ferment galactose, sucrose and maltose (Table S2.9).

###### Typus

**China**, Shaanxi Province, obtained from persimmon, *Q.-M. Wang* (**holotype** CGMCC 2.3286^T^ preserved in a metabolically inactive state).

***Pichia humicola*** Q.-M. Wang, A.-H. Li, M.-M. Liu, H.-H. Zhu & F.-Y. Bai, ***sp. nov.*** Fungal Names FN 572764. Fig. 34I-K.

###### Etymology

the specific epithet *humicola* refers to the type strain isolated from soil.

###### Culture characteristics

On YM agar, after 1 month at 25 °C, the streak culture is white to yellowish, butyrous and smooth. The margin is entire (Fig. 34I). In YM broth, after 3 days at 25 °C, cells are oval and fusiform, 1.05–3.22 × 3.11–5.53 µm and single, budding is polar (Fig. 34J), after 7 days at 25 °C, a sediment and a ring are formed. After 1 month at 25 °C, sediment is present, pellicle and ring do not appear. In Dalmau plate culture on corn meal agar, hyphae are formed. On Acetate agar, the ascus is oval and contains one ascospore (Fig. 34K). The ascospores are spherical or subspherical, 1.78–2.26 × 1.68–2.45 µm.

###### Physiological and biochemical characteristics

Glucose fermentation is present. Lactose, galactose, sucrose, maltose and raffinose fermentation are absent. Glucose, galactose, L-sorbose (variable), sucrose (latent), maltose (variable), cellobiose (latent), trehalose (variable), lactose (variable), melibiose (variable), raffinose (variable), melezitose (latent), inulin, soluble starch (variable), D-xylose, L-arabinose (variable), D-arabinose (variable), D-ribose (variable), L-rhamnose (variable), D-glucosamine (latent), N-acetyl-D-glucosamine (latent, weak), ethanol, glycerol (variable), erythritol (variable), ribitol (variable), galactitol (variable), D-mannitol, D-glucitol (variable), methyl-α-D-glucoside (variable), salicin (variable), DL-lactic acid, succinic acid (variable), inositol (variable) and D-glucuronic acid (variable) are assimilated as sole carbon sources. Methanol, citric acid and hexadecane are not assimilated. Ammonium sulfate, potassium nitrate, sodium nitrite, L-lysine, ethylamine and cadaverine are assimilated as sole nitrogen sources. The maximum growth temperature is 42 °C. Growth in the vitamin-free medium is positive. Starch-like substances are not produced. Growth on 50 % (w/w) glucose-yeast extract agar is negative.

Physiologically, *P. humicola* differs from the closely related species *P. pseudolambica* (Fig. 10) in its ability to assimilate galactose, sucrose, cellobiose, melezitose, inulin, D-mannitol, potassium nitrate and sodium nitrite (Table S2.9).

###### Typus

**China**, Longzi County, Shannan, Tibet, obtained from soil, Aug. 2019, *Q.-M. Wang* (**holotype** CGMCC 2.6999^T^ preserved in a metabolically inactive state, ex-type cultures are preserved in culture collections under the numbers XZ1989-1).

###### Other culture examined

**China**, Jiangzi County, Shigatze City, Tibet, obtained from soil, Aug. 2019, isolated by *Q.M. Wang*, culture CGMCC 2.6940 (= XZ18-36F1A). **China**, Longzi County, Shannan, Tibet, obtained from soil, Aug. 2019, isolated by *Q.M. Wang*, culture XZ1989-2. **China**, Kadinggou, Nyingchi City, Tibet, obtained from soil, Aug. 2019, isolated by *Q.M. Wang*, culture XZ186-3.

***Pichia planticola*** Q.-M. Wang, A.-H. Li, M.-M. Liu, H.-H. Zhu & F.-Y. Bai, *sp. nov.* Fungal Names FN 572765. Fig. 35A-C.

**Fig. 35.**
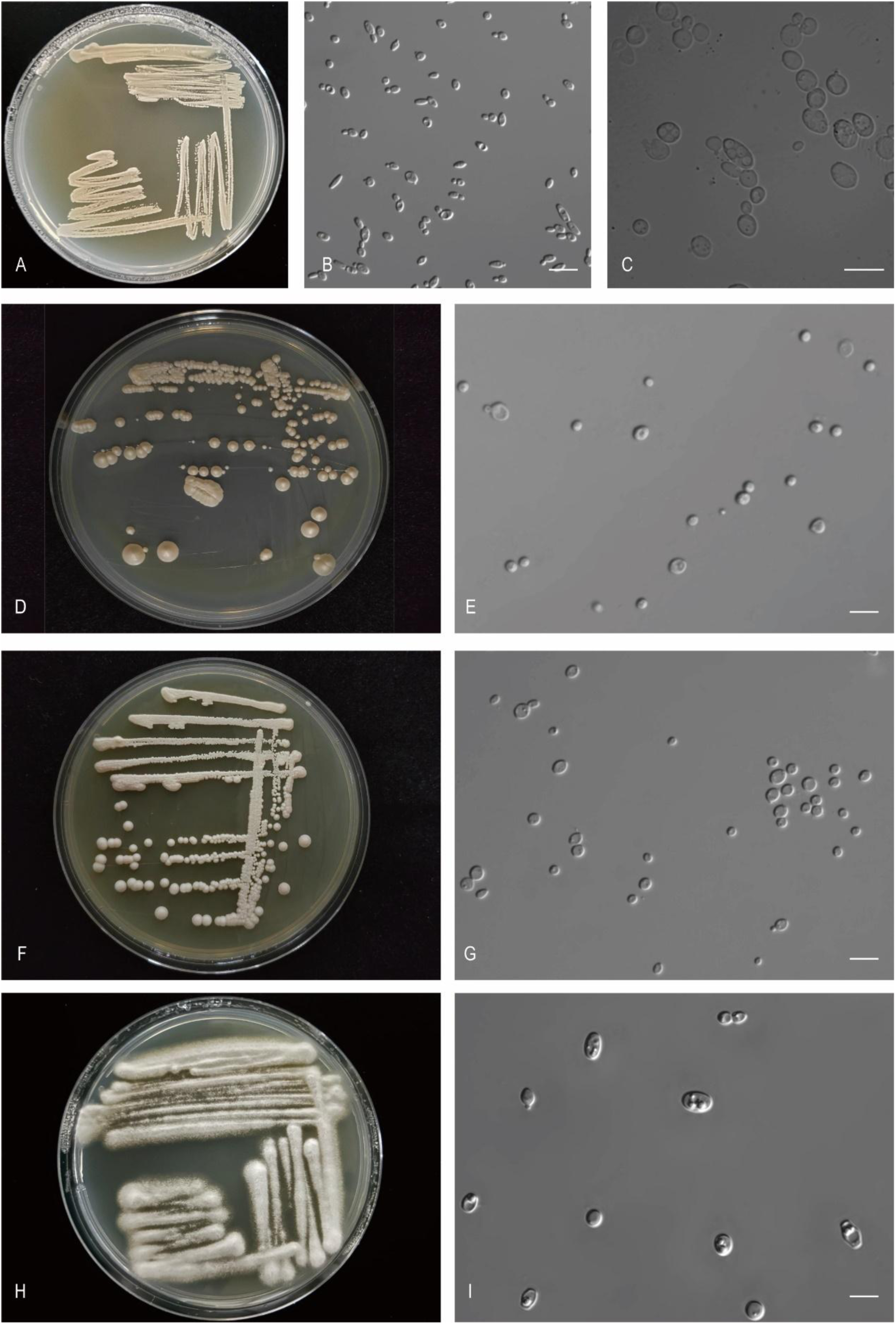
The streak culture grown in YM agar and vegetative cells grown in YM broth for 7 d at 25 °C. A-C. *Pic. planticola* CGMCC 2.3930^T^. D, E. *Pic. putredosilvae* CGMCC 2.7000^T^. F, G. *Pic. Xizangensis* CGMCC 2.7001^T^. H, I. *Sat. jianfenglinensis* CGMCC 2.3783^T^. Scale bars: B, C, E, G, I = 10 μm.

###### Etymology

the specific epithet *planticola* refers to the type strain isolated from a plant.

###### Culture characteristics

On YM agar, after 1 month at 25 °C, the streak culture is white, butyrous. The margin is fimbriate (Fig. 35A). In YM broth, after 3 days at 25 °C, cells are cylindrical, fusiform, ellipsoidal and oval, 2.64–4.30 × 3.86–6.69 µm and single, budding is polar (Fig. 35B), after 7 days at 25 °C, a sediment is formed. After 1 month at 25 °C, sediment is present, the ring and a pellicle do not appear. In Dalmau plate culture on corn meal agar, pseudohyphae are not formed. On Acetate agar, the ascus is fusiform and contains four ascospores (Fig. 35C). The ascospores are spherical or subspherical, 1.60–3.06 × 1.87–3.03 µm.

###### Physiological and biochemical characteristics

Glucose, lactose, galactose, sucrose, maltose and raffinose fermentation are absent. Glucose, galactose, L-sorbose, sucrose, maltose (latent, weak), trehalose (variable), melibiose (variable), melezitose (weak), inulin (latent), soluble starch (latent), D-xylose, D-arabinose (variable), D-ribose (latent), L-rhamnose (variable), D-glucosamine, N-acetyl-D-glucosamine, methanol (variable), ethanol, glycerol (variable), erythritol (variable), ribitol (variable), galactitol (latent, weak), D-mannitol, D-glucitol, methyl-α-D-glucoside (variable), salicin (latent, weak), DL-lactic acid, succinic acid and citric acid are assimilated as sole carbon sources. Cellobiose, lactose, raffinose, L-arabinose, inositol, hexadecane and D-glucuronic acid are not assimilated. Ammonium sulfate, potassium nitrate, sodium nitrite (variable), L-lysine, ethylamine and cadaverine are assimilated as sole nitrogen sources. The maximum growth temperature is 42 °C. Growth in the vitamin-free medium is positive. Starch-like substances are not produced. Growth on 50 % (w/w) glucose-yeast extract agar is negative.

Physiologically, *P. planticola* differs from the closely related species *P. xizangensis*, *P. manshurica*, *P. garciniae* and *P. membranifaciens* (Fig. 10) in its ability to assimilate D-ribose (Table S2.9).

###### Typus

**China**, Jilin Province, obtained from plant, *Q.- M. Wang* (**holotype** CGMCC 2.3930^T^ preserved in a metabolically inactive state).

###### Other culture examined

**China**, Jilin Province, obtained from plant, isolated by *Q.M. Wang*, culture CGMCC 2.3918.

***Pichia putredisilvae*** Q.-M. Wang, A.-H. Li, M.-M. Liu, H.-H. Zhu & F.-Y. Bai, ***sp. nov.*** Fungal Names FN 572766. Fig. 35D, E.

###### Etymology

the specific epithet *putredisilvae* refers to the type strain isolated from rotten wood.

###### Culture characteristics

On YM agar, after 1 month at 25 °C, the streak culture is white, butyrous, smooth and pale glossy. The margin is entire (Fig. 35D). In YM broth, after 3 days at 25 °C, cells are oval and fusiform, 2.61–6.65 × 2.90–7.09 µm and single, budding is polar (Fig. 35E), after 7 days at 25 °C, a sediment is formed. After 1 month at 25 °C, sediment is present, but a pellicle and a ring do not appear. In Dalmau plate culture on corn meal agar, pseudohyphae are not formed. Asci and ascospores are not observed on V8 and Acetate agar.

###### Physiological and biochemical characteristics

Glucose fermentation is present. Lactose, galactose, sucrose, maltose and raffinose fermentation are absent. Glucose, raffinose (latent, weak), D-ribose (latent), ethanol and D-mannitol are assimilated as sole carbon sources. Galactose, L-sorbose, sucrose, maltose, cellobiose, trehalose, lactose, melibiose, melezitose, inulin, soluble starch, D-xylose, L-arabinose, D-arabinose, L-rhamnose, D-glucosamine, N-acetyl-D-glucosamine, methanol, glycerol, erythritol, ribitol, galactitol, D-glucitol, methyl-α-D-glucoside, salicin, DL-lactic acid, succinic acid, citric acid, inositol, hexadecane and D-glucuronic acid are not assimilated. Ammonium sulfate, potassium nitrate, sodium nitrite, L-lysine, ethylamine and cadaverine are assimilated as sole nitrogen sources. The maximum growth temperature is 32 °C. Growth in the vitamin-free medium is positive (latent). Starch-like substances are not produced. Growth on 50 % (w/w) glucose-yeast extract agar is negative.

Physiologically, *P. putredosilvae* does not differ from the closely related species *P. sinensis*, *P. paraexigua*, *P. bovicola*, *P. exigua*, *P. scutulata* and *P. rugopelliculosa* (Fig. 10) in its ability to assimilate carbon sources and nitrogen sources (Table S2.9).

###### Typus

**China**, Basongcuo Lake, Nyingchi, Tibet, obtained from rotten wood, Aug. 2019, *Q.-M. Wang* (**holotype** CGMCC 2.7000^T^ preserved in a metabolically inactive state, ex-type cultures are preserved in culture collections under the numbers XZY838-3).

***Pichia xizangensis*** Q.-M. Wang, A.-H. Li, M.-M. Liu, H.-H. Zhu & F.-Y. Bai, ***sp. nov.*** Fungal Names FN 572767. Fig. 35F, G.

###### Etymology

the specific epithet *xizangensis* refers to the geographic origin of the type strain, Xizang.

###### Culture characteristics

On YM agar, after 1 month at 25 °C, the streak culture is white to cream. The margin is entire (Fig. 35F). In YM broth, after 3 days at 25 °C, cells are oval and fusiform, 2.92–4.95 × 3.97–6.83 µm and single, budding is polar (Fig. 35G), after 7 days at 25 °C, sediment and ring are formed. After 1 month at 25 °C, sediment is present, the ring and a pellicle do not appear. In Dalmau plate culture on corn meal agar, pseudohyphae are not formed. Asci and ascospores are not observed on V8 and Acetate agar.

###### Physiological and biochemical characteristics

Glucose, lactose, galactose, sucrose, maltose and raffinose fermentation are absent. Glucose, galactose (latent), L-sorbose (latent, weak), sucrose (latent, weak), maltose (latent), cellobiose (latent), trehalose (latent), melibiose (latent, weak), melezitose (latent, weak), inulin (weak), soluble starch (latent), L-rhamnose (latent, weak), D-glucosamine (latent), N-acetyl-D-glucosamine (latent, weak), ethanol, glycerol, D-mannitol, D-glucitol (latent), salicin (latent), DL-lactic acid (latent), succinic acid and citric acid (latent) are assimilated as sole carbon sources. Lactose, raffinose, D-xylose, L-arabinose, D-arabinose, D-ribose, methanol, erythritol, ribitol, galactitol, methyl-α-D-glucoside inositol, hexadecane and D-glucuronic acid are not assimilated. Ammonium sulfate, potassium nitrate (latent, weak), sodium nitrite (latent, weak), L-lysine, ethylamine and cadaverine are assimilated as sole nitrogen sources. The maximum growth temperature is 35 °C. Growth in the vitamin-free medium is positive. Starch-like substances are not produced. Growth on 50 % (w/w) glucose-yeast extract agar is negative.

Physiologically, *P. xizangensis* differs from the closely related species *P. planticola*, *P. manshurica*, *P. garciniae* and *P. membranifaciens* (Fig. 10) in its ability to assimilate cellobiose (Table S2.9).

###### Typus

**China**, Jiangzi County, Shigatze City, Tibet, obtained from soil, Aug. 2019, *Q.-M. Wang* (**holotype** CGMCC 2.7001^T^ preserved in a metabolically inactive state, ex-type cultures are preserved in culture collections under the numbers XZ17A5).

***Saturnispora jianfenglingensis*** Q.-M. Wang, A.-H. Li, M.-M. Liu, H.-H. Zhu & F.-Y. Bai, ***sp. nov*.**

Fungal Names FN 572768. Fig. 35H, I.

###### Etymology

the specific epithet *jianfenglingensis* refers to the geographic origin of the type strain, Jianfengling, Hainan province.

###### Culture characteristics

On YM agar, after 1 month at 25 °C, the streak culture is white, powdery and rough. The margin is entire (Fig. 35H). In YM broth, after 3 days at 25 °C, cells are oval and cylindrical, 3.17–6.22 × 5.75–11.90 µm and single, budding is polar (Fig. 35I), after 7 days at 25 °C, a sediment, pellicle and ring are formed. After 1 month at 25 °C, sediment, pellicle and ring are present. In Dalmau plate culture on corn meal agar, hyphae are formed. Asci and ascospores are not observed on V8 and Acetate agar.

###### Physiological and biochemical characteristics

Glucose, lactose, galactose, sucrose, maltose and raffinose fermentation are absent. Glucose, galactose, L-sorbose, trehalose, melibiose (latent, weak), raffinose, inulin, D-xylose, ethanol, glycerol, ribitol, D-mannitol, D-glucitol, DL-lactic acid and succinic acid are assimilated as sole carbon sources. Sucrose, maltose, cellobiose, lactose, melezitose, soluble starch, L-arabinose, D-arabinose, D-ribose, L-rhamnose, D-glucosamine, N-acetyl-D-glucosamine, methanol, erythritol, galactitol, methyl-α-D-glucoside, salicin, citric acid, inositol, hexadecane and D-glucuronic acid are not assimilated. Ammonium sulfate, potassium nitrate, sodium nitrite, L-lysine, ethylamine and cadaverine are assimilated as sole nitrogen sources. The maximum growth temperature is 35 °C. Growth in the vitamin-free medium is positive. Starch-like substances are not produced. Growth on 50 % (w/w) glucose-yeast extract agar is negative.

Physiologically, *S. jianfenglinensis* differs from the closely related species *S. linzhiensis* and *S. zaruensis* (Fig. 11) in its ability to assimilate melibiose, raffinose, D-xylose and glycerol (Table S2.10).

###### Typus

**China**, Jianfaling, Hainan province, obtained from rotten wood, Jan. 2008, *Q.-M. Wang* (**holotype** CGMCC 2.3783^T^ preserved in a metabolically inactive state).

###### Other culture examined

**China**, Hainan province, obtained from rotten wood, Jan. 2008, isolated by *Q.M. Wang*, culture XZJF10-2.

***Saturnispora linzhiensis*** Q.-M. Wang, A.-H. Li, M.-M. Liu, H.-H. Zhu & F.-Y. Bai, ***sp. nov.*** Fungal Names FN 572769. Fig. 36A, B.

**Fig. 36.**
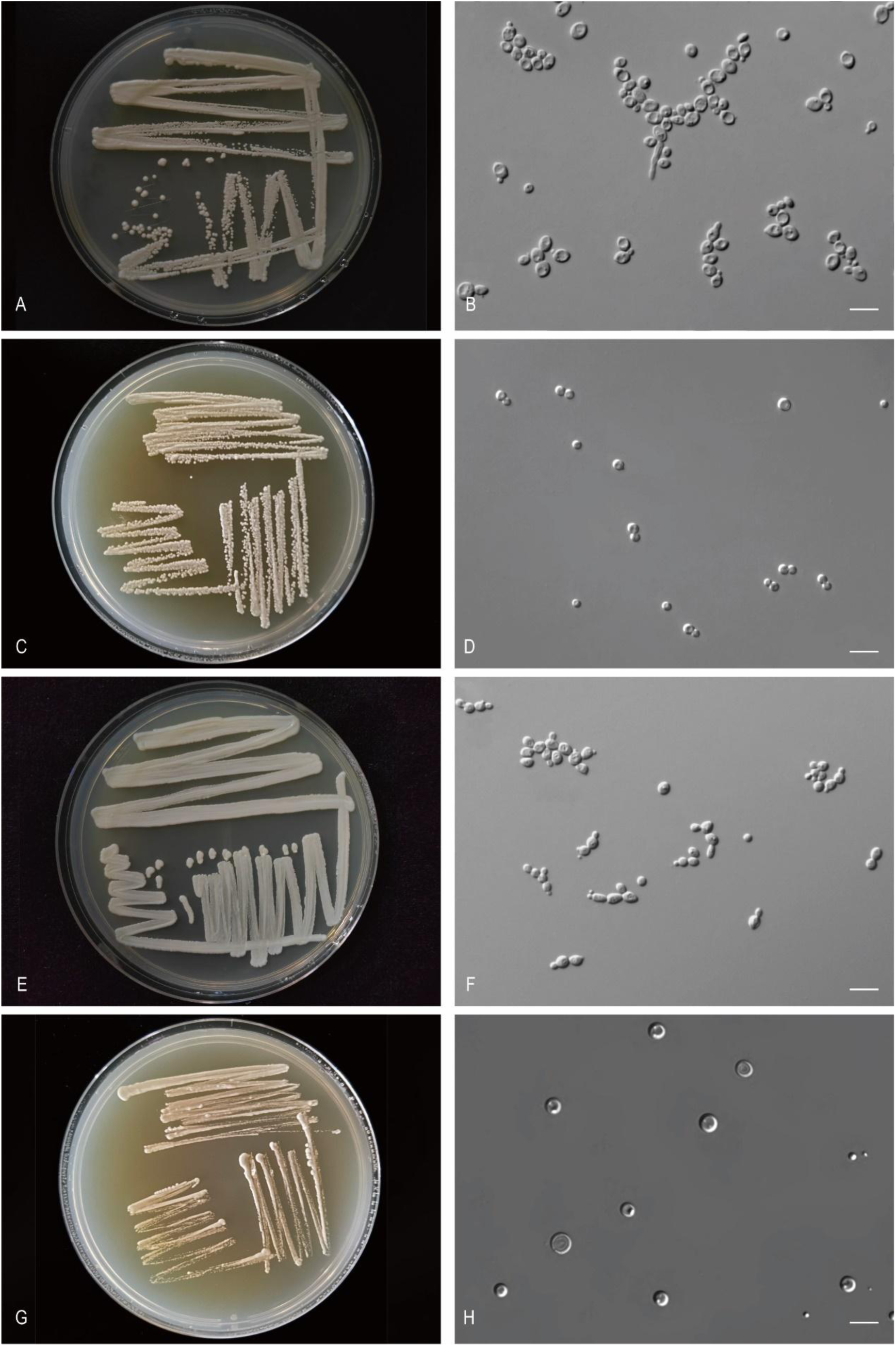
The streak culture grown in YM agar and vegetative cells grown in YM broth for 7 d at 25 °C. A, B. *Sat. linzhiensis* CGMCC 2.7003^T^. C, D. *Xiu. putredisilvae* CGMCC 2.3792^T^. E, F. *Che. Guizhouensis* CGMCC 2.6988^T^. G, H. *Hem. corticola* CGMCC 2.3790^T^. Scale bars: B, D, F, H = 10 μm.

###### Etymology

the specific epithet *linzhiensis* refers to the geographic origin of the type strain, Linzhi, Tibet.

###### Culture characteristics

On YM agar, after 1 month at 25 °C, the streak culture is white, butyrous, smooth and pale glossy. The margin is entire (Fig. 36A). In YM broth, after 3 days at 25 °C, cells are oval and fusiform, 3.42–5.33 × 4.80–6.34 µm and single, budding is polar (Fig. 36B), after 7 days at 25 °C, a sediment is formed. After 1 month at 25 °C, sediment is present, but a ring and pellicle do not appear. In Dalmau plate culture on corn meal agar, pseudohyphae are not formed. Asci and ascospores are not observed on V8 and Acetate agar.

###### Physiological and biochemical characteristics

Glucose fermentation is present. Lactose, galactose, sucrose, maltose and raffinose fermentation are absent. Glucose, galactose (latent, weak), L-sorbose, maltose, trehalose, inulin (weak), ethanol, ribitol (latent), D-mannitol, D-glucitol (latent), DL-lactic acid (latent, weak) and succinic acid are assimilated as sole carbon sources. Sucrose, cellobiose, lactose, melibiose, raffinose, melezitose, soluble starch, D-xylose, L-arabinose, D-arabinose, D-ribose, L-rhamnose, D-glucosamine, N-acetyl-D-glucosamine, methanol, glycerol, erythritol, galactitol, methyl-α-D-glucoside salicin, citric acid, inositol, hexadecane and D-glucuronic acid are not assimilated. Ammonium sulfate, potassium nitrate, sodium nitrite (weak), L-lysine, ethylamine and cadaverine are assimilated as sole nitrogen sources. The maximum growth temperature is 35 °C. Growth in the vitamin-free medium is positive. Starch-like substances are not produced. Growth on 50 % (w/w) glucose-yeast extract agar is negative.

Physiologically, *S. linzhiensis* differs from the closely related species *S. jianfenglinensis* and *S. zaruensis* (Fig. 11) in its ability to assimilate maltose (Table S2.10).

###### Typus

**China**, Jiangzi County, Shigatze City, Tibet, obtained from soil, Aug. 2019, *Q.-M. Wang* (**holotype** CGMCC 2.7003^T^ preserved in a metabolically inactive state, ex-type cultures are preserved in culture collections under the numbers XZ18-24-2).

***Xiuguozyma putredisilvae*** Q.-M. Wang, A.-H. Li, M.-M. Liu, H.-H. Zhu & F.-Y. Bai, ***sp. nov.*** Fungal Names FN 572770. Fig. 36C, D.

###### Etymology

the specific epithet *putredisilvae* refers to the type strain isolated from rotten wood.

###### Culture characteristics

On YM agar, after 1 month at 25 °C, the streak culture is white, butyrous, smooth and pale glossy. The margin is entire (Fig. 36C). In YM broth, after 3 days at 25 °C, cells are globose and oval, 2.15–4.38 × 2.68–4.33 µm and single, budding is polar (Fig. 36D), after 7 days at 25 °C, a sediment is formed. After 1 month at 25 °C, sediment is present, but a ring and pellicle do not appear. In Dalmau plate culture on corn meal agar, pseudohyphae are not formed. Asci and ascospores are not observed on V8 and Acetate agar.

###### Physiological and biochemical characteristics

Glucose, lactose, galactose, sucrose, maltose and raffinose fermentation are absent. Glucose, galactose, L-sorbose, sucrose (weak), maltose (latent), cellobiose (latent), trehalose, melibiose, raffinose, inulin, soluble starch (weak), D-xylose, D-ribose, D-glucosamine, N-acetyl-D-glucosamine, methanol (latent, weak), ethanol, glycerol, ribitol, D-mannitol, D-glucitol, DL-lactic acid and succinic acid are assimilated as sole carbon sources. Lactose, melezitose, L-arabinose, D-arabinose, L-rhamnose, erythritol, galactitol, methyl-α-D-glucoside, salicin, citric acid, inositol, hexadecane and D-glucuronic acid are not assimilated. Ammonium sulfate, potassium nitrate, sodium nitrite (latent), L-lysine, ethylamine and cadaverine are assimilated as sole nitrogen sources. The maximum growth temperature is 40 °C. Growth in the vitamin-free medium is positive. Starch-like substances are not produced. Growth on 50 % (w/w) glucose-yeast extract agar is negative.

Physiologically, *X. putredisilvae* differs from the closely related species *X. silvatica* and *X. insectalens* (Fig. 12) in its ability to assimilate L-sorbose, sucrose, maltose, melibiose, raffinose, inulin, soluble starch, D-xylose, D-ribose, methanol, potassium nitrate, sodium nitrite, and grow in vitamin-free medium (Table S2.11).

###### Typus

**China**, Hainan province, obtained from rotten wood, Jan. 2008, *Q.-M. Wang* (**holotype** CGMCC 2.3792^T^ preserved in a metabolically inactive state).

###### Other culture examined

**China**, Hainan province, obtained from rotten wood, Jan. 2008, isolated by *Q.M. Wang*, culture CGMCC 2.3784 (= G2WZ06-1).

New taxa in the *Debaryomycetaceae* (*Serinales*, *Pichiomycetes*, *Saccharomycotina*) *Chernovozyma guizhouensis* Q.-M. Wang, A.-H. Li, M.-M. Liu, H.-H. Zhu & F.-Y. Bai, *sp. nov.* Fungal Names FN 572771. Fig. 36E, F.

###### Etymology

the specific epithet *guizhouensis* refers to the geographic origin of the type strain, Guizhou province.

###### Culture characteristics

On YM agar, after 1 month at 25 °C, the streak culture is whitish-cream, butyrous, smooth and pale glossy. The margin is entire (Fig. 36E). In YM broth, after 3 days at 25 °C, cells are oval and fusiform, 2.61–3.89 × 3.10–5.59 µm and single, budding is polar (Fig. 36F), after 7 days at 25 °C, a sediment and ring are formed. After 1 month at 25 °C, sediment and ring are present, but pellicle does not appear. In Dalmau plate culture on corn meal agar, pseudohyphae are not formed. Asci and ascospores are not observed on V8 and Acetate agar.

###### Physiological and biochemical characteristics

Glucose (weak) fermentation is present. Lactose, galactose, sucrose, maltose and raffinose fermentation are absent. Glucose, galactose, L-sorbose, trehalose, melibiose, raffinose, melezitose (weak), inulin, D-arabinose (latent, weak), D-ribose (latent), D-glucosamine, N-acetyl-D-glucosamine, ethanol, glycerol, ribitol, D-mannitol, D-glucitol, DL-lactic acid, succinic acid, citric acid and D-glucuronic acid (latent, weak) are assimilated as sole carbon sources. Sucrose, maltose, cellobiose, lactose, soluble starch, D-xylose, L-arabinose, L-rhamnose, methanol, erythritol, galactitol, methyl-α-D-glucoside, salicin, inositol and hexadecane are not assimilated. Ammonium sulfate, potassium nitrate, sodium nitrite, L-lysine, ethylamine and cadaverine are assimilated as sole nitrogen sources. The maximum growth temperature is 32 °C. Growth in the vitamin-free medium is positive (weak). Starch-like substances are not produced. Growth on 50 % (w/w) glucose-yeast extract agar is positive.

Physiologically, *C. guizhouensis* differs from the closely related species *C. aurita*, *C. sophiae-reginae* and *C. palmyrensis* (Fig. 13) in its ability to assimilate melibiose, raffinose, inulin, D-arabinose, potassium nitrate, sodium nitrite, and grow with 50% glucose and vitamin-free medium (Table S2.12).

###### Typus

**China**, Maotai Town, Hunan province, obtained from soil, Aug. 2019, *Q.-M. Wang* (**holotype** CGMCC 2.6988^T^ preserved in a metabolically inactive state, ex-type cultures are preserved in culture collections under the numbers gmt3-3-4).

###### Other culture examined

**China**, Maotai Town, Hunan province, obtained from soil, Aug. 2019, isolated by *Q.M. Wang*, culture gmt3-3-5.

***Hemisphaericaspora corticola*** Q.-M. Wang, A.-H. Li, M.-M. Liu, H.-H. Zhu & F.-Y. Bai, ***sp. nov.*** Fungal Names FN 572772. Fig. 36G, H.

###### Etymology

the specific epithet *corticola* refers to the type strain isolated from bark.

###### Culture characteristics

On YM agar, after 1 month at 25 °C, the streak culture is white, butyrous and smooth. The margin is entire (Fig. 36G). In YM broth, after 3 days at 25 °C, cells are globose and oval, 3.70–6.74 × 3.72–5.90 µm and single, budding is polar (Fig. 36H), after 7 days at 25 °C, a sediment is formed. After 1 month at 25 °C, sediment is present, the ring and a pellicle do not appear. In Dalmau plate culture on corn meal agar, pseudohyphae are not formed. Asci and ascospores are not observed on V8 and Acetate agar.

###### Physiological and biochemical characteristics

Glucose (weak) and lactose (weak) fermentation are present. Galactose, sucrose, maltose and raffinose fermentation are absent. Glucose, L-sorbose, sucrose, maltose, cellobiose (latent), trehalose, lactose (latent, weak), melibiose (weak), raffinose (weak), melezitose (weak), inulin, D-xylose, D-arabinose (latent, weak), D-ribose (latent, weak), L-rhamnose (weak), D-glucosamine, N-acetyl-D-glucosamine, ethanol (weak), ribitol (latent), galactitol (latent, weak), D-mannitol, D-glucitol, methyl-α-D-glucoside (latent), salicin and succinic acid (latent) are assimilated as sole carbon sources. Galactose, soluble starch, L-arabinose, methanol, glycerol, erythritol, DL-lactic acid, citric acid, inositol, hexadecane and D-glucuronic acid are not assimilated. Ammonium sulfate, potassium nitrate (latent), sodium nitrite (latent, weak), L-lysine, ethylamine and cadaverine (weak) are assimilated as sole nitrogen sources. The maximum growth temperature is 37 °C. Growth in the vitamin-free medium is positive. Starch-like substances are not produced. Growth on 50 % (w/w) glucose-yeast extract agar is positive.

Physiologically, *H. corticola* differs from the closely related species *H. elongata*, *H. mengyangensis* and *H. subhashii* (Fig. 14) in its ability to assimilate L-sorbose, lactose, melibiose, raffinose, L-rhamnose, galactitol, potassium nitrate and sodium nitrite (Table S2.13).

###### Typus

**China**, Hainan province, obtained from rotten wood, *Q.-M. Wang* (**holotype** CGMCC 2.3790^T^ preserved in a metabolically inactive state).

***Insectozyma pseudocorydali*** Q.-M. Wang, A.-H. Li, M.-M. Liu, H.-H. Zhu & F.-Y. Bai, ***sp. nov.*** Fungal Names FN 572773. Fig. 37A, B.

**Fig. 37.**
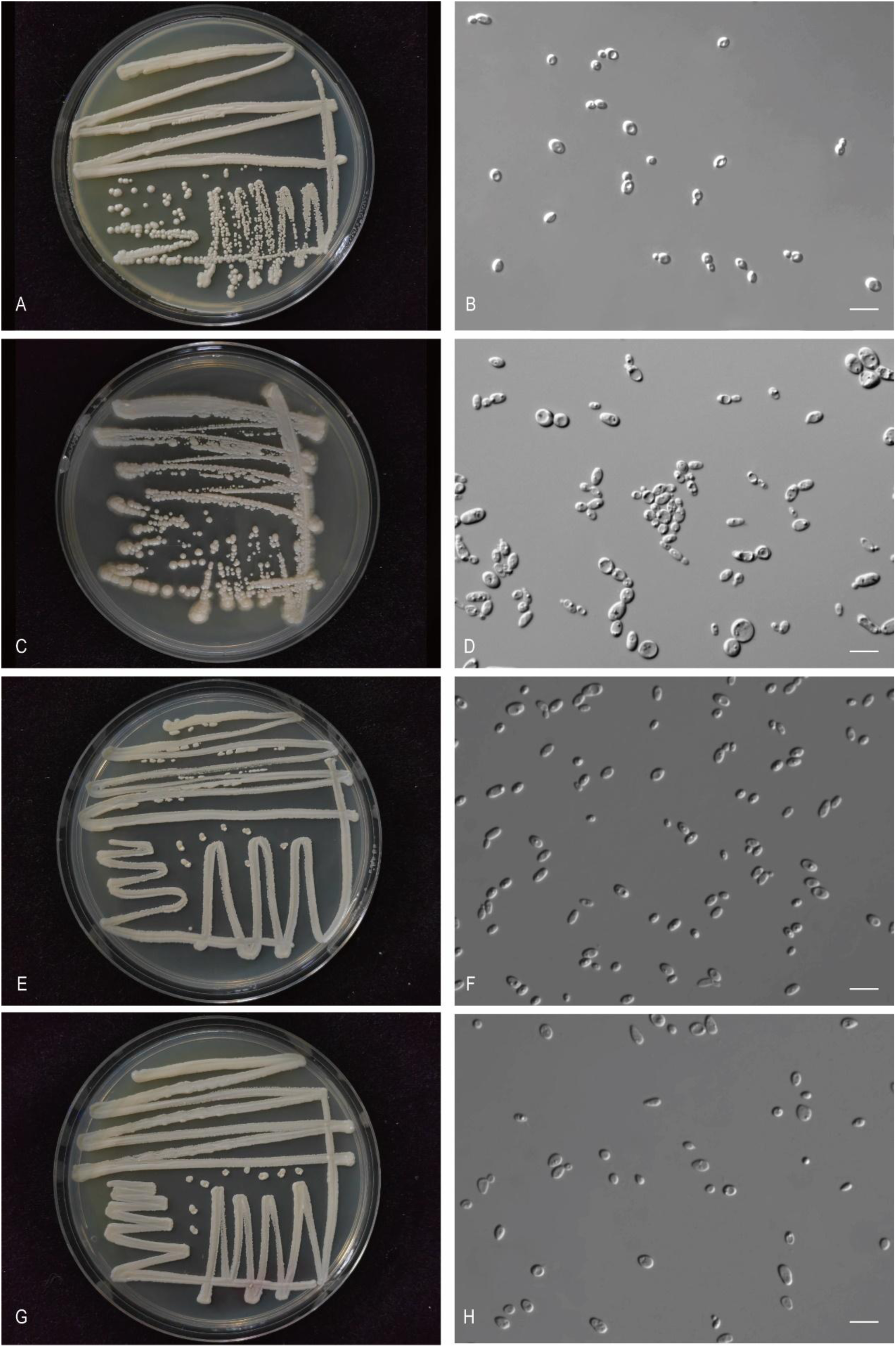
The streak culture grown in YM agar and vegetative cells grown in YM broth for 7 d at 25 °C. A, B. *Ins. pseudocorydali* CGMCC 2.3733^T^. C, D. *Suh. caprifoliacearum* CGMCC 2.7037^T^. E, F. *Suh. Ilexicola* CGMCC 2.6997^T^. G, H. *Suh. quercus* CGMCC 2.6996^T^. Scale bars: B, D, F, H = 10 μm.

###### Etymology

the specific epithet *pseudocorydali* refers to a similar colony morphology to that of *Insectozyma corydali*.

###### Culture characteristics

On YM agar, after 1 month at 25 °C, the streak culture is white, butyrous, and smooth. The margin is entire (Fig. 37A). In YM broth, after 3 days at 25 °C, cells are subglobose, oval and ellipsoidal, 2.52–4.23 × 3.70–5.96 µm and single, budding is polar (Fig. 37B), after 7 days at 25 °C, a sediment is formed. After 1 month at 25 °C, sediment and ring are present, but pellicle does not appear. In Dalmau plate culture on corn meal agar, pseudohyphae are not formed. Asci and ascospores are not observed on V8 and Acetate agar.

###### Physiological and biochemical characteristics

Glucose and galactose (variable) fermentation are present. Lactose, sucrose, maltose and raffinose fermentation are absent. Glucose, galactose, L-sorbose, sucrose, maltose, cellobiose (variable), trehalose (variable), melibiose (variable), raffinose (variable), melezitose, inulin, soluble starch(variable), D-xylose, L-arabinose, D-ribose (variable), D-glucosamine, N-acetyl-D-glucosamine, ethanol, glycerol (variable), ribitol, galactitol (variable), D-mannitol (variable), D-glucitol, methyl-α-D-glucoside, salicin, DL-lactic acid (variable), succinic acid and citric acid are assimilated as sole carbon sources. Lactose, D-arabinose, L-rhamnose, methanol, erythritol, inositol, hexadecane and D-glucuronic acid are not assimilated. Ammonium sulfate, potassium nitrate, sodium nitrite (variable), L-lysine, ethylamine and cadaverine are assimilated as sole nitrogen sources. The maximum growth temperature is 37 °C. Growth in the vitamin-free medium is positive. Starch-like substances are not produced. Growth on 50 % (w/w) glucose-yeast extract agar is positive (variable).

Physiologically, *I. pseudocorydali* differs from the closely related species *I. corydali* (Fig. 15) in its inability to assimilate D-glucuronic acid and the ability to assimilate inulin, potassium nitrate, and grow in vitamin-free medium (Table S2.14).

###### Typus

**China**, Beijing, obtained from the gut of *Anomala aulax*, Oct. 2007, *Q.-M. Wang* (**holotype** CGMCC 2.3733^T^ preserved in a metabolically inactive state, ex-type cultures are preserved in culture collections under the numbers LJG1.1).

###### Other culture examined

**China**, Shaanxi province, obtained from soil, isolated by *Q.M. Wang*, culture CGMCC 2.3277. **China**, Hainan province, obtained from bark, isolated by *Q.M. Wang*, culture CGMCC 2.3227. **China**, Hainan province, obtained from flower, isolated by *Q.M. Wang*, culture CGMCC 2.3494.

***Suhomyces caprifoliacearum*** Q.-M. Wang, A.-H. Li, M.-M. Liu, H.-H. Zhu & F.-Y. Bai, ***sp. nov.*** Fungal Names FN 572774. Fig. 37C, D.

###### Etymology

the specific epithet *caprifoliacearum* refers to *Caprifoliaceae*, the plant genus from which the type strain was isolated.

###### Culture characteristics

On YM agar, after 1 month at 25 °C, the streak culture is white, butyrous, and pale glossy. The margin is fimbriate (Fig. 37C). In YM broth, after 3 days at 25 °C, cells are globose, fusiform and cylindrical, 2.52–3.97 × 2.51–4.81 µm and single, budding is polar (Fig. 37D), after 7 days at 25 °C, a sediment is formed. After 1 month at 25 °C, sediment is present, but a ring and pellicle do not appear. In Dalmau plate culture on corn meal agar, hyphae are formed. Asci and ascospores are not observed on V8 and Acetate agar.

###### Physiological and biochemical characteristics

Glucose fermentation is present. Lactose, galactose, sucrose, maltose and raffinose fermentation are absent. Glucose, galactose (variable), L-sorbose (latent), sucrose, maltose, cellobiose, trehalose, lactose (variable), raffinose (variable), melezitose, inulin, soluble starch (variable), D-xylose (latent, weak), D-arabinose (variable), D-ribose (variable), D-glucosamine (latent, weak), N-acetyl-D-glucosamine, ethanol (latent), glycerol (variable), D-mannitol, D-glucitol, methyl-α-D-glucoside (variable), salicin (latent, weak), succinic acid and citric acid (latent) are assimilated as sole carbon sources. Melibiose, L-arabinose, L-rhamnose, methanol, erythritol, ribitol, galactitol, DL-lactic acid, inositol, hexadecane and D-glucuronic acid are not assimilated. Ammonium sulfate, potassium nitrate, sodium nitrite (variable), L-lysine, ethylamine and cadaverine are assimilated as sole nitrogen sources. The maximum growth temperature is 20 °C. Growth in the vitamin-free medium is positive (latent). Starch-like substances are not produced. Growth on 50 % (w/w) glucose-yeast extract agar is positive (variable).

Physiologically, *S. caprifoliacearum* differs from the closely related species *S. ilexicola* and *S. quercus* (Fig. 16) in its inability to assimilate ribitol (Table S2.15).

###### Typus

**China**, Sejila Mountain, Nyingchi, Tibet, obtained from the leaf of *Caprifoliaceae*, Aug. 2019, *Q.- M. Wang* (**holotype** CGMCC 2.7037^T^ preserved in a metabolically inactive state, ex-type cultures are preserved in culture collections under the numbers XZY145-10).

###### Other culture examined

**China**, Sejila Mountain, Nyingchi, Tibet, obtained from plant, Aug. 2019, isolated by *Q.M. Wang*, culture XZY144-8; **China**, Sejila Mountain, Nyingchi, Tibet, obtained from leaf of *Caprifoliaceae*, Aug. 2019, isolated by *Q.M. Wang*, culture XZY145M2.

***Suhomyces ilexicola*** Q.-M. Wang, A.-H. Li, M.-M. Liu, H.-H. Zhu & F.-Y. Bai, ***sp. nov.*** Fungal Names FN 572775. Fig. 37E, F.

###### Etymology

the specific epithet *ilexicola* refers to *Ilex*, the plant genus from which the type strain was isolated.

###### Culture characteristics

On YM agar, after 1 month at 25 °C, the streak culture is white, butyrous, smooth and pale glossy. The margin is entire (Fig. 37E). In YM broth, after 3 days at 25 °C, cells are ellipsoidal and fusiform, 2.65–3.56 × 3.59–6.15 µm and single, budding is polar (Fig. 37F), after 7 days at 25 °C, a sediment is formed. After 1 month at 25 °C, sediment is present, but a ring and pellicle do not appear. In Dalmau plate culture on corn meal agar, pseudohyphae are not formed. Asci and ascospores are not observed on V8 and Acetate agar.

###### Physiological and biochemical characteristics

Glucose fermentation is present. Lactose, galactose, sucrose, maltose and raffinose fermentation are absent. Glucose, galactose, L-sorbose, sucrose, maltose, cellobiose, trehalose, melibiose (variable), raffinose (variable), melezitose, inulin, soluble starch, D-xylose, D-ribose (latent), D-glucosamine (latent), N-acetyl-D-glucosamine (latent, weak), ethanol, glycerol (latent), erythritol, ribitol, D-mannitol, D-glucitol, methyl-α-D-glucoside, salicin, succinic acid and citric acid are assimilated as sole carbon sources. Lactose, L-arabinose, D-arabinose, L-rhamnose, methanol, galactitol, DL-lactic acid, inositol, hexadecane and D-glucuronic acid are not assimilated. Ammonium sulfate, potassium nitrate, sodium nitrite (latent, weak), L-lysine, ethylamine and cadaverine are assimilated as sole nitrogen sources. The maximum growth temperature is 25 °C. Growth in the vitamin-free medium is positive. Starch-like substances are not produced. Growth on 50 % (w/w) glucose-yeast extract agar is negative.

Physiologically, *S. ilexicola* differs from the closely related species *S. caprifoliacearum* and *S. quercus* (Fig. 15) in its ability to assimilate erythritol (Table S2.15).

###### Typus

**China**, Katinggou, Nyingchi City, Tibet, obtained from plant, Aug. 2019, *Q.-M. Wang* (**holotype** CGMCC 2.6997^T^ preserved in a metabolically inactive state, ex-type cultures are preserved in culture collections under the numbers XZY189-1).

###### Other culture examined

**China**, Sejila Mountain, Nyingchi, Tibet, obtained from plant, Aug. 2019, isolated by *Q.M. Wang*, culture XZY165-1.

***Suhomyces quercus*** Q.-M. Wang, A.-H. Li, M.-M. Liu, H.-H. Zhu & F.-Y. Bai, ***sp. nov.*** Fungal Names FN 572776. Fig. 37G, H.

###### Etymology

the specific epithet *quercus* refers to *Quercus*, the plant genus from which the type strain was isolated.

###### Culture characteristics

On YM agar, after 1 month at 25 °C, the streak culture is white, butyrous, and smooth. The margin is entire (Fig. 37G). In YM broth, after 3 days at 25 °C, cells are ellipsoidal and fusiform, 1.94–4.98 × 3.20–7.87 µm and single, budding is polar (Fig. 37H), after 7 days at 25 °C, a sediment is formed. After 1 month at 25 °C, sediment is present, but a ring and pellicle do not appear. In Dalmau plate culture on corn meal agar, pseudohyphae are not formed. Asci and ascospores are not observed on V8 and Acetate agar.

###### Physiological and biochemical characteristics

Glucose, lactose (variable) and galactose fermentation are present. Sucrose, maltose and raffinose fermentation are absent. Glucose, galactose (latent), L-sorbose (latent), sucrose, maltose, cellobiose, trehalose, lactose (variable), raffinose (variable), melezitose (latent), inulin (latent), soluble starch (variable), D-xylose (variable), D-glucosamine (latent), N-acetyl-D-glucosamine (latent), ethanol (latent), glycerol (latent), ribitol (latent), D-mannitol, D-glucitol (latent), methyl-α-D-glucoside, salicin, succinic acid and citric acid (variable) are assimilated as sole carbon sources. Melibiose, L-arabinose, D-arabinose, D-ribose, L-rhamnose, methanol, erythritol, galactitol, DL-lactic acid, inositol, hexadecane and D-glucuronic acid are not assimilated. Ammonium sulfate, potassium nitrate, sodium nitrite, L-lysine, ethylamine (latent) and cadaverine are assimilated as sole nitrogen sources. The maximum growth temperature is 30 °C. Growth in the vitamin-free medium is positive. Starch-like substances are not produced. Growth on 50 % (w/w) glucose-yeast extract agar is positive (variable).

Physiologically, *S. quercus* differs from the closely related species *S. caprifoliacearum* and *S. ilexicola* (Fig. 16) in its ability to ferment galactose (Table S2.15).

###### Typus

**China**, Baiba County, Nyingchi, Tibet, obtained from bark of *Quercus semecarpifolia*, Aug. 2019, *Q.-M. Wang* (**holotype** CGMCC 2.6996^T^ preserved in a metabolically inactive state, ex-type cultures are preserved in culture collections under the numbers XZY98-4).

###### Other culture examined

**China**, New basom lake, Linzhi, Tibet, obtained from bark of *Pinus*, Aug. 2019, isolated by *Q.M. Wang*, culture XZY480-1; **China**, New basom lake, Linzhi, Tibet, obtained from rotten wood, Aug. 2019, isolated by *Q.M. Wang*, culture XZY457-1; **China**, New basom lake, Linzhi, Tibet, obtained from rotten wood, Aug. 2019, isolated by *Q.M. Wang*, culture XZY477-3.

***Suhomyces xizangensis*** Q.-M. Wang, A.-H. Li, M.-M. Liu, H.-H. Zhu & F.-Y. Bai, ***sp. nov.*** Fungal Names FN 572777. Fig. 38A, B.

**Fig. 38.**
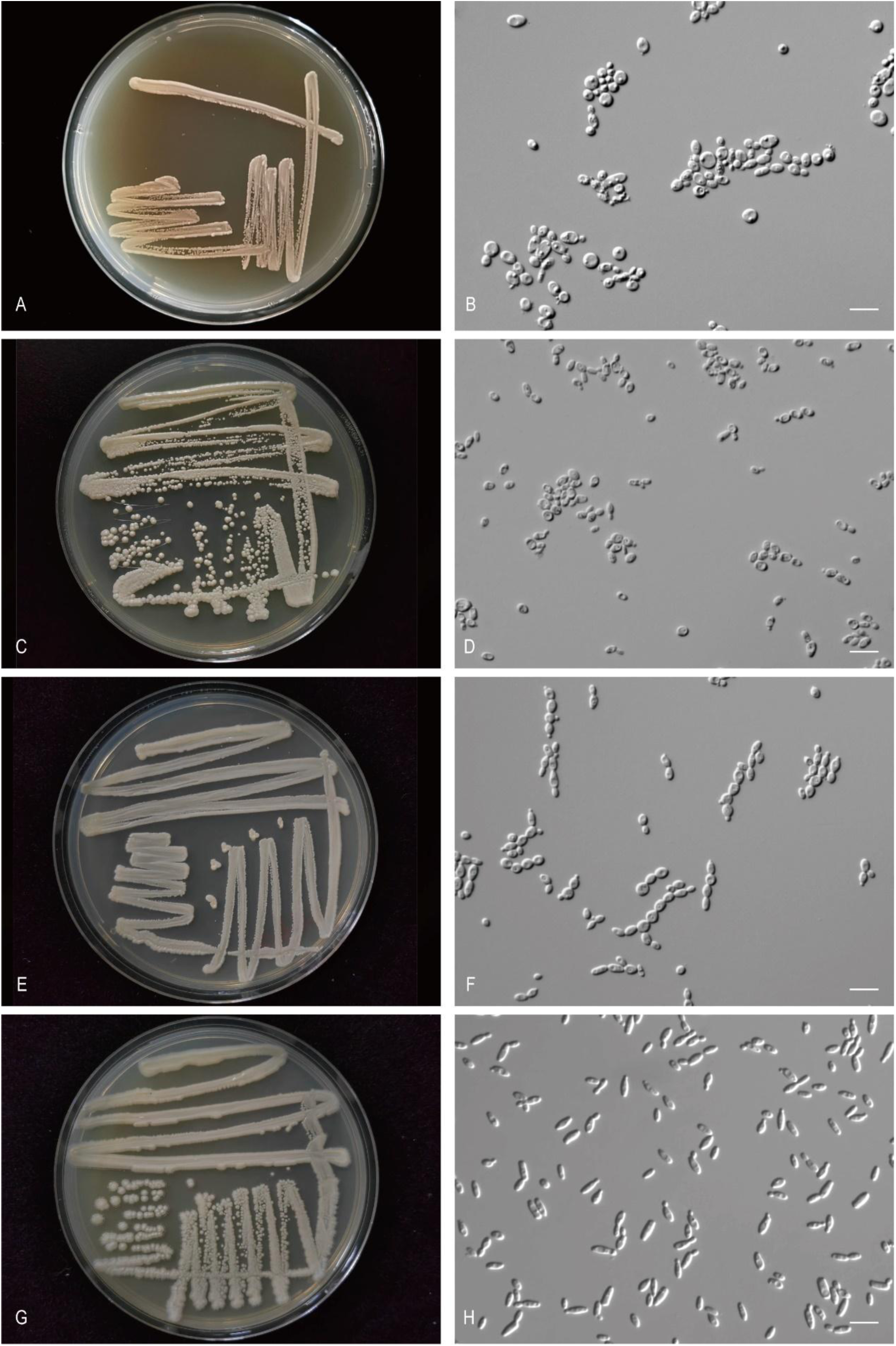
The streak culture grown in YM agar and vegetative cells grown in YM broth for 7 d at 25 °C. A, B. *Suh. xizangensis* CGMCC 2.8158^T^. C, D. *Yam. betulicola* CGMCC 2.6994^T^. E, F. *Yam. chuxiongensis* CGMCC 2.6995^T^. G, H. *Yam. follicola* CGMCC 2.3482^T^. Scale bars: B, D, F, H = 10 μm.

###### Etymology

the specific epithet *xizangensis* refers to the geographic origin of the type strain, Xizang.

###### Culture characteristics

On YM agar, after 1 month at 25 °C, the streak culture is white, butyrous, and pale glossy. The margin is fimbriate (Fig. 38A). In YM broth, after 3 days at 25 °C, cells are cylindrical, ellipsoidal and spherical, 3.43–6.62 × 3.14–7.62 µm and single, budding is polar (Fig. 38B), after 7 days at 25 °C, a sediment is formed. After 1 month at 25 °C, sediment is present, but a ring and pellicle do not appear. In Dalmau plate culture on corn meal agar, pseudohyphae are not formed. Asci and ascospores are not observed on V8 and Acetate agar.

###### Physiological and biochemical characteristics

Glucose fermentation is present. Lactose, galactose, sucrose, maltose and raffinose fermentation are absent. Glucose, galactose, L-sorbose, sucrose, maltose, cellobiose (latent), trehalose, melezitose, inulin, D-xylose (latent), D-glucosamine (latent), N-acetyl-D-glucosamine, ethanol, glycerol (latent), ribitol, D-mannitol, D-glucitol, methyl-α-D-glucoside (latent), salicin (latent) and succinic acid are assimilated as sole carbon sources. Lactose, melibiose, raffinose, soluble starch, L-arabinose, D-arabinose, D-ribose, L-rhamnose, methanol, erythritol, galactitol, DL-lactic acid, citric acid, inositol, hexadecane and D-glucuronic acid are not assimilated. Ammonium sulfate, potassium nitrate, sodium nitrite, L-lysine, ethylamine and cadaverine are assimilated as sole nitrogen sources. The maximum growth temperature is 25 °C. Growth in the vitamin-free medium is positive. Starch-like substances are not produced. Growth on 50 % (w/w) glucose-yeast extract agar is negative.

Physiologically, *S. xizangensis* differs from the closely related species *S. chickasaworum*, *S. emberorum*, *S. wounanorum*, *S. yuchorum*, *S. terraborum*, *S. guaymorum*, *S. coccinellae*, *S. bokatorum*, *S. kunorum*, *S. rilaensis*, *S. bolitotheri*, *S. choctaworum*, *S. canberraensis*, *S. drosophilae*, *S. panamericanus*, *S. atakaporum*, *S. tanzawaensis*, *S. ambrosiae*, *S. taliae*, *S. anneliseae*, *S. maxi*, *S. bribrorum*, *S. vadensis*, *S. faveliae*, *S. prunicola*, *S. pyralidae*, *S. kilbournensis*, *S. xylopsoci*, *S. quercus*, *S. caprifoliacearum* and *S. ilexicola* (Fig. 16) in its ability to assimilate carbon sources and nitrogen sources (Table S2.15).

###### Typus

**China**, Zhangmu, Linzhi City, Tibet, obtained from rotten wood, Aug. 2019, *Q.-M. Wang* (**holotype** CGMCC 2.8158^T^ preserved in a metabolically inactive state, ex-type cultures are preserved in culture collections under the numbers XZFM12-1).

###### Other culture examined

**China**, Gyantse County, Shigatse City, Tibet, obtained from soil, Aug. 2019, isolated by *Q.M. Wang*, culture XZ15-1, **China**, Zhangmu, Linzhi, Tibet, obtained from rotten wood, Aug. 2019, isolated by *Q.M. Wang*, culture XZFM10F2.

***Yamadazyma betulicola*** Q.-M. Wang, A.-H. Li, M.-M. Liu, H.-H. Zhu & F.-Y. Bai, ***sp. nov.*** Fungal Names FN 572778. Fig. 38C, D.

###### Etymology

the specific epithet *betulicola* refers to the substrate origin of the type strain, *Betula*.

###### Culture characteristics

On YM agar, after 1 month at 25 °C, the streak culture is amber, butyrous, smooth and glossy. The margin is entire (Fig. 38C). In YM broth, after 3 days at 25 °C, cells are ellipsoidal and fusiform, 2.12–4.38 × 3.98–6.73 µm and single, budding is polar (Fig. 38D), after 7 days at 25 °C, a sediment and ring are formed. After 1 month at 25 °C, sediment and ring are present, but pellicle does not appear. In Dalmau plate culture on corn meal agar, pseudohyphae are not formed. Asci and ascospores are not observed on V8 and Acetate agar.

###### Physiological and biochemical characteristics

Glucose and galactose fermentation are present. Lactose, sucrose, maltose and raffinose fermentation are absent. Glucose, galactose, L-sorbose, sucrose, maltose, cellobiose, trehalose, melezitose, inulin, soluble starch (weak), D-xylose, L-arabinose, D-arabinose (variable), D-ribose, L-rhamnose, D-glucosamine (variable), N-acetyl-D-glucosamine, ethanol (latent), glycerol, erythritol, ribitol, D-mannitol, D-glucitol, methyl-α-D-glucoside, salicin, succinic acid and citric acid are assimilated as sole carbon sources. Lactose, melibiose, raffinose, methanol, galactitol, DL-lactic acid, inositol, hexadecane and D-glucuronic acid are not assimilated. Ammonium sulfate, potassium nitrate, sodium nitrite (variable), L-lysine, ethylamine and cadaverine are assimilated as sole nitrogen sources. The maximum growth temperature is 42 °C. Growth in the vitamin-free medium is positive. Starch-like substances are not produced. Growth on 50 % (w/w) glucose-yeast extract agar is positive.

Physiologically, *Y. betulicola* differs from the closely related species *Y. planticola* (Fig. 17) in its inability to assimilate raffinose and DL-lactic acid, and the ability to ferment galactose and grow with 50% glucose (Table S2.16).

###### Typus

**China**, Basom lake, Linzhi, Tibet, obtained from leaf of *Betula platyphylla*, Aug. 2019, *Q.-M. Wang* (**holotype** CGMCC 2.6994^T^ preserved in a metabolically inactive state, ex-type cultures are preserved in culture collections under the numbers XZY130-3).

###### Other culture examined

**China**, Katinggou, Nyingchi City, Tibet, obtained from *Pteridophyta*, Aug. 2019, isolated by *Q.M. Wang*, culture XZY198-1; **China**, Katinggou, Nyingchi City, Tibet, obtained from leaf of *Lirianthe delavayi*, Aug. 2019, isolated by *Q.M. Wang*, culture XZY176-2; **China**, Katinggou, Nyingchi City, Tibet, obtained from *Pteridophyta*, Aug. 2019, isolated by *Q.M. Wang*, culture XZY198-9.

***Yamadazyma chuxiongensis*** Q.-M. Wang, A.-H. Li, M.-M. Liu, H.-H. Zhu & F.-Y. Bai, ***sp. nov.*** Fungal Names FN 572779. Fig. 38E, F.

###### Etymology

the specific epithet *chuxiongensis* refers to the geographic origin of the type strain, Chuxiong, Yunnan province.

###### Culture characteristics

On YM agar, after 1 month at 25 °C, the streak culture is white, butyrous, smooth and pale glossy. The margin is entire (Fig. 38E). In YM broth, after 3 days at 25 °C, cells are ellipsoidal, cylindrical and fusiform, 1.80–3.34 × 2.79–6.42 µm and single, budding is polar (Fig. 38F), after 7 days at 25 °C, a sediment is formed. After 1 month at 25 °C, sediment is present, a ring and pellicle do not appear. In Dalmau plate culture on corn meal agar, pseudohyphae are not formed. Asci and ascospores are not observed on V8 and Acetate agar.

###### Physiological and biochemical characteristics

Glucose and galactose fermentation are present. Lactose, sucrose, maltose and raffinose fermentation are absent. Glucose, galactose, L-sorbose (latent), sucrose (latent), maltose (latent), cellobiose (latent), trehalose, lactose (latent), melibiose (weak), raffinose (weak), melezitose (latent), inulin, soluble starch (latent), D-xylose (latent), L-arabinose (latent), D-arabinose (latent), D-ribose (variable), L-rhamnose, D-glucosamine (latent), N-acetyl-D-glucosamine, ethanol (latent), glycerol, erythritol, ribitol, D-mannitol, D-glucitol, methyl-α-D-glucoside (latent), salicin (latent), succinic acid and citric acid are assimilated as sole carbon sources. Methanol, galactitol, DL-lactic acid, inositol, hexadecane and D-glucuronic acid are not assimilated. Ammonium sulfate, potassium nitrate (latent), sodium nitrite (latent, weak), L-lysine (latent), ethylamine and cadaverine are assimilated as sole nitrogen sources. The maximum growth temperature is 42 °C. Growth in the vitamin-free medium is positive. Starch-like substances are not produced. Growth on 50 % (w/w) glucose-yeast extract agar is negative.

Physiologically, *Y. chuxiongensis* differs from the closely related species *Y. insectora* and *Y. riverae* (Fig. 17) in its inability to assimilate hexadecane and D-glucuronic acid and the ability to assimilate melibiose, inulin, soluble starch, potassium nitrate, sodium nitrite and grow in vitamin-free medium (Table S2.16).

###### Typus

**China**, Lufeng County, Chuxiong, Yunnan Province, obtained from rotten wood, Oct. 2019, *Q.-M. Wang* (**holotype** CGMCC 2.6995^T^ preserved in a metabolically inactive state, ex-type cultures are preserved in culture collections under the numbers LF3-2).

###### Other culture examined

**China**, Lufeng County, Chuxiong, Yunnan Province, obtained from rotten wood, Oct. 2019, isolated by *Q.M. Wang*, culture LF2-1.

***Yamadazyma follicola*** Q.-M. Wang, A.-H. Li, M.-M. Liu, H.-H. Zhu & F.-Y. Bai, ***sp. nov.*** Fungal Names FN 572780. Fig. 38G, H.

###### Etymology

the specific epithet *follicola* refers to the type strain isolated from leaf of an unidentified plant.

###### Culture characteristics

On YM agar, after 1 month at 25 °C, the streak culture is white, butyrous, and pale glossy. The margin is fimbriate (Fig. 38G). In YM broth, after 3 days at 25 °C, cells are cylindrical, fusiform and ellipsoidal, 2.07–3.23 × 3.47–6.07 µm and single, budding is polar (Fig. 38H), after 7 days at 25 °C, a sediment is formed. After 1 month at 25 °C, sediment is present, a ring and pellicle do not appear. In Dalmau plate culture on corn meal agar, pseudohyphae are not formed. Asci and ascospores are not observed on V8 and Acetate agar.

###### Physiological and biochemical characteristics

Glucose and galactose (weak) fermentation are present. Lactose, sucrose, maltose and raffinose fermentation are absent. Glucose, galactose, L-sorbose, sucrose, maltose, cellobiose, trehalose, lactose, melibiose, raffinose, melezitose, inulin, soluble starch, D-xylose, L-arabinose, D-arabinose, D-ribose, L-rhamnose, D-glucosamine, N-acetyl-D-glucosamine, ethanol, erythritol, ribitol, galactitol, D-glucitol, salicin, DL-lactic acid, succinic acid and citric acid are assimilated as sole carbon sources. Methanol, glycerol, D-mannitol, methyl-α-D-glucoside inositol, hexadecane and D-glucuronic acid are not assimilated. Ammonium sulfate, potassium nitrate, L-lysine, ethylamine (latent) and cadaverine are assimilated as sole nitrogen sources. Sodium nitrite is not assimilated. The maximum growth temperature is 35 °C. Growth in the vitamin-free medium is positive. Starch-like substances are not produced. Growth on 50 % (w/w) glucose-yeast extract agar is negative.

Physiologically, *Y. follicola* differs from the closely related species *Y. tenuis* and *Y. laniorum* (Fig. 17) in its inability to assimilate glycerol and the ability to assimilate melibiose, raffinose and potassium nitrate (Table S2.16).

###### Typus

**China**, Hainan province, obtained from leaf, *Q.-M. Wang* (**holotype** CGMCC 2.3482^T^ preserved in a metabolically inactive state).

***Yamadazyma planticola*** Q.-M. Wang, A.-H. Li, M.-M. Liu, H.-H. Zhu & F.-Y. Bai, ***sp. nov.*** Fungal Names FN 572781. Fig. 39A, B.

**Fig. 39.**
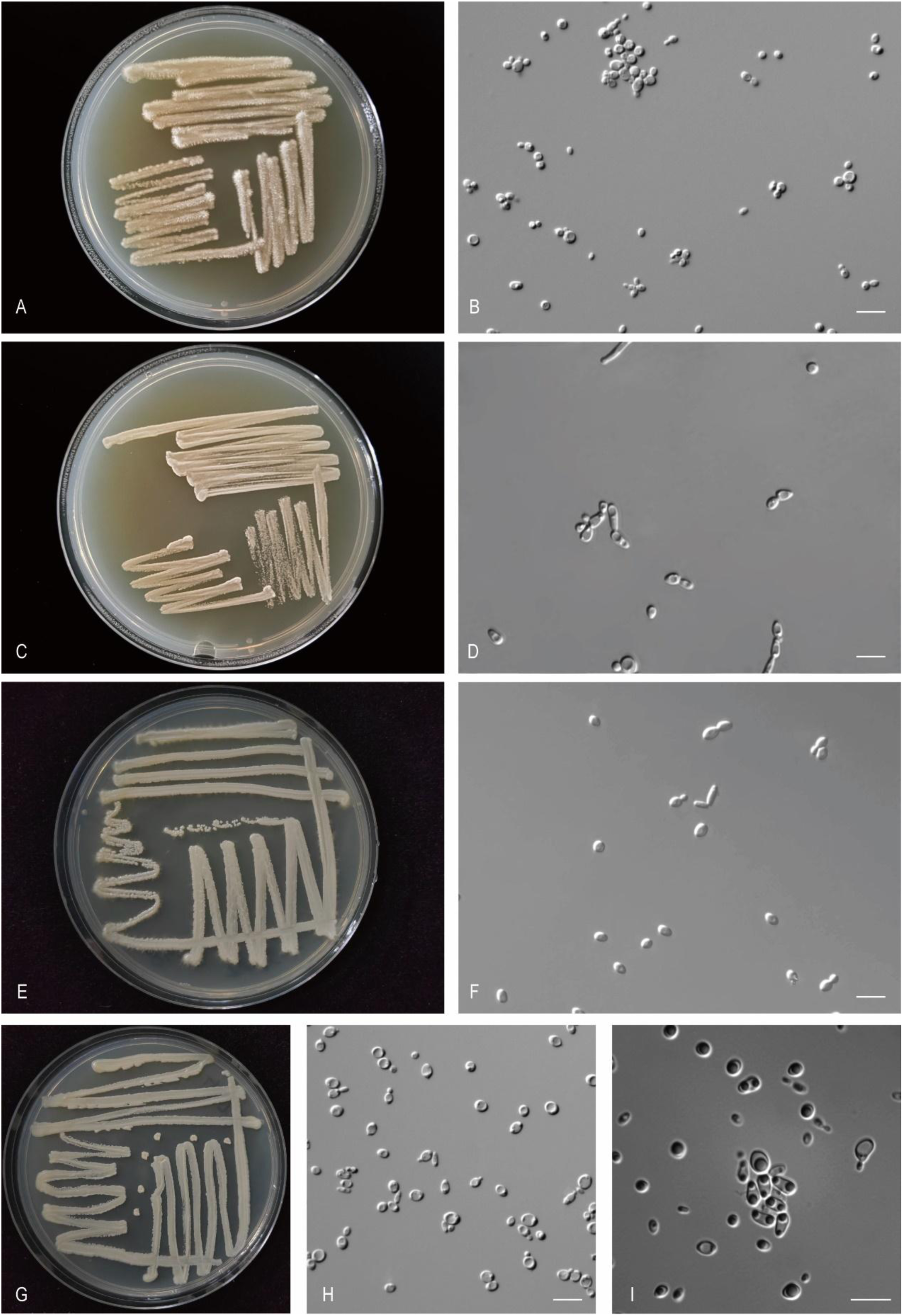
The streak culture grown in YM agar and vegetative cells grown in YM broth for 7 d at 25 °C. A, B. *Yam. planticola* CGMCC 2.8167^T^. C, D. *Yam. putredisilvae* CGMCC 2.3789^T^. E, F. *Yam. sympetra* CGMCC 2.3642^T^. G-I. *Aus. ficea* CGMCC 2.6956^T^. Scale bars: B, D, F, H, I = 10 μm.

###### Etymology

the specific epithet *planticola* refers to the type strain isolated from plant.

###### Culture characteristics

On YM agar, after 1 month at 25 °C, the streak culture is white, butyrous, rough. The margin is fimbriate (Fig. 39A). In YM broth, after 3 days at 25 °C, cells are oval, ellipsoidal and fusiform, 1.14–3.69 × 2.24–4.85 µm and single, budding is polar (Fig. 39B), after 7 days at 25 °C, a sediment, ring and pellicle are formed. After 1 month at 25 °C, sediment, ring and pellicle are present. In Dalmau plate culture on corn meal agar, hyphae are formed. Asci and ascospores are not observed on V8 and Acetate agar.

###### Physiological and biochemical characteristics

Glucose (weak) fermentation is present. Lactose, galactose, sucrose, maltose and raffinose fermentation are absent.Glucose, galactose, L-sorbose, sucrose, maltose, cellobiose (latent), trehalose, raffinose (latent, weak), melezitose (latent), inulin (latent), soluble starch (latent, weak), D-xylose, L-arabinose (latent), D-arabinose (latent), D-ribose (latent), L-rhamnose (latent), D-glucosamine (latent), N-acetyl-D-glucosamine (latent), ethanol, glycerol, erythritol, ribitol (latent), D-mannitol, D-glucitol, methyl-α-D-glucoside, salicin (latent), DL-lactic acid, succinic acid and citric acid (latent, weak) are assimilated as sole carbon sources. Lactose, melibiose, methanol, galactitol, inositol, hexadecane and D-glucuronic acid are not assimilated. Ammonium sulfate, potassium nitrate, sodium nitrite, L-lysine, ethylamine and cadaverine are assimilated as sole nitrogen sources. The maximum growth temperature is 32 °C. Growth in the vitamin-free medium is positive. Starch-like substances are not produced. Growth on 50 % (w/w) glucose-yeast extract agar is negative.

Physiologically, *Y. planticola* differs from the closely related species *Y. betulicola* (Fig. 17) in its inability to ferment galactose and grow with 50% glucose and the ability to assimilate raffinose and DL-lactic acid (Table S2.16).

###### Typus

**China**, Sejila Mountain, Nyingchi City, Tibet, obtained from the plant of *Liliaceae*, Aug. 2019, *Q.- M. Wang* (**holotype** CGMCC 2.8167^T^ preserved in a metabolically inactive state, ex-type cultures are preserved in culture collections under the numbers XZY144M2).

***Yamadazyma putredisilvae*** Q.-M. Wang, A.-H. Li, M.-M. Liu, H.-H. Zhu & F.-Y. Bai, ***sp. nov.*** Fungal Names FN 572782. Fig. 39C, D.

###### Etymology

the specific epithet *putredisilvae* refers to the type strain isolated from rotten wood.

###### Culture characteristics

On YM agar, after 1 month at 25 °C, the streak culture is white, butyrous, smooth and pale glossy. The margin is entire (Fig. 39C). In YM broth, after 3 days at 25 °C, cells are subglobose, oval and ellipsoidal, 2.48–4.94 × 3.76–6.21 µm and single, budding is polar (Fig. 39D), after 7 days at 25 °C, a sediment is formed. After 1 month at 25 °C, sediment is present, but a ring and pellicle do not appear. In Dalmau plate culture on corn meal agar, pseudohyphae are not formed. Asci and ascospores are not observed on V8 and Acetate agar.

###### Physiological and biochemical characteristics

Glucose and galactose (weak) fermentation are present. Lactose, sucrose, maltose and raffinose fermentation are absent. Glucose, galactose, L-sorbose, sucrose, maltose, cellobiose, trehalose (variable), lactose, melibiose, raffinose, melezitose, inulin, soluble starch, D-xylose, L-arabinose, D-arabinose (latent), D-ribose, L-rhamnose, D-glucosamine, N-acetyl-D-glucosamine, ethanol, glycerol, erythritol, ribitol, galactitol (variable), D-mannitol, D-glucitol, methyl-α-D-glucoside, salicin, DL-lactic acid (variable), succinic acid (weak) and citric acid (variable) are assimilated as sole carbon sources. Methanol, inositol, hexadecane and D-glucuronic acid are not assimilated. Ammonium sulfate, potassium nitrate, sodium nitrite, L-lysine, ethylamine and cadaverine are assimilated as sole nitrogen sources. The maximum growth temperature is 37 °C. Growth in the vitamin-free medium is positive. Starch-like substances are not produced. Growth on 50 % (w/w) glucose-yeast extract agar is positive (variable).

Physiologically, *Y. putredisilvae* differs from the closely related species *Y. ubonensis* (Fig. 17) in its inability to assimilate D-glucuronic acid, and the ability to ferment galactose and to assimilate L-sorbose, lactose, melibiose, raffinose, inulin, soluble starch, D-arabinose, D-ribose, N-acetyl-D-glucosamine, succinic acid, potassium nitrate, sodium nitrite, and grow in vitamin-free medium (Table S2.16).

###### Typus

**China**, Hainan province, obtained from rotten wood, Jan. 2008, *Q.-M. Wang* (**holotype** CGMCC 2.3789^T^ preserved in a metabolically inactive state, ex-type cultures are preserved in culture collections under the numbers CBS 18411 and X2WZ03-6).

###### Other culture examined

**China**, Wuzhishan, Hainan province, obtained from rotten wood, Jan. 2008, isolated by *Q.M. Wang*, culture CGMCC 2.3793.

***Yamadazyma sympetra*** Q.-M. Wang, A.-H. Li, M.-M. Liu, H.-H. Zhu & F.-Y. Bai, ***sp. nov.*** Fungal Names FN 572783. Fig. 39E, F.

###### Etymology

the specific epithet *sympetra* refers to *Sympetrum,* the insect genus from which the type strain was isolated.

###### Culture characteristics

On YM agar, after 1 month at 25 °C, the streak culture is white, butyrous, smooth and glossy. The margin is fimbriate (Fig. 39E). In YM broth, after 3 days at 25 °C, cells are subglobose, oval, ellipsoidal and cylindrical, 2.82–3.51 × 4.04–5.97 µm and single, budding is polar (Fig. 39F), after 7 days at 25 °C, a sediment and incomplete pellicle are formed. After 1 month at 25 °C, sediment and incomplete pellicle are present, and a ring does not appear. In Dalmau plate culture on corn meal agar, pseudohyphae are not formed. Asci and ascospores are not observed on V8 and Acetate agar.

###### Physiological and biochemical characteristics

Glucose, galactose (variable) and raffinose (variable) fermentation are present. Lactose, sucrose and maltose fermentation are absent. Glucose, galactose, L-sorbose, sucrose, maltose, cellobiose (variable), trehalose, lactose, melibiose, raffinose, melezitose, inulin, soluble starch, D-xylose, L-arabinose, D-arabinose, D-ribose, L-rhamnose, D-glucosamine, N-acetyl-D-glucosamine, methanol (variable), ethanol, glycerol, erythritol, ribitol, galactitol (variable), D-mannitol, D-glucitol, methyl-α-D-glucoside (latent), salicin, DL-lactic acid (variable), succinic acid, citric acid (variable), inositol (variable), hexadecane (variable) and D-glucuronic acid (variable) are assimilated as sole carbon sources. Ammonium sulfate, potassium nitrate, sodium nitrite (variable), L-lysine, ethylamine and cadaverine are assimilated as sole nitrogen sources. The maximum growth temperature is 42 °C. Growth in the vitamin-free medium is positive. Starch-like substances are not produced. Growth on 50 % (w/w) glucose-yeast extract agar is positive (variable).

Physiologically, *Y. sympetra* differs from the closely related species *Y. diddensiae*, *Y. putridisilvae*, *Y. ubonensis*, *Y. phyllophila*, *Y. paraphyllophila*, *Y. akitaensis*, *Y. nakazawae*, *Y. philogaea*, *Y. siamensis*, *Y. kanchanaburiensis*, *Y. dendronema*, *Y. germanica*, *Y. vaughaniae*, *Y. naeodendra*, *Y. laniorum*, *Y. tenuis* and *Y. follicola* (Fig. 17) in its ability to assimilate carbon sources and nitrogen sources (Table S2.16).

###### Typus

**China**, Beijing, obtained from *Sympetrum kunckeli*, Jul. 2007, *F.Y. Bai* (**holotype** CGMCC 2.3642^T^ preserved in a metabolically inactive state, ex-type cultures are preserved in culture collections under the numbers CBS 18382 and HQ3.4).

###### Other culture examined

**China**, Beijing, obtained from *Sympetrum kunckeli*, Feb. 2007, isolated by *F.Y. Bai*, culture CGMCC 2.3657; **China**, Beijing, obtained from inside an insect, isolated by *F.Y. Bai*, culture CGMCC 2.3716; **China**, Beijing, obtained from inside an insect, isolated by *F.Y. Bai*, culture CGMCC 2.3714 (= CBS 18412).

New taxa in the *Metschnikowiaceae* (*Serinales*, *Pichiomycetes*, *Saccharomycotina*) *Australozyma ficea* Q.-M. Wang, A.-H. Li, M.-M. Liu, H.-H. Zhu & F.-Y. Bai, *sp. nov.* Fungal Names FN 572784. Fig. 39G-I.

###### Etymology

the specific epithet *ficea* refers to *Ficus*, the plant genus from which the type strain was isolated.

###### Culture characteristics

On YM agar, after 1 month at 25 °C, the streak culture is yellowish, butyrous, smooth. The margin is entire (Fig. 39G). In YM broth, after 3 days at 25 °C, cells are oval and cylindrical, 1.63–4.73 × 4.02–5.87 µm and single, budding is polar (Fig. 39H), after 7 days at 25 °C, a sediment is formed. After 1 month at 25 °C, sediment, ring and pellicle are present. In Dalmau plate culture on corn meal agar, pseudohyphae are not formed. On V8 agar, the ascus is fusiform and contains one or two ascospores (Fig. 39I). The ascospores are spherical or subspherical, 1.24–2.44 × 1.64–2.97 µm.

###### Physiological and biochemical characteristics

Glucose, galactose and sucrose fermentation are present. Lactose, maltose and raffinose fermentation are absent. Glucose, galactose, L-sorbose, sucrose, maltose, trehalose, raffinose (weak), melezitose (latent), inulin, soluble starch (weak), D-xylose (latent), D-arabinose (latent), D-ribose (weak), L-rhamnose (weak), ethanol (latent), glycerol (latent), ribitol (latent), D-mannitol, D-glucitol, methyl-α-D-glucoside (latent), succinic acid, citric acid and inositol (weak) are assimilated as sole carbon sources. Cellobiose, lactose, melibiose, L-arabinose, D-glucosamine, N-acetyl-D-glucosamine, methanol, erythritol, galactitol, salicin, DL-lactic acid, hexadecane and D-glucuronic acid are not assimilated. Ammonium sulfate, potassium nitrate (latent), sodium nitrite (latent), L-lysine, ethylamine and cadaverine are assimilated as sole nitrogen sources. The maximum growth temperature is 32 °C. Growth in the vitamin-free medium is positive. Starch-like substances are not produced. Growth on 50 % (w/w) glucose-yeast extract agar is positive (weak).

Physiologically, *A. ficea* differs from the closely related species *A. saopauloensis*, *A. picinguabensis*, *A. tocantinsensis*, *A. robnettiae*, *A. touchengensis*, *A. bambusicola*, *A. nongkhaiensis*, *A. succicola* and *A. saccharicola* (Fig. 18) in its ability to assimilate raffinose, inulin, soluble starch, inositol, potassium nitrate and sodium nitrite (Table S2.17).

###### Typus

**China**, Lufeng County, Chuxiong, Yunnan Province, obtained from *Ficus microcarpa*, Oct. 2019, *Q.-M. Wang* (**holotype** CGMCC 2.6956^T^ preserved in a metabolically inactive state, ex-type culture is preserved in culture collections under the numbers LF17-2).

###### Other culture examined

**China**, Lufeng County, Chuxiong, Yunnan Province, obtained from *Ficus microcarpa*, Oct. 2019, isolated by *Q.M. Wang*, culture LF117-1.

***Clavispora pieris*** Q.-M. Wang, A.-H. Li, M.-M. Liu, H.-H. Zhu & F.-Y. Bai, ***sp. nov.*** Fungal Names FN 572785. Fig. 40A, B.

**Fig. 40.**
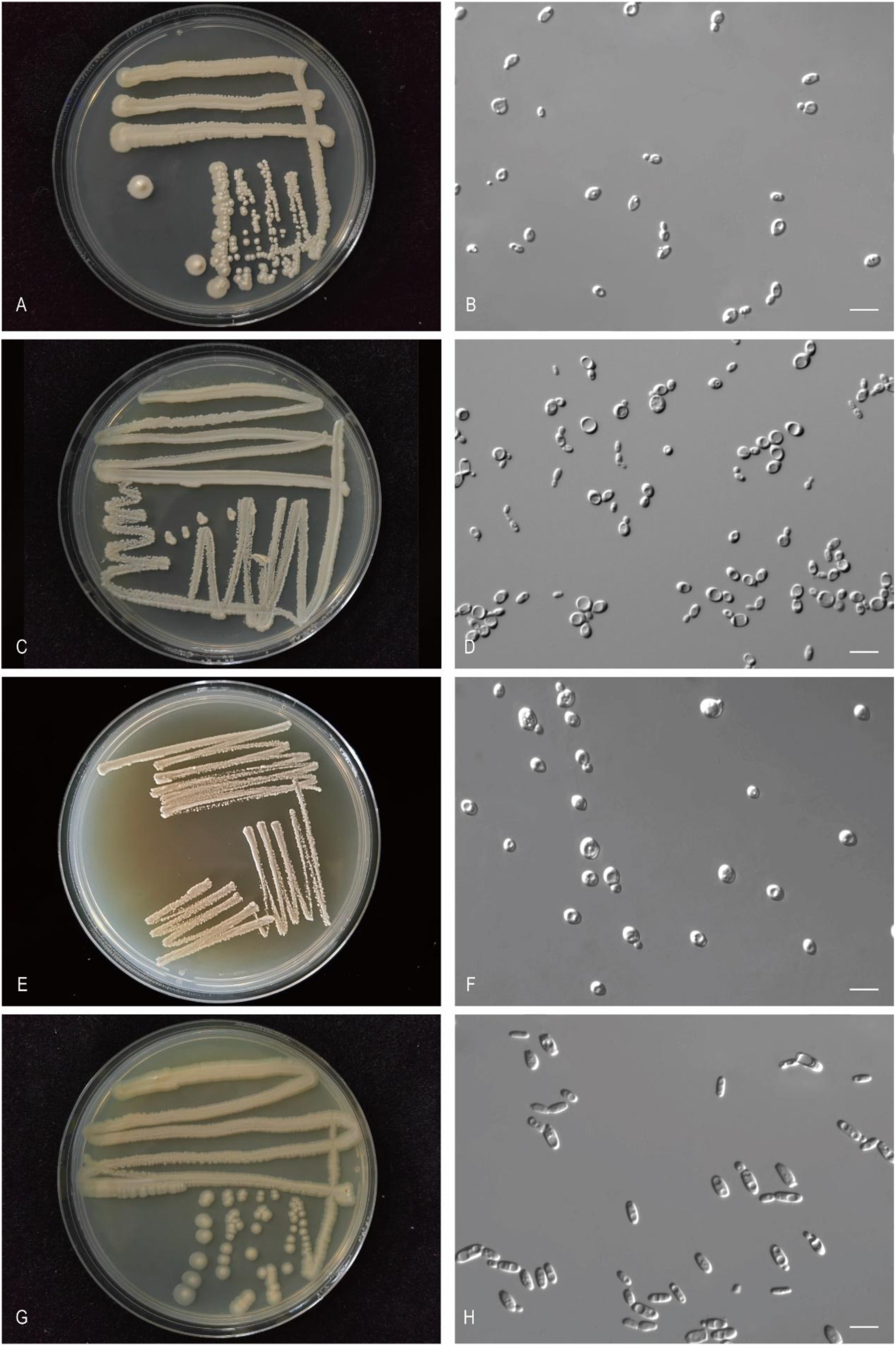
The streak culture grown in YM agar and vegetative cells grown in YM broth for 7 d at 25 °C. A, B. *Cla. pieris* CGMCC 2.3228^T^. C, D. *Cla. yunnanensis* CGMCC 2.6998^T^. E, F. *Met. allorhynchi* CGMCC 2.3708^T^. G, H. *Met. hunanensis* CGMCC 2.3430^T^. Scale bars: B, D, F, H = 10 μm.

###### Etymology

the specific epithet *pieris* refers to *Pieris*, the plant genus from which the type strain was isolated.

###### Culture characteristics

On YM agar, after 1 month at 25 °C, the streak culture is white, butyrous, smooth and pale glossy. The margin is entire (Fig. 40A). In YM broth, after 3 days at 25 °C, cells are oval and ellipsoidal, 2.37–4.58 × 4.13–6.82 µm and single, budding is polar (Fig. 40B), after 7 days at 25 °C, a sediment is formed. After 1 month at 25 °C, sediment is present, pellicle and ring do not appear. In Dalmau plate culture on corn meal agar, pseudohyphae are not formed. Asci and ascospores are not observed on V8 and Acetate agar.

###### Physiological and biochemical characteristics

Glucose, sucrose and maltose fermentation are present. Lactose, galactose and raffinose fermentation are absent. Glucose, galactose, L-sorbose, sucrose, maltose, trehalose, melibiose (weak), raffinose, melezitose, inulin, soluble starch (latent, weak), D-xylose (weak), D-glucosamine, N-acetyl-D-glucosamine, ethanol, glycerol, ribitol, D-mannitol, D-glucitol, methyl-α-D-glucoside, DL-lactic acid, succinic acid and citric acid are assimilated as sole carbon sources. Cellobiose, lactose, L-arabinose, D-arabinose, D-ribose, L-rhamnose, methanol, erythritol, galactitol, salicin, inositol, hexadecane and D-glucuronic acid are not assimilated. Ammonium sulfate, potassium nitrate, sodium nitrite, L-lysine, ethylamine and cadaverine are assimilated as sole nitrogen sources. The maximum growth temperature is 35 °C. Growth in the vitamin-free medium is positive. Starch-like substances are not produced. Growth on 50 % (w/w) glucose-yeast extract agar is negative.

Physiologically, *C. pieris* does not differ from the closely related species *C. yunnanensis*, *C. carvajalis*, *C. phyllophila*, *C. fructus*, *C. asparagi*, *C. vitiphila*, *C. santaluciae*, *C. pruni*, *C. paralusitaniae* and *C. lusitaniae* (Fig. 18) in its ability to utilize carbon sources and nitrogen sources (Table S2.18).

###### Typus

**China**, Hunan province, obtained from *Pieris rapae*, Nov. 2006, *Q.-M. Wang* (**holotype** CGMCC 2.3228^T^ preserved in a metabolically inactive state, ex-type cultures are preserved in culture collections under the numbers CFD1.4).

###### Other culture examined

**China**, Lufeng County, Chuxiong, Yunnan Province, obtained from leaf of *Morus alba*, Oct. 2019, isolated by *Q.M. Wang*, culture CGMCC 2.8160 (= LF23-1).

***Clavispora yunnanensis*** Q.-M. Wang, A.-H. Li, M.-M. Liu, H.-H. Zhu & F.-Y. Bai, ***sp. nov.*** Fungal Names FN 572786. Fig. 40C, D.

###### Etymology

the specific epithet *yunnanensis* refers to the geographic origin of the type strain, Yunnan province.

###### Culture characteristics

On YM agar, after 1 month at 25 °C, the streak culture is white, butyrous, smooth and glossy. The margin is entire (Fig. 40C). In YM broth, after 3 days at 25 °C, cells are oval and cylindrical, 1.87–5.28 × 5.31–7.08 µm and single, budding is polar (Fig. 40D), after 7 days at 25 °C, a sediment is formed. After 1 month at 25 °C, sediment is present, pellicle and ring do not appear. In Dalmau plate culture on corn meal agar, pseudohyphae are formed. Asci and ascospores are not observed on V8 and Acetate agar.

###### Physiological and biochemical characteristics

Glucose and galactose fermentation are present. Lactose, sucrose, maltose and raffinose fermentation are absent. Glucose, L-sorbose, sucrose (latent), maltose (latent), cellobiose (latent), trehalose, raffinose (weak), melezitose (latent), inulin (latent), soluble starch (weak), D-xylose (latent), L-rhamnose (weak), D-glucosamine (latent), N-acetyl-D-glucosamine, ethanol (latent), glycerol (latent), ribitol (latent), D-mannitol, D-glucitol (latent), methyl-α-D-glucoside (latent), salicin (latent), DL-lactic acid (weak), succinic acid, citric acid and D-glucuronic acid (weak) are assimilated as sole carbon sources. Galactose, lactose, melibiose, L-arabinose, D-arabinose, D-ribose, methanol, erythritol, galactitol, inositol and hexadecane are not assimilated. Ammonium sulfate, potassium nitrate, sodium nitrite, L-lysine, ethylamine and cadaverine are assimilated as sole nitrogen sources. The maximum growth temperature is 30 °C. Growth in the vitamin-free medium is positive. Starch-like substances are not produced. Growth on 50 % (w/w) glucose-yeast extract agar is negative.

Physiologically, *C. yunnanensis* differs from the closely related species *C. carvajalis*, *C. phyllophila*, *C. fructus*, *C. asparagi*, *C. vitiphila*, *C. santaluciae* and *C. pruni* (Fig. 18) in its ability to assimilate L-rhamnose (Table S2.18).

###### Typus

**China**, Xiongxian County, Zhaotong city, Yunnan Province, obtained from flower, Oct. 2019, *Q.- M. Wang* (**holotype** CGMCC 2.6998^T^ preserved in a metabolically inactive state, ex-type cultures are preserved in culture collections under the numbers YN87-1).

###### Other culture examined

**China**, Lufeng County, Chuxiong, Yunnan Province, obtained from rotten wood, Oct. 2019, isolated by *Q.M. Wang*, culture LF5-1B.

***Metschnikowia allorhynchi*** Q.-M. Wang, A.-H. Li, M.-M. Liu, H.-H. Zhu & F.-Y. Bai, ***sp. nov.*** Fungal Names FN 572787. Fig. 40E, F.

###### Etymology

the specific epithet *allorhynchi* refers to *Allorhynchium*, the insect genus from which the type strain was isolated.

###### Culture characteristics

On YM agar, after 1 month at 25 °C, the streak culture is yellowish-white, butyrous, smooth and pale glossy. The margin is entire (Fig. 40E). In YM broth, after 3 days at 25 °C, cells are subglobose and oval, 3.24–6.39 × 4.18–7.17 µm and single, budding is polar (Fig. 40F), after 7 days at 25 °C, a sediment is formed. After 1 month at 25 °C, the sediment ring and are present, pellicle does not appear. In Dalmau plate culture on corn meal agar, pseudohyphae are not formed. Asci and ascospores are not observed on V8 and Acetate agar.

###### Physiological and biochemical characteristics

Glucose and sucrose (weak) fermentation are present. Lactose, galactose, maltose and raffinose fermentation are absent. Glucose, galactose, L-sorbose, sucrose, maltose, cellobiose, trehalose, melezitose, inulin, soluble starch (latent, weak), D-glucosamine (latent, weak), N-acetyl-D-glucosamine (latent, weak), ethanol (latent), glycerol (latent), ribitol (weak), D-mannitol, D-glucitol and citric acid are assimilated as sole carbon sources. Lactose, melibiose, raffinose, D-xylose, L-arabinose, D-arabinose, D-ribose, L-rhamnose, methanol, erythritol, galactitol, methyl-α-D-glucoside, salicin, DL-lactic acid, succinic acid, inositol, hexadecane and D-glucuronic acid are not assimilated. L-lysine, ethylamine and cadaverine are assimilated as sole nitrogen sources. Ammonium sulfate, potassium nitrate and sodium nitrite are not assimilated. The maximum growth temperature is 30 °C. Growth in the vitamin-free medium is positive (weak). Starch-like substances are not produced. Growth on 50 % (w/w) glucose-yeast extract agar is negative.

Physiologically, *M. allorhynchi* differs from the closely related species *M. noctiluminum*, *M. kofuensis* and *M. viticola* (Fig. 19) in its inability to assimilate salicin and the ability grow in vitamin-free medium (Table S2.19).

###### Typus

**China**, Beijing, obtained from *Allorhynchium argentatum*, Oct. 2007, *F.-Y. Bai* (**holotype** CGMCC 2.3708^T^ preserved in a metabolically inactive state, ex-type culture is preserved in culture collections under the numbers NHF4.1).

###### Other culture examined

**China**, Beijing, obtained from *Allorhynchium argentatum*, Oct. 2007, isolated by *F.-Y. Bai*, culture NHF4.2 (= CGMCC 2.3707).

***Metschnikowia hunanensis*** Q.-M. Wang, A.-H. Li, M.-M. Liu, H.-H. Zhu & F.-Y. Bai, ***sp. nov.*** Fungal Names FN 572788. Fig. 40G, H.

###### Etymology

the specific epithet *hunanensis* refers to the geographic origin of the type strain, Hunan province.

###### Culture characteristics

On YM agar, after 1 month at 25 °C, the streak culture is yellowish-white, butyrous, warty and pale glistening. The margin is eroded (Fig. 40G). In YM broth, after 3 days at 25 °C, cells are ellipsoidal and cylindrical, 2.01–5.13 × 4.76–6.97 µm and single, budding is polar (Fig. 40H), after 7 days at 25 °C, a sediment is formed. After 1 month at 25 °C, sediment and ring are present, but pellicle does not appear. In Dalmau plate culture on corn meal agar, pseudohyphae are not formed. Asci and ascospores are not observed on V8 and Acetate agar.

###### Physiological and biochemical characteristics

Glucose fermentation is present. Lactose, galactose, sucrose, maltose and raffinose fermentation are absent. Glucose, galactose, L-sorbose, sucrose, maltose, cellobiose, trehalose (weak), melezitose, inulin, soluble starch (weak), D-xylose (latent, weak), D-ribose (latent), D-glucosamine (weak), N-acetyl-D-glucosamine, ethanol, glycerol, ribitol (latent), D-mannitol, D-glucitol, methyl-α-D-glucoside (latent), salicin and succinic acid are assimilated as sole carbon sources. Lactose, melibiose, raffinose, L-arabinose, D-arabinose, L-rhamnose, methanol, erythritol, galactitol, DL-lactic acid, citric acid, Inositol, hexadecane and D-glucuronic acid are not assimilated. Potassium nitrate, sodium nitrite (latent, weak), L-lysine, ethylamine and cadaverine are assimilated as sole nitrogen sources. Ammonium sulfate is not assimilated. The maximum growth temperature is 35 °C. Growth in the vitamin-free medium is positive. Starch-like substances are not produced. Growth on 50 % (w/w) glucose-yeast extract agar is negative.

Physiologically, *M. hunanensis* does not differ from the closely related species *M. follicola*, *M. reukaufii* and *M. cibodasensis* (Fig. 19) in its ability to assimilate carbon sources and nitrogen sources (Table S2.19).

###### Typus

**China**, Hunan province, obtained from the flower of *Pharbitis nil*, Jan. 2007, *F.-Y. Bai* (**holotype** CGMCC 2.3430^T^ preserved in a metabolically inactive state, ex-type culture is preserved in culture collections under the numbers QN2.1).

###### Other culture examined

**China**, Hunan province, obtained from a bee, Jan. 2007, isolated by *F.-Y. Bai*, culture BeeM1.7 (= CGMCC 2.3437).

***Metschnikowia follicola*** Q.-M. Wang, A.-H. Li, M.-M. Liu, H.-H. Zhu & F.-Y. Bai, ***sp. nov.*** Fungal Names FN 572789. Fig. 41A, B.

**Fig. 41.**
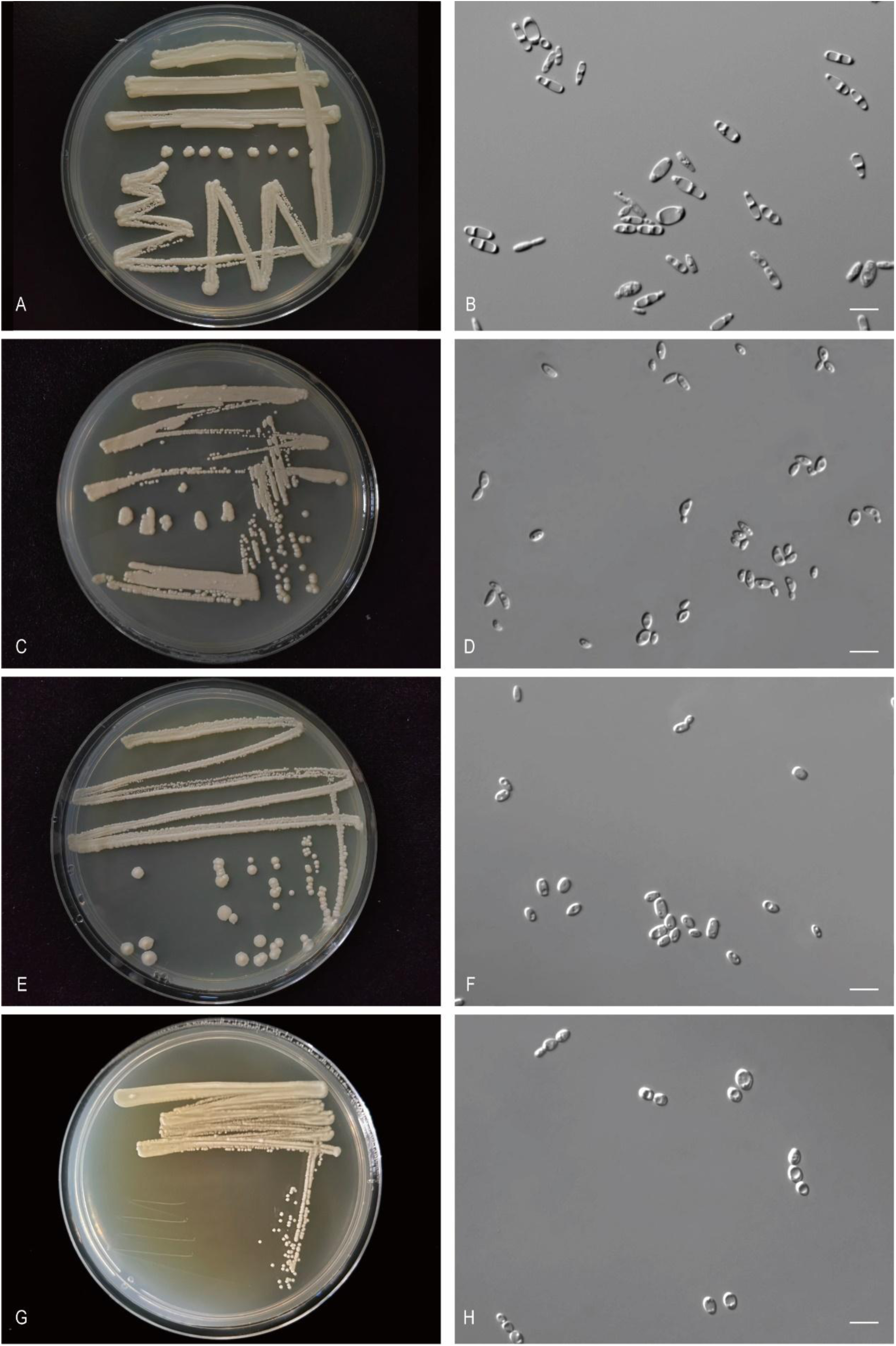
The streak culture grown in YM agar and vegetative cells grown in YM broth for 7 d at 25 °C. A, B. *Met. follicola* CGMCC 2.6949^T^. C, D. *Met. lycaenae* CGMCC 2.3740^T^. E, F. *Met. mantia* CGMCC 2.3753^T^. G, H. *Met. planticola* CGMCC 2.2746^T^. Scale bars: B, D, F, H = 10 μm.

###### Etymology

the specific epithet *follicola* refers to the type strain isolated from leaf.

###### Culture characteristics

On YM agar, after 1 month at 25 °C, the streak culture is white, butyrous, smooth and pale glossy. The margin is entire (Fig. 41A). In YM broth, after 3 days at 25 °C, cells are fusiform and cylindrical, 2.59–4.66 × 3.37–5.44 µm and single, budding is polar (Fig. 41B), after 7 days at 25 °C, a sediment and ring are formed. After 1 month at 25 °C, sediment and ring are present, but a pellicle does not appear. In Dalmau plate culture on corn meal agar, pseudohyphae are not formed. Asci and ascospores are not observed on V8 and Acetate agar.

###### Physiological and biochemical characteristics

Glucose, galactose (variable), sucrose (variable), maltose (variable) and raffinose (variable) fermentation are present. Lactose fermentation is absent. Glucose, galactose (variable), L-sorbose (variable), sucrose, maltose, cellobiose, trehalose (latent), lactose (variable), melibiose (variable), raffinose (variable), melezitose (latent), inulin (latent), soluble starch (variable), D-xylose (variable), L-arabinose (variable), D-arabinose (variable), D-ribose (variable), L-rhamnose (variable), D-glucosamine (latent), N-acetyl-D-glucosamine (latent), ethanol (latent), glycerol (latent), erythritol (variable), ribitol (variable), D-mannitol, D-glucitol (variable), methyl-α-D-glucoside (latent), salicin (variable), succinic acid (variable), citric acid(variable), inositol (variable) and D-glucuronic acid (variable) are assimilated as sole carbon sources. Methanol, galactitol, DL-lactic acid and hexadecane are not assimilated. Ammonium sulfate (variable), potassium nitrate (variable), sodium nitrite, L-lysine, ethylamine and cadaverine are assimilated as sole nitrogen sources. The maximum growth temperature is 42 °C. Growth in the vitamin-free medium is positive. Starch-like substances are not produced. Growth on 50 % (w/w) glucose-yeast extract agar is positive(variable).

Physiologically, *M. follicola* differs from the closely related species *M. hunanensis*, *M. reukaufii* and *M. cibodasensis* (Fig. 19) in its ability to ferment maltose (Table S2.19).

###### Typus

**China**, Katinggou, Nyingchi City, Tibet, obtained from flower, Aug. 2019, *Q.-M. Wang* (**holotype** CGMCC 2.6949^T^ preserved in a metabolically inactive state, ex-type cultures are preserved in culture collections under the numbers XZY30-5).

###### Other culture examined

**China**, Katinggou, Nyingchi City, Tibet, obtained from plant of *Liliaceae*, Aug. 2019, isolated by *Q.M. Wang*, culture XZY30-2. **China**, Sejila Mountain, Nyingchi, Tibet, obtained from plant, Aug. 2019, isolated by *Q.M. Wang*, culture XZY40A. **China**, Sejila Mountain, Nyingchi, Tibet, obtained from leaf of *Piptanthus concolor*, Aug. 2019, isolated by *Q.M. Wang*, culture XZY279-3. **China**, Gongbujiangda, Nyingchi, Tibet, obtained from plant of *Brassicaceae*, Aug. 2019, isolated by *Q.M. Wang*, culture XZY3F2. **China**, Katinggou, Nyingchi City, Tibet, obtained from flower, Aug. 2019, isolated by *Q.M. Wang*, culture XZY19-6. **China**, Katinggou, Nyingchi City, Tibet, obtained from flower, Aug. 2019, isolated by *Q.M. Wang*, culture XZY19-5 (= CGMCC 2.6945). **China**, Sejila Mountain, Nyingchi, Tibet, obtained from *Gentiana crassicaulis*, Aug. 2019, isolated by *Q.M. Wang*, culture XZY39-4. **China**, Xincuo Village, Nyingchi City, Tibet, obtained from *Sanguisorba officinalis*, Aug. 2019, isolated by *Q.M. Wang*, culture XZY257-2-1. **China**, Gongbujiangda, Nyingchi, Tibet, obtained from flower, Aug. 2019, isolated by *Q.M. Wang*, culture XZY3-2. **China**, Gongbujiangda, Nyingchi, Tibet, obtained from flower, Aug. 2019, isolated by *Q.M. Wang*, culture XZY3-3. **China**, Sejila Mountain, Nyingchi, Tibet, obtained from *Gentiana crassicaulis*, Aug. 2019, isolated by *Q.M. Wang*, culture XZY39-2. **China**, Sejila Mountain, Nyingchi, Tibet, obtained from plant, Aug. 2019, isolated by *Q.M. Wang*, culture XZY40-1 (= CGMCC 2.6951). **China**, Katinggou, Nyingchi City, Tibet, obtained from flower, Aug. 2019, isolated by *Q.M. Wang*, culture XZY30 (= CGMCC 2.6947). **China**, Baiba County, Linzhi, Tibet, obtained from flower, Aug. 2019, isolated by *Q.M. Wang*, culture XZY87 (= CGMCC 2.6948). **China**, Katinggou, Nyingchi City, Tibet, obtained from *Pilea swinglei*, Aug. 2019, isolated by *Q.M. Wang*, culture XZY14A (= CGMCC 2.6946). **China**, Tibet, obtained from leaf, Dec. 2004, isolated by *Q.M. Wang*, culture CGMCC 2.2651. **China**, Gongbujiangda, Nyingchi, Tibet, obtained from flower, Aug. 2019, isolated by *Q.M. Wang*, culture XZY3-4 (= CGMCC 2.6950). **China**, Sejila Mountain, Nyingchi, Tibet, obtained from *Piptanthus nepalensis*, Aug. 2019, isolated by *Q.M. Wang*, culture XZY279-2 (= CGMCC 2.6952). **China**, Xincuo Village, Nyingchi City, Tibet, obtained from *Sanguisorba officinalis*, Aug. 2019, isolated by *Q.M. Wang*, culture XZY257-1 (= CGMCC 2.6953). **China**, Tibet, obtained from leaf, Apr. 2005, isolated by *Q.M. Wang*, culture CGMCC 2.2800 (= CBS 18394).

***Metschnikowia lycaenae*** Q.-M. Wang, A.-H. Li, M.-M. Liu, H.-H. Zhu & F.-Y. Bai, ***sp. nov.*** Fungal Names FN 572790. Fig. 41C, D.

###### Etymology

the specific epithet *lycaenae* refers to *Lycaena*, the insect genus from which the type strain was isolated.

###### Culture characteristics

On YM agar, after 1 month at 25 °C, the streak culture is yellowish-white, butyrous, and pale glossy. The margin is entire (Fig. 41C). In YM broth, after 3 days at 25 °C, cells are ellipsoidal, 2.06–3.80 × 3.78–6.23 µm and single, budding is polar (Fig. 41D), after 7 days at 25 °C, sediment is formed. After 1 month at 25 °C, sediment and ring are present, but pellicle does not appear. In Dalmau plate culture on corn meal agar, pseudohyphae are not formed. Asci and ascospores are not observed on V8 and Acetate agar.

###### Physiological and biochemical characteristics

Glucose fermentation is present. Lactose, galactose, sucrose, maltose and raffinose fermentation are absent. Glucose, galactose, L-sorbose, sucrose, maltose, cellobiose, trehalose, raffinose (variable), melezitose, inulin, soluble starch (latent, weak), D-xylose (variable), D-glucosamine, N-acetyl-D-glucosamine, ethanol (variable), glycerol, ribitol, D-mannitol, D-glucitol, methyl-α-D-glucoside (latent), salicin, DL-lactic acid (variable), succinic acid and citric acid (variable) are assimilated as sole carbon sources. Lactose, melibiose, L-arabinose, D-arabinose, D-ribose, L-rhamnose, methanol, erythritol, galactitol, inositol, hexadecane and D-glucuronic acid are not assimilated. Ammonium sulfate (latent), potassium nitrate (latent, weak), sodium nitrite, L-lysine (weak), ethylamine and cadaverine are assimilated as sole nitrogen sources. The maximum growth temperature is 42 °C. Growth in the vitamin-free medium is positive. Starch-like substances are not produced. Growth on 50 % (w/w) glucose-yeast extract agar is negative.

Physiologically, *M. lycaenae* differs from the closely related species *C. golubevii* and *M. lunata* (Fig. 19) in its inability to assimilate hexadecane and D-glucuronic acid, and the ability to assimilate inulin, soluble starch, potassium nitrate, sodium nitrite and grow in vitamin-free medium (Table S2.19).

###### Typus

**China**, Beijing, obtained from gut of *Lycaena dispar*, Oct. 2007, *F.-Y. Bai* (**holotype** CGMCC 2.3740^T^ preserved in a metabolically inactive state, ex-type culture is preserved in culture collections under the numbers HHDE1.2.1).

###### Other culture examined

**China**, Beijing, obtained from the gut of *Lycaena dispar*, Oct. 2007, isolated by *F.-Y. Bai*, culture HHDE1.2.2(= CGMCC2.3741).

***Metschnikowia mantia*** Q.-M. Wang, A.-H. Li, M.-M. Liu, H.-H. Zhu & F.-Y. Bai, ***sp. nov.*** Fungal Names FN 572791. Fig. 41E, F.

###### Etymology

the specific epithet ***mantia*** refers to *Mantodea*, the insect genus from which the type strain was isolated.

###### Culture characteristics

On YM agar, after 1 month at 25 °C, the streak culture is whitish-cream, butyrous, smooth and glossy. The margin is entire (Fig. 41E). In YM broth, after 3 days at 25 °C, cells are ellipsoidal and cylindrical, 2.59–4.16 × 3.25–6.70 µm and single, budding is polar (Fig. 41F), after 7 days at 25 °C, a sediment is formed. After 1 month at 25 °C, sediment is present, pellicle and ring do not appear. In Dalmau plate culture on corn meal agar, pseudohyphae are not formed. Asci and ascospores are not observed on V8 and Acetate agar.

###### Physiological and biochemical characteristics

Glucose and galactose fermentation are present. Lactose, sucrose, maltose and raffinose fermentation are absent. Glucose, galactose, D-glucosamine (latent, weak), N-acetyl-D-glucosamine, D-mannitol and D-glucitol are assimilated as sole carbon sources. L-sorbose, sucrose, maltose, cellobiose, trehalose, lactose, melibiose, raffinose, melezitose, inulin, soluble starch, D-xylose, L-arabinose, D-arabinose, D-ribose, L-rhamnose, methanol, ethanol, glycerol, erythritol, ribitol, galactitol, methyl-α-D-glucoside, salicin, DL-lactic acid, succinic acid, citric acid, inositol, hexadecane and D-glucuronic acid are not assimilated. L-lysine and cadaverine are assimilated as sole nitrogen sources. Ammonium sulfate, potassium nitrate, sodium nitrite and ethylamine are not assimilated. The maximum growth temperature is 42 °C. Growth in the vitamin-free medium is negative. Starch-like substances are not produced. Growth on 50 % (w/w) glucose-yeast extract agar is negative.

Physiologically, *M. mantia* differs from the closely related species *C. golubevii*, *M. lunata*, *M. kunwiensis* and *M. lycaenae* (Fig. 19) in its inability to assimilate L-sorbose, maltose, cellobiose, trehalose, melezitose, glycerol, ribitol, methyl-α-D-glucoside, salicin and ethylamine (Table S2.19).

###### Typus

**China**, Beijing, obtained from *Mantodea* sp., Oct. 2007, *F.-Y. Bai* (**holotype** CGMCC 2.3753^T^ preserved in a metabolically inactive state).

***Metschnikowia planticola*** Q.-M. Wang, A.-H. Li, M.-M. Liu, H.-H. Zhu & F.-Y. Bai, ***sp. nov.*** Fungal Names FN 572792. Fig. 41G, H.

###### Etymology

the specific epithet *planticola* refers to the type strain isolated from a plant.

###### Culture characteristics

On YM agar, after 1 month at 25 °C, the streak culture is whitish, butyrous, smooth and glossy. The margin is entire (Fig. 41G). In YM broth, after 3 days at 25 °C, cells are ellipsoidal and oval, 3.27–5.00 × 5.04–7.25 µm and single, budding is polar (Fig. 41H), after 7 days at 25 °C, a sediment is formed. After 1 month at 25 °C, sediment is present, and the ring and pellicle do not appear. In Dalmau plate culture on corn meal agar, pseudohyphae are not formed. Asci and ascospores are not observed on V8 and Acetate agar.

###### Physiological and biochemical characteristics

Glucose (weak), galactose (variable), sucrose (variable) and maltose (weak) fermentation are present. Lactose and raffinose fermentation are absent. Glucose, galactose (variable), L-sorbose (latent), sucrose (latent), maltose (latent), cellobiose (variable), trehalose (latent), melezitose, inulin (latent), D-glucosamine (latent), N-acetyl-D-glucosamine (latent), ethanol (latent, weak), glycerol (latent, weak), ribitol (variable), galactitol (latent, weak), D-mannitol, D-glucitol (latent), methyl-α-D-glucoside (variable), salicin (latent, weak) and succinic acid (latent) are assimilated as sole carbon sources. Lactose, melibiose, raffinose, soluble starch, D-xylose, L-arabinose, D-arabinose, D-ribose, L-rhamnose, methanol, erythritol, DL-lactic acid, citric acid, inositol, hexadecane and D-glucuronic acid are not assimilated. Ammonium sulfate (latent, weak), potassium nitrate (variable), L-lysine (latent), ethylamine (latent) and cadaverine (latent) are assimilated as sole nitrogen sources. Sodium nitrite is not assimilated. The maximum growth temperature is 25 °C. Growth in the vitamin-free medium is positive. Starch-like substances are not produced. Growth on 50 % (w/w) glucose-yeast extract agar is positive.

Physiologically, *M. planticola* differs from the closely related species *M. rancensis* (Fig. 19) in its inability to assimilate D-ribose, citric acid, hexadecane and D-glucuronic acid, and the ability to ferment maltose and to assimilate galactitol and grow in vitamin-free medium (Table S2.19).

###### Typus

**China**, Tibet, obtained from plant leaves, Apr. 2005, *Q.-M. Wang* (**holotype** CGMCC 2.2746^T^ preserved in a metabolically inactive state, ex-type culture is preserved in culture collections under the numbers XZ67C2).

###### Other culture examined

**China**, Tibet, obtained from plant leaves, Apr. 2005, isolated by *Q.M. Wang*, culture CGMCC 2.2747 (= XZ63A4). **China**, Tibet, obtained from plant leaves, isolated by *Q.M. Wang*, culture CGMCC 2.2748. **China**, Sejila Mountain Forest Monitoring Station, Nyingchi, Tibet, obtained from rotten wood, Aug. 2019, isolated by *Q.M. Wang*, culture XZY148-4 (=CGMCC 2.6943).

***Metschnikowia taianensis*** Q.-M. Wang, A.-H. Li, M.-M. Liu, H.-H. Zhu & F.-Y. Bai, ***sp. nov.*** Fungal Names FN 572793. Fig. 42A, B.

**Fig. 42.**
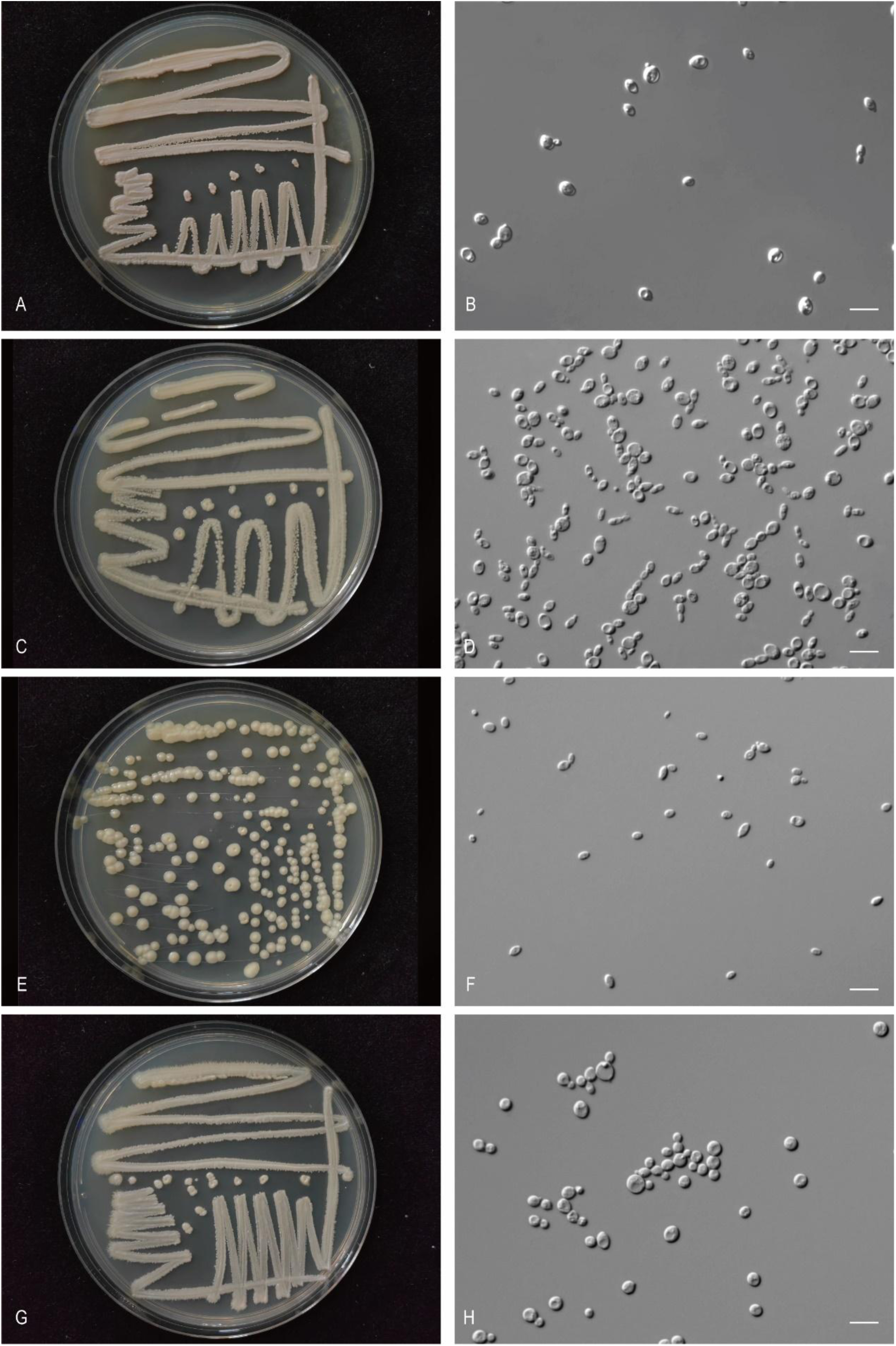
The streak culture grown in YM agar and vegetative cells grown in YM broth for 7 d at 25 °C. A, B. *Met. taianensis* CGMCC 2.4474^T^. C, D. *Met. betulae* CGMCC 2.7036^T^. E, F. *Met. xizangensis* CGMCC 2.6957^T^. G, H. *Sou. violae* CGMCC 2.6990^T^. Scale bars: B, D, F, H = 10 μm.

###### Etymology

the specific epithet *taianensis* refers to the geographic origin of the type strain, Taian, Shandong province.

###### Culture characteristics

On YM agar, after 1 month at 25 °C, the streak culture is pale-pink, butyrous, smooth. The margin is eroded (Fig. 42A). In YM broth, after 3 days at 25 °C, cells are ellipsoidal and oval, 2.75–4.10 × 3.68–6.15 µm and single, budding is polar (Fig. 42B), after 7 days at 25 °C, a sediment is formed. After 1 month at 25 °C, a sediment is present, ring and pellicle do not appear. In Dalmau plate culture on corn meal agar, pseudohyphae are not formed. Asci and ascospores are not observed on V8 and Acetate agar.

###### Physiological and biochemical characteristics

Glucose and galactose (weak) fermentation are present. Lactose, sucrose, maltose and raffinose fermentation are absent. Glucose, galactose, L-sorbose, sucrose, maltose, cellobiose, trehalose, melezitose, inulin, soluble starch, D-xylose, D-ribose (variable), D-glucosamine, N-acetyl-D-glucosamine, ethanol, glycerol, erythritol (variable), ribitol, D-mannitol, D-glucitol, methyl-α-D-glucoside, salicin, DL-lactic acid (latent, weak), succinic acid and citric acid are assimilated as sole carbon sources. Lactose, melibiose, raffinose, L-arabinose, D-arabinose, L-rhamnose, methanol, galactitol, inositol, hexadecane and D-glucuronic acid are not assimilated. Ammonium sulfate (variable), potassium nitrate, sodium nitrite, L-lysine, ethylamine and cadaverine are assimilated as sole nitrogen sources. The maximum growth temperature is 42 °C. Growth in the vitamin-free medium is positive.

Starch-like substances are not produced. Growth on 50 % (w/w) glucose-yeast extract agar is negative. Physiologically, *M. taianensis* differs from the closely related species *M. andauensis*, *M. chrysoperlae*, *M. leonuri*, *M. shanxiensis*, *M. sinensis*, *M. fructicola*, *M. rubicola*, *M. pulcherrima*, *M. ziziphicola*, *M. pulcherrima*, *M. pimensis* and *M. picachoensis* (Fig. 19) in its ability to assimilate potassium nitrate (Table S2.19).

###### Typus

**China**, obtained from bark, Aug. 2011, *Q.-M. Wang* (**holotype** CGMCC 2.4474^T^ preserved in a metabolically inactive state, ex-type cultures are preserved in culture collections under the numbers TAHT66-3 and CBS 18395).

###### Other culture examined

**China**, Taian, Shandong province, obtained from bark, Aug. 2011, isolated by *Q.M. Wang*, culture CGMCC 2.4473 (= TAHT66-2); **China**, obtained from bark, Aug. 2011, isolated by *Q.M. Wang*, culture CGMCC 2.4472 (= TAHT66-1).

***Metschnikowia betulae*** Q.-M. Wang, A.-H. Li, M.-M. Liu, H.-H. Zhu & F.-Y. Bai, ***sp. nov.*** Fungal Names FN 572794. Fig. 42C, D.

###### Etymology

the specific epithet *betulae* refers to *Betula*, the plant genus from which the type strain was isolated.

###### Culture characteristics

On YM agar, after 1 month at 25 °C, the streak culture is pale-yellow, butyrous, smooth and pale glossy. The margin is entire (Fig. 42C). In YM broth, after 3 days at 25 °C, cells are fusiform, ellipsoidal and oval, 1.58–3.53 × 2.71–5.16 µm and single, budding is polar (Fig. 42D), after 7 days at 25 °C, sediment and ring are formed. After 1 month at 25 °C, sediment and ring are present, but pellicle does not appear. In Dalmau plate culture on corn meal agar, pseudohyphae are not formed. Asci and ascospores are not observed on V8 and Acetate agar.

###### Physiological and biochemical characteristics

Glucose (weak) fermentation is present. Lactose, galactose, sucrose, maltose and raffinose fermentation are absent. Glucose, galactose, L-sorbose, sucrose, maltose, cellobiose, trehalose (latent), melibiose (variable), raffinose (variable), melezitose (latent), inulin (latent), soluble starch (variable), D-xylose (latent), L-arabinose (variable), D-glucosamine, N-acetyl-D-glucosamine, ethanol (latent), glycerol (latent), ribitol (latent), D-mannitol, D-glucitol, methyl-α-D-glucoside (latent), salicin (latent), DL-lactic acid (variable), succinic acid (latent) and citric acid (latent) are assimilated as sole carbon sources. Lactose, D-arabinose, D-ribose, L-rhamnose, methanol, erythritol, galactitol, inositol, hexadecane and D-glucuronic acid are not assimilated. Ammonium sulfate, potassium nitrate, sodium nitrite (weak), L-lysine, ethylamine and cadaverine are assimilated as sole nitrogen sources. The maximum growth temperature is 25 °C. Growth in the vitamin-free medium is positive. Starch-like substances are not produced. Growth on 50 % (w/w) glucose-yeast extract agar is positive (weak).

Physiologically, *M. betulae* differs from the closely related species *M. corniflorae* (Fig. 19) in its inability to assimilate D-ribose, hexadecane and D-glucuronic acid, and the ability to assimilate inulin, D-xylose, glycerol, succinic acid, potassium nitrate, sodium nitrite, and grow in vitamin-free medium and grow with 50% glucose (Table S2.19).

###### Typus

**China**, Basom lake, Linzhi, Tibet, obtained from bark of *Betula platyphylla*, Aug. 2019, *Q.-M. Wang* (**holotype** CGMCC 2.7036^T^ preserved in a metabolically inactive state, ex-type culture is preserved in culture collections under the numbers XZY138-7).

###### Other culture examined

**China**, Basom lake, Linzhi, Tibet, obtained from bark of *Betula platyphylla*, Aug. 2019, isolated by *Q.M. Wang*, culture XZY138M1.

***Metschnikowia xizangensis*** Q.-M. Wang, A.-H. Li, M.-M. Liu, H.-H. Zhu & F.-Y. Bai, ***sp. nov.*** Fungal Names FN 572795. Fig. 42E, F.

###### Etymology

the specific epithet *tibetensis* refers to the geographic origin of the type strain, Xizang.

###### Culture characteristics

On YM agar, after 1 month at 25 °C, the streak culture is white, butyrous, smooth and glossy. The margin is fringed (Fig. 42E). In YM broth, after 3 days at 25 °C, cells are ellipsoidal and oval, 1.49–3.13 × 3.22–5.42 µm and single, budding is polar (Fig. 42F), after 7 days at 25 °C, sediment and ring are formed. After 1 month at 25 °C, sediment and ring are present, and pellicle does appear. In Dalmau plate culture on corn meal agar, pseudohyphae are not formed. Asci and ascospores are not observed on V8 and Acetate agar.

###### Physiological and biochemical characteristics

Glucose (weak) fermentation is present. Lactose, galactose, sucrose, maltose and raffinose fermentation are absent. Glucose, galactose (variable), L-sorbose (latent), sucrose (latent), maltose (latent), cellobiose (latent), trehalose (latent), raffinose (variable), melezitose (latent), inulin (latent), soluble starch (variable), D-xylose (latent), D-ribose (latent), L-rhamnose (variable), D-glucosamine (latent), N-acetyl-D-glucosamine (latent), ethanol (latent), glycerol (latent), ribitol (latent, weak), D-mannitol, D-glucitol, methyl-α-D-glucoside (latent), salicin, DL-lactic acid (variable), succinic acid (latent) and citric acid (variable) are assimilated as sole carbon sources. Lactose, melibiose, L-arabinose, D-arabinose, methanol, erythritol, galactitol, inositol, hexadecane and D-glucuronic acid are not assimilated. Ammonium sulfate, potassium nitrate, sodium nitrite (latent), L-lysine, ethylamine and cadaverine are assimilated as sole nitrogen sources. The maximum growth temperature is 35 °C. Growth in the vitamin-free medium is positive. Starch-like substances are not produced. Growth on 50 % (w/w) glucose-yeast extract agar is positive.

Physiologically, *M. xizangensis* differs from the closely related species *M. miensis* (Fig. 19) in its ability to ferment glucose and to assimilate methyl-α-D-glucoside, sodium nitrite and grow in vitamin-free medium (Table S2.19).

###### Typus

**China**, Sejila Mountain, Nyingchi, Tibet, obtained from flower, Aug. 2019, *Q.-M. Wang* (**holotype** CGMCC 2.6957^T^ preserved in a metabolically inactive state, ex-type cultures are preserved in culture collections under the numbers XZY287-2).

###### Other culture examined

**China**, Sejila Mountain, Nyingchi, Tibet, obtained from flower, Aug. 2019, isolated by *Q.M. Wang*, culture XZY314-2. **China**, Sejila Mountain, Nyingchi, Tibet, obtained from flower, Aug. 2019, isolated by *Q.M. Wang*, culture XZY287-3.

***Soucietia violae*** Q.-M. Wang, A.-H. Li, M.-M. Liu, H.-H. Zhu & F.-Y. Bai, ***sp. nov.*** Fungal Names FN 572796. Fig. 42G, H.

###### Etymology

the specific epithet *violae* refers to *Viola*, the plant genus from which the type strain was isolated.

###### Culture characteristics

On YM agar, after 1 month at 25 °C, the streak culture is white, butyrous, rough. The margin is fimbriate (Fig. 42G). In YM broth, after 3 days at 25 °C, cells are globose and oval, 2.43–5.54 × 4.17–7.50 µm and single, budding is polar (Fig. 42H), after 7 days at 25 °C, a sediment and ring are formed. After 1 month at 25 °C, sediment and ring are present, but pellicle does not appear. In Dalmau plate culture on corn meal agar, hyphae are formed. Asci and ascospores are not observed on V8 and Acetate agar.

###### Physiological and biochemical characteristics

Glucose and galactose (weak) fermentation are present. Lactose, sucrose, maltose and raffinose fermentation are absent. Glucose, galactose, L-sorbose, sucrose (latent), maltose, cellobiose (latent), trehalose, raffinose (latent, weak), melezitose (latent), inulin, soluble starch, D-xylose (latent), D-ribose (latent, weak), D-glucosamine, N-acetyl-D-glucosamine, ethanol (latent), glycerol, erythritol (latent, weak), ribitol (latent), D-mannitol, D-glucitol, methyl-α-D-glucoside(latent), salicin (latent) and citric acid (latent) are assimilated as sole carbon sources. Lactose, melibiose, L-arabinose, D-arabinose, L-rhamnose, methanol, galactitol, DL-lactic acid, succinic acid, inositol, hexadecane and D-glucuronic acid are not assimilated. Ammonium sulfate, potassium nitrate, sodium nitrite, L-lysine, ethylamine and cadaverine are assimilated as sole nitrogen sources. The maximum growth temperature is 30 °C. Growth in the vitamin-free medium is positive. Starch-like substances are not produced. Growth on 50 % (w/w) glucose-yeast extract agar is positive.

Physiologically, *S. violae* differs from the closely related species *S. linzhiensis* and *S. sequanensis* (Fig. 18) in its inability to assimilate L-arabinose and succinic acid, and the ability to assimilate sucrose, melezitose, inulin, potassium nitrate and sodium nitrite (Table S2.20).

###### Typus

**China**, Basom lake, Linzhi, Tibet, obtained from *Viola tricolor*, Aug. 2019, *Q.-M. Wang* (**holotype** CGMCC 2.6990^T^ preserved in a metabolically inactive state, ex-type cultures are preserved in culture collections under the numbers XZY238F3).

###### Other culture examined

**China**, Basom lake, Linzhi, Tibet, obtained from *Viola tricolor*, Aug. 2019, isolated by *Q.M. Wang*, culture XZY238F4.

New taxon in the *Saccharomycopsidaceae* (*Ascoideales*, *Saccharomycetes, Saccharomycotina*) *Saccharomycopsis parafodiens* Q.-M. Wang, A.-H. Li, M.-M. Liu, H.-H. Zhu & F.-Y. Bai, *sp. nov.* Fungal Names FN 572797. Fig. 43A, B.

**Fig. 43.**
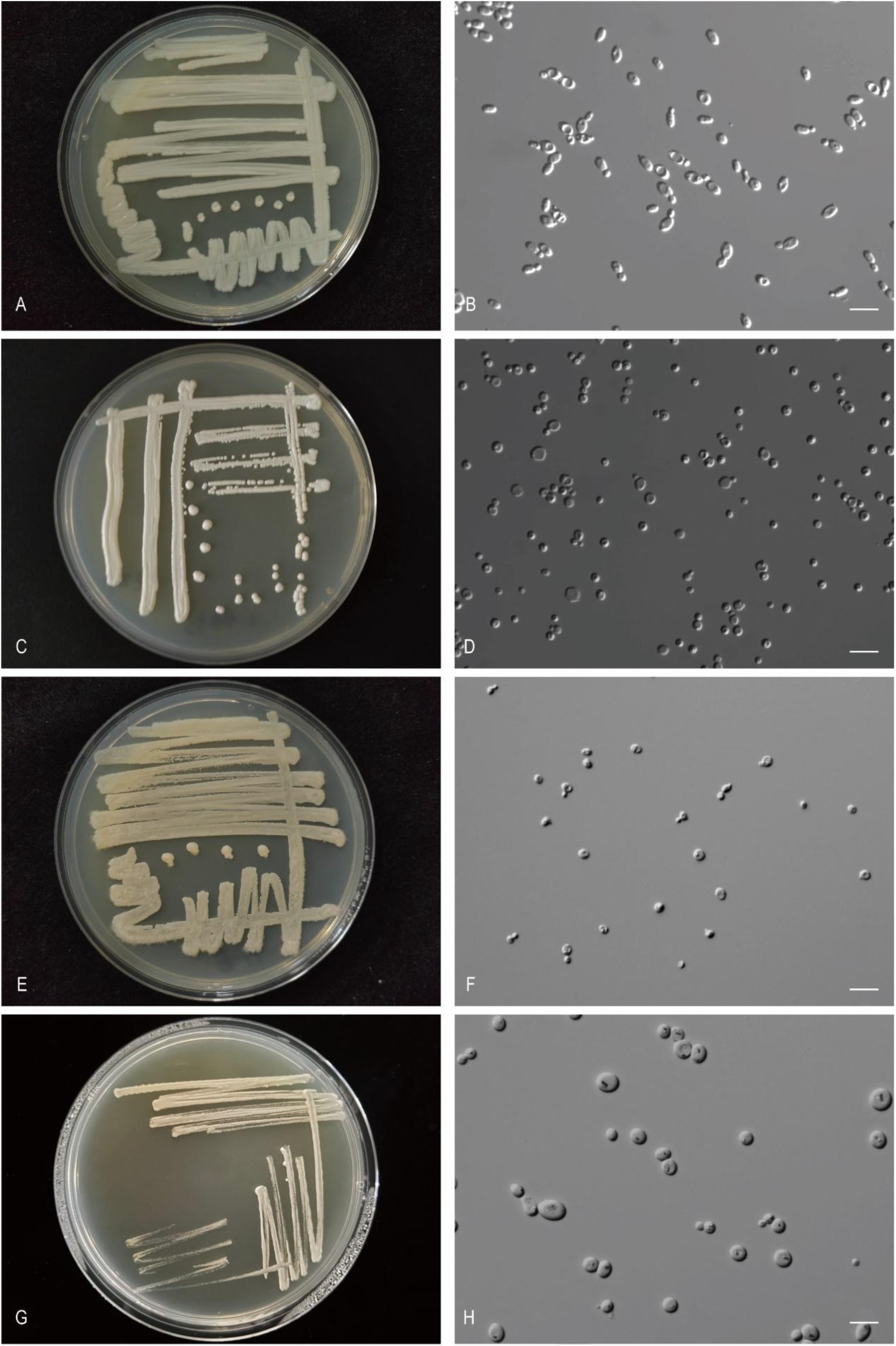
The streak culture grown in YM agar and vegetative cells grown in YM broth for 7 d at 25 °C. A, B. *Sac. parafodiens* CGMCC 2.2749^T^. C, D. *Pet. putridasilvae* CGMCC 2.4335^T^. E, F. *Pet. soli* CGMCC 2.6942^T^. G, H. *Yur. pini* CGMCC 2.7035^T^. Scale bars: B, D, F, H = 10 μm.

###### Etymology

the specific epithet *parafodiens* refers to a similar colony morphology to that of *Saccharomycopsis fodiens*.

###### Culture characteristics

On YM agar, after 1 month at 25 °C, the streak culture is white, butyrous, smooth and glossy. The margin is entire (Fig. 43A). In YM broth, after 3 days at 25 °C, cells are ellipsoidal and cylindrical, 2.36–3.36 × 3.71–5.86 µm and single, budding is polar (Fig. 43B), after 7 days at 25 °C, a sediment is formed. After 1 month at 25 °C, sediment is present, and the ring and pellicle do not appear. In Dalmau plate culture on corn meal agar, hyphae are formed. Asci and ascospores are not observed on V8 and Acetate agar.

###### Physiological and biochemical characteristics

Glucose, lactose, galactose, sucrose, maltose and raffinose fermentation are absent. Maltose, melezitose (latent, weak), L-rhamnose, N-acetyl-D-glucosamine, glycerol, galactitol and methyl-α-D-glucoside are assimilated as sole carbon sources. Glucose, galactose, L-sorbose, sucrose, cellobiose, trehalose, lactose, melibiose, raffinose, inulin, soluble starch, D-xylose, L-arabinose, D-arabinose, D-ribose, D-glucosamine, methanol, ethanol, erythritol, ribitol, D-mannitol, D-glucitol, salicin, DL-lactic acid, succinic acid, citric acid, inositol, hexadecane and D-glucuronic acid are not assimilated. Ammonium sulfate, potassium nitrate, sodium nitrite, L-lysine (weak), ethylamine (weak) and cadaverine are assimilated as sole nitrogen sources. The maximum growth temperature is 32 °C. Growth in the vitamin-free medium is positive. Starch-like substances are not produced. Growth on 50 % (w/w) glucose-yeast extract agar is negative.

Physiologically, *S. parafodiens* differs from the closely related species *S. fodiens* (Fig. 20) in its inability to assimilate glucose, L-sorbose, cellobiose, trehalose, soluble starch, D-xylose, L-arabinose, D-arabinose, D-ribose, ethanol, erythritol, ribitol, D-mannitol, D-glucitol, salicin, DL-lactic acid, succinic acid, citric acid, inositol, D-glucuronic acid and grow with 50% glucose, and the ability to assimilate maltose, melezitose, L-rhamnose, N-acetyl-D-glucosamine, galactitol, methyl-α-D-glucoside, potassium nitrate, sodium nitrite, L-lysine, ethylamine, cadaverine and grow in vitamin-free medium (Table S2.21).

###### Typus

**China**, Tibet, obtained from leaf, Apr. 2005, *Q.-M. Wang* (**holotype** CGMCC 2.2749^T^ preserved in a metabolically inactive state, ex-type cultures are preserved in culture collections under the numbers XZ71A2).

New taxa in the *Phaffomycetaceae* (*Phaffomycetales*, *Saccharomycetes, Saccharomycotina*) *Petasospora putridisilvae* Q.-M. Wang, A.-H. Li, M.-M. Liu, H.-H. Zhu & F.-Y. Bai, *sp. nov.* Fungal Names FN 572798. Fig. 43C, D.

###### Etymology

the specific epithet *putridisilvae* refers to the type strain isolated from rotten wood.

###### Culture characteristics

On YM agar, after 1 month at 25 °C, the streak culture is whitish, butyrous, smooth and pale glossy. The margin is entire (Fig. 43C). In YM broth, after 3 days at 25 °C, cells are globose, subglobose and oval, 2.34–3.55 × 2.63–3.87 µm and single, budding is polar (Fig. 43D), after 7 days at 25 °C, a sediment is formed. After 1 month at 25 °C, sediment is present, pellicle and ring do not appear. In Dalmau plate culture on corn meal agar, hyphae are formed. Asci and ascospores are not observed on V8 and Acetate agar.

###### Physiological and biochemical characteristics

Glucose and sucrose fermentation are present. Lactose, galactose, maltose and raffinose fermentation are absent. Glucose, L-sorbose, sucrose, maltose, cellobiose, trehalose, melibiose, raffinose, melezitose, inulin, soluble starch (latent), D-xylose, L-arabinose (weak), D-arabinose, D-ribose (latent), L-rhamnose, N-acetyl-D-glucosamine, ethanol, glycerol, galactitol, D-mannitol, D-glucitol, methyl-α-D-glucoside, salicin, DL-lactic acid, succinic acid and citric acid are assimilated as sole carbon sources. Galactose, lactose, D-glucosamine, methanol, erythritol, ribitol, inositol, hexadecane and D-glucuronic acid are not assimilated. Ammonium sulfate, potassium nitrate, sodium nitrite, L-lysine, ethylamine and cadaverine are assimilated as sole nitrogen sources. The maximum growth temperature is 40 °C. Growth in the vitamin-free medium is positive. Starch-like substances are not produced. Growth on 50 % (w/w) glucose-yeast extract agar is negative.

Physiologically, *P. putridasilvae* differs from the closely related species *P. maritima* (Fig. 21) in its inability to assimilate D-glucuronic acid, and the ability to assimilate L-sorbose, melibiose, inulin, D-ribose, N-acetyl-D-glucosamine, galactitol, potassium nitrate, sodium nitrite, and grow in vitamin-free medium (Table S2.22).

###### Typus

**China**, Fujian, obtained from rotten wood, *Q.-M. Wang* (**holotype** CGMCC 2.4335^T^ preserved in a metabolically inactive state).

***Petasospora soli sp. nov.*** Q.-M. Wang, A.-H. Li, M.-M. Liu, H.-H. Zhu & F.-Y. Bai, ***sp. nov.*** Fungal Names FN 572799. Fig. 43E, F.

###### Etymology

the specific epithet *soli* refers to the type strain isolated from soil.

###### Culture characteristics

On YM agar, after 1 month at 25 °C, the streak culture is yellowish-cream, butyrous, smooth and pale glossy. The margin is entire (Fig. 43E). In YM broth, after 3 days at 25 °C, cells are ellipsoidal and oval, 1.67–3.51 × 2.24–4.55 µm and single, budding is polar (Fig. 43F), after 7 days at 25 °C, a sediment is formed. After 1 month at 25 °C, sediment is present, and the ring and pellicle do not appear. In Dalmau plate culture on corn meal agar, pseudohyphae are not formed. Asci and ascospores are not observed on V8 and Acetate agar.

###### Physiological and biochemical characteristics

Glucose, lactose, galactose, sucrose, maltose and raffinose fermentation are absent. Glucose, L-sorbose (latent), sucrose, maltose (latent), cellobiose (latent), trehalose (latent), melezitose (latent), inulin (latent), D-xylose, L-rhamnose (latent), N-acetyl-D-glucosamine (latent), ethanol, glycerol, D-mannitol (latent), D-glucitol(latent), methyl-α-D-glucoside, salicin, DL-lactic acid (latent) and succinic acid (latent) are assimilated as sole carbon sources. Galactose, lactose, melibiose, raffinose, soluble starch, L-arabinose, D-arabinose, D-ribose, D-glucosamine, methanol, erythritol, ribitol, galactitol, citric acid, inositol, hexadecane and D-glucuronic acid are not assimilated. Ammonium sulfate, potassium nitrate, sodium nitrite (latent), L-lysine (latent), ethylamine and cadaverine (latent) are assimilated as sole nitrogen sources. The maximum growth temperature is 42 °C. Growth in the vitamin-free medium is positive. Starch-like substances are not produced. Growth on 50 % (w/w) glucose-yeast extract agar is negative.

Physiologically, *P. soli* differs from the closely related species *P. easanensis*, *P. xylosilytica*, *P. maesa* and *P. japonica* (Fig. 21) in its ability to assimilate inulin, potassium nitrate and sodium nitrite (Table S2.22).

###### Typus

**China**, National Forest Reserve, Inner Mongolia, obtained from soil, Oct. 2019, *Q.-M. Wang* (**holotype** CGMCC 2.6942^T^ preserved in a metabolically inactive state, ex-type cultures are preserved in culture collections under the numbers Y1-6).

###### Other culture examined

**China**, National Forest Reserve, Inner Mongolia, obtained from soil, Oct. 2019, isolated by *Q.M. Wang*, culture Y1-5. ***Yurkovozyma*** Q.-M. Wang & M.-M. Liu, ***gen. nov.*** Fungal Names FN 572822.

###### Etymology

The genus is named in honor of A. Yurkov for his contribution to yeast research.

###### Type species

*Yurkovozyma pini* Q.-M. Wang & M.-M. Liu.

This genus is proposed for two Chinese strains, namely XZY480-2 and XZY480-10, which is in a separate branch closely related to *Gotozyma*, *Millerago, Komagataea* in the phylogenetic tree (Fig. 22). Member of the *Phaffomycetaceae* (*Phaffomycetales*, *Saccharomycetes*). The genus is mainly circumscribed by rDNA phylogenetic analysis.

Sexual reproduction is not known. Colonies are white to yellowish, butyrous, smooth. Budding is multilateral. Fermentation is present.

###### Species accepted

***Yurkovozyma pini*** Q.-M. Wang & M.-M. Liu, ***sp. nov.*** Fungal Names FN 572800. Fig. 43G, H.

###### Etymology

the specific epithet pini refers to *Pinus*, the plant genus from which the type strain was isolated.

###### Culture characteristics

On YM agar, after 1 month at 25 °C, the streak culture is white to yellowish, butyrous, smooth and pale glossy. The margin is entire (Fig. 43G). In YM broth, after 3 days at 25 °C, cells are globosal, subglobosal and oval, 3.58–8.67 × 3.94–9.82 µm and single, budding is polar (Fig. 43H), after 7 days at 25 °C, a sediment is formed. After 1 month at 25 °C, a sediment is present, pellicle and ring do not appear. In Dalmau plate culture on corn meal agar, pseudohyphae are not formed. Asci and ascospores are not observed on V8 and Acetate agar.

###### Physiological and biochemical characteristics

Glucose, lactose, galactose and maltose (weak) fermentation are present. Sucrose and raffinose fermentation are absent. Glucose, galactose, L-sorbose, sucrose, maltose, cellobiose, trehalose, melezitose, inulin, soluble starch, D-xylose, L-rhamnose, N-acetyl-D-glucosamine, ethanol, glycerol, erythritol (latent), ribitol, D-mannitol, D-glucitol, methyl-α-D-glucoside, salicin and succinic acid are assimilated as sole carbon sources. Lactose, melibiose, raffinose, L-arabinose, D-arabinose, D-ribose, D-glucosamine, methanol, galactitol, DL-lactic acid, citric acid, inositol, hexadecane and D-glucuronic acid are not assimilated. Ammonium sulfate, potassium nitrate (weak), sodium nitrite, L-lysine, ethylamine and cadaverine are assimilated as sole nitrogen sources. The maximum growth temperature is 20 °C. Growth in the vitamin-free medium is positive. Starch-like substances are not produced. Growth on 50 % (w/w) glucose-yeast extract agar is negative.

Physiologically, *Y. pini* differs from the closely related species *K. salicaria* (Fig. 22) in its inability to assimilate DL-lactic acid, citric acid and D-glucuronic acid, and the ability to ferment glucose, lactose, galactose, maltose and to assimilate L-sorbose, sucrose, maltose, trehalose, melezitose, inulin, soluble starch, N-acetyl-D-glucosamine, erythritol, ribitol, methyl-α-D-glucoside, potassium nitrate and grow in vitamin-free medium (Table S2.23).

###### Typus

**China**, Basongcuo, Tibet, obtained from pine, Aug. 2019, *Q.-M. Wang* (**holotype** CGMCC 2.7035^T^ preserved in a metabolically inactive state, ex-type cultures are preserved in culture collections under the numbers XZY480-2).

###### Other culture examined

**China**, Basongcuo, Nyingchi, Tibet, obtained from pine, Oct. 2019, isolated by *Q.M. Wang*, culture XZY480-10.

New taxa in the *Wickerhamomycetaceae* (*Phaffomycetales*, *Saccharomycetes, Saccharomycotina*) *Hansenula foliicola,* Q.-M. Wang, A.-H. Li, M.-M. Liu, H.-H. Zhu & F.-Y. Bai, *sp. nov.* Fungal Names FN 572801. Fig. 44A, B.

**Fig. 44.**
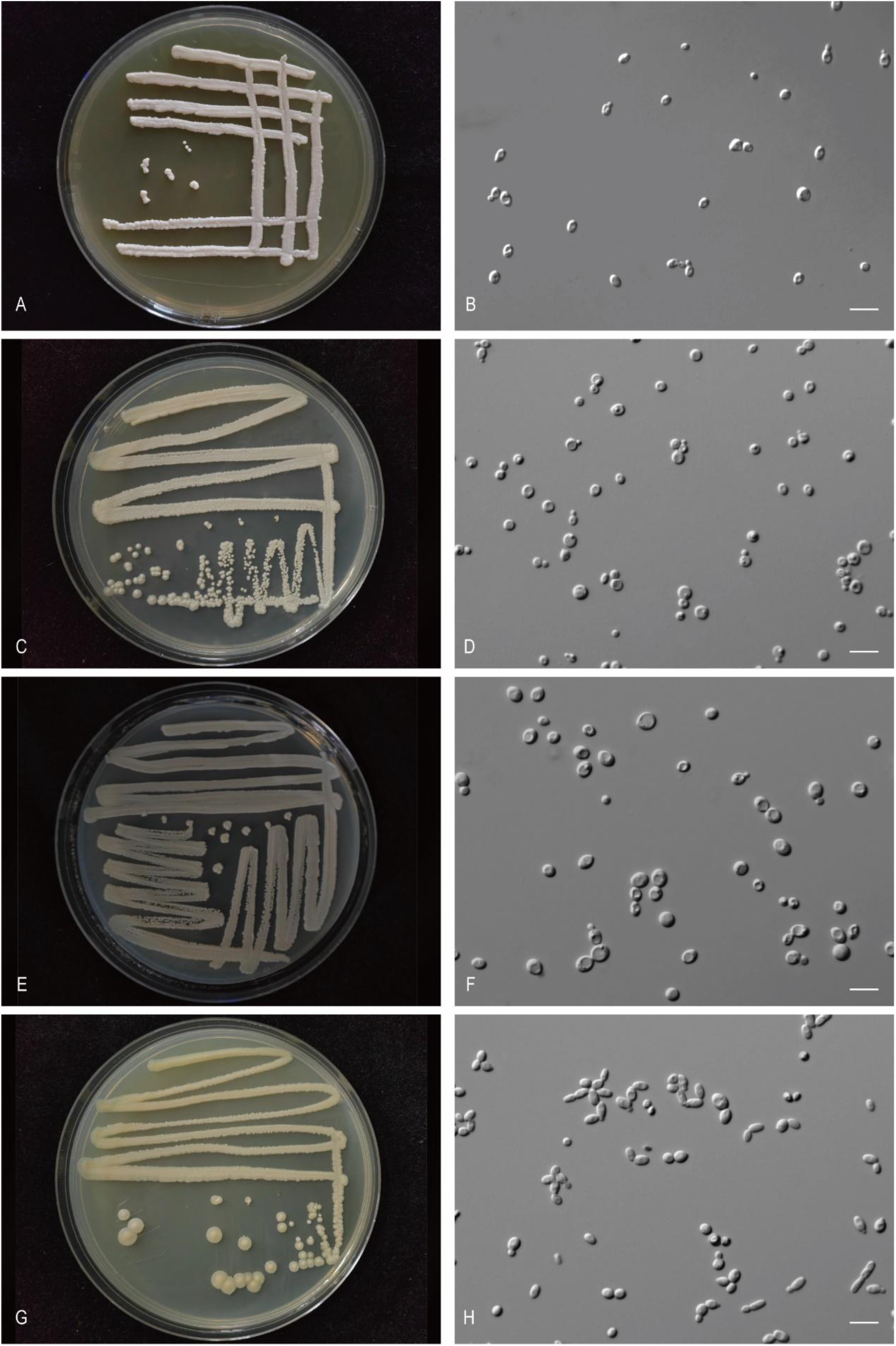
The streak culture grown in YM agar and vegetative cells grown in YM broth for 7 d at 25 °C. A, B. *Han. foliicola* CGMCC 2.3630^T^. C, D. *Han. linzhienesis* CGMCC 2.6968^T^. E, F. *Wal. humusiae* CGMCC 2.6967^T^. G, H. *Xin. betulae* CGMCC 2.6970^T^. Scale bars: B, D, F, H = 10 μm.

###### Etymology

the specific epithet *foliicola* refers to the type strain isolated from a leaf.

###### Culture characteristics

On YM agar, after 1 month at 25 °C, the streak culture is white, butyrous, smooth and pale glossy. The margin is entire (Fig. 44A). In YM broth, after 3 days at 25 °C, cells are oval and ellipsoidal, 2.10–3.66 × 3.21–5.15 µm and single, budding is polar (Fig. 44B), after 7 days at 25 °C, a sediment is formed. After 1 month at 25 °C, sediment is present, the ring and pellicle do not appear. In Dalmau plate culture on corn meal agar, pseudohyphae are not formed. Asci and ascospores are not observed on V8 and Acetate agar.

###### Physiological and biochemical characteristics

Glucose, galactose, sucrose and maltose fermentation are present. Lactose and raffinose fermentation are absent. Glucose, galactose, L-sorbose, sucrose, maltose, cellobiose, trehalose, melibiose, raffinose, melezitose, inulin, soluble starch, D-xylose, L-arabinose, D-ribose, L-rhamnose, D-glucosamine, N-acetyl-D-glucosamine, methanol, ethanol, glycerol, erythritol, ribitol, galactitol, D-mannitol, D-glucitol, methyl-α-D-glucoside, salicin, DL-lactic acid, succinic acid and citric acid are assimilated as sole carbon sources. Lactose, D-arabinose, inositol, hexadecane and D-glucuronic acid are not assimilated. Ammonium sulfate, potassium nitrate, sodium nitrite, L-lysine, ethylamine and cadaverine are assimilated as sole nitrogen sources. The maximum growth temperature is 35 °C. Growth in the vitamin-free medium is positive. Starch-like substances are not produced. Growth on 50 % (w/w) glucose-yeast extract agar is positive (weak).

Physiologically, *H. foliicola* does not differ from the closely related species *H. linzhiensis*, *H. sydowiora*, *H. arboraria*, *H. lynferdii*, *H. myanmarensis*, *H. anomala*, *H. subpelliculosa*, *H. edaphica*, *H. silvicultrix*, *H. ciferrii* and *H. siamensis* (Fig. 23) in its ability to assimilate carbon sources and nitrogen sources (Table S2.24).

###### Typus

**China**, Hainan province, obtained from leaf, Jun. 2007, *Q.-M. Wang* (**holotype** CGMCC 2.3630^T^ preserved in a metabolically inactive state).

***Hansenula linzhienesis*** Q.-M. Wang, A.-H. Li, M.-M. Liu, H.-H. Zhu & F.-Y. Bai, ***sp. nov.*** Fungal Names FN 572802. Fig. 44C, D.

###### Etymology

the specific epithet *linzhienesis* refers to the geographic origin of the type strain, Linzhi, Tibet.

###### Culture characteristics

On YM agar, after 1 month at 25 °C, the streak culture is white, butyrous and smooth. The margin is entire (Fig. 44C). In YM broth, after 3 days at 25 °C, cells are globose, subglobose and oval, 2.39–3.93 × 3.90–5.84 µm and single, budding is polar (Fig. 44D), after 7 days at 25 °C, sediment, ring and pellicle are formed. After 1 month at 25 °C, sediment, ring and pellicle are present. In Dalmau plate culture on corn meal agar, pseudohyphae are not formed. Asci and ascospores are not observed on V8 and Acetate agar.

###### Physiological and biochemical characteristics

Glucose and sucrose (weak) fermentation are present. Lactose, galactose, maltose and raffinose fermentation are absent. Glucose, galactose (latent), L-sorbose (latent), sucrose, maltose, trehalose (latent), raffinose (latent), melezitose (latent), inulin, soluble starch (latent), D-xylose (latent), L-arabinose (latent, weak), L-rhamnose (latent), ethanol (latent), glycerol (latent), erythritol (latent), ribitol (latent), D-mannitol, D-glucitol (latent),methyl-α-D-glucoside (latent), DL-lactic acid (latent),succinic acid (latent) and citric acid (latent) are assimilated as sole carbon sources. Cellobiose, lactose, melibiose, D-arabinose, D-ribose, D-glucosamine, N-acetyl-D-glucosamine, methanol, galactitol, salicin, inositol, hexadecane and D-glucuronic acid are not assimilated. Ammonium sulfate, potassium nitrate, sodium nitrite, L-lysine, ethylamine and cadaverine are assimilated as sole nitrogen sources. The maximum growth temperature is 35 °C. Growth in the vitamin-free medium is positive. Starch-like substances are not produced. Growth on 50 % (w/w) glucose-yeast extract agar is positive.

Physiologically, *H. linzhiensis* does not differ from the closely related species *H. foliicola*, *H. sydowiora*, *H. arboraria*, *H. lynferdii*, *H. myanmarensis*, *H. anomala*, *H. subpelliculosa*, *H. edaphica*, *H. silvicultrix*, *H. ciferrii* and *H. siamensis* (Fig. 23) in its ability to assimilate carbon sources and nitrogen sources (Table S2.24).

###### Typus

**China**, Basongcuo Lake, Nyingchi City, Tibet, obtained from rotten wood, Aug. 2019, *Q.-M. Wang* (**holotype** CGMCC 2.6968^T^ preserved in a metabolically inactive state, ex-type cultures are preserved in culture collections under the numbers XZY836-1).

###### Other culture examined

**China**, Basongcuo, Nyingchi, Tibet, obtained from rotten wood, Aug. 2019, isolated by *Q.M. Wang*, culture XZY836-1B.

***Waltiozyma humusiae*** Q.-M. Wang, A.-H. Li, M.-M. Liu, H.-H. Zhu & F.-Y. Bai, ***sp. nov.*** Fungal Names FN 572803. Fig. 44E, F.

###### Etymology

the specific epithet *humusiae* refers to the type strain isolated from soil.

###### Culture characteristics

On YM agar, after 1 month at 25 °C, the streak culture is white-cream, butyrous, smooth and pale glossy. The margin is entire (Fig. 44E). In YM broth, after 3 days at 25 °C, cells are ellipsoidal and oval, 3.86–5.42 × 4.95–6.98 µm and single, budding is polar (Fig. 44F), after 7 days at 25 °C, a sediment, ring and pellicle are formed. After 1 month at 25 °C, sediment, ring and pellicle are present. In Dalmau plate culture on corn meal agar, pseudohyphae are not formed. Asci and ascospores are not observed on V8 and Acetate agar.

###### Physiological and biochemical characteristics

Glucose and sucrose (variable) fermentation are present. Lactose, galactose, maltose and raffinose fermentation are absent. Glucose, galactose (variable), L-sorbose (variable), sucrose, maltose (variable), cellobiose, trehalose, lactose (variable), raffinose (variable), melezitose, inulin, soluble starch (latent, weak), D-xylose, L-rhamnose (variable), N-acetyl-D-glucosamine (variable), ethanol, glycerol (latent), erythritol (variable), ribitol (variable), D-mannitol, methyl-α-D-glucoside, salicin (latent), DL-lactic acid (latent) and succinic acid (latent) are assimilated as sole carbon sources. Melibiose, L-arabinose, D-arabinose, D-ribose, D-glucosamine, methanol, galactitol, D-glucitol, citric acid, inositol, hexadecane and D-glucuronic acid are not assimilated. Ammonium sulfate, potassium nitrate, sodium nitrite, L-lysine, ethylamine and cadaverine are assimilated as sole nitrogen sources. The maximum growth temperature is 32 °C. Growth in the vitamin-free medium is positive. Starch-like substances are not produced. Growth on 50 % (w/w) glucose-yeast extract agar is negative.

Physiologically, *W. humusiae* differs from the closely related species *W. solani* (Fig. 23) in its ability to assimilate inulin, soluble starch, potassium nitrate, sodium nitrite and grow in vitamin-free medium (Table S2.25).

###### Typus

**China**, Motuo County, Linzhi, Tibet, obtained from soil, Aug. 2019, *Q.-M. Wang* (**holotype** CGMCC 2.6967^T^ preserved in a metabolically inactive state, ex-type cultures are preserved in culture collections under the numbers M14F1).

###### Other culture examined

**China**, Jiangzi County, Shigatze City, Tibet, obtained from soil, Aug. 2019, isolated by *Q.M. Wang*, culture XZ18-4F1; **China**, Jiangzi County, Shigatze City, Tibet, obtained from soil, Aug. 2019, isolated by *Q.M. Wang*, culture XZ18-4F8; **China**, Jiangzi County, Shigatze City, Tibet, obtained from soil, Aug. 2019, isolated by *Q.M. Wang*, culture XZ18-41F3A; **China**, Motuo County, Linzhi, Tibet, obtained from soil, Aug. 2019, isolated by *Q.M. Wang*, culture M14F6; **China**, Motuo County, Linzhi, Tibet, obtained from soil, Aug. 2019, isolated by *Q.M. Wang*, culture M14F3.

***Xingzhongia betulae*** Q.-M. Wang, A.-H. Li, M.-M. Liu, H.-H. Zhu & F.-Y. Bai, ***sp. nov.*** Fungal Names FN 572804. Fig. 44G, H.

###### Etymology

the specific epithet *betulae* refers to *Betula*, the plant genus from which the type strain was isolated.

###### Culture characteristics

On YM agar, after 1 month at 25 °C, the streak culture is whitish-cream, butyrous, smooth and pale glossy. The margin is entire (Fig. 44G). In YM broth, after 3 days at 25 °C, cells are globose, ellipsoidal and oval, 2.18–3.89 × 3.88–6.39 µm and single, budding is polar (Fig.44H), after 7 days at 25 °C, a sediment and ring are formed. After 1 month at 25 °C, sediment and ring and are present, pellicle does not appear. In Dalmau plate culture on corn meal agar, pseudohyphae are formed. Asci and ascospores are not observed on V8 and Acetate agar.

###### Physiological and biochemical characteristics

Glucose fermentation is present. Lactose, galactose, sucrose, maltose and raffinose fermentation are absent. Glucose, L-sorbose (latent), sucrose, maltose, cellobiose, trehalose (latent), melezitose (latent), inulin (latent), soluble starch (latent, weak), D-xylose (latent), L-rhamnose (latent), ethanol, glycerol, D-mannitol, D-glucitol, methyl-α-D-glucoside, salicin, succinic acid and citric acid (latent) are assimilated as sole carbon sources. Galactose, lactose, melibiose, raffinose, L-arabinose, D-arabinose, D-ribose, D-glucosamine, N-acetyl-D-glucosamine, methanol, erythritol, ribitol, galactitol, DL-lactic acid, inositol, hexadecane and D-glucuronic acid are not assimilated. Ammonium sulfate, potassium nitrate, sodium nitrite, L-lysine, ethylamine and cadaverine are assimilated as sole nitrogen sources. The maximum growth temperature is 35 °C. Growth in the vitamin-free medium is positive. Starch-like substances are not produced. Growth on 50 % (w/w) glucose-yeast extract agar is negative.

Physiologically, *X. betulae* differs from the closely related species *X. pteridiae* and *X. hampshirensis* (Fig. 23) in its inability to assimilate DL-lactic acid (Table S2.26).

###### Typus

**China**, New basom lake, Linzhi, Tibet, obtained from bark of *Betula platyphylla*, Aug. 2019, *Q.- M. Wang* (**holotype** CGMCC 2.6970^T^ preserved in a metabolically inactive state, ex-type cultures are preserved in culture collections under the numbers XZY448-3).

###### Other culture examined

**China**, New basom lake, Linzhi, Tibet, obtained from bark of *Betula platyphylla*, Aug. 2019, isolated by *Q.M. Wang*, culture XZY448-3B.

***Xingzhongia insecticola*** Q.-M. Wang, A.-H. Li, M.-M. Liu, H.-H. Zhu & F.-Y. Bai, ***sp. nov.*** Fungal Names FN 572805. Fig. 45A-C.

**Fig. 45.**
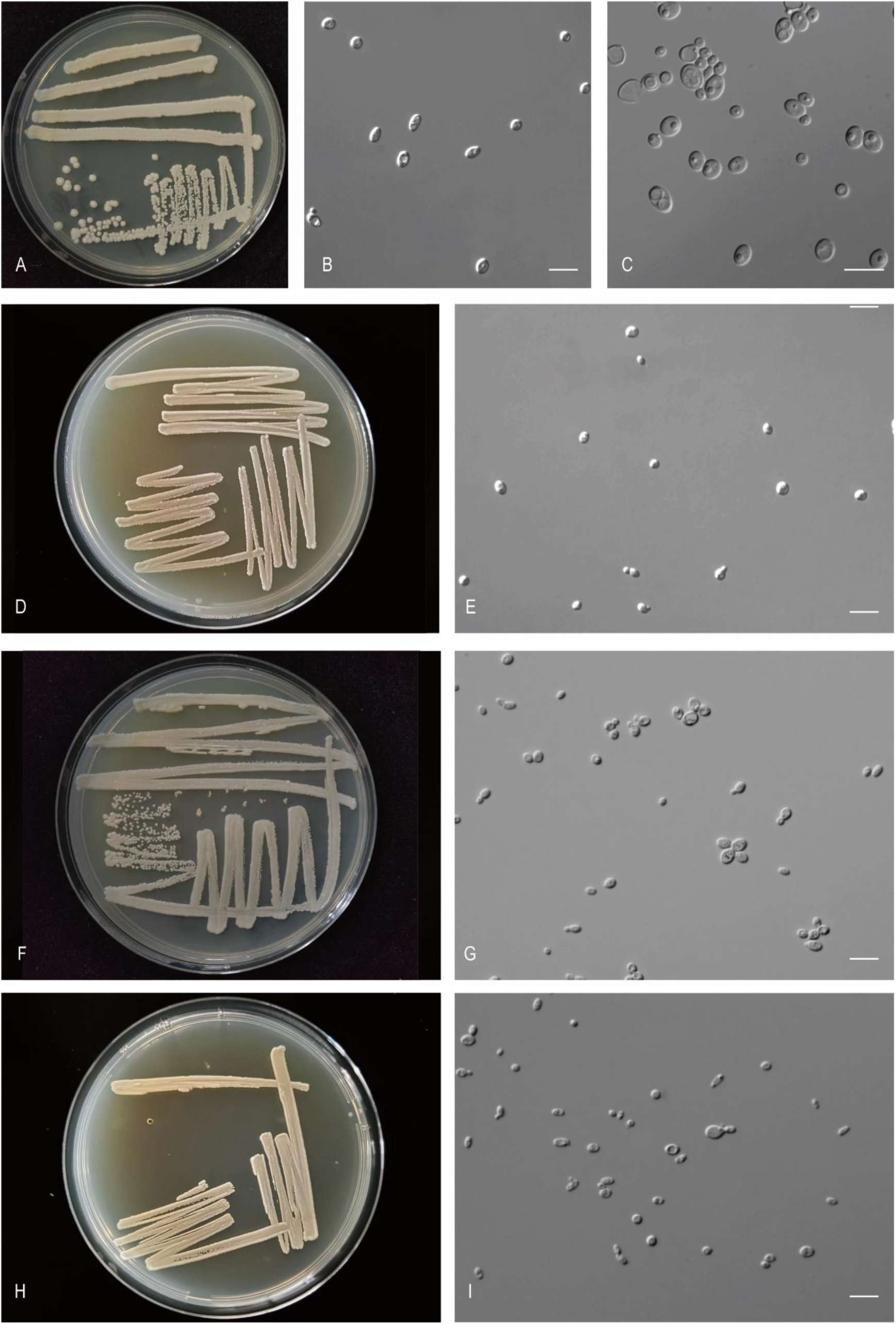
The streak culture grown in YM agar and vegetative cells grown in YM broth for 7 d at 25 °C. A-C. *Xin. insecticola* CGMCC 2.3637^T^. D, E. *Xin. pteridiae* CGMCC 2.3436^T^. F, G. *Wic. caprifoliaceae* CGMCC 2.6973^T^. H, I. *Wic. corticola* CGMCC 2.6972^T^. Scale bars: B, C, E, G, I = 10 μm.

###### Etymology

the specific epithet *insecticola* refers to the type strain isolated from an insect.

###### Culture characteristics

On YM agar, after 1 month at 25 °C, the streak culture is white, butyrous, smooth and glossy. The margin is entire (Fig. 45A). In YM broth, after 3 days at 25 °C, cells are oval, ellipsoidal and fusiform, 2.46–4.45 × 4.21–6.78 µm and single, budding is polar (Fig. 45B), after 7 days at 25 °C, a sediment is formed. After 1 month at 25 °C, sediment is present, the ring and pellicle do not appear. In Dalmau plate culture on corn meal agar, pseudohyphae are not formed. On Acetate agar, the ascus is oval and contains two ascospores (Fig. 45C). The ascospores are spherical or subspherical, 2.17–3.27 × 2.45–3.42 µm.

###### Physiological and biochemical characteristics

Glucose, sucrose and maltose (weak) fermentation are present. Lactose, galactose and raffinose fermentation are absent. Glucose, galactose (latent, weak), L-sorbose, sucrose, maltose, cellobiose, trehalose, melibiose (variable), raffinose (weak), melezitose, inulin, soluble starch (latent, weak), D-xylose, L-arabinose (variable), D-ribose (variable), L-rhamnose (variable), D-glucosamine (variable), N-acetyl-D-glucosamine (variable), methanol (variable), ethanol, glycerol (variable), ribitol, galactitol (variable), D-mannitol, D-glucitol, methyl-α-D-glucoside (latent, weak), salicin, DL-lactic acid and succinic acid are assimilated as sole carbon sources. Lactose, D-arabinose, erythritol, citric acid, inositol, hexadecane and D-glucuronic acid are not assimilated. Potassium nitrate (variable), L-lysine, ethylamine and cadaverine are assimilated as sole nitrogen sources. Ammonium sulfate and sodium nitrite are not assimilated. The maximum growth temperature is 37 °C. Growth in the vitamin-free medium is positive. Starch-like substances are not produced. Growth on 50 % (w/w) glucose-yeast extract agar is negative.

Physiologically, *X. insecticola* differs from the closely related species *X. scolytoplatypi* and *X. strasburgensis* (Fig. 23) in its ability to ferment maltose and to assimilate L-sorbose, inulin, and grow in vitamin-free medium (Table S2.26).

###### Typus

**China**, Beijing, obtained from inside an insect, *F.-Y. Bai* (**holotype** CGMCC 2.3637^T^ preserved in a metabolically inactive state, ex-type cultures are preserved in culture collections under the numbers CBS 18410).

###### Other culture examined

**China**, Beijing, obtained from *Sympetrum kunckeli*, Feb. 2007, isolated by *F.-Y. Bai*, culture CGMCC 2.3710(= HQ3.1).

***Xingzhongia pteridiae*** Q.-M. Wang, A.-H. Li, M.-M. Liu, H.-H. Zhu & F.-Y. Bai, ***sp. nov.*** Fungal Names FN 572806. Fig. 45D, E.

###### Etymology

the specific epithet *pteridiae* refers to *Pteridium*, the plant genus from which the strain was isolated.

###### Culture characteristics

On YM agar, after 1 month at 25 °C, the streak culture is white, butyrous, smooth. The margin is entire (Fig. 45D). In YM broth, after 3 days at 25 °C, cells are subglobose, ellipsoidal and oval, 1.93–3.42 × 2.81–4.78 µm and single, budding is polar (Fig. 45E), after 7 days at 25 °C, sediment, ring and pellicle are formed. After 1 month at 25 °C, sediment, ring and pellicle are present. In Dalmau plate culture on corn meal agar, pseudohyphae are not formed. Asci and ascospores are not observed on V8 and Acetate agar.

###### Physiological and biochemical characteristics

Glucose (variable) fermentation is present. Lactose, galactose, sucrose, maltose and raffinose fermentation are absent. Glucose, galactose (variable), L-sorbose, sucrose, maltose, cellobiose (variable), trehalose, lactose (variable), melibiose, raffinose, melezitose, inulin, soluble starch, D-xylose, L-arabinose (variable), D-ribose (latent, weak), L-rhamnose (variable), ethanol, glycerol, ribitol (variable), galactitol (variable), D-mannitol, D-glucitol, methyl-α-D-glucoside, salicin, DL-lactic acid, succinic acid and citric acid (variable) are assimilated as sole carbon sources. D-arabinose, D-glucosamine, N-acetyl-D-glucosamine, methanol, erythritol, inositol, hexadecane and D-glucuronic acid are not assimilated. Ammonium sulfate (variable), potassium nitrate (variable), sodium nitrite, L-lysine, ethylamine and cadaverine are assimilated as sole nitrogen sources. The maximum growth temperature is 40 °C. Growth in the vitamin-free medium is positive. Starch-like substances are not produced. Growth on 50 % (w/w) glucose-yeast extract agar is negative.

Physiologically, *X. pteridiae* differs from the closely related species *X. betulae* and *X. hampshirensis* (Fig. 23) in its ability to assimilate melibiose, raffinose and D-ribose (Table S2.26).

###### Typus

**China**, Hunan province, obtained from leaf of *Pteridium* sp., Jan. 2007, *Q.-M. Wang* (**holotype** CGMCC 2.3436^T^ preserved in a metabolically inactive state, ex-type cultures are preserved in culture collections under the numbers JUE3.4).

###### Other culture examined

**China**, Hunan Province, obtained from *Ganoderma applanatum*, isolated by *Q.M. Wang*, culture CGMCC 2.3429. **China**, Hunan Province, obtained from leaf *of Pteridium* sp., Jan. 2007, isolated by *Q.M. Wang*, culture JUE3.4B.

***Wickerhamomyces caprifoliacearum*** Q.-M. Wang, A.-H. Li, M.-M. Liu, H.-H. Zhu & F.-Y. Bai, ***sp. nov.*** Fungal Names FN 572807. Fig. 45F, G.

###### Etymology

the specific epithet *caprifoliacearum* refers to *Caprifoliaceae*, the plant genus from which the type strain was isolated.

###### Culture characteristics

On YM agar, after 1 month at 25 °C, the streak culture is white, cream, smooth and glossy. The margin is entire (Fig. 45F). In YM broth, after 3 days at 25 °C, cells are ellipsoidal and oval, 1.98–4.40 × 3.20–5.72 µm and single, budding is polar (Fig. 45G), after 7 days at 25 °C, a sediment and ring are formed. After 1 month at 25 °C, sediment and ring are present, but pellicle does not appear. In Dalmau plate culture on corn meal agar, pseudohyphae are not formed. Asci and ascospores are not observed on V8 and Acetate agar.

###### Physiological and biochemical characteristics

Glucose fermentation is present. Lactose, galactose, sucrose, maltose and raffinose fermentation are absent. Glucose, galactose (latent, weak), L-sorbose, sucrose, maltose, cellobiose, trehalose, melezitose, inulin, soluble starch (latent, weak), D-xylose, L-rhamnose, N-acetyl-D-glucosamine (variable), ethanol, glycerol, D-mannitol, D-glucitol, methyl-α-D-glucoside, salicin, DL-lactic acid (latent, weak) and succinic acid are assimilated as sole carbon sources. Lactose, melibiose, raffinose, L-arabinose, D-arabinose, D-ribose, D-glucosamine, methanol, erythritol, ribitol, galactitol, citric acid, inositol, hexadecane and D-glucuronic acid are not assimilated. Ammonium sulfate, potassium nitrate, sodium nitrite, L-lysine (latent), ethylamine and cadaverine are assimilated as sole nitrogen sources. The maximum growth temperature is 32 °C. Growth in the vitamin-free medium is positive. Starch-like substances are not produced. Growth on 50 % (w/w) glucose-yeast extract agar is negative.

Physiologically, *W. caprifoliacearum* does not differ from the closely related species *W. terricola* and *W. canadensis* (Fig. 23) in its inability to assimilate carbon sources and nitrogen sources (Table S2.27).

###### Typus

**China**, Katinggou, Nyingchi City, Tibet, obtained from the leaf of *Caprifoliaceae*, Aug. 2019, *Q.-M. Wang* (**holotype** CGMCC 2.6973^T^ preserved in a metabolically inactive state, ex-type cultures are preserved in culture collections under the numbers XZY167-6).

###### Other culture examined

**China**, Katinggou, Nyingchi City, Tibet, obtained from leaf of *Caprifoliaceae*, Aug. 2019, isolated by *Q.M. Wang*, culture XZY167-2.

***Wickerhamomyces corticola*** Q.-M. Wang, A.-H. Li, M.-M. Liu, H.-H. Zhu & F.-Y. Bai, ***sp. nov.*** Fungal Names FN 572808. Fig. 45H, I.

###### Etymology

the specific epithet *corticola* refers to the type strain isolated from bark.

###### Culture characteristics

On YM agar, after 1 month at 25 °C, the streak culture is whitish-cream, butyrous, smooth and pale glistening. The margin is entire (Fig. 45H). In YM broth, after 3 days at 25 °C, cells are ellipsoidal and oval, 1.91–4.29 × 2.56–5.88 µm and single, budding is polar (Fig. 45I), after 7 days at 25 °C, a sediment and ring are formed. After 1 month at 25 °C, sediment and ring are present, pellicle does not appear. In Dalmau plate culture on corn meal agar, pseudohyphae are not formed. Asci and ascospores are not observed on V8 and Acetate agar.

###### Physiological and biochemical characteristics

Glucose (weak) fermentation is present. Lactose, galactose, sucrose, maltose and raffinose fermentation are absent. Glucose, galactose (variable), L-sorbose, sucrose, maltose, cellobiose, trehalose, melibiose (variable), raffinose (variable), melezitose, inulin, soluble starch (weak), D-xylose, D-ribose (variable), L-rhamnose (latent), N-acetyl-D-glucosamine (variable), ethanol, glycerol, erythritol (variable), ribitol (variable), galactitol (variable), D-mannitol, D-glucitol, methyl-α-D-glucoside, salicin, DL-lactic acid (variable), succinic acid, citric acid (variable), inositol (variable), hexadecane (variable) and D-glucuronic acid (variable) are assimilated as sole carbon sources. Lactose, L-arabinose, D-arabinose, D-glucosamine and methanol are not assimilated. Ammonium sulfate, potassium nitrate, sodium nitrite, L-lysine, ethylamine and cadaverine are assimilated as sole nitrogen sources. The maximum growth temperature is 40 °C. Growth in the vitamin-free medium is positive. Starch-like substances are not produced. Growth on 50 % (w/w) glucose-yeast extract agar is negative.

Physiologically, *W. corticola* differs from the closely related species *W. psychrolipolyticus* (Fig. 23) in its inability to assimilate D-arabinose, and the ability to assimilate L-sorbose, inulin and succinic acid (Table S2.27).

###### Typus

**China**, Greater Khinganling National Forest Reserve, Heilongjiang Province, obtained from bark, Sep. 2017, *Q.-M. Wang* (**holotype** CGMCC 2.6972^T^ preserved in a metabolically inactive state, ex-type cultures are preserved in culture collections under the numbers HSP215-3).

###### Other culture examined

**China**, Greater Khinganling National Forest Reserve, Heilongjiang Province, obtained from bark, Sep. 2017, isolated by *Q.M. Wang*, culture HSP255-2; **China**, Greater Khinganling National Forest Reserve, Heilongjiang Province, obtained from bark, Sep. 2017, isolated by *Q.M. Wang*, culture HSP215-1.

***Wickerhamomyces terricola*** Q.-M. Wang, A.-H. Li, M.-M. Liu, H.-H. Zhu & F.-Y. Bai, ***sp. nov.*** Fungal Names FN 572809. Fig. 46A, B.

**Fig. 46.**
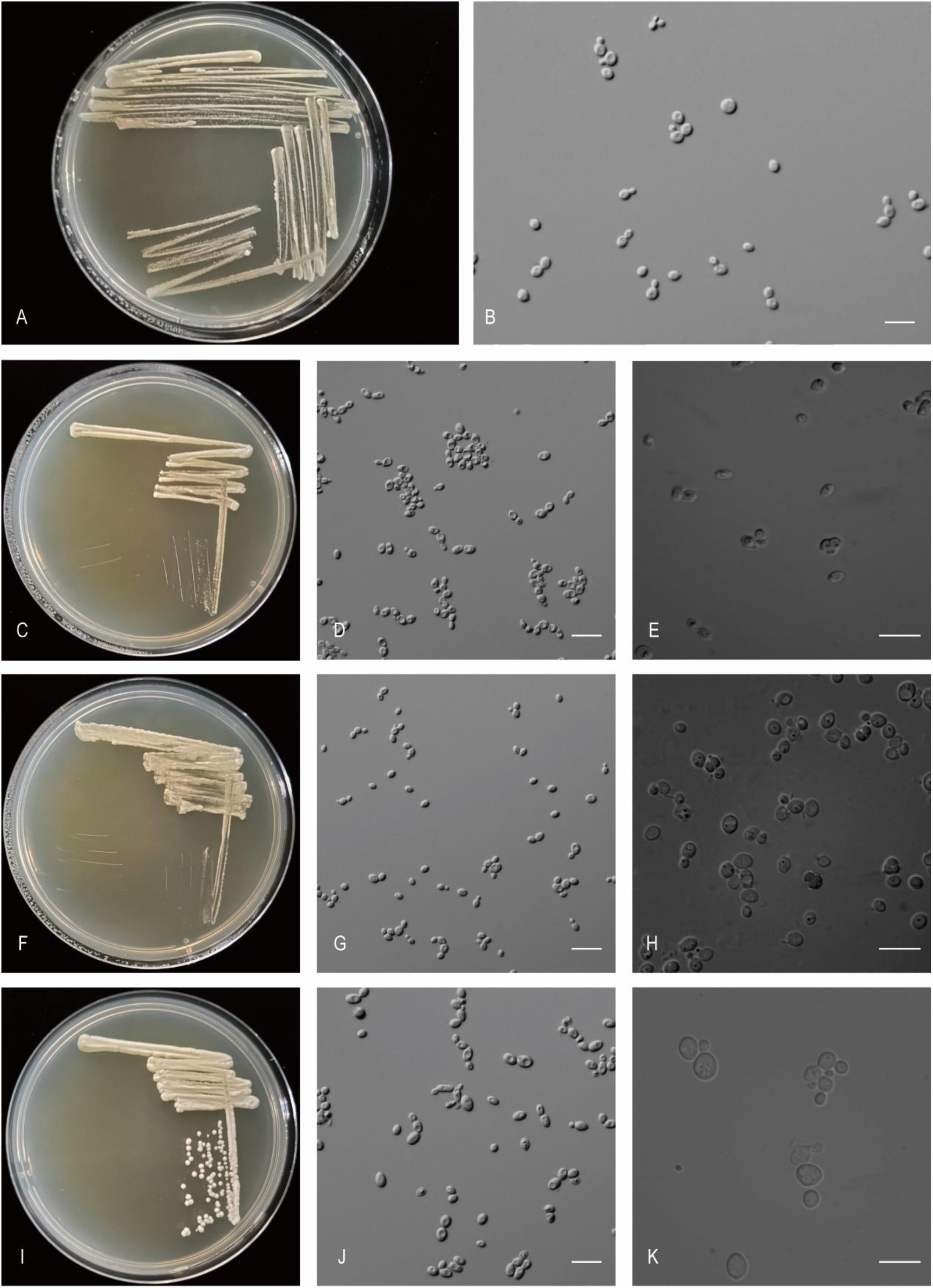
The streak culture grown in YM agar and vegetative cells grown in YM broth for 7 d at 25 °C. A, B. *Wic. terricola* CGMCC 2.8739^T^. C-E. *Gab. putridisilvae* CGMCC 2.4518^T^. F-H. *Hui. daliensis* CGMCC 2.4475^T^. I-K. *Jam. terricola* CGMCC 2.3906^T^. Scale bars: B, D, E, G, H, J, K = 10 μm.

###### Etymology

the specific epithet *terricola* refers to the type strain isolated from soil.

###### Culture characteristics

On YM agar, after 1 month at 25 °C, the streak culture is white, butyrous and smooth. The margin is fringed (Fig. 46A). In YM broth, after 3 days at 25 °C, cells are ellipsoidal and oval, 1.76–3.40 × 2.59–4.11 µm and single, budding is polar (Fig. 46B), after 7 days at 25 °C, a sediment and ring are formed. After 1 month at 25 °C, sediment, pellicle and ring are present. In Dalmau plate culture on corn meal agar, pseudohyphae are not formed. Asci and ascospores are not observed on V8 and Acetate agar.

###### Physiological and biochemical characteristics

Glucose fermentation is present. Lactose, galactose, sucrose, maltose and raffinose fermentation are absent. Glucose, galactose (latent), L-sorbose, sucrose, maltose, trehalose (latent), melezitose, inulin, soluble starch (weak), D-xylose, L-rhamnose, N-acetyl-D-glucosamine (latent, weak), ethanol, glycerol, ribitol (latent), D-mannitol, D-glucitol, methyl-α-D-glucoside, salicin, DL-lactic acid (latent, weak) and succinic acid are assimilated as sole carbon sources. Cellobiose, lactose, melibiose, raffinose, L-arabinose, D-arabinose, D-ribose, D-glucosamine, methanol, erythritol, galactitol, citric acid, inositol, hexadecane and D-glucuronic acid are not assimilated. Ammonium sulfate, potassium nitrate, sodium nitrite, L-lysine, ethylamine and cadaverine are assimilated as sole nitrogen sources. The maximum growth temperature is 30 °C. Growth in the vitamin-free medium is positive. Starch-like substances are not produced. Growth on 50 % (w/w) glucose-yeast extract agar is negative.

Physiologically, *W. terricola* differs from the closely related species *W. caprifoliacearum* and *W. canadensis* in its inability to assimilate cellobiose (Table S2.27).

###### Typus

**China**, Naidong District, Shannan City, Tibet, obtained from soil, Aug. 2019, *Q.-M. Wang* (**holotype** CGMCC 2.8739^T^ preserved in a metabolically inactive state, ex-type cultures are preserved in culture collections under the numbers XZY78-1).

New taxa in the Saccharomycetaceae (Saccharomycetales, Saccharomycetes, Saccharomycotina) Gaboromyces Q.-M. Wang & M.-M. Liu, *gen. nov.* Fungal Names FN 572823.

###### Etymology

The genus is named in honor of G. Péter for his contribution to yeast taxonomy.

###### Type species

*Gaboromyces psychrophila* Kabisch, Höning, Böhnlein, P. Pichner, Gareis & Wenning ex Q.-M. Wang & M.-M. Liu

This genus is proposed for *Kazachstania psychrophila* and a new species represented by two Chinese strains, namely CGMCC 2.4518 and BW12-4, which form a clade closely related to *Jamesozyma* and *Kazachstania kunashirensis* in the phylogenetic tree (Fig. 24). Member of the *Saccharomycetaceae* (*Saccharomycetales*, *Saccharomycetes*). The genus is mainly circumscribed by rDNA phylogenetic analysis.

Asci contains one or two globose to ovoid ascospores. Colonies are white to tan, butyrous, smooth. Budding is multilateral. Fermentation is present.

###### Species accepted

***Gaboromyces psychrophila*** Kabisch, Höning, Böhnlein, P. Pichner, Gareis & Wenning ex Q.-M. Wang & M.-M. Liu, ***sp. nov.*** Fungal Names FN 572824.

For a detailed description see Kabisch *et al*., Antonie van Leeuwenhoek 104: 930. 2013. *Holotype*: CBS 12689, preserved in a metabolically inactive state at Westerdijk Institute. *Synonym*: *Kazachstania psychrophila* Kabisch, Höning, Böhnlein, P. Pichner, Gareis & Wenning, Antonie van Leeuwenhoek 104: 930. 2013, nom. inval., Art. 40.7 (Melbourne).

***Gaboromyces putridisilvae*** Q.-M. Wang & M.-M. Liu, ***sp. nov.*** Fungal Names FN 572810. Fig. 46C-E.

###### Etymology

the specific epithet *putridisilvae* refers to the type strain isolated from rotten wood.

###### Culture characteristics

On YM agar, after 1 month at 25 °C, the streak culture is white, butyrous and smooth. The margin is entire (Fig. 46C). In YM broth, after 3 days at 25 °C, cells are ellipsoidal and fusiform, 1.81–3.43 × 2.16–4.07 µm and single, budding is polar (Fig. 46D), after 7 days at 25 °C, a sediment, pellicle and ring are formed. After 1 month at 25 °C, sediment, pellicle and ring are present. In Dalmau plate culture on corn meal agar, pseudohyphae are not formed. On V8 agar, the ascus is fusiform and contains two ascospores (Fig. 46E). The ascospores are spherical or subspherical, 3.06–3.88 × 3.02–3.93 µm.

###### Physiological and biochemical characteristics

Glucose, galactose, sucrose and maltose fermentation are present. Lactose and raffinose fermentation are absent. Glucose, galactose, sucrose, trehalose, lactose (weak), melibiose (weak), raffinose, inulin, D-ribose, ethanol (latent, weak), glycerol, DL-lactic acid and succinic acid are assimilated as sole carbon sources. L-sorbose, maltose, cellobiose, melezitose, soluble starch, D-xylose, L-arabinose, D-arabinose, L-rhamnose, D-glucosamine, N-acetyl-D-glucosamine, methanol, erythritol, ribitol, galactitol, D-mannitol, D-glucitol, methyl-α-D-glucoside, salicin, citric acid, inositol, hexadecane and D-glucuronic acid are not assimilated. Ammonium sulfate, potassium nitrate and sodium nitrite are assimilated as sole nitrogen sources. L-lysine, ethylamine and cadaverine are not assimilated. The maximum growth temperature is 30 °C. Growth in the vitamin-free medium is negative. Starch-like substances are not produced. Growth on 50 % (w/w) glucose-yeast extract agar is positive.

Physiologically, *G. putridisilvae* differs from the closely related species *G. psychrophila* (Fig. 24) in its inability to ferment raffinose and assimilate D-glucuronic acid, and the ability to assimilate lactose, melibiose, inulin, D-ribose, ethanol, potassium nitrate, sodium nitrite, and grow in vitamin-free medium and grow with 50% glucose (Table S2.28).

###### Typus

**China**, obtained from rotten wood, Mar. 2012, *Q.-M. Wang* (**holotype** CGMCC 2.4518^T^ preserved in a metabolically inactive state).

###### Other culture examined

**China**, Hainan province, obtained from rotten wood, Mar. 2012, isolated by *Q.M. Wang*, culture BW12-4.

***Huiozyma daliensis*** Q.-M. Wang, A.-H. Li, M.-M. Liu, H.-H. Zhu & F.-Y. Bai, ***sp. nov.*** Fungal Names FN 572811. Fig. 46F-H.

###### Etymology

the specific epithet *daliensis* refers to the geographic origin of the type strain, Dali, Yunnan province.

###### Culture characteristics

On YM agar, after 1 month at 25 °C, the streak culture is white, butyrous, smooth and glossy. The margin is entire (Fig. 46F). In YM broth, after 3 days at 25 °C, cells are ellipsoidal and fusiform, 1.55–2.85 × 2.25–5.00 µm and single, budding is polar (Fig. 46G), after 7 days at 25 °C, a sediment and ring are formed. After 1 month at 25 °C, the sediment and ring are present, but a pellicle does not appear. In Dalmau plate culture on corn meal agar, pseudohyphae are not formed. On V8 agar, the ascus is fusiform and contains two ascospores (Fig. 46H). The ascospores are ellipsoidal, 2.36–4.01 × 2.98–4.45 µm.

###### Physiological and biochemical characteristics

Glucose, galactose and sucrose fermentation are present. Lactose, maltose and raffinose fermentation are absent. Glucose, galactose, sucrose, maltose, trehalose, lactose (latent, weak), melibiose (variable), raffinose, inulin, soluble starch (variable), L-arabinose (variable), D-arabinose (variable), D-ribose (latent), L-rhamnose (variable), N-acetyl-D-glucosamine (variable), methanol (variable), ethanol, glycerol (latent), erythritol (variable), galactitol (variable), D-mannitol, D-glucitol (variable), methyl-α-D-glucoside (variable), DL-lactic acid (latent), succinic acid (latent), citric acid (variable), inositol (variable) and hexadecane(variable) are assimilated as sole carbon sources. L-sorbose, cellobiose, melezitose, D-xylose, D-glucosamine, ribitol, salicin and D-glucuronic acid are not assimilated. Ammonium sulfate, potassium nitrate, sodium nitrite, L-lysine, ethylamine and cadaverine are assimilated as sole nitrogen sources. The maximum growth temperature is 35 °C. Growth in the vitamin-free medium is positive. Starch-like substances are not produced. Growth on 50 % (w/w) glucose-yeast extract agar is positive.

Physiologically, *H. daliensis* differs from the closely related species *H. naganishii* and *H. sinensis* (Fig. 24) in its inability to ferment raffinose, and the ability to assimilate maltose, lactose, D-ribose, glycerol, D-mannitol, DL-lactic acid, succinic acid, potassium nitrate, sodium nitrite, and grow in vitamin-free medium (Table S2.29).

###### Typus

**China**, Dali, Yunnan province, obtained from rotten wood, Aug. 2011, *Q.-M. Wang* (**holotype** CGMCC 2.4475^T^ preserved in a metabolically inactive state, ex-type cultures are preserved in culture collections under the numbers CBS 18393).

###### Other culture examined

**China**, obtained from rotten wood, isolated by *Q.M. Wang*, culture CGMCC 2.4476.

***Jamesozyma terricola*** Q.-M. Wang, A.-H. Li, M.-M. Liu, H.-H. Zhu & F.-Y. Bai, ***sp. nov.*** Fungal Names FN 572812. Fig. 46I-K.

###### Etymology

the specific epithet *terricola* refers to the type strain isolated from soil.

###### Culture characteristics

On YM agar, after 1 month at 25 °C, the streak culture is white, butyrous, smooth. The margin is entire (Fig. 46I). In YM broth, after 3 days at 25 °C, cells are ellipsoidal and fusiform, 2.08–4.53 × 2.29–6.09 µm and single, budding is polar (Fig. 46J), after 7 days at 25 °C, a sediment and ring are formed. After 1 month at 25 °C, the sediment and ring are present, but a pellicle does not appear. In Dalmau plate culture on corn meal agar, pseudohyphae are not formed. On V8 agar, the ascus is fusiform and contains two or three ascospores (Fig. 46K). The ascospores are ellipsoidal, 8.62–13.48 × 12.12–17.74 µm.

###### Physiological and biochemical characteristics

Glucose and galactose fermentation are present. Lactose, sucrose, maltose and raffinose fermentation are absent. Glucose, galactose, L-sorbose (weak), maltose, cellobiose (weak), trehalose, melibiose, raffinose, inulin, soluble starch, D-ribose, glycerol, D-mannitol, DL-lactic acid, succinic acid, citric acid (weak) and hexadecane are assimilated as sole carbon sources. Sucrose, lactose, melezitose, D-xylose, L-arabinose, D-arabinose, L-rhamnose, D-glucosamine, N-acetyl-D-glucosamine, methanol, ethanol, erythritol, ribitol, galactitol, D-glucitol, methyl-α-D-glucoside, salicin, inositol and D-glucuronic acid are not assimilated. Ammonium sulfate, potassium nitrate, sodium nitrite, L-lysine, ethylamine and cadaverine are assimilated as sole nitrogen sources. The maximum growth temperature is 30 °C. Growth in the vitamin-free medium is positive. Starch-like substances are not produced. Growth on 50 % (w/w) glucose-yeast extract agar is positive.

Physiologically, *J. terricola* differs from the closely related species *J. xiongxianensis* and *J. jinghongensis* (Fig. 24) in its ability to assimilate L-sorbose, maltose, cellobiose, soluble starch and potassium nitrate (Table S2.30).

###### Typus

**China**, Xinjiang province, obtained from sand soil, Jul. 2008, *Q.-M. Wang* (**holotype** CGMCC 2.3906^T^ preserved in a metabolically inactive state).

###### Other culture examined

**China**, Yunnan province, obtained from soil, Jul. 2004, isolated by *Q.M. Wang*, culture YN0733-3.

***Jamesozyma xiongxianensis*** Q.-M. Wang, A.-H. Li, M.-M. Liu, H.-H. Zhu & F.-Y. Bai, ***sp. nov.*** Fungal Names FN 572813. Fig. 47A-C.

**Fig. 47.**
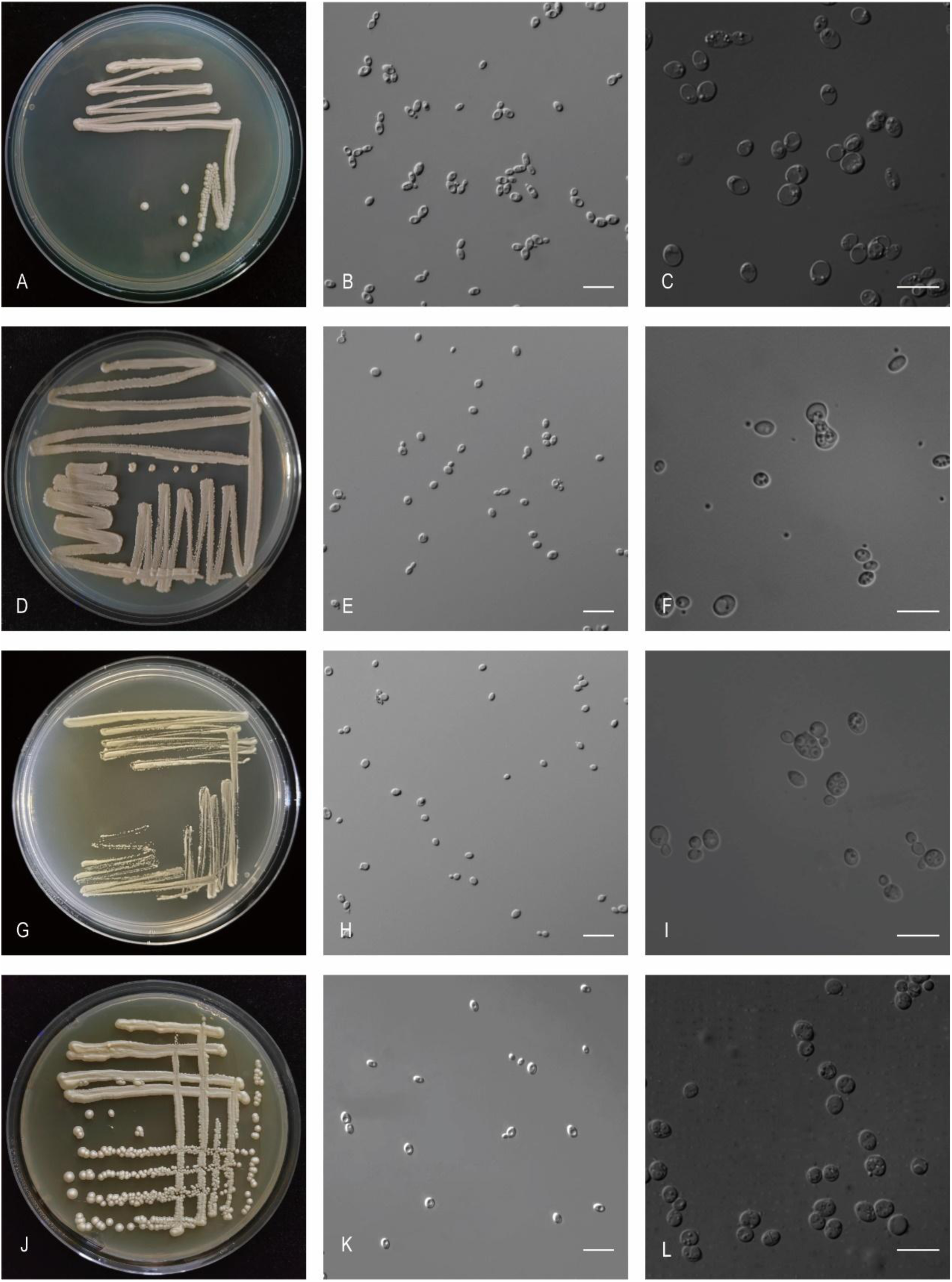
The streak culture grown in YM agar and vegetative cells grown in YM broth for 7 d at 25 °C. A-C. *Jam. xiongxianensis* CGMCC 2.8154^T^. D-F. *Klu. silvicola* CGMCC 2.6963^T^. G-I. *Mau. corticola* CGMCC 2.4466^T^. J-L. *Mau. yunnanensis* CGMCC 2.4425^T^. Scale bars: B, C, E, F, H, I, K, L = 10 μm.

###### Etymology

the specific epithet *xiongxianensis* refers to the geographic origin of the type strain, Xiongxian county, Yunnan province.

###### Culture characteristics

On YM agar, after 1 month at 25 °C, the streak culture is yellowish-white, butyrous, smooth and pale glossy. The margin is entire (Fig. 47A). In YM broth, after 3 days at 25 °C, cells are ellipsoidal and fusiform, 2.01–2.81 × 3.60–5.25 µm and single, budding is polar (Fig. 47B), after 7 days at 25 °C, a sediment is formed. After 1 month at 25 °C, sediment is present, but a ring and pellicle do not appear. In Dalmau plate culture on corn meal agar, pseudohyphae are not formed. On V8 agar, the ascus is fusiform and contains two ascospores (Fig. 47C). The ascospores are ellipsoidal, 3.69–3.77 × 5.27–6.11 µm.

###### Physiological and biochemical characteristics

Glucose, galactose and sucrose fermentation are present. Lactose, maltose and raffinose fermentation are absent. Glucose, galactose, sucrose, trehalose, melibiose (variable), raffinose, inulin (variable), D-ribose (latent), L-rhamnose (variable), D-glucosamine (variable), ethanol (variable), glycerol, ribitol (variable), D-mannitol, D-glucitol (variable), salicin (variable), DL-lactic acid (variable) and citric acid (variable) are assimilated as sole carbon sources. L-sorbose, maltose, cellobiose, lactose, melezitose, soluble starch, D-xylose, L-arabinose, D-arabinose, *N*-acetyl-D-glucosamine, methanol, erythritol, galactitol, methyl-α-D-glucoside, succinic acid, inositol, hexadecane and D-glucuronic acid are not assimilated. Ammonium sulfate (variable), sodium nitrite (variable), L-lysine, ethylamine (variable) and cadaverine (variable) are assimilated as sole nitrogen sources. Potassium nitrate is not assimilated. The maximum growth temperature is 30 °C. Growth in the vitamin-free medium is positive (weak). Starch-like substances are not produced. Growth on 50 % (w/w) glucose-yeast extract agar is negative.

Physiologically, *J. xiongxianensis* differs from the closely related species *J. terricola* and *J. jinghongensis* (Fig. 24) in its ability to ferment sucrose and to assimilate sucrose (Table S2.30).

###### Typus

**China**, Xiongxian County, Zhaotong city, Yunnan Province, obtained from soil, Oct. 2019, *Q.-M. Wang* (**holotype** CGMCC 2.8154^T^ preserved in a metabolically inactive state, ex-type cultures are preserved in culture collections under the numbers YN113-1).

###### Other culture examined

**China**, Xiongxian County, Zhaotong city, Yunnan Province, obtained from soil, Oct. 2019, isolated by *Q.M. Wang*, G.S. Wang, culture YN118D1; **China**, Xiongxian County, Zhaotong city, Yunnan Province, obtained from leaf of *Hovenia acerba*, Oct. 2019, isolated by *Q.M. Wang*, *G.S. Wang*, culture YN103-2.

***Kluyveromyces silvicola*** Q.-M. Wang, A.-H. Li, M.-M. Liu, H.-H. Zhu & F.-Y. Bai, ***sp. nov.*** Fungal Names FN 572814. Fig. 47D-F.

###### Etymology

the specific epithet *silvicola* refers to the type strain isolated from rotten wood.

###### Culture characteristics

On YM agar, after 1 month at 25 °C, the streak culture is pinkish, butyrous, smooth and pale glossy. The margin is entire (Fig. 47D). In YM broth, after 3 days at 25 °C, cells are ellipsoidal and oval, 1.49–3.61 × 2.41–4.19 µm and single, budding is polar (Fig. 47E), after 7 days at 25 °C, a sediment is formed. After 1 month at 25 °C, sediment is present, but a ring and pellicle do not appear. In Dalmau plate culture on corn meal agar, pseudohyphae are not formed. On V8 agar, the ascus is fusiform and contains four ascospores (Fig. 47F). The ascospores are spherical or subspherical, 2.23–3.90 × 2.74–4.14 µm.

###### Physiological and biochemical characteristics

Glucose, galactose (variable), sucrose, maltose (variable) and raffinose (variable) fermentation are present. Lactose fermentation is absent. Glucose, galactose, L-sorbose (latent), sucrose, maltose, cellobiose, trehalose (latent), melibiose (variable), raffinose, melezitose, inulin, soluble starch (variable), D-xylose (latent), L-arabinose (variable), D-ribose (variable), L-rhamnose (variable), N-acetyl-D-glucosamine (variable), ethanol, glycerol (variable), erythritol (variable), ribitol (variable), D-mannitol, D-glucitol, methyl-α-D-glucoside, salicin, DL-lactic acid (variable) and succinic acid (variable) are assimilated as sole carbon sources. Lactose, D-arabinose, D-glucosamine, methanol, galactitol, citric acid, inositol, hexadecane and D-glucuronic acid are not assimilated. Ammonium sulfate, potassium nitrate, sodium nitrite (variable), L-lysine, ethylamine and cadaverine are assimilated as sole nitrogen sources. The maximum growth temperature is 35 °C. Growth in the vitamin-free medium is positive. Starch-like substances are not produced. Growth on 50 % (w/w) glucose-yeast extract agar is negative.

Physiologically, *K. silvicola* differs from the closely related species *K. dobzhanskii* (Fig. 25) in its inability to assimilate citric acid and grow with 50% glucose, and the ability to assimilate inulin and potassium nitrate (Table S2.31).

###### Typus

**China**, Shengfeng Mountains Heilongjiang Province, obtained from bark, Sep. 2017, *Q.-M. Wang* (**holotype** CGMCC 2.6963^T^ preserved in a metabolically inactive state, ex-type cultures are preserved in culture collections under the numbers HSP168-2).

###### Other culture examined

**China**, Diaoluoshan, Hainan province, obtained from rotten wood, Aug. 2011, isolated by *Q.M. Wang*, culture CGMCC 2.4459; **China**, Shengfeng Mountains Heilongjiang Province, obtained from bark, Sep. 2017, isolated by *Q.M. Wang*, culture HSP168-1; **China**, Shengfeng Mountains Heilongjiang Province, obtained from bark, Sep. 2017, isolated by *Q.M. Wang*, culture HSP168-3. **China**, Angren County, Shigatze City, Tibet, obtained from rotten wood, Aug. 2019, isolated by *Q.M. Wang*, culture XZY710-1. **China**, Gongbujiangda, Nyingchi, Tibet, obtained from rotten wood, Aug. 2019, isolated by *Q.M. Wang*, culture XZY769-3.

***Maudiozyma corticola*** Q.-M. Wang, A.-H. Li, M.-M. Liu, H.-H. Zhu & F.-Y. Bai, ***sp. nov.*** Fungal Names FN 572815. Fig. 47G-I.

###### Etymology

the specific epithet *corticola* refers to the type strain isolated from bark.

###### Culture characteristics

On YM agar, after 1 month at 25 °C, the streak culture is white, butyrous and smooth. The margin is entire (Fig. 47G). In YM broth, after 3 days at 25 °C, cells are subrotund and oval, 1.19–2.46 × 1.31–3.78 µm and single, budding is polar (Fig. 47H), after 7 days at 25 °C, a sediment and ring are formed. After 1 month at 25 °C, sediment and ring are present, but a pellicle does not appear. In Dalmau plate culture on corn meal agar, pseudohyphae are not formed. On V8 agar, the ascus is oval and contains four ascospores (Fig. 47I). The ascospores are spherical or subspherical, 2.63–4.57 × 2.15–5.03 µm.

###### Physiological and biochemical characteristics

Glucose, galactose and sucrose fermentation are present. Lactose, maltose and raffinose fermentation are absent. Glucose, galactose, L-sorbose, trehalose, raffinose, inulin (latent), soluble starch, ethanol, galactitol (latent, weak), D-mannitol and succinic acid are assimilated as sole carbon sources. Sucrose, maltose, cellobiose, lactose, melibiose, melezitose, D-xylose, L-arabinose, D-arabinose, D-ribose, L-rhamnose, D-glucosamine, N-acetyl-D-glucosamine, methanol, glycerol, erythritol, ribitol, D-glucitol, methyl-α-D-glucoside, salicin, DL-lactic acid, citric acid, inositol, hexadecane and D-glucuronic acid are not assimilated. Ammonium sulfate, potassium nitrate, sodium nitrite, L-lysine, ethylamine and cadaverine are assimilated as sole nitrogen sources. The maximum growth temperature is 35 °C. Growth in the vitamin-free medium is negative. Starch-like substances are not produced. Growth on 50 % (w/w) glucose-yeast extract agar is positive.

Physiologically, *M. corticola* differs from the closely related species *M. yunnanensis*, *M. australis* and *M. exigua* (Fig. 24) in its ability to assimilate soluble starch and grow with 50% glucose (Table S2.32).

###### Typus

**China**, obtained from bark, Aug. 2011, *Q.-M. Wang* (**holotype** CGMCC 2.4466^T^ preserved in a metabolically inactive state).

***Maudiozyma yunnanensis*** Q.-M. Wang, A.-H. Li, M.-M. Liu, H.-H. Zhu & F.-Y. Bai, ***sp. nov.*** Fungal Names FN 572816. Fig. 47J-L.

###### Etymology

the specific epithet *yunnanensis* refers to the geographic origin of the type strain, Yunnan province.

###### Culture characteristics

On YM agar, after 1 month at 25 °C, the streak culture is pink-white, butyrous and smooth. The margin is entire (Fig. 47J). In YM broth, after 3 days at 25 °C, cells are oval, 1.96–3.89 × 2.65–5.12 µm and single, budding is polar (Fig. 47K), after 7 days at 25 °C, a sediment and ring are formed. After 1 month at 25 °C, sediment and ring are present, but a pellicle does not appear. In Dalmau plate culture on corn meal agar, pseudohyphae are not formed. On V8 agar, the ascus is subglobose and contains two or three ascospores (Fig. 47L). The ascospores are ellipsoidal, 2.38–3.76 × 3.69–4.42 µm.

###### Physiological and biochemical characteristics

Glucose, galactose and sucrose fermentation are present. Lactose, maltose and raffinose fermentation are absent. Glucose, galactose, L-sorbose, sucrose, maltose, cellobiose (variable), trehalose (variable), melibiose, raffinose (variable), melezitose (weak), inulin (weak), D-xylose (variable), L-arabinose (variable), D-arabinose (variable), D-glucosamine, N-acetyl-D-glucosamine (variable), ethanol, erythritol (variable), ribitol (weak), galactitol (weak), D-mannitol (variable), D-glucitol, methyl-α-D-glucoside (variable), salicin (variable), DL-lactic acid (latent), succinic acid (latent) and citric acid (variable) are assimilated as sole carbon sources. Lactose, soluble starch, D-ribose, L-rhamnose, methanol, glycerol, inositol, hexadecane and D-glucuronic acid are not assimilated. potassium nitrate (variable), sodium nitrite (weak), L-lysine, ethylamine (variable) and cadaverine (latent, weak) are assimilated as sole nitrogen sources. Ammonium sulfate is not assimilated. The maximum growth temperature is 32 °C. Growth in the vitamin-free medium is positive. Starch-like substances are not produced. Growth on 50 % (w/w) glucose-yeast extract agar is negative.

Physiologically, *M. yunnanensis* differs from the closely related species *M. corticola*, *M. australis* and *M. exigua* (Fig. 24) in its ability to assimilate maltose, melibiose, melezitose, ribitol, D-glucitol, citric acid, and grow in vitamin-free medium (Table S2.32).

###### Typus

**China**, obtained from the flower of *Benincasa hispida*, *Q.-M. Wang* (**holotype** CGMCC 2.4425^T^ preserved in a metabolically inactive state).

###### Other culture examined

**China**, Hainan Province, obtained from rotten wood, isolated by *Q.M. Wang*, culture CGMCC 2.4308.

***Naumovozyma terricola*** Q.-M. Wang, A.-H. Li, M.-M. Liu, H.-H. Zhu & F.-Y. Bai, ***sp. nov.*** Fungal Names FN 572817. Fig. 48A, B.

**Fig. 48.**
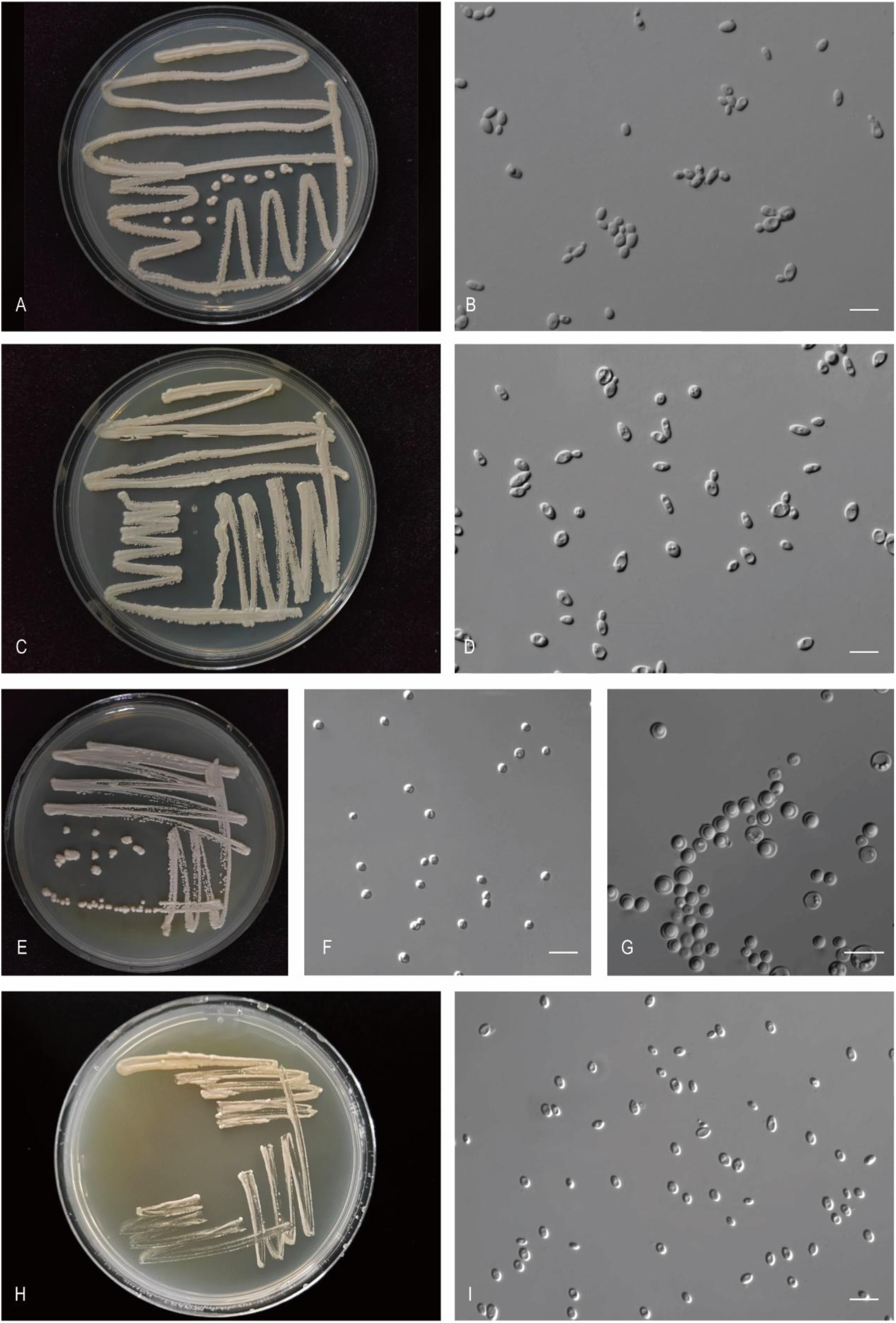
The streak culture grown in YM agar and vegetative cells grown in YM broth for 7 d at 25 °C. A, B. *Nau. terricola* CGMCC 2.6960^T^. C, D. *Tor. xizangensis* CGMCC 2.6961^T^. E-G. *Zyg. silvicola* CGMCC 2.4531^T^. H, I. *Tap. follicola* CGMCC 2.2537^T^. Scale bars: B, D, F, G, I = 10 μm.

###### Etymology

the specific epithet *terricola* refers to the type strain isolated from soil.

###### Culture characteristics

On YM agar, after 1 month at 25 °C, the streak culture is white, butyrous, rough and dimmed. The margin is fringed (Fig. 48A). In YM broth, after 3 days at 25 °C, cells are ellipsoidal and fusiform, 1.94–5.03 × 4.58–6.14 µm and single, budding is polar (Fig. 48B), after 7 days at 25 °C, a sediment, pellicle and ring are formed. After 1 month at 25 °C, sediment, pellicle and ring are present. In Dalmau plate culture on corn meal agar, pseudohyphae are not formed. Asci and ascospores are not observed on V8 and Acetate agar.

###### Physiological and biochemical characteristics

Glucose and galactose fermentation are present. Lactose, sucrose, maltose and raffinose fermentation are absent. Glucose, galactose, maltose (variable), raffinose (weak), inulin (weak), soluble starch (variable), L-rhamnose (variable), D-glucosamine (variable), N-acetyl-D-glucosamine (variable), methanol (variable), ethanol (latent), glycerol (latent), erythritol (variable), ribitol (variable), D-mannitol, methyl-α-D-glucoside (weak), DL-lactic acid (variable), succinic acid (latent, weak) and citric acid (latent, weak) are assimilated as sole carbon sources. L-sorbose, sucrose, cellobiose, trehalose, lactose, melibiose, melezitose, D-xylose, L-arabinose, D-arabinose, D-ribose, galactitol, D-glucitol, salicin, inositol, hexadecane and D-glucuronic acid are not assimilated. Ammonium sulfate, potassium nitrate (variable), sodium nitrite, L-lysine, ethylamine and cadaverine are assimilated as sole nitrogen sources. The maximum growth temperature is 37 °C. Growth in the vitamin-free medium is positive (variable). Starch-like substances are not produced. Growth on 50 % (w/w) glucose-yeast extract agar is negative.

Physiologically, *N. terricola* differs from the closely related species *N. castellii*, *N. baii* and *N. dairenensis* (Fig. 26) in its ability to assimilate raffinose, inulin, glycerol, D-mannitol, methyl-α-D-glucoside, succinic acid, citric acid, sodium nitrite, L-lysine, ethylamine and cadaverine (Table S2.33).

###### Typus

**China**, Xiongxian County, Zhaotong city, Yunnan Province, obtained from soil, Oct. 2019, *Q.-M. Wang* (**holotype** CGMCC 2.6960^T^ preserved in a metabolically inactive state, ex-type cultures are preserved in culture collections under the numbers YN125-1).

###### Other culture examined

**China**, obtained from soil, isolated by *Q.M. Wang*, culture CGMCC 2.4461.

***Torulaspora xizengensis*** Q.-M. Wang, A.-H. Li, M.-M. Liu, H.-H. Zhu & F.-Y. Bai, ***sp. nov.*** Fungal Names FN 572818. Fig. 48C, D.

###### Etymology

the specific epithet *xizengensis* refers to the geographic origin of the type strain, Xizang.

###### Culture characteristics

On YM agar, after 1 month at 25 °C, the streak culture is white, butyrous, smooth and pale glossy. The margin is entire (Fig. 48C). In YM broth, after 3 days at 25 °C, cells are ellipsoidal and fusiform, 2.09–5.04 × 3.89–7.58 µm and single, budding is polar (Fig. 48D), after 7 days at 25 °C, a sediment is formed. After 1 month at 25 °C, sediment is present, but a ring and pellicle do not appear. In Dalmau plate culture on corn meal agar, pseudohyphae are not formed. Asci and ascospores are not observed on V8 and Acetate agar.

###### Physiological and biochemical characteristics

Glucose, galactose (variable), sucrose and maltose (variable) fermentation are present. Lactose and raffinose fermentation are absent. Glucose, galactose (variable), L-sorbose (variable), sucrose (variable), maltose (variable), cellobiose (variable), trehalose, lactose (variable), raffinose (variable), melezitose, inulin, soluble starch (variable), D-xylose (variable), D-arabinose (variable), D-glucosamine (variable), N-acetyl-D-glucosamine (variable), methanol (variable), ethanol (variable), glycerol (variable), erythritol (variable), ribitol (variable), D-mannitol, D-glucitol (latent), methyl-α-D-glucoside, salicin (variable), DL-lactic acid (variable), succinic acid (variable) and citric acid (variable) are assimilated as sole carbon sources. Melibiose, L-arabinose, D-ribose, L-rhamnose, galactitol, inositol, hexadecane and D-glucuronic acid are not assimilated. Ammonium sulfate, potassium nitrate, sodium nitrite (variable), L-lysine, ethylamine (variable) and cadaverine (variable) are assimilated as sole nitrogen sources. The maximum growth temperature is 35 °C. Growth in the vitamin-free medium is positive. Starch-like substances are not produced. Growth on 50 % (w/w) glucose-yeast extract agar is positive.

Physiologically, *T. xizengensis* differs from the closely related species *T. delbrueckii* (Fig. 27) in its ability to assimilate potassium nitrate (Table S2.34).

###### Typus

**China**, Angren County, Shigatze City, Tibet, obtained from rotten wood, Aug. 2019, *Q.-M. Wang* (**holotype** CGMCC 2.6961^T^ preserved in a metabolically inactive state, ex-type cultures are preserved in culture collections under the numbers XZY711-3).

###### Other culture examined

**China**, Gongbujiangda, Nyingchi, Tibet, obtained from rotten wood, Aug. 2019, isolated by *Q.M. Wang*, culture XZY765-1 (= CGMCC 2.6962); **China**, Baiba County, Linzhi, Tibet, obtained from rotten wood, Aug. 2019, isolated by *Q.M. Wang*, culture XZY735-1; **China**, Basongcuo Lake, Nyingchi, Tibet, obtained from leaf of *Viola tricolor*, Aug. 2019, isolated by *Q.M. Wang*, culture XZY238F1; **China**, Basongcuo Lake, Nyingchi, Tibet, obtained from leaf of *Viola tricolor*, Aug. 2019, isolated by *Q.M. Wang*, culture XZY238F2; **China**, Angren County, Shigatze City, Tibet, obtained from rotten wood, Aug. 2019, isolated by *Q.M. Wang*, culture XZY719-1.

***Zygotorulaspora silvicola*** Q.-M. Wang, A.-H. Li, M.-M. Liu, H.-H. Zhu & F.-Y. Bai, ***sp. nov.*** Fungal Names FN 572819. Fig. 48E-G.

###### Etymology

the specific epithet silvicola refers to the type strain isolated from rotten wood.

###### Culture characteristics

On YM agar, after 1 month at 25 °C, the streak culture is white, butyrous, smooth and pale glossy. The margin is entire (Fig. 48E). In YM broth, after 3 days at 25 °C, cells are oval and globose, 2.42–3.77 × 2.79–4.67 µm and single, budding is polar (Fig. 48F), after 7 days at 25 °C, a sediment is formed. After 1 month at 25 °C, sediment is present, but a ring and pellicle do not appear. In Dalmau plate culture on corn meal agar, pseudohyphae are not formed. On V8 agar, the ascus is oval and contains one to three ascospores (Fig. 48G). The ascospores are spherical or subspherical, 1.43–2.91 x 1.33–2.94 µm.

###### Physiological and biochemical characteristics

Glucose, galactose, sucrose, maltose and raffinose fermentation are present. Lactose fermentation is absent. Glucose, galactose, L-sorbose, sucrose, maltose, trehalose, melibiose, raffinose, melezitose, inulin, ethanol, glycerol, ribitol, D-mannitol, D-glucitol, methyl-α-D-glucoside, DL-lactic acid, succinic acid and citric acid are assimilated as sole carbon sources. Cellobiose, lactose, soluble starch, D-xylose, L-arabinose, D-arabinose, D-ribose, L-rhamnose, D-glucosamine, N-acetyl-D-glucosamine, methanol, erythritol, galactitol, salicin, inositol, hexadecane and D-glucuronic acid are not assimilated. Ammonium sulfate, potassium nitrate, sodium nitrite, L-lysine, ethylamine and cadaverine are assimilated as sole nitrogen sources. The maximum growth temperature is 42 °C. Growth in the vitamin-free medium is positive. Starch-like substances are not produced. Growth on 50 % (w/w) glucose-yeast extract agar is negative.

Physiologically, *Z. silvicola* differs from the closely related species *Z. cariocana* (Fig. 28) in its ability to assimilate glycerol, ribitol, citric acid, potassium nitrate and sodium nitrite (Table S2.35).

###### Typus

**China**, Bawangling, Hainan province, obtained from rotten wood, Mar. 2012, *Q.-M. Wang* (**holotype** CGMCC 2.4531^T^ preserved in a metabolically inactive state).

###### Other culture examined

**China**, Bawangling, Hainan province, obtained from rotten wood, Mar. 2012, isolated by *Q.M. Wang*, culture BW31-7.

New taxa in the *Taphrina* (*Taphrinaceae*, *Taphrinales*, *Taphrinomycetes*, *Taphrinomycotina*) *Taphrina follicola* Q.-M. Wang, A.-H. Li, M.-M. Liu, H.-H. Zhu & F.-Y. Bai, *sp. nov.* Fungal Names FN 572820. Fig. 48H, I.

###### Etymology

the specific epithet *follicola* refers to the type strain isolated from a leaf.

###### Culture characteristics

On YM agar, after 1 month at 25 °C, the streak culture is pale-pink, butyrous, smooth and glossy. The margin is entire (Fig. 48H). In YM broth, after 3 days at 25 °C, cells are ellipsoidal, 2.30–3.36 × 3.41–5.15 µm and single, budding is polar (Fig. 48I), after 7 days at 25 °C, a sediment is formed. After 1 month at 25 °C, sediment and pellicle are present, but a ring does not appear. In Dalmau plate culture on corn meal agar, pseudohyphae are not formed. Asci and ascospores are not observed on V8 and Acetate agar.

###### Physiological and biochemical characteristics

Glucose, lactose, galactose, sucrose, maltose and raffinose fermentation are absent. Glucose, galactose, L-sorbose, sucrose, maltose, cellobiose, trehalose, raffinose, melezitose, inulin, soluble starch, D-xylose, L-arabinose, D-arabinose, N-acetyl-D-glucosamine, ethanol, glycerol, ribitol (latent, weak), D-mannitol and D-glucitol are assimilated as sole carbon sources. Lactose, melibiose, D-ribose, L-rhamnose, D-glucosamine, methanol, erythritol, galactitol, methyl-α-D-glucoside, salicin, DL-lactic acid, succinic acid, citric acid, inositol, hexadecane and D-glucuronic acid are not assimilated. Ammonium sulfate, potassium nitrate, sodium nitrite and cadaverine are assimilated as sole nitrogen sources. L-lysine and ethylamine are not assimilated. The maximum growth temperature is 32 °C. Growth in the vitamin-free medium is positive. Starch-like substances are produced. Growth on 50 % (w/w) glucose-yeast extract agar is negative.

Physiologically, *T. follicola* does not differ from the closely related species *T. betulina*, *T. americana*, *T. tormentillae*, *T. antarctica*, *T. populina*, *T. populi-salicis* and *T. johansonii* (Fig. 29) in its ability to utilize carbon sources and nitrogen sources (Table S2.36).

###### Typus

**China**, Shaanxi province, obtained from leaf, Dec. 2003, *Q.-M. Wang* (**holotype** CGMCC 2.2537^T^ preserved in a metabolically inactive state, ex-type cultures are preserved in culture collections under the numbers CB91).

***Taphrina planticola*** Q.-M. Wang, A.-H. Li, M.-M. Liu, H.-H. Zhu & F.-Y. Bai, ***sp. nov.*** Fungal Names FN 572821. Fig. 49A, B.

**Fig. 49.**
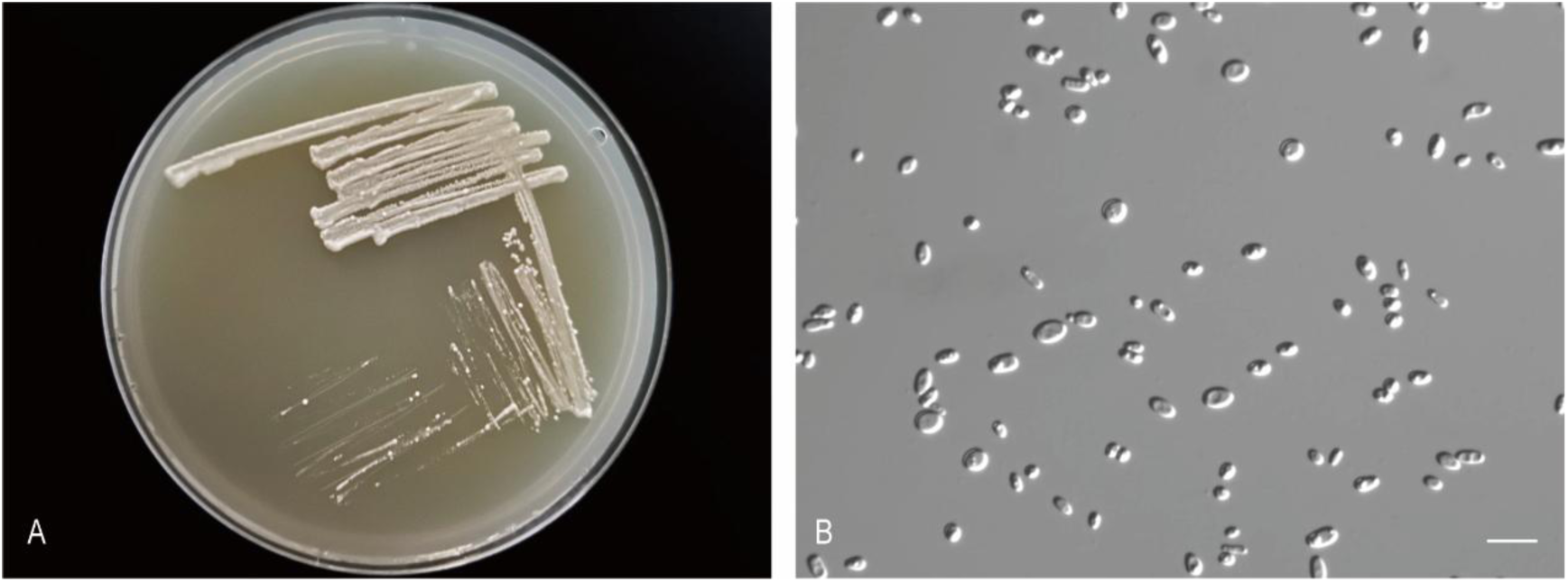
The streak culture grown in YM agar and vegetative cells grown in YM broth for 7 d at 25 °C. A, B. *Tap. planticola* CGMCC 2.5682^T^. Scale bars: B = 10 μm.

###### Etymology

the specific epithet *planticola* refers to the type strain isolated from a plant.

###### Culture characteristics

On YM agar, after 1 month at 25 °C, the streak culture is white, butyrous, smooth. The margin is eroded (Fig. 49A). In YM broth, after 3 days at 25 °C, cells are ellipsoidal, oval and fusiform, 1.50–2.82 × 3.18–4.67 µm and single, budding is polar (Fig. 49B), after 7 days at 25 °C, a sediment is formed. After 1 month at 25 °C, sediment and pellicle are present, but a ring does not appear. In Dalmau plate culture on corn meal agar, pseudohyphae are not formed. Asci and ascospores are not observed on V8 and Acetate agar.

###### Physiological and biochemical characteristics

Glucose, sucrose (weak) and maltose (latent) fermentation are present. Lactose, galactose and raffinose fermentation are absent. Glucose, galactose, L-sorbose (latent), sucrose, maltose, cellobiose (latent), trehalose, raffinose (latent), melezitose (latent, weak), inulin, soluble starch (weak), D-xylose (latent), L-arabinose (weak), D-arabinose (latent, weak), D-ribose (weak), D-glucosamine, N-acetyl-D-glucosamine, ethanol (weak), glycerol (latent, weak), erythritol (latent, weak), ribitol (latent, weak), galactitol (latent), D-mannitol, D-glucitol (latent), methyl-α-D-glucoside (latent), salicin (latent), succinic acid and citric acid (weak) are assimilated as sole carbon sources. Lactose, melibiose, L-rhamnose, methanol, DL-lactic acid, inositol, hexadecane and D-glucuronic acid are not assimilated. Ammonium sulfate, potassium nitrate, sodium nitrite (latent), L-lysine, ethylamine and cadaverine are assimilated as sole nitrogen sources. The maximum growth temperature is 30 °C. Growth in the vitamin-free medium is positive. Starch-like substances are not produced. Growth on 50 % (w/w) glucose-yeast extract agar is negative.

Physiologically, *T. planticola* differs from the closely related species *T. kurtzmanii*, *T. caerulescens* and *T. arrabidae* (Fig. 29) in its ability to ferment glucose, sucrose and maltose, and the ability to assimilate galactose, maltose, raffinose, inulin, L-arabinose, D-arabinose, D-ribose, erythritol, galactitol, methyl-α-D-glucoside and ethylamine (Table S2.36).

###### Typus

**China**, obtained from leaf, Apr. 2016, *Q.-M. Wang* (**holotype** CGMCC 2.5682^T^ preserved in a metabolically inactive state, ex-type cultures are preserved in culture collections under the numbers 133131).

## Resource Availability

Tables S1–S2 and Figures S1–S2 are deposited at Figshare repository: https://doi.org/10.6084/m9.figshare.29278250.

## CONTRIBUTIONS

Q.-M.W. conceived and designed the project. M.-M.L., Y.-L.J., H.-H.Z. and Y.-T.G. performed phenotypic characterization. Q.-M.W., M.-M.L. and A.-H.L. analyzed the molecular data. X.-M.Z. performed the phylogenomic and genomic metrics analyses. J.-H.L., X.-H.Z., J.-C.W. and Y.-J.S. edited the rDNA phylogenetic trees and checked the physiological data. Q.-M.W., A.-H.L., D.-P. and G.-S.W. performed sampling and yeast isolation. Q.-M.W., M.-M.L. and A.-H.L. wrote the paper. Q.-M.W., M.-M.L., A.-H.L. and F.-Y.B. revised and edited the paper.

## ACKNOWLEDGEMENTS

This study was supported by grants No. 32370015 and No. 31961133020 from the National Natural Science Foundation of China (NSFC), No. XZ202501ZY009 from Science and Technology Project of Xizang Autonomous Region, China, No. 2021FY100900 and No. 2023FY101300 from the Science & Technology Fundamental Resources Investigation Program of the Ministry of Science and Technology of China, No. 521000981388 from Advanced Talents Incubation Program of Hebei University. The authors are solely responsible for the content of this work.

## DECLARATIONS

The authors declare that there are no conflicts of interest.

## Supporting Information

**Table S1.** List of yeasts and sequence numbers retrieved from GenBank (provided as a separate Excel file).

**Table S2.** Physiological and biochemical characteristics of new species and their closest relatives (provided as a separate Excel file).

**Fig. S1.** The proportion of strains isolated from different substrates.

**Fig. S2.** Phylogeny of new taxa (in bold) in the *Suhomyces* inferred from the sequences of ITS region (including 5.8S rDNA) by maximum likelihood analysis and over 50 % from 1 000 bootstrap replicates is shown. Bar = 0.2 substitutions per nucleotide position.

## REFERENCES

Altschul SF, Madden TL, Schäffer AA, et al. (1997). Gapped BLAST and PSI-BLAST: a new generation of protein database search programs. Nucleic Acids Research 25: 3389–3402.

Bankevich A, Nurk S, Antipov D, et al. (2012). SPAdes: A new genome as sembly algorithm and its applications to single-cell sequencing. Journal of Computational Biology 19: 455–477.

Boekhout T, Aime MC, Begerow D, et al. (2021). The evolving species concepts used for yeasts: from phenotypes and genomes to speciation networks. Fungal Diversity 109: 27–55.

Boekhout T, Amend AS, El Baidouri F, et al. (2022). Trends in yeast diversity discovery. Fungal Diversity 114: 491–537.

Chen S, Zhou Y, Chen Y, et al. 2018. Fastp: An ultra-fast all-in-one FASTQ preprocessor. Bioinformatics 34: i884–i890.

Cortimiglia C, Alonso-Del-Real J, Daza MV, et al. (2024). Evaluating the genome-based average nucleotide identity calculation for identification of twelve yeast species. Journal of Fungi 10: 646.

Felsenstein J (1985) Confidence limits on phylogenies: An approach using the bootstrap. Evolution 39: 783–791.

Fell JW, Boekhout T, Fonseca A, et al. (2000). Biodiversity and systematics of basidiomycetous yeasts as determined by large-subunit rDNA D1/D2 domains sequence analysis. International Journal of Systematic and Evolutionary Microbiology 50: 1351–1371.

Groenewald M, Hittinger CT, Bensch K, et al. (2023). A genome-informed higher rank classification of the biotechnologically important fungal subphylum *Saccharomycotina*. Studies in Mycology 105: 1–22.

Goris J, Konstantinidis KT, Klappenbach JA, et al. 2007. DNA-DNA hybridization values and their relationship to whole-genome sequence similarities. International Journal of Systematic and Evolutionary Microbiology 57: 81–91.

Jiang YL, Bao WJ, Liu F, et al. (2024). Proposal of one new family, seven new genera and seventy new basidiomycetous yeast species mostly isolated from Tibet and Yunnan provinces, China. Studies in Mycolog 109: 57–153.

Kurtzman CP, Fell JW, Boekhout T (2011a). In: *The Yeasts*, *A Taxonomic Study* (Kurtzman CP, Fell JW, Boekhout T, eds), 5th edn. Elsevier, The Netherlands.

Kurtzman CP, Fell JW, Boekhout T, et al. (2011b). Methods for isolation, phenotypic characterization and maintenance of yeasts. In: The Yeasts, A Taxonomic Study, 5th edn. (Kurtzman CP, Fell JW, Boekhout T, eds). Elsevier, Netherland: 87–110.

Kumar S, Stecher G, Tamura K (2016). MEGA7: Molecular evolutionary genetics analysis version 7.0 for bigger datasets. Molecular Biology and Evolution 33:1870–1874.

Kurtzman CP (2011). Discussion of teleomorphic and anamorphic ascomycetous yeasts and yeast-like taxa. In: The Yeasts, A Taxonomic Study (Kurtzman CP, Fell JW, Boekhout T, eds), 5th edn. Elsevier, The Netherland: 293–307.

Kurtzman CP, Robnett CJ. (1998). Identification and phylogeny of ascomycetous yeasts from analysis of nuclear large subunit (26S) ribosomal DNA partial sequences. Antonie Van Leeuwenhoek 73: 331–371.

Lachance MA, Boekhout T, Scorzetti G, et al. (2011). *Candida* Berkhout. In: The Yeasts, A Taxonomic Study (Kurtzman CP, Fell JW, Boekhout T, eds), 5th edn. Elsevier, The Netherlands: 987–1278.

Lachance MA, Lee DK, Hsiang T. (2020). Delineating yeast species with genome average nucleotide identity: a calibration of ANI with haplontic, heterothallic *Metschnikowia* species. Antonie Van Leeuwenhoek 113: 2097–2106.

Li AH, Yuan FX, Groenewald M, et al. (2020). Diversity and phylogeny of basidiomycetous yeasts from plant leaves and soil: proposal of two new orders, three new families, eight new genera and one hundred and seven new species. Studies in Mycology 96: 17–140.

Li YY, Wang MM, Groenewald M, et al. (2022). Proposal of two new combinations, twenty new species, four new genera, one new family, and one new order for the anamorphic basidiomycetous yeast species in *Ustilaginomycotina*. Frontiers in Microbiology 12: 777338.

Liu MM, Zhao XM, Bai J, et al. (2025). Taxogenomic reclassification of *Candida* and related genera in Saccharomycotina. Fungal Diversity (submitted).

Liu F, Hu ZD, Yurkov A, et al. (2024a). *Saccharomycetaceae*: delineation of fungal genera based on phylogenomic analyses, genomic relatedness indices and genomics-based synapomorphies. Persoonia 52: 1–21.

Liu F, Hu ZD, Zhao XM, et al. (2024b). Phylogenomic analysis of the *Candida auris*-*Candida haemuli* clade and related taxa in the *Metschnikowiaceae*, and proposal of thirteen new genera, fifty-five new combinations and nine new species. Persoonia 52: 22–43.

Libkind D, Čadež N, Opulente DA, et al. (2020). Towards yeast taxogenomics: lessons from novel species descriptions based on complete genome sequences. FEMS Yeast Research 20: foaa042.

Nakase T, Takashima M (1993). A simple procedure for the high frequency isolation of new taxa of ballistosporous yeasts living on the surfaces of plants. RIKEN Review 3: 33–34.

Opulente DA, LaBella AL, Harrison MC, et al. (2024). Genomic factors shape carbon and nitrogen metabolic niche breadth across Saccharomycotina yeasts. Science 384: eadj4503.

Scorzetti G, Fell JW, Fonseca A, et al. (2002). Systematics of basidiomycetous yeasts: a comparison of large subunit D1/D2 and internal transcribed spacer rDNA regions. FEMS Yeast Research 2: 495– 517.

Shen XX, Opulente DA, Kominek J, et al. (2018). Tempo and Mode of Genome Evolution in the Budding Yeast Subphylum. Cell 175: 1533–1545.e20.

Sugita T, Nakase T (1999). Non-universal usage of the leucine CUG codon and the molecular phylogeny of the genus Candida. Systematic and Applied Microbiology 22: 79–86.

Ter-Hovhannisyan V, Lomsadze A, Chernoff YO, et al. 2008. Gene prediction in novel fungal genomes using an ab initio algorithm with unsupervised training. Genome Research 18: 1979–1990.

Troiano E, Larini I, Binati RL, et al. (2023). Finding a correct species assignment for a *Metschnikowia* strain: insights from the genome sequencing of strain DBT012. FEMS Yeast Research 23: foad024.

Vu D, Groenewald M, Szöke S, et al. (2016). DNA barcoding analysis of more than 9000 yeast isolates contributes to quantitative thresholds for yeast species and genera delimitation. Studies in Mycology 85: 91–105.

Wang QM, Bai FY (2008). Molecular phylogeny of basidiomycetous yeasts in the *Cryptococcus luteolus* lineage (*Tremellales*) based on nuclear rDNA and mitochondrial cytochrome b gene sequence analyses: proposal of *Derxomyces* gen. nov. and *Hannaella* gen. nov., and eight novel *Derxomyces* species. FEMS Yeast Research 8: 799–814.

Wang QM, Theelen B, Groenewald M, et al. (2014). *Moniliellomycetes* and *Malasseziomycetes*, two new classes in *Ustilaginomycotina*. Persoonia 33: 41–47.

