## Supplementary figures and images for "Proposal of two new genera and seventy-seven new species of ascomycetous yeasts isolated from China"

### Fig. S1

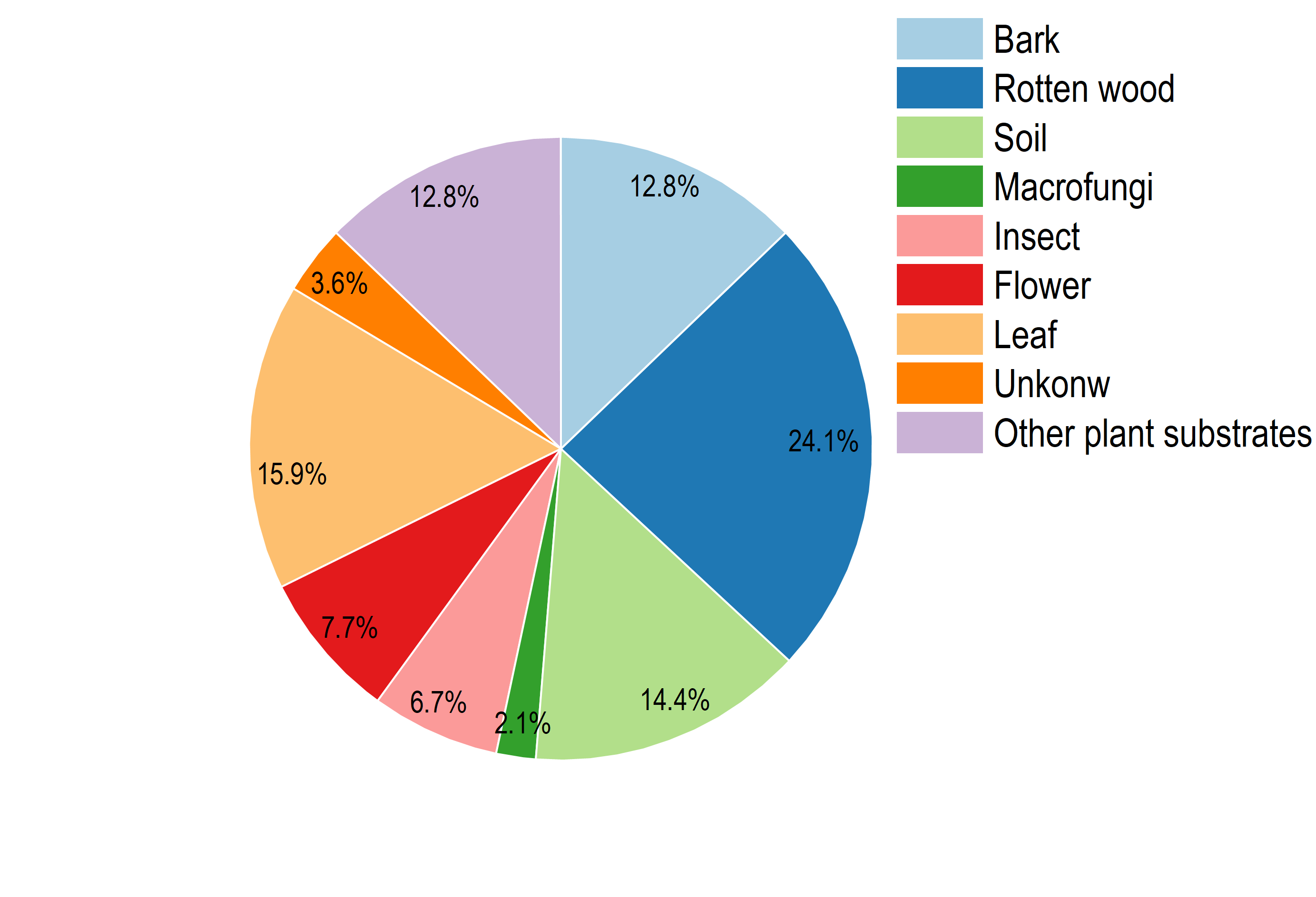
