## Supplementary material for "Proposal of two new genera and seventy-seven new species of ascomycetous yeasts isolated from China": Fig. S2

### Slide 1
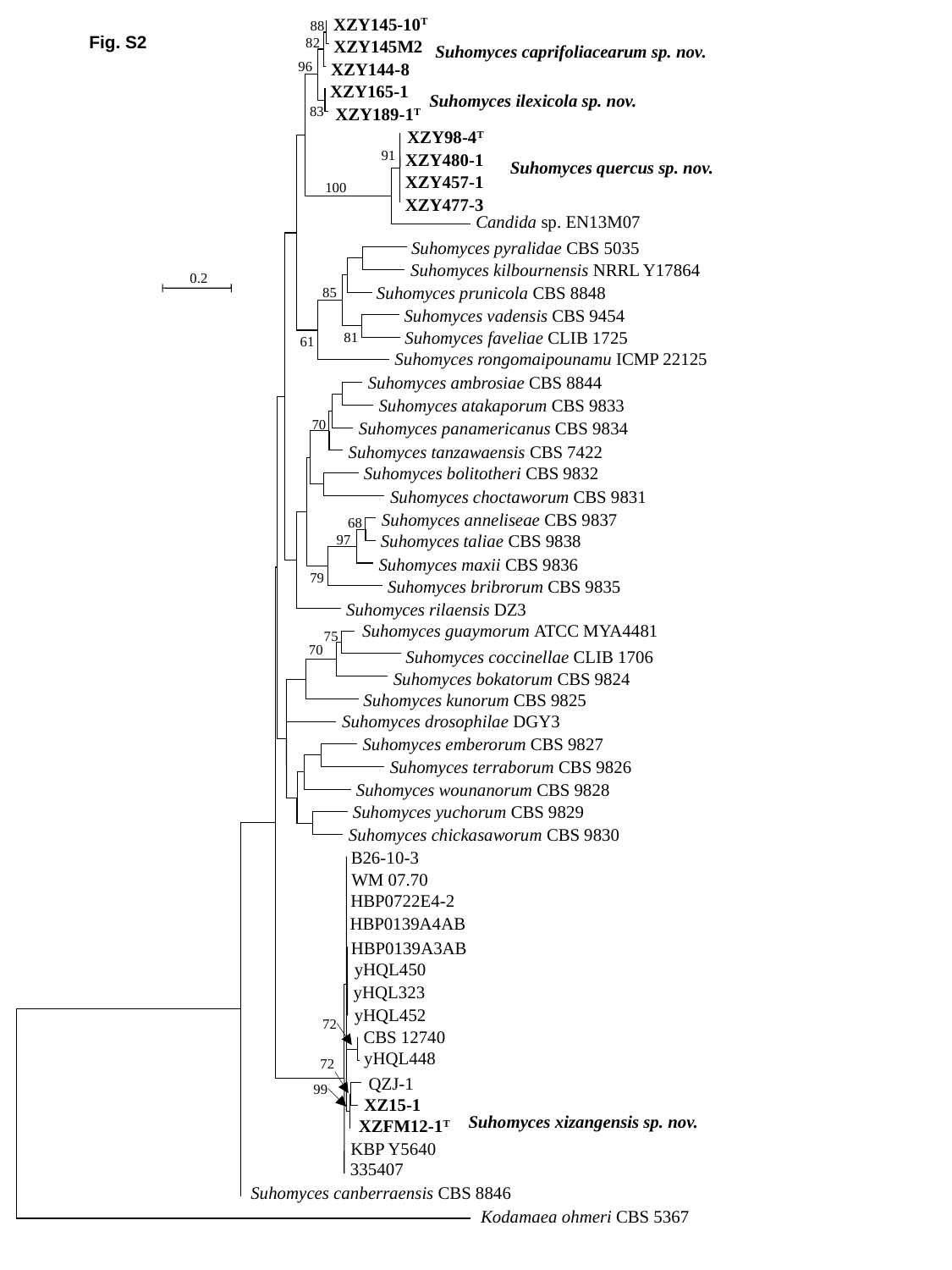

XZY145-10^T^
88
82
Suhomyces caprifoliacearum sp. nov.
 XZY145M2
96
 XZY144-8
 XZY165-1
Suhomyces ilexicola sp. nov.
83
 XZY189-1^T^
 XZY98-4^T^
91
 XZY480-1
Suhomyces quercus sp. nov.
 XZY457-1
100
 XZY477-3
 Candida sp. EN13M07
 Suhomyces pyralidae CBS 5035
 Suhomyces kilbournensis NRRL Y17864
0.2
 Suhomyces prunicola CBS 8848
85
 Suhomyces vadensis CBS 9454
 Suhomyces faveliae CLIB 1725
81
61
 Suhomyces rongomaipounamu ICMP 22125
 Suhomyces ambrosiae CBS 8844
 Suhomyces atakaporum CBS 9833
70
 Suhomyces panamericanus CBS 9834
 Suhomyces tanzawaensis CBS 7422
 Suhomyces bolitotheri CBS 9832
 Suhomyces choctaworum CBS 9831
 Suhomyces anneliseae CBS 9837
68
 Suhomyces taliae CBS 9838
97
 Suhomyces maxii CBS 9836
79
 Suhomyces bribrorum CBS 9835
 Suhomyces rilaensis DZ3
 Suhomyces guaymorum ATCC MYA4481
75
70
 Suhomyces coccinellae CLIB 1706
 Suhomyces bokatorum CBS 9824
 Suhomyces kunorum CBS 9825
 Suhomyces drosophilae DGY3
 Suhomyces emberorum CBS 9827
 Suhomyces terraborum CBS 9826
 Suhomyces wounanorum CBS 9828
 Suhomyces yuchorum CBS 9829
 Suhomyces chickasaworum CBS 9830
B26-10-3
WM 07.70
HBP0722E4-2
HBP0139A4AB
HBP0139A3AB
yHQL450
yHQL323
yHQL452
72
CBS 12740
yHQL448
72
QZJ-1
99
 XZ15-1
Suhomyces xizangensis sp. nov.
XZFM12-1^T^
 KBP Y5640
 335407
 Suhomyces canberraensis CBS 8846
 Kodamaea ohmeri CBS 5367
Fig. S2
